# Characterization of prevalence and health consequences of uniparental disomy in four million individuals from the general population

**DOI:** 10.1101/540955

**Authors:** Priyanka Nakka, Samuel Pattillo Smith, Anne H. O’Donnell-Luria, Kimberly F. McManus, 23andMe Research Team, Joanna L. Mountain, Sohini Ramachandran, J. Fah Sathirapongsasuti

## Abstract

Meiotic nondisjunction and resulting aneuploidy can lead to severe health consequences in humans. Aneuploidy rescue can restore euploidy but may result in uniparental disomy (UPD), the inheritance of both homologs of a chromosome from one parent with no representative copy from the other. Current understanding of UPD is limited to ~3,300 cases for which UPD was associated with clinical presentation due to imprinting disorders or recessive diseases. Thus, the prevalence of UPD and its phenotypic consequences in the general population are unknown. We searched for instances of UPD in over four million consented research participants from the personal genetics company 23andMe, Inc., and 431,094 UK Biobank participants. Using computationally detected DNA segments identical-by-descent (IBD) and runs of homozygosity (ROH), we identified 675 instances of UPD across both databases. Here we present the first characterization of UPD prevalence in the general population, a machine-learning framework to detect UPD using ROH, and a novel association between autism and UPD of chromosome 22.

## Introduction

Meiotic nondisjunction can have severe consequences for human reproduction and health. For example, nondisjunction can lead to aneuploidy, which is the leading cause of both spontaneous miscarriages and severe developmental disabilities^1–5^. Because determining the etiology of aneuploidy is extremely difficult in humans, many studies have instead focused on either studying the consequences of aneuploidy in individual cases presenting in the clinic or studying recombination events using population-genomic datasets in order to understand meiotic processes^3,6,7^. Recombination is an integral part of meiosis, facilitating alignment and then proper segregation of homologous chromosomes. Thus recombination at each chromosome pair in a human genome is generally regarded as necessary to prevent aneuploidy (with some exceptions^3,6^).

However, viable, euploid humans can result from aneuploid gametes if trisomic rescue, monosomic rescue, or gametic complementation restore normal ploidy during early development^8–12^. These processes can result in uniparental disomy (UPD), which is the inheritance of both homologs of a chromosome from only one parent with no representative copy from the other parent (Figure 1).

**Figure 1.**
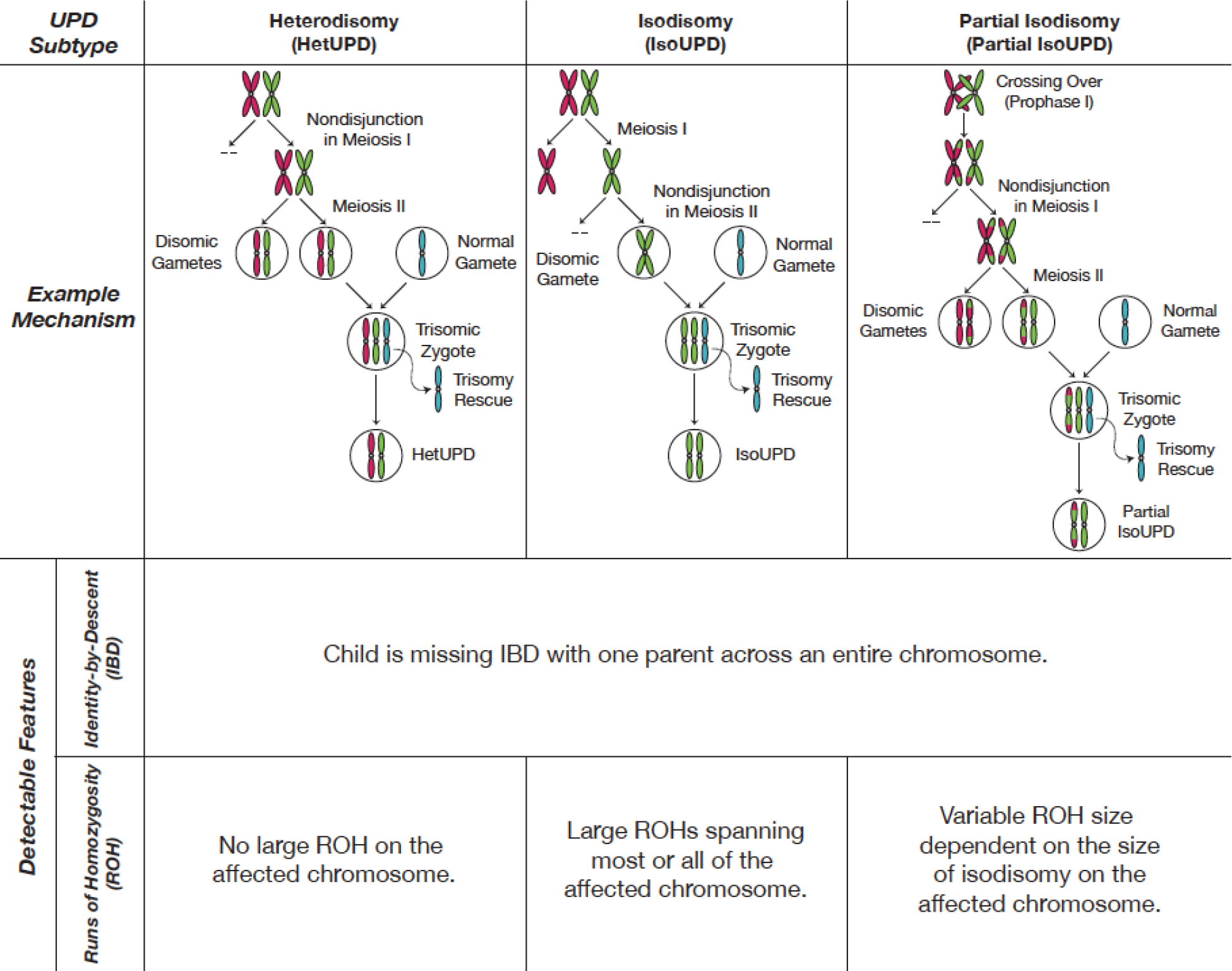
Subtypes of UPD with example mechanisms and detectable genomic features for each UPD subtype. There are three subtypes of UPD: heterodisomy (hetUPD), isodisomy (isoUPD), and partial isodisomy (partial isoUPD). HetUPD is caused by nondisjunction in meiosis I, and an affected individual will inherit both homologs of a chromosome from the same parent. IsoUPD is caused by nondisjunction in meiosis II, and an affected individual will inherit two identical copies of one homolog from one parent. Partial isoUPD is caused by nondisjunction in either meiosis I or meiosis II after crossing over has happened, resulting in sections of isodisomy and heterodisomy on the UPD chromosome. Given genomic data from a parent-child pair, all UPD subtypes can be detected in the same way based on identity-by-descent (IBD): a parent-child pair will be missing IBD across an entire chromosome. Lastly, isoUPD and some types of partial isoUPD will show large runs of homozygosity (ROHs), which can be detected computationally without the need for parental genotype data.

Since the first report of UPD in 1987^13,14^, ~3,300 cases of UPD have been described in the scientific literature (http://upd-tl.com/upd.html^10^). To date, UPD of each of the autosomes and the X chromosome has been documented^10,12,15^. UPD can cause clinical consequences by disrupting genomic imprinting, or by unmasking harmful recessive alleles in large blocks of homozygosity on the affected chromosome. Detecting UPD is a useful diagnostic tool for specific imprinting disorders and for rare Mendelian diseases caused by homozygosity^12,16–19^. UPD has also been implicated in tumorigenesis, particularly in cases of genome-wide UPD, which affects all the chromosomes in the genome^11,20^. Thus, current understanding of UPD is based largely on case reports of individuals in which UPD is detected following suspicion of an imprinting or other clinical disorder, and, in some instances (typically <10 confirmed cases) within larger case-control studies^12,16–19,21^.

There are three subtypes of UPD resulting from nondisjunction during different stages of meiosis (which we refer to as “meiotic-origin UPD”): isodisomy (“isoUPD”), heterodisomy (“hetUPD”) and partial isodisomy (“partial isoUPD”), which involves meiotic crossover (Figure 1). UPD can also be classified according to the parent of origin; when the disomic pair originates from the mother, the resulting case is termed maternal UPD (“matUPD”), and when the disomic pair originates from the father, the case is termed paternal UPD (“patUPD”). Despite the wealth of clinical UPD cases, prevalence and per-chromosome rates of UPD and its subtypes are not characterized in the general population. Past estimates of UPD prevalence include rates of one in 3500 and one in 5000^10,22^; these estimates were determined by extrapolation from UPD events causing clinical presentation and so do not account for variation in prevalence across chromosomes or for UPD associated with healthy phenotypes^22^. Therefore, to obtain an accurate estimate of UPD prevalence, hundreds of thousands of samples from the general population are needed^12^. And while chromosome recombination and segregation are regarded as highly constrained processes, population genetic datasets are now reaching large enough sizes to yield insight into normal variability in recombination within and among human genomes.

To address this gap, we detected instances of UPD in consented research participants from the direct to consumer genetics company 23andMe, Inc., whose database consists of single-nucleotide polymorphism (SNP) data from over 4.4 million individuals, and in 431,094 northern European UK Biobank participants. Here we present the first estimates of UPD prevalence in the general population, a new machine learning method to identify UPD in individuals without parental genotypes, and previously unrecognized phenotypes associated with UPD. We used both identity-by-descent (IBD) and a new supervised classification framework based on runs of homozygosity (ROH) to identify UPD while accounting for parental relatedness and differences in ROH length distributions between ancestral populations. We found that UPD is twice as common in the general population (estimated rate: one in 2000 births) than was previously thought and that, contrary to expectation, many individuals with long isodisomy events (ranging between 12 - 227 Mb) appear to have healthy phenotypes.

## Results

### UPD prevalence estimated from parent-child genotypes

Unlike typical parent-child pairs, individuals with UPD lack IBD segments across an entire chromosome with one parent; thus, we can use parent-child data to identify UPD cases. In the 23andMe dataset, we analyzed 916,712 parent-child pairs, which include 214,915 trios, to estimate the prevalence of UPD. We found 199 cases of UPD distributed across all 23 chromosomes except chromosome 18 (Figure 2A). Within 214,915 trios, we found 105 cases of UPD and estimate that UPD occurs with an overall prevalence rate of roughly one in 2000 births (rate: 0.05%; 99% CI: [0.04%, 0.06%]). Thus, we found that UPD is more common than previously thought (previous estimate based on UPD15 cases: one in 3500 births or 0.03%)^22^. Of the 105 true positives observed in 23andMe trios, 26 are patUPD cases and 79 are matUPD, suggesting that maternal origin UPD is three times as prevalent as paternal origin UPD. Within the 23andMe trios, four were “double UPD” cases, where two chromosomes in the same individual were inherited uniparentally; thus, we estimate that double UPD occurs at a rate of roughly one in 50,000 births. We also found that paternal partial isoUPD is the least common UPD subtype, and we observed more hetUPD and partial isoUPD cases than isoUPD cases (Figure 2B).

**Figure 2.**
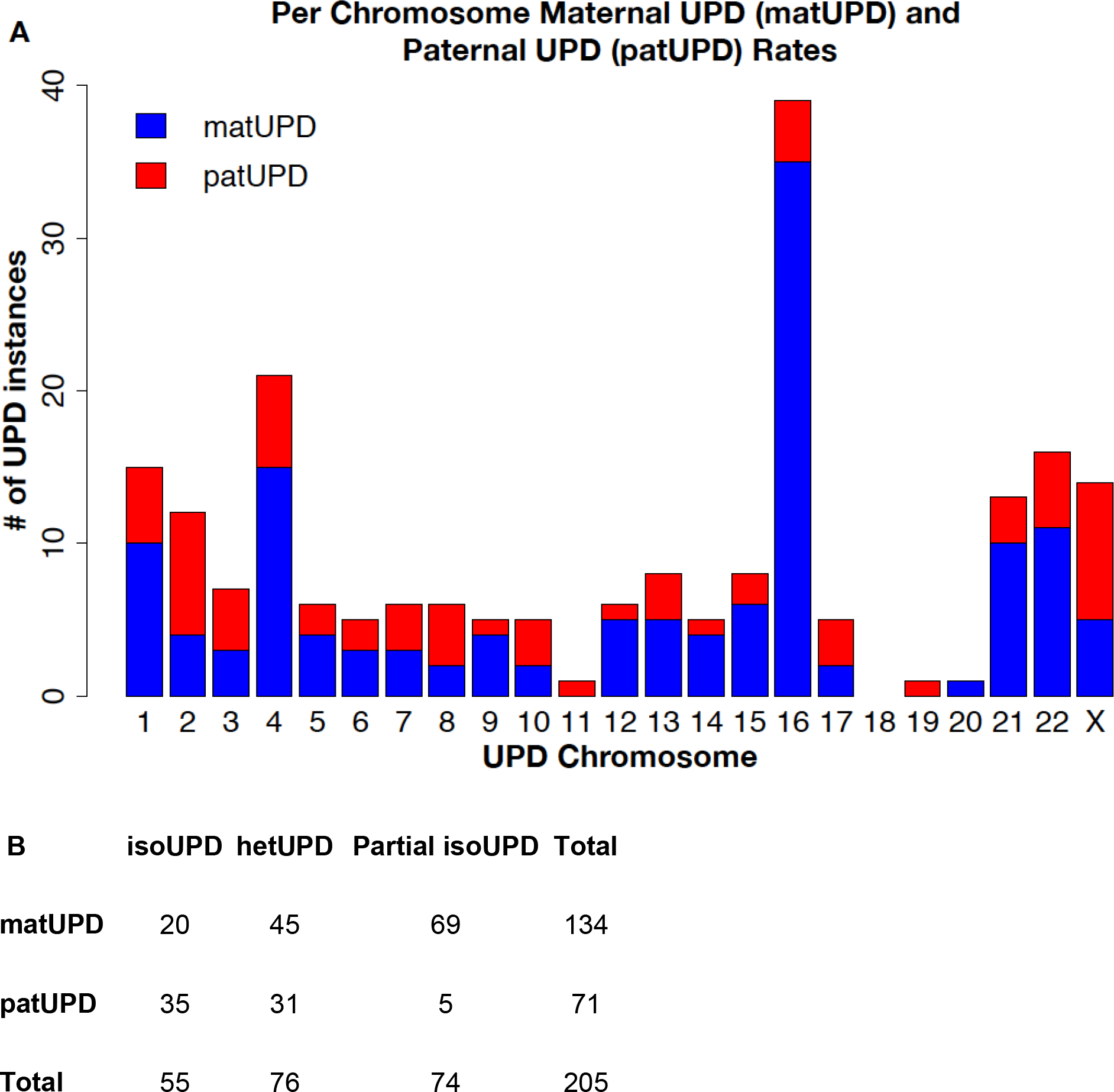
Using IBD-based UPD detection, we found 199 individuals with 205 incidences of UPD in 916,712 parent-child duos from the 23andMe dataset. We identified six individuals with cases of double UPD, when two chromosomes in one individual are inherited uniparentally. A) The per-chromosome distribution of true positives for UPD in the 23andMe dataset; in these true positives, UPD occurs most frequently on chromosomes 1, 4, 16, 21, 22 and X. We also observed three times as many maternal UPD (matUPD) cases as paternal UPD (patUPD) cases. B) We found that paternal partial isoUPD is the least common subtype of UPD. We also observed overall more hetUPD and partial isoUPD cases than isoUPD cases.

We compared the per-chromosome rates of UPD true positives in the 23andMe database to those from published reports of UPD in the literature (Supplementary Figure 1). We found that, while UPD true positives in the 23andMe database occur most frequently on chromosomes 1, 4, 16, 21, 22, and X, published UPD cases are most common on chromosomes 6, 7, 11, 14 and 15, which each contain clusters of imprinted genes that cause clinical phenotypes (http://upd-tl.com/upd.html^10^). We failed to reject the null hypothesis of independence between per-chromosome rates of 23andMe UPD true positives and published UPD cases (Fisher’s exact test; *p*-value = 1), and the two per-chromosome distributions are not significantly correlated (Pearson’s correlation; *p*-value = 0.72). Thus, we conclude that the clinical UPD cases are biased towards chromosomes where UPD causes clinical presentation, and do not represent the true distribution of UPD in the general population.

### New ROH-based UPD detection without parental genotypes

In many studies, parental genotypes for all probands may be too costly or logistically difficult to generate, and so classification of putative UPD cases may be made based on singleton genotypes only. Several clinical guidelines exist for prioritizing putative UPD cases for further analysis; all of these methods look for a large ROH confined to a single chromosome^23^.

However, multiple population-genetic studies have shown that relatively large ROH (>1Mb) are common even in outbred populations^24–27^; we recapitulate this result in eight cohorts in the 23andMe dataset (Supplementary Figure 2). Thus, an effective ROH-based method for UPD detection must be able to identify UPD chromosomes in the presence of large ROH on non-UPD chromosomes. Further, such a method must be able to distinguish between partial isoUPD and ROH blocks resulting from consanguinity.

To address these challenges, here we introduce a new supervised logistic regression classification framework that accounts for ROH length distributions within ancestral populations and is able to identify partial isoUPD and isoUPD on all autosomes and the X chromosome. In simulations across five cohorts in the 23andMe dataset (northern European, southern European, Latino, African American and East Asian individuals), we demonstrated that our classifiers achieve high power while minimizing the false positive rate (auROC > 0.9 for all classifiers; Figure 3A, Supplementary Figure 3). We also found that classifiers for larger chromosomes perform better than those for smaller chromosomes (Figure 3A; Supplementary Figure 3). In three cohorts (Ashkenazi Jewish, Middle Eastern and South Asians), we found that our classifiers performed poorly on simulated genotype data (auROC < 0.9; Supplementary Figure 3); when analyzing individual genotype data from these three cohorts in the 23andMe dataset, we classified thousands of putative cases which appeared to be false positives.

**Figure 3.**
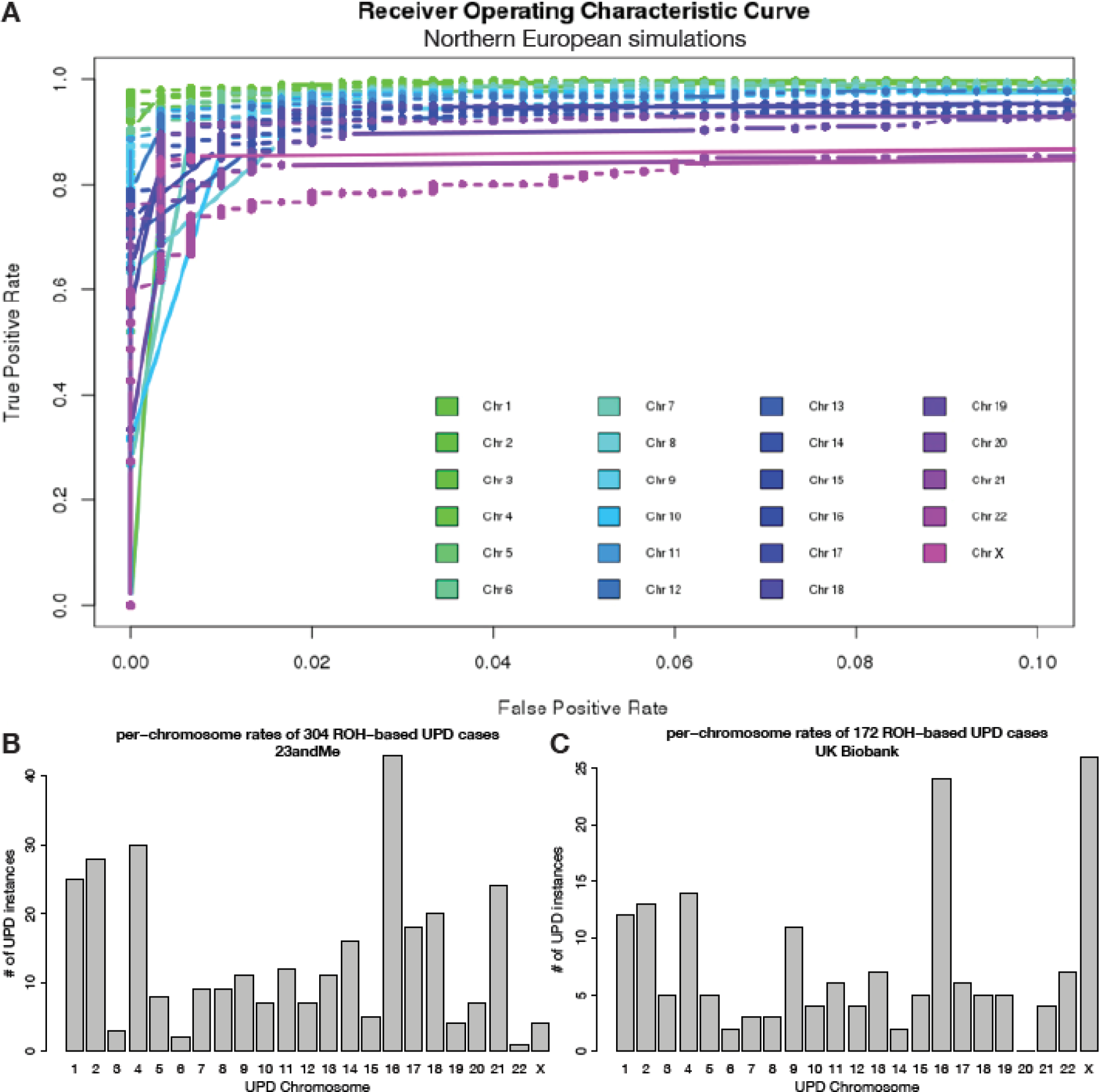
Our per-chromosome simulation-based classification framework identified 304 UPD cases using runs of homozygosity (ROH) across five cohorts (northern European, southern European, Latino, African American and East Asian individuals) in the 23andMe dataset and 172 ROH-based UPD cases in northern Europeans from the UK Biobank. A) Receiver Operating Characteristic (ROC) curves show the performance of our per-chromosome UPD classifiers on simulated testing data, based on genotype data from northern Europeans in the 23andMe dataset. Our classifiers identified UPD with high accuracy (area under the ROC curve (auROC) > 0.9; TPR between 0.75-0.98 when FPR is fixed at 0.01). At fixed FPR, power is inversely related to chromosome length. B) The chromosome distribution of the ROH-based cases found in the 23andMe dataset recapitulates features of the chromosome distribution of true positives for UPD, which are identified through IBD analysis (Figure 2A; Pearson’s correlation = 0.67; *p*-value = 0.0005). C) The chromosome distribution of the ROH-based cases found in the UK Biobank also recapitulates features of the chromosome distribution of true positives for UPD identified through IBD analysis (Figure 2A; Pearson’s correlation = 0.74; *p*-value = 4.79 x 10^−5^). We note that parent-of-origin cannot be identified for ROH-based cases.

Therefore, we ignored these three cohorts for further ROH-based UPD detection. As another form of validation, we applied our classifiers to northern European true positives (IBD-based UPD cases). We found that our classifiers identify 67% of true positives with greater than 20% isodisomy across the chromosome (Supplementary Table 1). In 1,371,138 singletons from five cohorts in the 23andMe dataset (northern European, southern European, Latino, African American and East Asian individuals), we classified 304 putative ROH-based UPD cases using our ROH-based method, 297 of which were newly discovered using ROH analysis (Figure 3B). The chromosome distribution of the ROH-based cases (Figure 3B) recapitulates the chromosome distribution of true positives identified through IBD analysis (Figure 2A; Pearson’s correlation = 0.67; *p*-value = 0.0005).

We also applied these 23 classifiers to data from 431,094 northern European individuals from the UK Biobank Project and identified 172 ROH-based UPD cases, observing cases of each chromosome except chromosome 20 (Figure 3C). The chromosome distribution of ROH-based cases (Figure 3C) in the UK Biobank also recapitulates the chromosome distribution of UPD cases identified through IBD analysis in the 23andMe database (Figure 2A; Pearson’s correlation = 0.74; *p*-value = 4.785 x 10^−5^). Karyograms of ROH in the 172 putative UPD cases from the UK Biobank show large blocks of homozygosity ranging from 12.7 Mb to 231 Mb (Supplementary Figure 4; Supplementary File 1).

**Figure 4.**
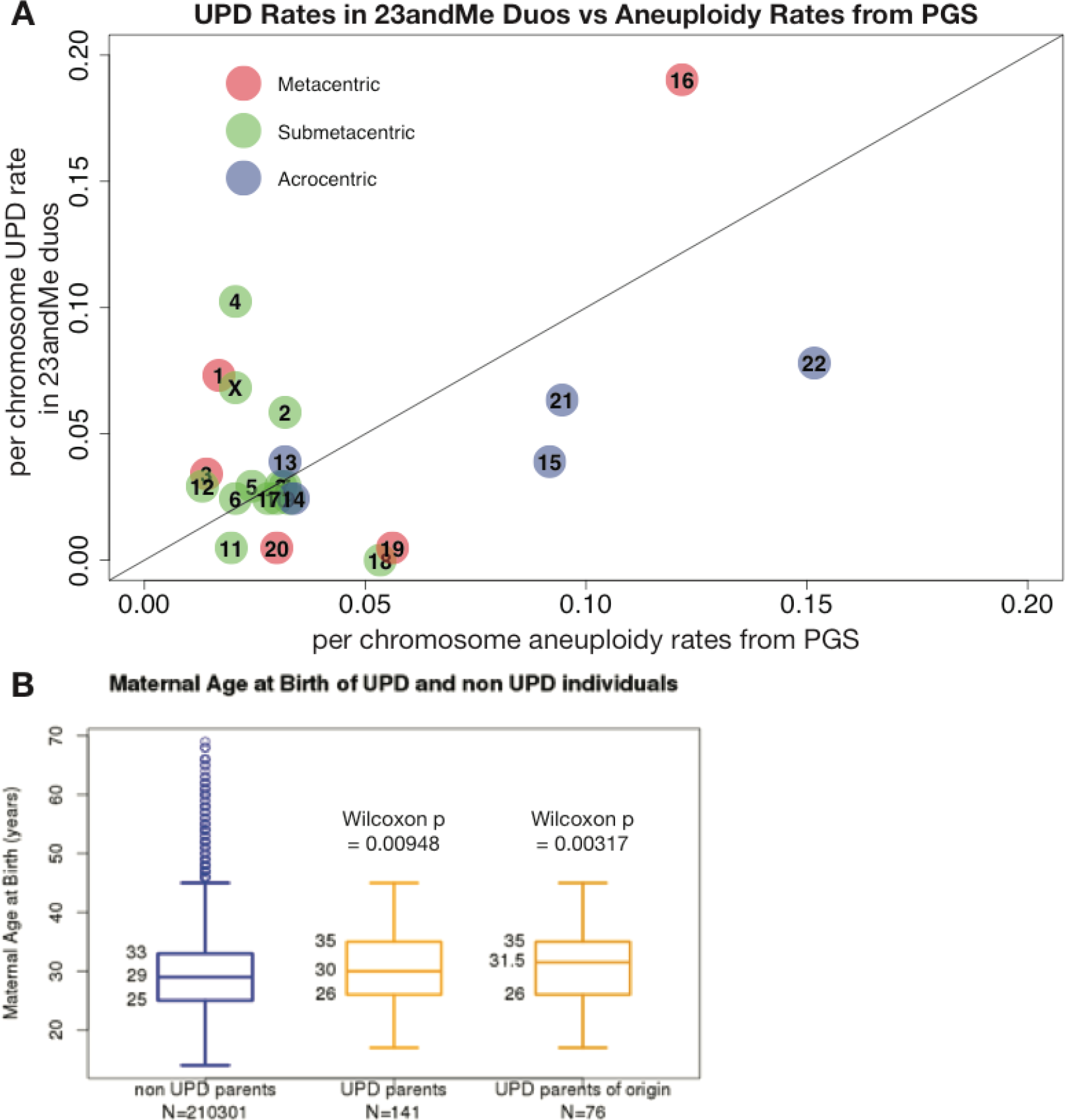
Aneuploidy rates and maternal age are correlated with UPD. A) The correlation between per-chromosome UPD rate in true positives from the 23andMe database and per chromosome aneuploidy rate in published pre-implantation genetic screening data^4^; chromosomes are colored by centromeric type: metacentric chromosomes are shown in red, submetacentric chromosomes in green and acrocentric chromosomes in blue. These two rates are significantly correlated (Pearson’s correlation = 0.49; *p*-value = 0.02) and this correlation remains significant after correction for chromosome length and centromeric type (Pearson’s correlation = 0.73; *p*-value = 0.006), suggesting that meiotic nondisjunction occurs more frequently on some chromosomes (such as 15, 16, 21, and 22) than others, resulting in more instances of both UPD and aneuploidy on these chromosomes. We also note that the acrocentric chromosomes have among the highest per-chromosome rates of both UPD and aneuploidy. B) The age distribution of mothers of UPD true negatives (blue) and that of mothers who are parents of origin of UPD true positives (matUPD cases, yellow) in the 23andMe dataset. We find that mothers of UPD cases are significantly older than mothers of UPD true negatives (Wilcoxon *p*-value = 0.00948) and that this associations holds when restricted to cases of matUPD, where mothers are the parents of origin of the UPD cases (Wilcoxon *p*-value = 0.00317).

### Phenotypic Consequences of UPD

UPD can cause phenotypic consequences in multiple ways, including 1) disrupting imprinting and 2) uncovering recessive alleles in blocks of isodisomy. We tested for phenotypic associations between UPD of each of the 23 chromosomes in true positives in the 23andMe dataset and 206 phenotypes across five categories (cognitive, personality, morphology, obesity and metabolic traits) obtained from self-reported survey answers. We found 23 nominally significant (*p*-value < 0.01) phenotype associations with UPD of chromosomes 1, 3, 6, 7, 8, 15, 16, 21 and 22 (Supplementary Table 2). While some of these 23 associations were driven by a single UPD case, three associations had multiple cases (or multiple measurements, in the case of quantitative traits), representing a more robust signal: we found that UPD6 is associated with lower weight (*p*-value = 0.0038) and shorter height (*p*-value = 0.0055), and UPD22 is associated with a higher risk for autism (*p*-value = 2.557 x 10^−5^) (Table 1).

**Table 1.** PheWAS in the 23andMe dataset identified phenotypes significantly associated with UPD of chromosomes 6 and 22 (*p*-value < 0.01). Only traits with at least two cases (or two measurements for quantitative traits) are shown; the full list of all associations is shown in Supplementary Table 2. Effect sizes shown are odd ratios. We tested for association between UPD on each of the autosomes and 206 self-reported phenotypes across five categories (cognitive, personality, morphology, obesity and metabolic traits). We uncover novel associations between UPD of chromosome 6 and weight and height, and UPD of chromosome 22 and autism spectrum (this last association remains significant after correcting for the number of self-reported phenotypes tested).

| UPD type | Phenotype | Effect Size (95% CI) | P-values (Uncorrected) |
| --- | --- | --- | --- |
| UPD6 | Weight | -2.02 (-3.38 -0.65) | 0.0038 |
| UPD6 | Height | -1.99 (-3.40 -0.59) | 0.0055 |
| UPD22 | Autism Spectrum | 3.61 (1.93 5.30) | 2.557 x 10<sup>-5</sup> |

### Variants Associated with UPD Incidence

Although heritability of UPD, or chromosomal aneuploidy, has not been reported, there may be genetic variants that predispose individuals to produce aneuploid germ cells, thus increasing the likelihood of giving birth to an offspring with UPD. We tested this hypothesis by performing a genome-wide association study (GWAS) comparing 221 parents of UPD cases to 205,141 parents of UPD true negatives from IBD analysis. We performed the analysis in all parents, adjusted for age, and stratified by parental sex. No association reached genome-wide significance (*p*-value = 5 x 10^−8^), and the heritability estimated by LD score regression^28^ was non-significant across all three analyses (Supplementary Figure 5; *p*-value > 0.05). Given the small sample size, this likely reflects our lack of power to detect genetic associations even for common variants associated with UPD.

In order to further investigate the etiology of UPD, we assess the relationship between per-chromosome UPD rates and per-chromosomes rates of aneuploidy in pre-implantation embryos (PGS). We found that UPD rates in the 23andMe database are significantly correlated with published aneuploidy rates from PGS^4^ (Figure 4A; Pearson’s correlation = 0.49; *p*-value = 0.02). We also found that mothers who are parents of origin of UPD true positives in the 23andMe dataset are significantly older than those of UPD true negatives (Figure 4B; Wilcoxon *p*-value = 0.00317), whereas paternal age does not show a robust association with UPD (Supplementary Figure 6; Wilcoxon *p*-value = 0.286).

## Discussion

The recombination process has long been studied by evolutionary, medical, molecular, and population geneticists, in part to gain insight into meiotic nondisjunction. It is difficult to directly study meiotic nondisjunction in humans because errors in meiosis often lead to fetal loss or serious health consequences. However, UPD is a detectable genomic signature of meiotic nondisjunction and aneuploidy in euploid, liveborn individuals. In this study we show that, given large genomic datasets, detecting UPD offers new insight into recombination and meiosis in humans.

Firstly, using 214,915 trios in the 23andMe customer database, we obtained the first estimate of UPD prevalence in the general population: one in 2000 births, 1.75 times higher than the current clinical estimate of one in 3500 births. The current estimate of UPD prevalence is derived from UPD15 prevalence in clinical cohorts, which might not be representative of the general population and also does not account for differences in UPD prevalence between chromosomes^22^. The 23andMe customer base comprises, for the most part, healthy individuals from the general population and so our estimate is more representative of overall UPD prevalence. We also found that the per-chromosome prevalence rate of UPD is significantly correlated with per-chromosome aneuploidy rates calculated from published PGS data (Figure 4A; Pearson’s correlation = 0.49; *p*-value = 0.02) whereas per-chromosome rates from clinical UPD cases are not (Supplementary Figure 7; Pearson’s correlation = 0.2; *p*-value = 0.34).

Since a liveborn individual with UPD results from the restoration of euploidy in an aneuploid zygote, we expect the true per-chromosome rates of UPD to be correlated with those of aneuploidy, providing further evidence that our estimated rates are closer to the true prevalence and per chromosome distribution of UPD than existing clinical rates. We note that participation in 23andMe may be cost-prohibitive for many, and also that the customer base may be biased toward geographic regions or other covariates. Furthermore, individuals with severe health problems may be unlikely or unable to participate in 23andMe, and so the UPD cases in this study may be depleted for UPD causing serious health consequences.

Secondly, we have introduced a new method to find UPD cases using genomic data from singleton data only, without requiring parental genotypes. Existing guidelines for classification of putative UPD cases without parental genotypes consist of a hard ROH length threshold for all chromosomes^23^. These putative cases can then be further investigated by genotyping parents, cytogenetic techniques or DNA methylation studies. However, ROH length distributions vary by 1) the demographic history of an individual’s ancestral population(s), 2) the history of consanguinity in the individual’s recent ancestors, and 3) by chromosome. Our method learns the distributions of ROH lengths on each chromosome from simulated data based on each of eight global population cohorts in the 23andMe dataset while also modeling recent consanguinity and is able to classify UPD with high accuracy. Using our method, we were able to find 297 additional UPD cases in 1,371,138 individuals in the 23andMe cohort. Our classifiers can also be readily applied to other genomic datasets such as the UK Biobank^29,30^, in which we identified 172 additional ROH-based UPD cases (Figure 3C, Supplementary Figure 4, Supplementary File 1). This underscores that an effective ROH-based method for UPD detection offers crucial insight into UPD when combined with large-scale genomic datasets. One limitation of our ROH-based detection method is that we can only identify isoUPD and partial isoUPD cases that contain large blocks of homozygosity (spanning greater than 30% of the chromosome). In that respect, UPD per-chromosome rates estimated using our ROH-based method are conservative. In order to minimize the false positive rate, we did not try to refine classification of small partial isoUPDs (Supplementary Table 1; FPR: 7 x 10^−5^). Also, we were unable to identify UPD in populations that are historically known to practice endogamy and thus have higher than average levels of homozygosity (Ashkenazi Jewish, Middle Eastern and South Asian individuals; Supplementary Figure 3).

Errors in recombination typically, with few exceptions^3,6^, lead to aneuploidy and severe health consequences, and so are largely viewed as deleterious. However, the majority of UPD types, including the most common UPD (UPD16), did not show significant, plausible associations with deleterious traits in the 23andMe database (Table 1). Our work challenges the typical view that errors in recombination are strongly deleterious, showing that even in extreme cases where individuals are homozygous for an entire chromosome, those individuals can be, to the best of our knowledge, phenotypically normal and healthy (Table 1). We do find a novel association between autism and UPD22. We note that phenotype data in the 23andMe database is self-reported and so depends on customers answering surveys about their health and traits. We also note that there has yet to be a prospective study of the long-term consequences of UPD since most current studies focus on special syndromes and recessive disorders that are apparent in childhood; future studies could extend our work in this direction.

To interrogate the role of genetics in UPD etiology, we performed the first GWAS of UPD. Though our results are mostly suggestive, with increased sample sizes or deep sequencing, future studies may find plausible, significant loci underlying UPD incidence.

Lastly, we expect the etiology of meiotic nondisjunction and UPD to be similar since UPD is caused by rescue of aneuploid zygotes. Here, we found that UPD rates in the 23andMe database are significantly correlated with aneuploidy rates from PGS (Figure 4A; Pearson’s correlation = 0.49; *p*-value = 0.02). Also, similarly to aneuploidy, we found that mothers who are parents of origin of UPD true positives in the 23andMe dataset are significantly older than those of UPD true negatives (Figure 4B; Wilcoxon *p*-value = 0.00317). Previous studies have shown elevated escape from crossover inference on certain chromosomes (8, 9, and 16) and especially in older mothers; future studies could test whether crossover interference rates vary between UPD cases and UPD true negatives^7^. And though we focused in this study on meiotic-origin UPD, future studies could also extend our work to characterize the prevalence and chromosomal distribution of segmental (or mitotic-origin) UPD cases in the general population; segmental UPD is also currently only studied in clinical settings^8^.

## Online Methods

### Samples

In this study, we analyzed genome-wide SNP genotypes from 4,400,363 research participants from the 23andMe customer base; this research platform has been previously described^31,32^. All research participants included in these analyses provided consent and answered surveys online according to a human subjects protocol approved by Ethical and Independent Review Services, an independent institutional review board (http://www.eanireview.com). We also analyzed genotype data from 500,000 participants in the UK Biobank Project^29,30^ (www.ukbiobank.ac.uk). Phenotype data for these individuals were collected through questionnaires, interviews, health records, physical measurements, and imaging carried out at assessment centers across the UK.

### Genotyping and Quality Control

For the 23andMe dataset, DNA extraction and genotyping were performed on saliva samples by clinical laboratories at Laboratory Corporation of America, which is certified by Clinical Laboratory Improvement Amendments and accredited by College of American Pathologists. Samples were genotyped on one of five Illumina platforms: 1) and 2) two versions of the Illumina HumanHap550+ BeadChip, plus about 25,000 custom SNPs selected by 23andMe (~560,000 SNPs total), 3) a variation on the Illumina OmniExpress+ BeadChip, with custom SNPs (~950,000 SNPs total), 4) a fully customized array, including a lower redundancy subset of v2 and v3 SNPs with additional coverage of lower-frequency coding variation (~570,000 SNPs total) and 5) a customized array based on Illumina’s Global Screening Array (~640,000 SNPs total), supplemented with ~50,000 SNPs of custom content. Samples that failed to reach 98.5% call rate were re-analyzed. Individuals whose analyses failed repeatedly were re-contacted by 23andMe customer service to provide additional samples. For all ROH analyses, we limited our analyses to SNPs that are shared between the Illumina platforms 1 - 4 described above, and then we removed SNPs with a minor allele frequency (MAF) less than 5% and SNPs with genotyping rate less than 99%, resulting in 381,379 SNPs in total. For IBD analyses, we analyzed 579,957 SNPs for all individuals.

For the UK Biobank dataset, quality control was carried out as described in the UK Biobank genotyping quality control document^30^. As was done in the 23andMe dataset, we then removed all SNPs with MAF less than 5% and SNPs with genotyping rate less than 99%, resulting in 360,540 SNPs total.

### Ancestry Classification

23andMe’s ancestry analysis has been described previously^33^. Briefly, the algorithm first partitions phased genomic data into short windows of about 300 SNPs. Within each window, a support vector machine (SVM) was used to classify individual haplotypes into one of 25 reference populations (https://www.23andme.com/ancestry-composition-guide/). The SVM classifications are then fed into a hidden Markov model (HMM) that accounts for switch errors and incorrect assignments and gives probabilities for each reference population in each window. Finally, simulated admixed individuals were used to recalibrate the HMM probabilities so that the reported assignments are consistent with the simulated admixture proportions. The reference population data is derived from public datasets (the Human Genome Diversity Project, HapMap, and 1000 Genomes), as well as 23andMe research participants who have reported having four grandparents from a single country.

### Population Structure

For population-specific analyses such as ROH detection in the 23andMe dataset, research participants were divided into eight cohorts based on genome-wide ancestry proportions from reference populations, as determined by 23andMe’s Ancestry Composition method: northern Europeans, southern Europeans, African Americans, Ashkenazi Jewish, East Asians, South Asians, Latino/as, and Middle Eastern individuals. The classification criteria have been previously described in Campbell *et al.*^7^. Briefly, individuals labeled as northern European met all of the following criteria: greater than 97% European and Middle Eastern/northern African ancestry combined, greater than 90% European ancestry, and greater than 85% northern European ancestry. Southern European individuals satisfied the following requirements: greater than 97% European and Middle Eastern/northern African ancestry combined, greater than 90% European ancestry, and greater than 85% southern European ancestry. Ashkenazi Jewish individuals had greater than 97% European and Middle Eastern/northern African ancestry combined, greater than 90% European ancestry, and greater than 85% Ashkenazi Jewish ancestry. Middle Eastern individuals had greater than 97% European and Middle Eastern/northern African ancestry combined and greater than 70% Middle Eastern/northern African ancestry. East Asian individuals had greater than 97% East Asian and Southeast Asian ancestry combined. South Asians had greater than 97% South Asian ancestry. Individuals were classified as African Americans or Latinos/Latinas if they had greater than 90% European and African and East Asian/Native American and Middle Eastern/northern African ancestry combined as well as greater than 1% African and American ancestry. African Americans and Latino/as were distinguished using a logistic regression classifier trained on self-identified “Black African” and “Hispanic” individuals^7^.

In the UK Biobank dataset, we focused our analyses on 431,094 individuals of northern European ancestry identified by principal components analysis (PCA) on the genotype data following QC. We arrived at this sample as follows: we first performed PCA using FlashPCA2^34^ (version: 2.0) on 2,504 individuals from the 1000 Genomes Project Phase 3 database^35^. We pruned the genotype data from the UK Biobank for linkage disequilibrium, resulting in 70,527 SNPs, and we then projected genotype data from 431,102 UK Biobank participants who self-identified as “white British” onto the principal components space derived from PCA of the 1000 Genomes individuals. Lastly, we removed eight individuals who were outliers in PCA with the following thresholds: first principal component value less than 0.03 (PC1 < 0.03), and second principal component value greater than 0.15 (PC2 > 0.15). These filtering steps resulted in 431,094 northern European individuals.

### Identification of Parent-Child Duos from Identity-by-Descent Segments

We identified identical-by-descent (IBD) DNA segments for every pair of individuals in the 23andMe dataset, according to a method that has been previously described by Henn *et al.*^36^. Briefly, we compared a pair of individuals’ genotypes at 579,957 SNPs and identified SNPs where the individuals are homozygous for different alleles (also called “opposite homozygotes”). Long regions (>5 cM) lacking opposite homozygotes were characterized as “IBD segments”^36^.

Pairs of individuals that share more than 85% of their genome IBD were classified as parent and child. Theoretically, parent-child pairs should share 100% of their genome IBD on one homologous chromosome, but the threshold is lowered here to 85% to account for the possibility of UPD of chromosome 1, which accounts for ~10% of the genome, and for the possibility of error or lack of SNP coverage over ~5% of the genome. Using these criteria, we identified 916,712 parent-child duos in the 23andMe database.

### Identification of Runs of Homozygosity

We calculated runs of homozygosity (ROH) using GARLIC^26,37^. Briefly, GARLIC implements a model-based method for identifying ROH and classifying ROH into length classes. In this method, logarithm of the odds (LOD) scores for autozygosity are calculated in sliding windows of SNPs across the genome; SNP window sizes were chosen automatically by GARLIC based on SNP density. The LOD scores are functions of a user-specified error rate to account for genotyping error and mutation rate, as well as population-specific allele frequencies. The distribution of LOD scores is used to determine a threshold for ROH-calling. After ROH are identified, contiguous ROH windows are concatenated. Lastly, we performed Gaussian mixture modeling using the Mclust function from the mclust R package^38^ (Version 5.4) with the same parameters used by Kang *et al.*^27^ to classify ROH into three length classes: 1) Class A, which are the shortest ROH; 2) Class B; and 3) Class C, which are the longest (class boundaries shown in Supplementary Table 3).

We applied GARLIC to the eight cohorts in the 23andMe database described earlier: 974,511 northern Europeans, 34,508 southern Europeans, 90,349 African Americans, 70,144 Ashkenazi Jewish, 63,683 East Asians, 19,493 South Asians, 208,087 Latino/as, and 16,013 Middle Eastern individuals as well as 431,094 northern Europeans in the UK Biobank dataset. In the 23andMe dataset, we used a window size of 60 SNPs for autosomes and 30 SNPs for the X chromosome, which were automatically chosen by GARLIC as the best window size given SNP density, and an error rate of 0.001, which was used in previous studies of ROH^26,37^. In the UK Biobank dataset, we used a window size of 60 SNPs for both autosomes and the X chromosome, which were automatically chosen by GARLIC as the best window size for these data. In each dataset, only females were included in analyses of ROH on the X chromosome.

In the 23andMe database, population-specific allele frequencies were calculated from individuals who are true negatives for UPD (identified as described in “IBD-based UPD Detection”); sample sizes of true negatives in the 23andMe cohorts are as follows: 28,338 northern Europeans, 1,018 southern Europeans, 1,500 African Americans, 2,066 Ashkenazi Jewish, 2031 East Asians, 982 South Asians, 7,639 Latino/as, and 437 Middle Eastern individuals. In the UK Biobank cohort, we calculated allele frequencies for GARLIC from 431,094 individuals of northern European ancestry. All Class C ROH were then filtered for deletions as described in “Filtering of Deleted Genomic Regions”.

### IBD-based UPD Detection

To detect UPD events in children, we looked for parent-child duos who lack IBD segments across an entire chromosome. For the X chromosome, only the following parent-child pairs, which would normally be expected to share IBD on the X chromosome, were considered: mother-daughter pairs, father-daughter pairs, and mother-son pairs. The putative UPD cases were then tested for a deletion spanning the putative UPD chromosome in both the parent and child according to the method described in “Filtering of Deleted Genomic Regions” below; we refer to children in parent-child pairs without deletion of the putative UPD chromosome as true positives for UPD. IBD segments can also be used to determine true negatives for UPD; we identified children in trios who are completely half identical to both parents and refer to these as true negatives. In order to calculate prevalence of UPD, we focus on trio data since only then can both parent-child pairs be tested for missing IBD.

To distinguish between maternal UPD (matUPD; when the disomic chromosome pair originates from the mother) and paternal UPD (patUPD; when the disomic chromosome pair originates from the father), we labeled the older individual in a parent-child duo as the parent and the younger individual as the child using self-reported age data. If an individual was missing IBD across a chromosome with the father, we labeled the case as maternal UPD (matUPD). If an individual was missing IBD with their mother, we labeled the case as paternal UPD (patUPD).

For UPD cases with a mother and father genotyped in the 23andMe database, we were able to use IBD with the parent-of-origin of the disomic chromosome pair to differentiate between the three subtypes of UPD: isodisomy (isoUPD), heterodisomy (hetUPD), and partial isodisomy (partial isoUPD). IsoUPD chromosomes are completely half-identical to the parent-of-origin, and hetUPD chromosomes are completely identical to the parent-of-origin. Partial isoUPD chromosomes are some fraction half-identical and some fraction fully identical to the parent-of-origin. For UPD cases detected in parent-child duos and lacking genotype data for the parent-of-origin, we use ROH to differentiate between the three subtypes. UPD chromosomes with ROH spanning 100% of the chromosome are labeled isoUPD, UPD chromosomes with 0% Class C ROH are labeled hetUPD, and UPD chromosomes with between 0% and 100% Class C ROH are labeled partial isoUPD.

### Filtering of Deleted Genomic Regions

Large deletions — which can arise from somatic events or be prevalent in low-quality saliva samples — can manifest in genotype data as large regions of homozygosity and missing IBD that confound UPD detection. Thus, we screened all putative UPD cases for deletions. We filtered for deletions in one of two ways: by testing for significantly decreased Log R Ratio (LRR) across an ROH, and by using the CNV caller in BCFtools^39^ (version: 1.4.1). LRR, which is a measure of probe intensity, can be used to detect several types of copy number variants^40^. Theoretically, LRR is 0 at all loci across the genome, and decreases across a deleted region. Therefore, we tested whether the average LRR of a given region (i.e. a run of homozygosity) was significantly lower than the genome-wide average LRR using a two-sample t-test. Runs of homozygosity with significantly lower LRR than the genome-wide average (*p*-value < 0.05) were filtered out of all ROH-based analyses.

For a chromosome missing IBD between a parent-child pair, we used the CNV calling function of BCFtools to identify whole chromosome deletions and trisomies/mosaic trisomies in the parent and child at the putative UPD chromosome^39^. The command line option “-l 0.8” was used to upweight LRR within the HMM model relative to BAF; when BAF is given equal weight with LRR, we found that blocks of isodisomy were called as deletions even without a corresponding decrease in LRR across the homozygous region. If more than 40% of a putative chromosome in either parent or child is called with copy number of 1 (deletion), the pair was excluded from further analysis. If more than 50% of the chromosome in either parent or child is called with copy number of 3 (trisomy), the pair was also excluded.

### Simulation of Training Data for ROH-based UPD Classifiers

We generated training data for 23 logistic regression-based classifiers for eight cohorts (described in “Population Structure”) to detect UPD of each of the autosomes and the X chromosome without parental genotype data, as follows. We simulated 46,000 individuals, consisting of 1,000 UPD cases and 1,000 controls for each chromosome for each of the eight cohorts, by randomly pairing 46,000 pairs of individuals across all cohorts. Only individuals without any Class C ROH-length deletions and pairs sharing less than 930cM IBD were considered for selection as “parents” for the simulated children. To model consanguinity within our training data, we forced at least 100 pairs of 1000 controls for each chromosome to share between 100 and 930 cM IBD. For our X chromosome classifiers, only female children were simulated since two X chromosomes are required to detect homozygosity on the X chromosome.

We generated 2000 independent trios with one child each to train each classifier. We randomly sampled recombination breakpoints according to a distribution of crossover probabilities for each locus. The probability of at least one crossover between every pair of adjacent loci was calculated using Equation 1 below, in which we assume that crossovers are Poisson distributed with a rate equal to the difference in genetic distance in cM between two given loci multiplied by 0.01 (since there is approximately one crossover event in 100 cM).

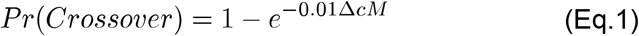

To calculate genetic distances, we used recombination maps ascertained from the 23andMe research cohort^7^. The genetic maps are publicly available at the following URL: https://github.com/auton1/Campbell_et_al. We then simulated meiosis by randomly choosing one homolog from each parent to be inherited by the child. For each chromosome’s classifier, we simulated UPD cases by deriving both homologs of that chromosome from only one parent. Lastly, we copied genotypes from parental homologs to generate genotypes for the child.

We used GARLIC to detect ROH for each simulated child using the same parameters and population allele frequencies specified in “Identification of Runs of Homozygosity and Genomic Hot/Coldspots for ROH”. We found that ROH length distributions were not significantly different between the simulated UPD cases and true positive UPD cases ascertained through IBD-based UPD detection (*p*-value > 0.05) (Supplementary Figure 8A). However, the distribution of the number of class C ROH differed significantly between the simulated UPD cases and real UPD cases (*p*-value < 0.05), and thus, we did not use the number of Class C ROHs to train the classifiers (Supplementary Figure 8B). Our simulations produced every subtype of UPD (Supplementary Figures 9-10).

### ROH-based UPD Detection and Performance Assessment

We developed 23 logistic regression classifiers, one for each autosome and the X chromosome, with two independent variables, trained on the simulations described in the previous section. For a given chromosome *i*, where *i* ∈ {1 … *n*} and *n* = 23 in females and *n* = 22 in males, let *c*_*i*_ be the total class C ROH length in base pairs. Also, let *c*_(*i*)_ be the ith order statistic for n class C ROH lengths across all chromosomes, where *c*_(*n*)_ is the maximum ROH length across all chromosomes. The two variables we trained each classifier on are *c*_*i*_ for *i* ∈ {1 … *n*} and *c*_(*n*−1)_⁄*c*_(*n*)_. We focused on Class C ROH for training the classifiers because, in comparing the distributions of ROH lengths between true positives (UPD cases detected through IBD) and true negatives for UPD, we found that only Class C ROH length is significantly different (*p*-value < 0.05) between the true positives and true negatives (Supplementary Figure 11).

To assess the performance of our classifiers, we generated Receiver Operating Characteristic (ROC) curves by testing each classifier on: (1) a simulated set of 11,500 individuals, consisting of 250 cases and 250 controls for each chromosome, and independent from the training simulations described in the previous section; and (2) the set of true positives and true negatives for UPD ascertained from IBD-based UPD detection. In testing, we found that performance of the classifiers, as measured by area under a ROC curve (auROC), increases with increased proportion of isodisomy on the UPD chromosome (Supplementary Figure 12). Specifically, detecting hetUPD without parental data is not possible due to the lack of large ROH blocks, and partial isoUPD detection is dependent on the size of the ROH. Thus, we restricted the training set further to only comprise true positives with at least 30% isoUPD on the UPD chromosome and a randomly sampled set of simulated controls equal in number to the cases for a given chromosome. We then chose initial probability cutoffs for each classifier from its ROC curve; we chose the probability threshold that minimized the False Positive Rate (FPR) and if there were multiple cutoffs that satisfied these criteria, we then chose the cutoff that also maximized the True Positive Rate (TPR). We used these cutoffs to classify putative UPD cases of each chromosome and remove duplicate cases; this last step removes individuals with large blocks of ROH on multiple chromosomes due to recent consanguinity. We then chose a final probability cutoff for our classifiers of 0.9 to classify ROH-based UPD cases; when applied to the known northern European true positives and true negatives, this cutoff allows us to minimize false positive rate (7 x 10^−5^) while maintaining a true positive rate of 67% for isoUPD spanning greater than 20% of the chromosome (Supplementary Table 1).

### Phenotypic Association Studies (PheWAS) in the 23andMe dataset

We regressed 206 phenotypes across five categories (cognitive, personality, morphology, obesity and metabolic traits) onto UPD status, using children with UPD (true positives detected using IBD analysis) as cases and true negatives for UPD as controls. We tested each subtype of UPD (by chromosome and parent-of-origin) separately, and we restricted these analyses to individuals of European ancestry. We performed logistic regression for binary traits and linear regression for quantitative traits with the following covariates: age, sex, genotyping platform and the first five principal components (see “Genome-Wide Association Studies (GWAS) in the 23andMe dataset” section) to adjust for population substructure. We also tested for differences in parental age between UPD true positives and true negatives.

### Genome-Wide Association Studies (GWAS) in the 23andMe dataset

We conducted GWAS on parents of UPD true positives identified by IBD-based analysis in order to find loci associated with the risk of giving birth to children with UPD. The GWAS was performed on all SNPs that passed quality control by running a logistic regression model correcting for the effects of age, parental age at birth of the child with UPD, first five genetic principal components, and genotype platform, performed separately by sex of the parents (mother or father). For more details on GWAS, imputation, and PCA, please refer to the Supplementary Methods.

## Acknowledgments

We thank the 23andMe research participants who made this work possible. We also thank the employees of 23andMe who developed the infrastructure that made this research possible. The research has been conducted using the UK Biobank Resource under Application Number 44606. We gratefully acknowledge Uta Francke, Shai Carmi, Aaron Carrel, Kirk Lohmueller, Priya Moorjani, John Novembre, Ben Raphael, Suyash Shringarpure, Janie Shelton, and the Ramachandran Lab for helpful conversations. This research was supported in part by US National Institutes of Health (NIH) grant R01GM118652, NIH COBRE award P20GM109035, and National Science Foundation (NSF) CAREER award DBI-1452622 to S. Ramachandran. Support was also provided by the NIH funded by the National Child Health and Development Institute training grant K12HD052896 to A. O’Donnell-Luria.

Members of the 23andMe Research Team: Michelle Agee, Babak Alipanahi, Adam Auton, Robert K. Bell, Katarzyna Bryc, Sarah L. Elson, Pierre Fontanillas, Nicholas A. Furlotte, David A. Hinds, Karen E. Huber, Aaron Kleinman, Nadia K. Litterman, Jennifer C. McCreight, Matthew H. McIntyre, Elizabeth S. Noblin, Carrie A.M. Northover, Steven J. Pitts, Olga V. Sazonova, Janie F. Shelton, Suyash Shringarpure, Chao Tian, Joyce Y. Tung, Vladimir Vacic, and Catherine H. Wilson.

## Supplementary Information

**Supplementary Figure 1.**
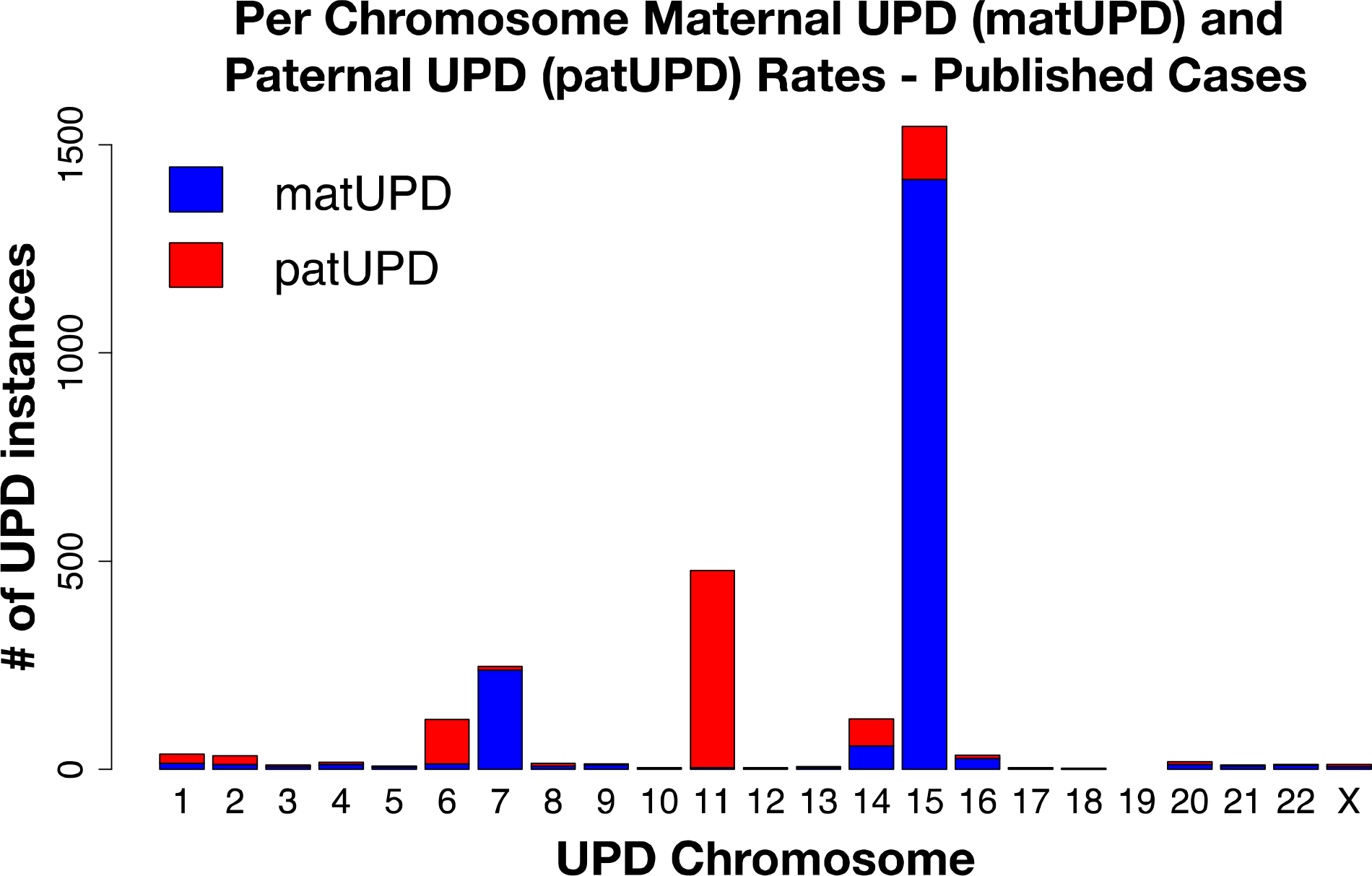
The per chromosome distribution of clinical UPD cases published in the literature to date (http://upd-tl.com/upd.html^1^, accessed 11/29/18). More than one case has been observed on each autosome except 19 and the X chromosome. Published UPD cases seem to cluster on chromosomes 6, 7, 11, 14 and 15, which contain clusters of imprinted genes that cause clinical phenotypes. There are 1869 matUPD cases in total and 881 patUPD cases in total, suggesting that matUPD is about twice as common as patUPD.

**Supplementary Figure 2.**
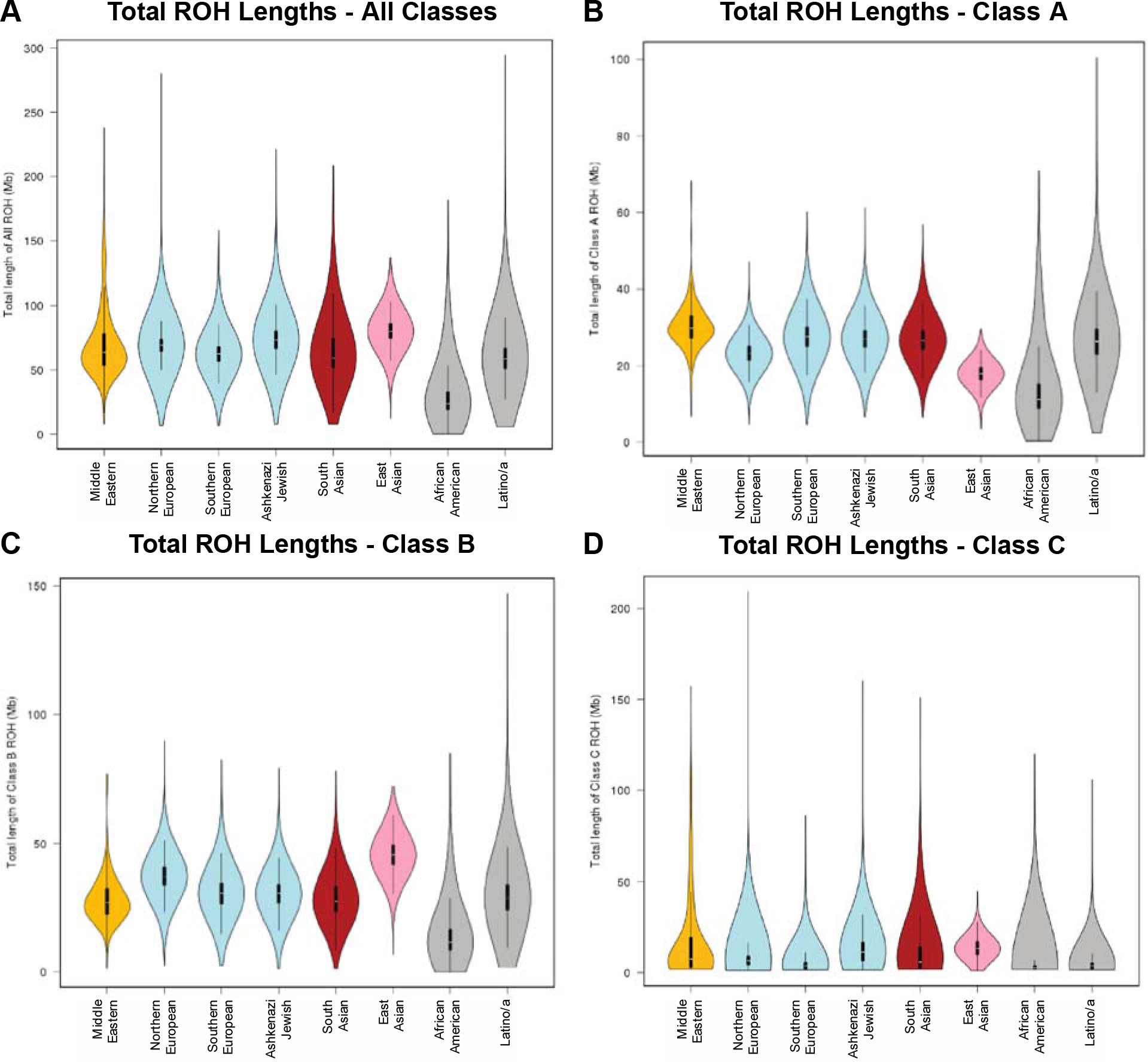
Violin plots of ROH length distributions (in Mb) for eight cohorts in the 23andMe database. The cohorts are colored by continental ancestry group: Middle Eastern (yellow), European (blue), South Asian (red), East Asian (pink), and admixed (grey). ROH were identified using GARLIC, which divides inferred ROH into three classes based on length. These plots show A) all classes combined, B) class A, the shortest ROH, C) class B, intermediate length ROH, and D) class C, the longest ROH. These plots recapitulate patterns of ROH length distributions seen in published analyses of ROH across global populations^2,3^.

**Supplementary Figure 3.**
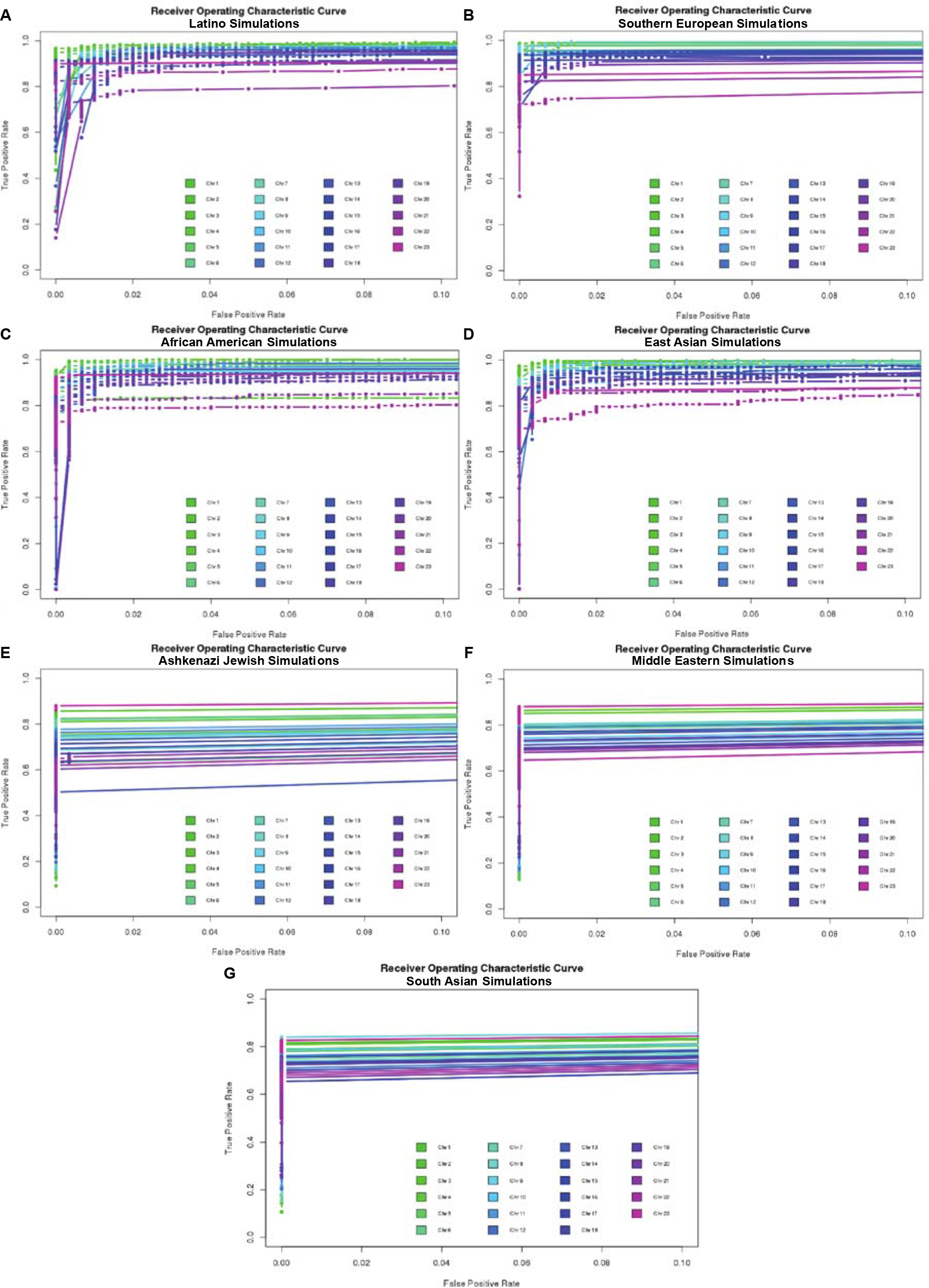
Receiver operating characteristic curves for 23 UPD classifiers (one per chromosome) trained on simulated individuals based on. A) Latino cohorts (TPR between 0.76 and 0.98 when FPR is fixed at 0.01), B) South European cohorts (TPR between 0.75 and 0.99 when FPR is fixed at 0.01), C) African American cohorts (TPR between 0.79 and 0.99 when FPR is fixed at 0.01), D) East Asian cohorts (TPR between 0.74 and 1 when FPR is fixed at 0.01), E) Ashkenazi Jewish cohorts (TPR between 0.51 and 0.899 when FPR is fixed at 0.01), F) Middle Eastern cohorts (TPR between 0.62 and 0.88 when FPR is fixed at 0.01), and G) South Asian cohorts (TPR between 0.63 and 0.84 when FPR is fixed at 0.01). Plots A-D show ROC curves with auROC > 0.9, which lead to successful classification in real data, whereas plots E-G show classifiers that perform relatively poorly on simulated testing data (auROC < 0.9). The cohorts in plots E-G, Ashkenazi Jewish, Middle Eastern and South Asian, are known to have practiced endogamy and so high levels of consanguinity may be confounding UPD detection for these classifiers.

**Supplementary Figure 4.**
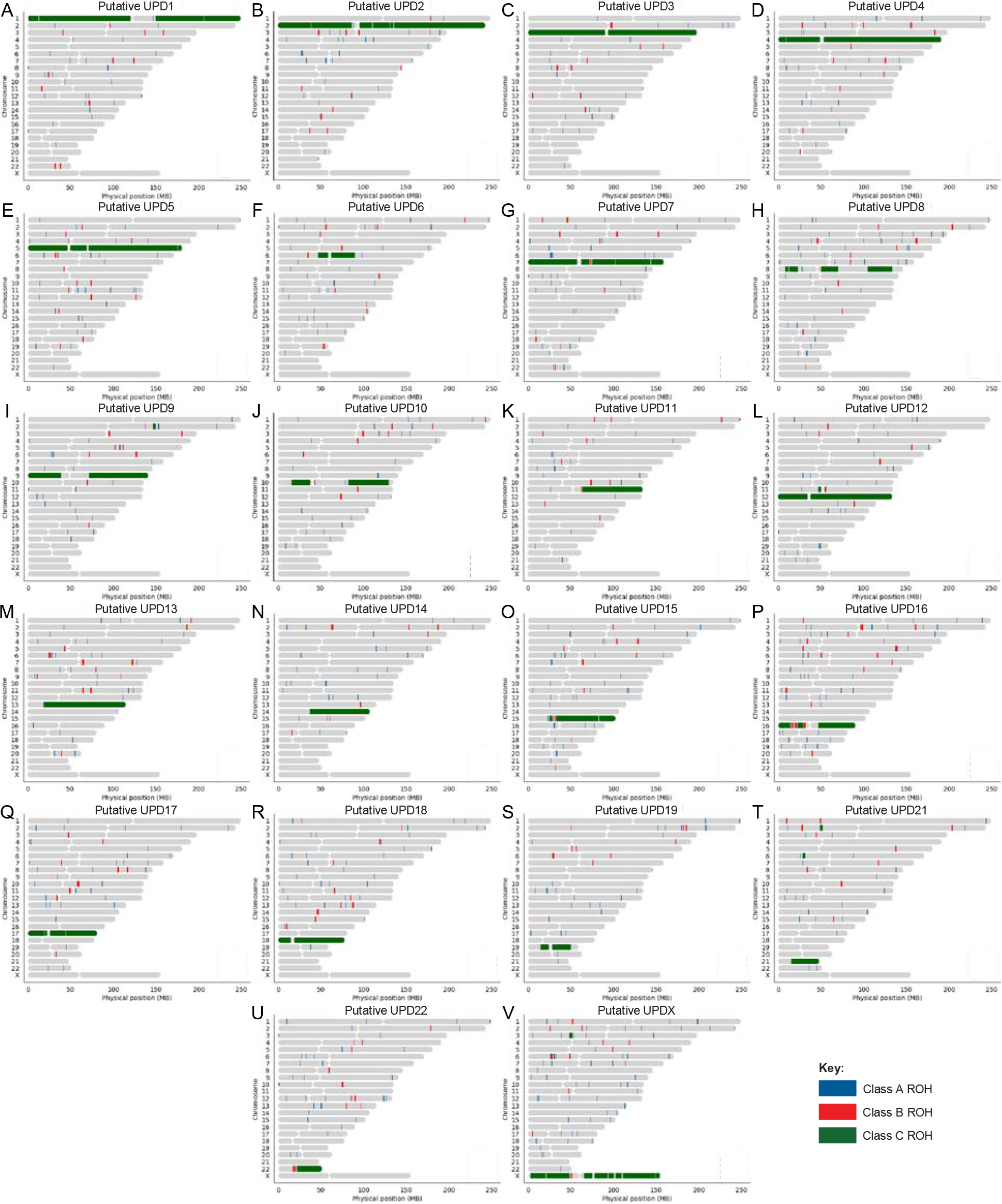
Ideograms of ROH for 22 ROH-based UPD cases we identified in the UK Biobank. We applied our ROH-based classifiers to 431,094 northern European individuals from the UK Biobank and identified 172 putative cases of UPD across 21 autosomes and the X chromosome (we did not classify any UPD cases on chromosome 20). This figure shows ideograms of ROH for 22 of the 172 putative cases, randomly drawn to illustrate ROH patterns of UPD cases of each chromosome for which we classify UPD; blue rectangles along the chromosomes represent Class A ROH, red rectangles represent Class B ROH, and green rectangles represent Class C ROH.

**Supplementary Figure 5.**
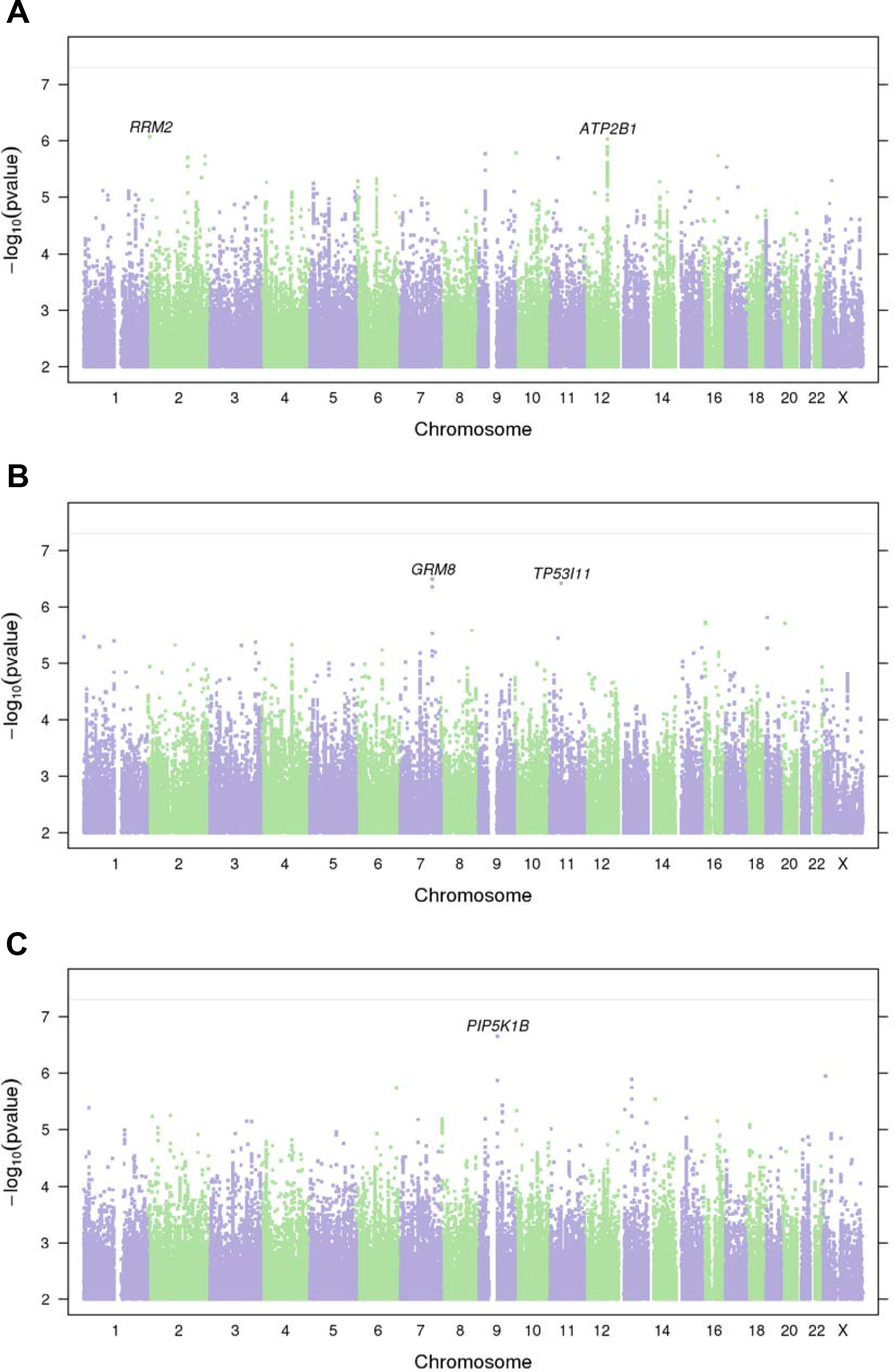
Manhattan plots of GWAS of UPD incidence, stratified by parental sex. (A) GWAS of all parents of UPD cases (both mothers and fathers), adjusted for sex; (B) GWAS of mothers of UPD cases only; and (C) GWAS of fathers of UPD cases only. There are no variants reaching genome-wide significance and the few hits reaching suggestive association level (*p*-value < 1×10^−6^) are likely false positives based on gene annotations.

**Supplementary Figure 6.**
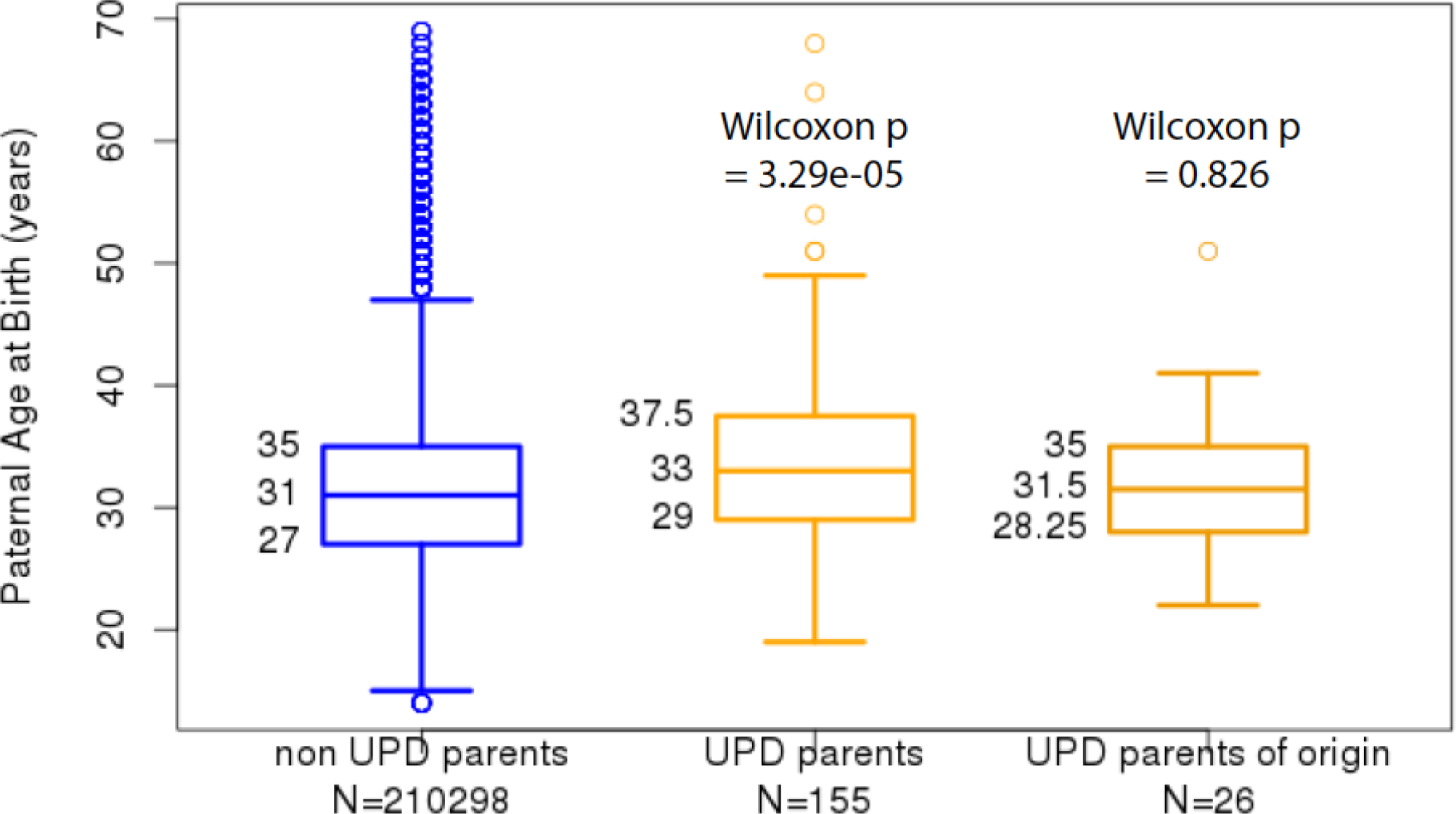
The age distribution of fathers of UPD true negatives (blue) and that of fathers who are parents of origin of UPD true positives (patUPD cases, yellow) in the 23andMe dataset. Fathers of UPD cases are significantly older than fathers of true negatives (Wilcoxon *p*-value = 3.29 x 10^−5^). However, we do not observe a significant difference in paternal age when we restrict analysis to fathers who are parents of origin of UPD children (Wilcoxon *p*-value = 0.286).

**Supplementary Figure 7.**
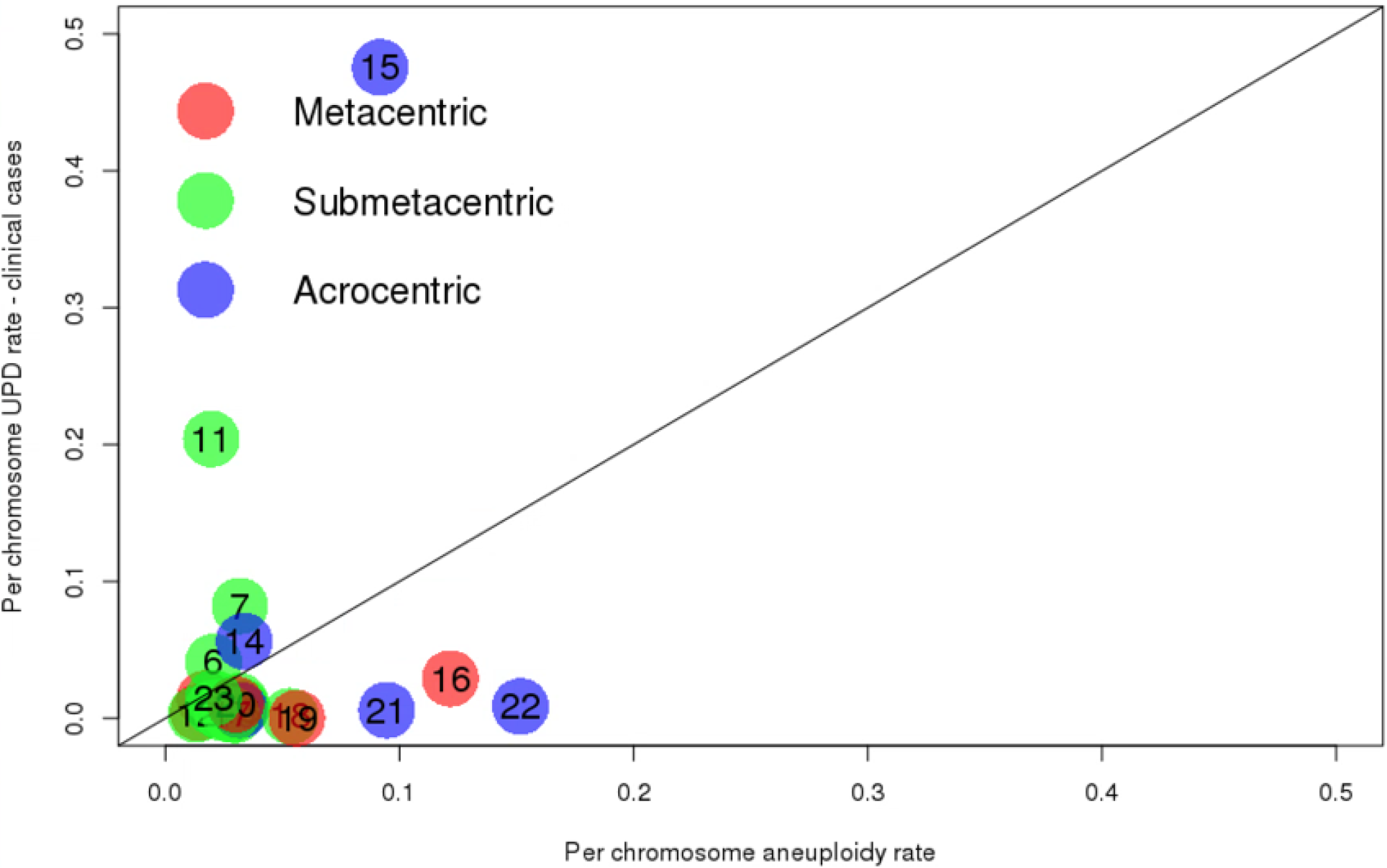
Correlation between per-chromosome UPD rate in published clinical cases (http://upd-tl.com/upd.html^1^) and per-chromosome aneuploidy rate in published pre-implantation embryo data^4^. Chromosomes are colored by centromeric type: metacentric chromosomes are shown in red, submetacentric chromosomes in green and acrocentric chromosomes in blue. In contrast to our results of correlation between per chromosome UPD rates in the 23andMe dataset and those from PGS data (Figure 4A), these two rates are not significantly correlated (Pearson’s correlation = 0.2; *p*-value = 0.34). This is expected since clinical cases are likely biased towards chromosomes causing serious medical phenotypes.

**Supplementary Figure 8.**
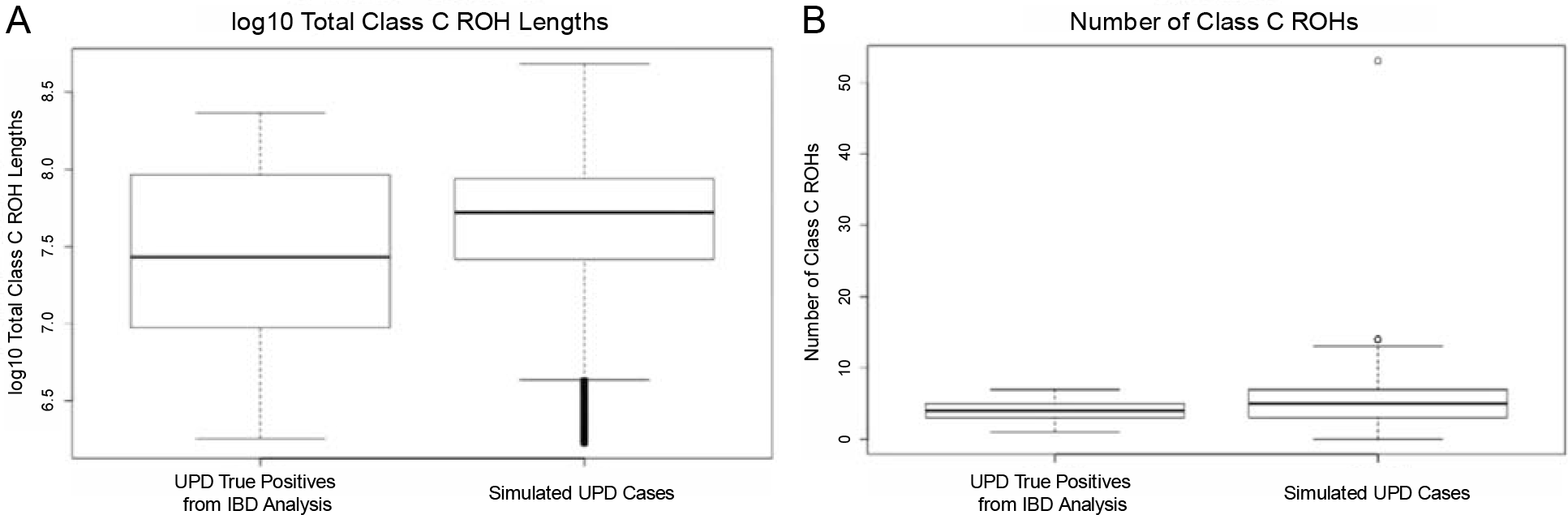
Boxplots comparing A) log10 total Class C ROH lengths of UPD true positives in the 23andMe dataset, which are identified using IBD analysis (left) and of simulated UPD cases (right) (t-test *p*-value = 0.94) and B) total number of Class C ROHs of UPD true positives (left) and of simulated UPD cases (right) (t-test *p*-value = 0.04). Because the number of Class C ROH differed significantly between real UPD cases and simulated UPD cases (panel B, t-test *p*-value < 0.05), we did not train on this variable in our classifiers and instead train on total Class C ROH length (panel A, t-test *p*-value > 0.05).

**Supplementary Figure 9.**
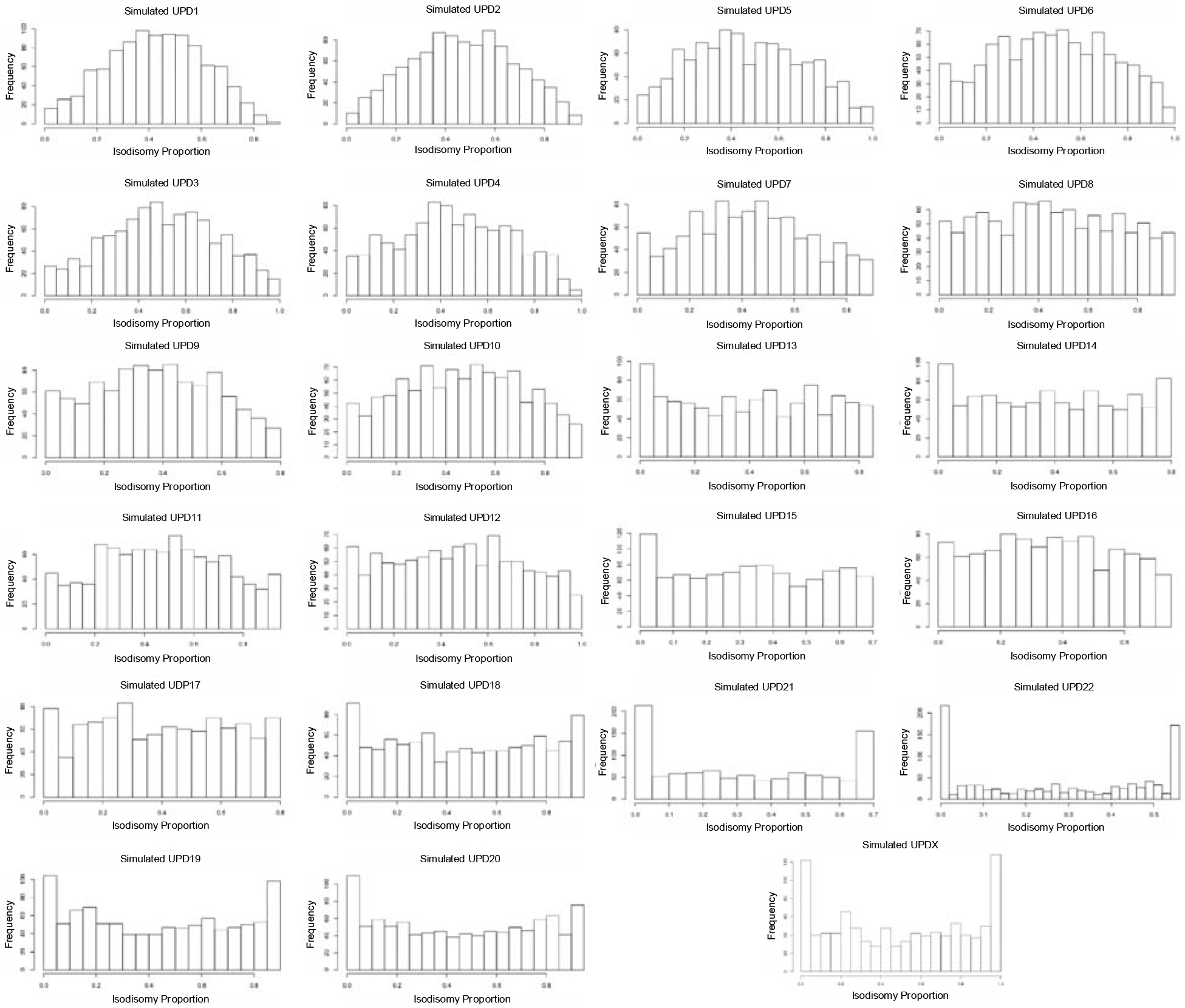
Distributions of isodisomy proportion in simulated UPD cases for each of 23 chromosomes. We simulated 1000 cases of UPD for each chromosome for each cohort based on genotype data from 23andMe. We see that, though the distribution of cases varies between chromosomes, every subtype of UPD (hetUPD, in which 0% isodisomy occurs; isoUPD, in which 100% isodisomy occurs; and partial isoUPD, in which an intermediate proportion between 0 and 100% isodisomy occurs) is produced on each chromosome by our simulation method.

**Supplementary Figure 10.**
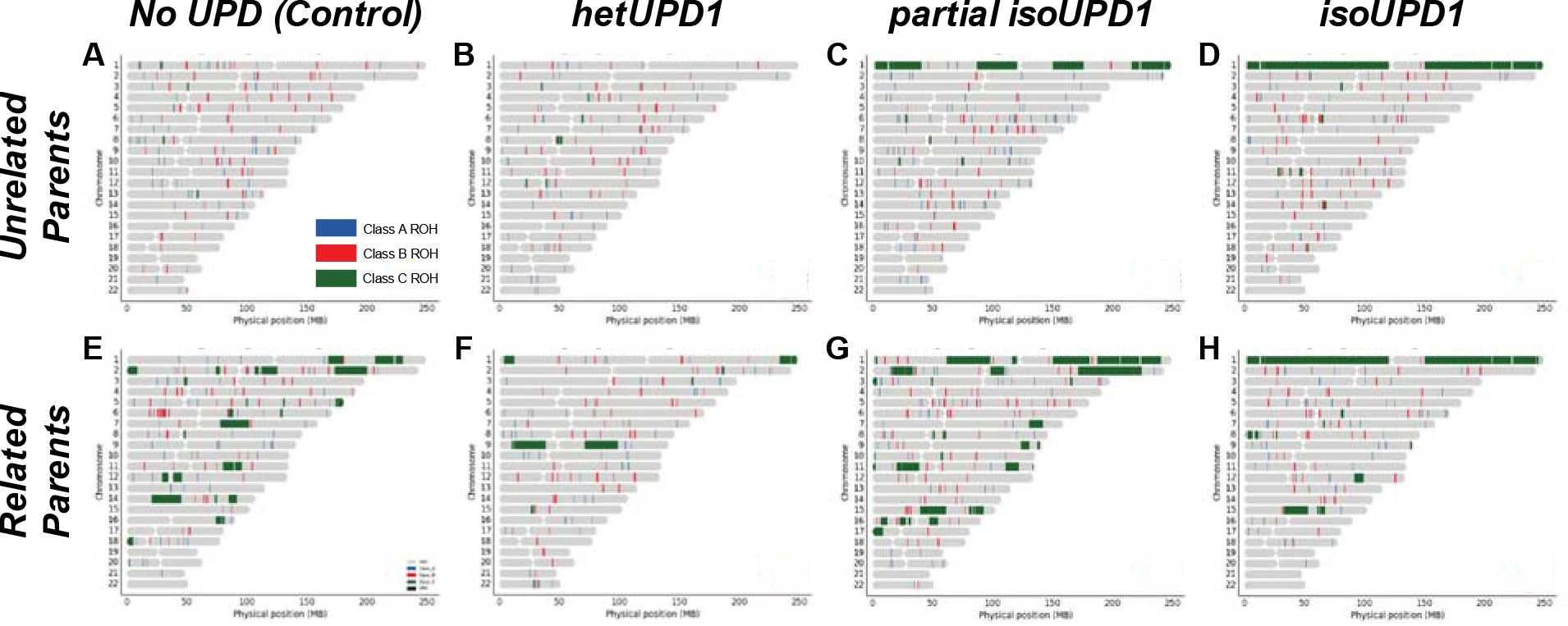
Example ideograms showing ROH locations (Class A ROH in blue, Class B ROH in red and Class C ROH in green). These were drawn from six simulated UPD cases and two simulated controls to illustrate the classification problem our ROH-based supervised classifiers faced. A) Ideogram of ROH in a control with unrelated parents, B) hetUPD of chromosome 1 with unrelated parents, C) partial isoUPD of chromosome 1 with unrelated parents, D) isoUPD of chromosome 1 with unrelated parents, E) control with related parents, F) hetUPD of chromosome 1 with related parents, G) partial isoUPD of chromosome 1 with related parents, and H) isoUPD of chromosome 1 with related parents. These figures show that long ROH can occur in partial isoUPD and isoUPD cases as well as individuals with related parents, and further illustrate that hetUPD cannot be identified based on ROH.

**Supplementary Figure 11.**
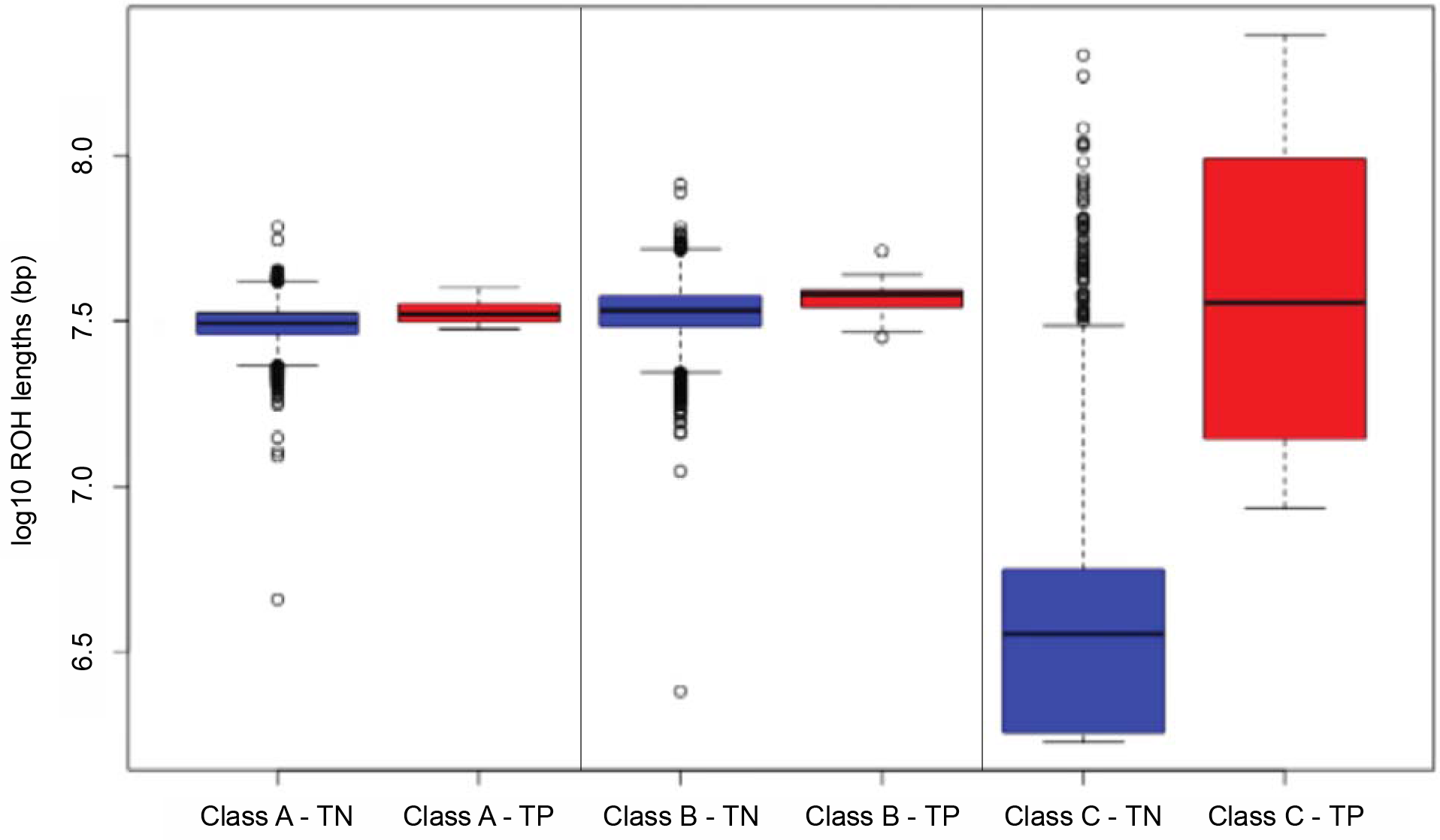
Boxplots comparing log10 ROH lengths (in bp) between UPD true negatives (TN, blue) and UPD true positives (TP, red) in the 23andMe dataset across the three length classes of ROH, from left to right: 1) Class A, the shortest ROH; 2) Class B, intermediate length ROH; 3) Class C, the longest ROH. ROH length class boundaries for each cohort are determined by GARLIC using gaussian mixture modeling (Supplementary Table 3). Only Class C ROH lengths differ significantly between UPD true negatives and true positives (t-test *p*-value < 0.05).

**Supplementary Figure 12.**
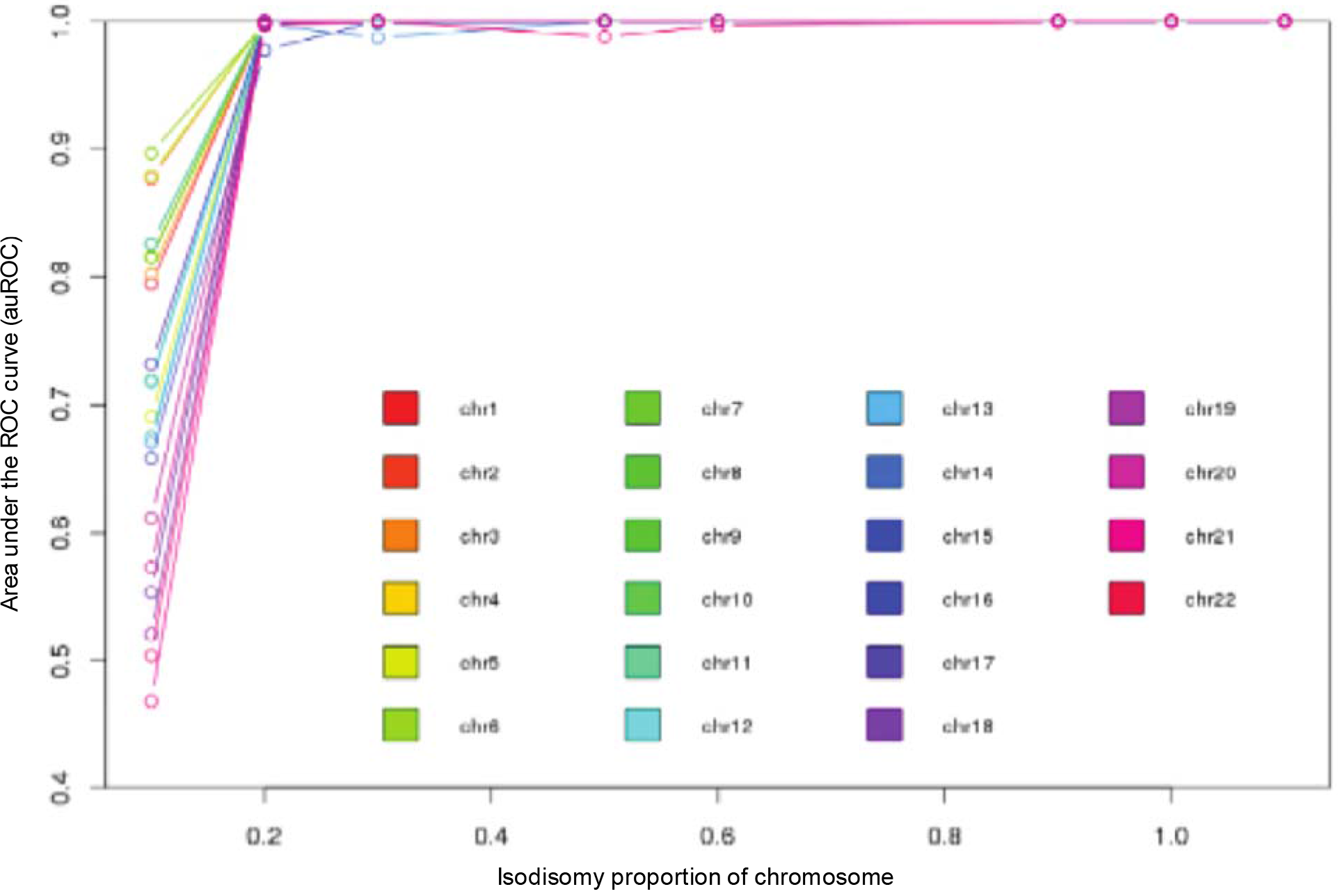
Area under the ROC curve (auROC) versus isodisomy proportion on the simulated chromosome. We found that auROC increases with isodisomy proportion on the simulated chromosome. Our classifiers perform best when isodisomy spans at least 20-50% of the chromosome.

**Supplementary Table 1.**
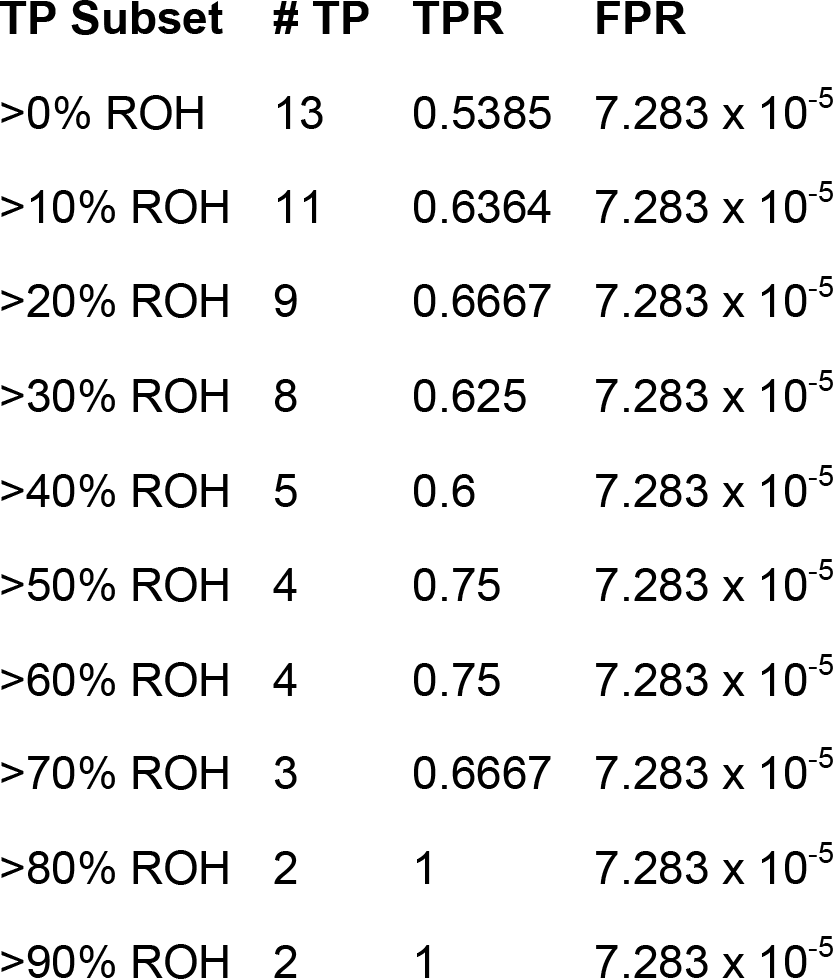
True Positive Rate (TPR) and False Positive Rate (FPR) when our classifier is applied to northern European true positives (TP) in the 23andMe dataset, identified from IBD analysis. The TP are subdivided by ROH percent across the UPD chromosome. We find that our classifiers have an TPR of 67% for cases with ROH spanning more than 20% of the chromosome while maintaining an FPR of 7 x 10^−5^.

**Supplementary Table 2.** Phenotypes significantly associated with UPD of chromosomes 1, 3, 6, 7, 8, 15, 16, 21 and 22 (*p*-value < 0.01). Traits with at least two cases (or two measurements for quantitative traits) are shown in bold. We tested for association between UPD of each of the chromosomes and 206 phenotypes across five categories (cognitive, personality, morphology, obesity and metabolic traits). Where possible, we also tested for association between matUPD and patUPD of each of the chromosomes separately. Effect sizes shown are odds ratios.

| **UPD type** | **Phenotype** | **Effect Size (95% CI)** | **P-values (Uncorrected)** |
| --- | --- | --- | --- |
| patUPD1 | Any Bariatric Surgery | 3.79 (1.39 6.20) | 0.0020 |
| UPD1 | Type 2 Diabetes | 3.19 (0.87 5.50) | 0.0070 |
| UPD3 | Hyperglycemia | 4.73 (2.32 7.13) | 0.0001 |
| UPD3 | Type 2 Diabetes | 4.04 (1.08 7.00) | 0.0076 |
| UPD6 | Hyperglycemia | 4.38 (1.60 7.15) | 0.0020 |
| UPD6 | Type 2 Diabetes | 3.90 (1.12 6.68) | 0.0060 |
| **UPD6** | **Weight** | **-2.02 (-3.38 -0.65)** | **0.0038** |
| **UPD6** | **Height** | **-1.99 (-3.40 -0.59)** | **0.0055** |
| UPD7 | Self-rated attractiveness | -4.20 (-7.08 -1.32) | 0.0042 |
| UPD7 | Life satisfaction | -3.75 (-6.53 -0.97) | 0.0081 |
| UPD8 | Birth weight | -5.31 (-8.21 -2.41) | 0.0003 |
| UPD8 | Autism | 3.73 (1.36 6.11) | 0.0021 |
| UPD8 | Memory Loss | 4.19 (1.37 7.02) | 0.0036 |
| UPD8 | Altitude Sickness | -4.11 (-6.90 -1.32) | 0.0038 |
| UPD8 | Likes to play with ideas | -8.36 (-14.59 -2.13) | 0.0086 |
| UPD8 | Autism Spectrum | 3.14 (0.78 5.49) | 0.0090 |
| matUPD15 | Autism Spectrum | 5.47 (2.42 8.51) | 0.0004 |
| UPD15 | Self-rated math ability | -3.53 (-6.09 -0.96) | 0.0071 |
| patUPD16 | Type 2 Diabetes | 7.72 (4.72 10.73) | 4.596 x 10<sup>-7</sup> |
| patUPD16 | Hyperglycemia | 6.05 (3.22 8.88) | 2.778 x 10<sup>-5</sup> |
| patUPD16 | High Cholesterol | 3.29 (0.84 5.74) | 0.0084 |
| matUPD21 | Feels left out of social activity | -3.52 (-5.93 -1.11) | 0.0042 |
| **UPD22** | **Autism Spectrum** | **3.61 (1.93 5.30)** | **2.557 x 10<sup>-5</sup>** |

**Supplementary Table 3.**
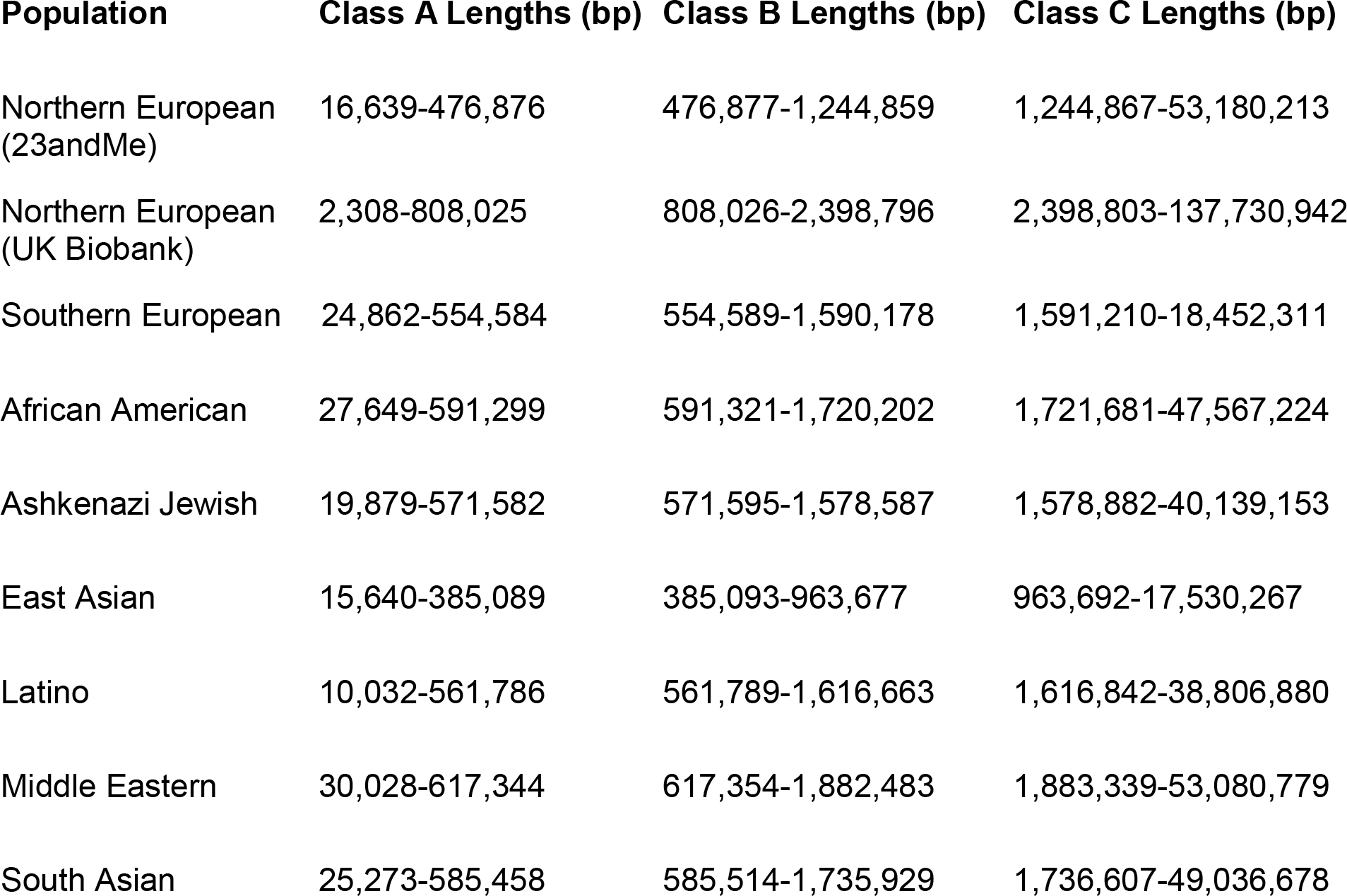
Boundaries (in base pairs) for each of three length classes of ROH across nine cohorts in the 23andMe database and the UK Biobank. Class boundaries were calculated using Gaussian mixture modeling on ROH length distributions in UPD true negatives from each of the eight cohorts in the 23andMe dataset and from all northern Europeans in the UK Biobank.

| Population | Class A Lengths (bp) | Class B Lengths (bp) | Class C Lengths (bp) |
| --- | --- | --- | --- |
| Northern European (23andMe) | 16,639-476,876 | 476,877-1,244,859 | 1,244,867-53,180,213 |
| Northern European (UK Biobank) | 2,308-808,025 | 808,026-2,398,796 | 2,398,803-137,730,942 |
| Southern European | 24,862-554,584 | 554,589-1,590,178 | 1,591,210-18,452,311 |
| African American | 27,649-591,299 | 591,321-1,720,202 | 1,721,681-47,567,224 |
| Ashkenazi Jewish | 19,879-571,582 | 571,595-1,578,587 | 1,578,882-40,139,153 |
| East Asian | 15,640-385,089 | 385,093-963,677 | 963,692-17,530,267 |
| Latino | 10,032-561,786 | 561,789-1,616,663 | 1,616,842-38,806,880 |
| Middle Eastern | 30,028-617,344 | 617,354-1,882,483 | 1,883,339-53,080,779 |
| South Asian | 25,273-585,458 | 585,514-1,735,929 | 1,736,607-49,036,678 |

## Supplementary Methods

### Genome-Wide Association Study

For the genome-wide association study (GWAS) of UPD, we restricted participants to a set of individuals who have European ancestry determined through an analysis of local ancestry described in the Methods section. A maximal set of unrelated individuals was chosen for each analysis using a segmental identity-by-descent (IBD) estimation algorithm^5^. Individuals were defined as related if they shared more than 700 cM IBD, including regions where the two individuals share either one or both genomic segments IBD. This level of relatedness (roughly 20% of the genome) corresponds approximately to the minimal expected sharing between first cousins in an outbred population. When selecting individuals for case/control phenotype analyses, the selection process is designed to maximize case sample size by preferentially retaining cases over controls. Specifically, if both an individual case and an individual control are found to be related, then the case is retained in the analysis.

Imputation panels created by combining multiple smaller panels have been shown to give better imputation performance than the individual constituent panels alone^6^. To that end, we combined the May 2015 release of the 1000 Genomes Phase 3 haplotypes^7^ with the UK10K imputation reference panel^8^ to create a single unified imputation reference panel. To do this, multiallelic sites with *N* alternate alleles were split into *N* separate biallelic sites. We then removed any site whose minor allele appeared in only one sample. For each chromosome, we used Minimac3^9^ to impute the reference panels against each other, reporting the best-guess genotype at each site. This gave us calls for all samples over a single unified set of variants. We then joined these together to get, for each chromosome, a single file with phased calls at every site for 6,285 samples. Throughout, we treated structural variants and small indels in the same way as SNPs.

In preparation for imputation we split each chromosome of the reference panel into chunks of no more than 300,000 variants, with 10,000 variants overlapping on each side. We used a single batch of 10,000 individuals to estimate Minimac3 imputation model parameters for each chunk. To generate phased participant data for the v1 to v4 platforms, we used an internally-developed tool at 23andMe, Inc., Finch, which implements the Beagle graph-based haplotype phasing algorithm^10^, modified to separate the haplotype graph construction and phasing steps. Finch extends the Beagle model to accommodate genotyping error and recombination, in order to handle cases where there are no consistent paths through the haplotype graph for the individual being phased. We constructed haplotype graphs for all participants from a representative sample of genotyped individuals, and then performed out-of-sample phasing of all genotyped individuals against the appropriate graph. For the X chromosome, we built separate haplotype graphs for the non-pseudoautosomal region and each pseudoautosomal region, and these regions were phased separately. For the 23andMe participants genotyped on the Illumina Global Screening Array-based platform (see “Genotyping and Quality Control” section), we used a similar approach, but using a new phasing algorithm, Eagle2^11^.

We imputed phased participant data against the merged reference panel using Minimac3, treating males as homozygous pseudo-diploids for the non-pseudoautosomal region. We computed association test results for the genotyped and the imputed SNPs. We assessed association by logistic regression assuming additive allelic effects. For tests using imputed data, we used the imputed dosages rather than best-guess genotypes. We also included covariates for age, gender, the top five principal components to account for residual population structure, and indicators for genotype platforms to account for genotype batch effects. The association test *p*-value we report was computed using a likelihood ratio test, which in our experience is better behaved than a Wald test on the regression coefficient. For quantitative traits, association tests were performed by linear regression. Results for the X chromosome were computed similarly, with male genotypes coded as if they were homozygous diploid for the observed allele.

Principal component analysis was performed independently for each ancestry, using ~65,000 high quality genotyped variants present in all five genotyping platforms. It was computed on a subset of one million participants randomly sampled across all the genotyping platforms. PC scores for participants not included in the analysis were obtained by projection, combining the eigenvectors of the analysis and the SNP weights.

