## Supplementary File 1 for "Characterization of prevalence and health consequences of uniparental disomy in four million individuals from the general population"

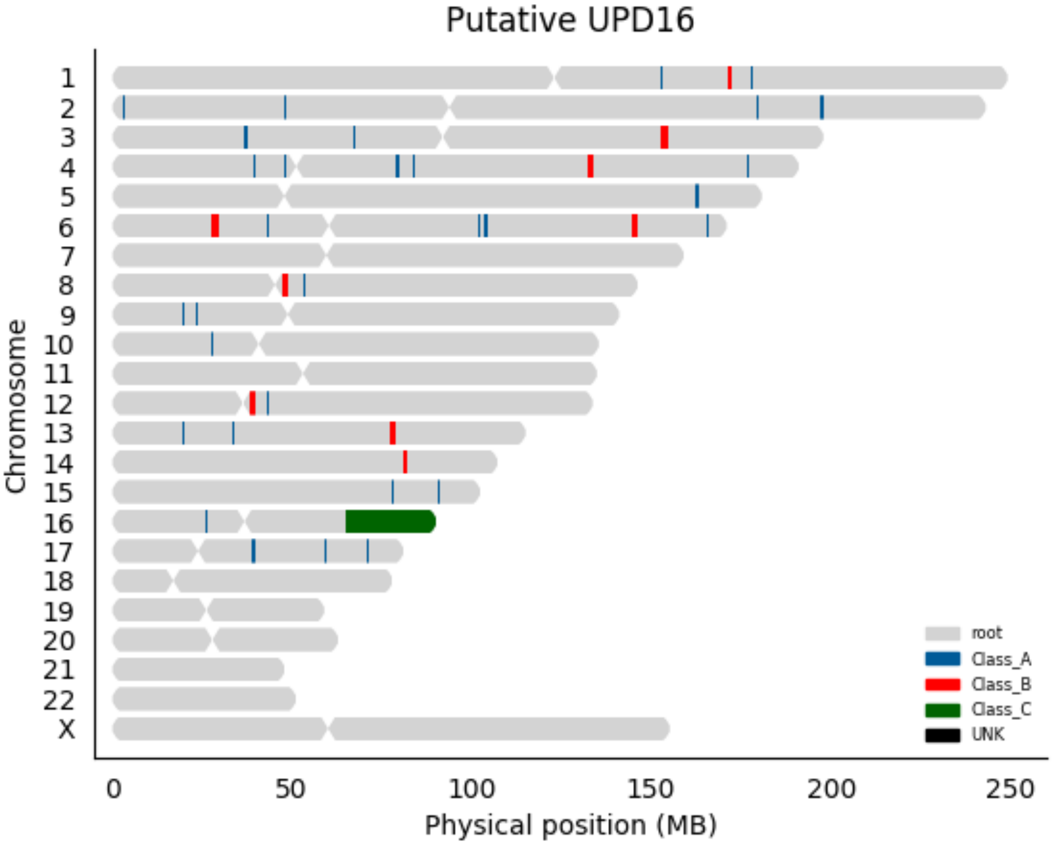

Putative UPD16
Chromosome
X
0
50
100
150
200
250
Physical position (MB)
root
Class_A
Class_B
Class_C
UNK

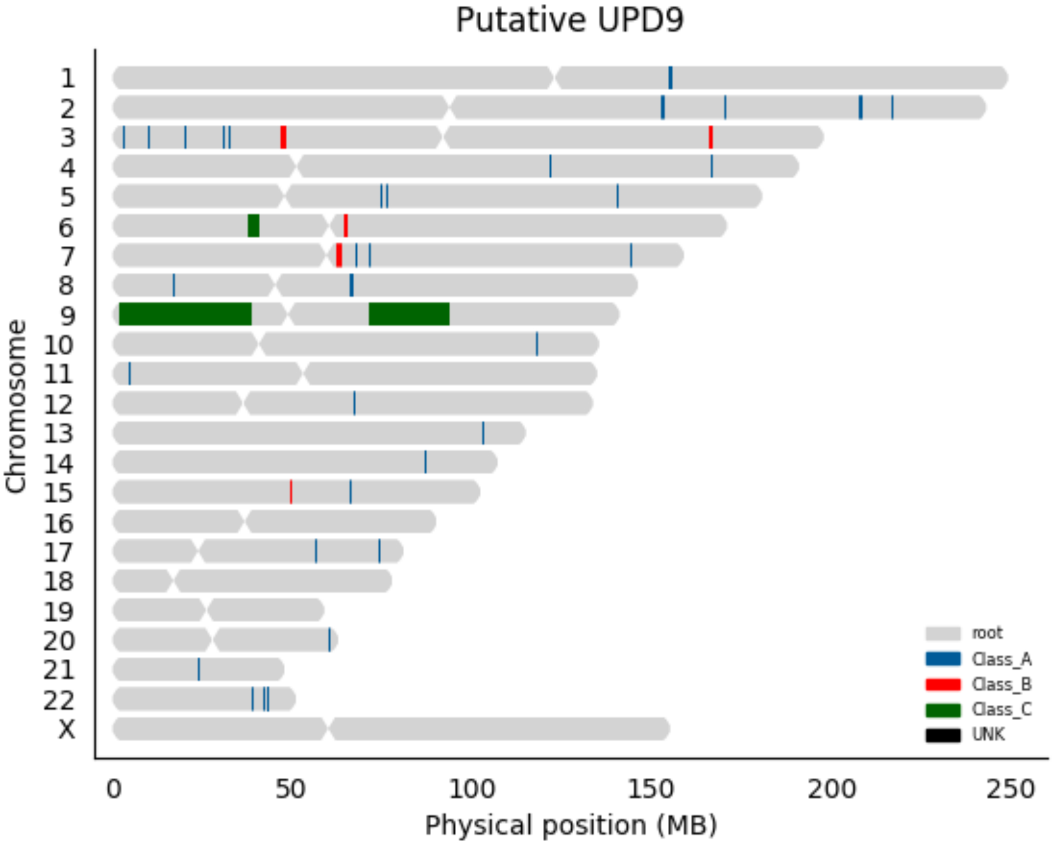

Putative UPD9
Chromosome
X
0
50
100
150
200
250
Physical position (MB)
root
Class_A
Class_B
Class_C
UNK

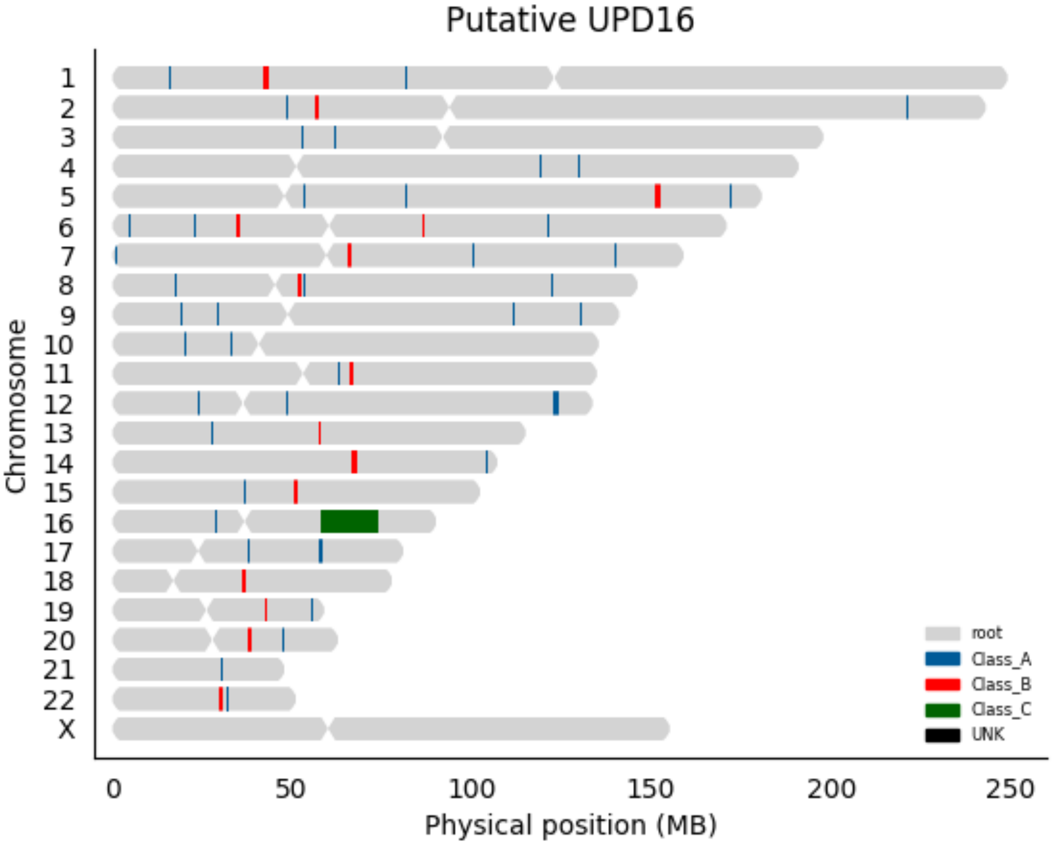

Putative UPD16
Chromosome
X
0
50
100
150
200
250
Physical position (MB)
root
Class_A
Class_B
Class_C
UNK

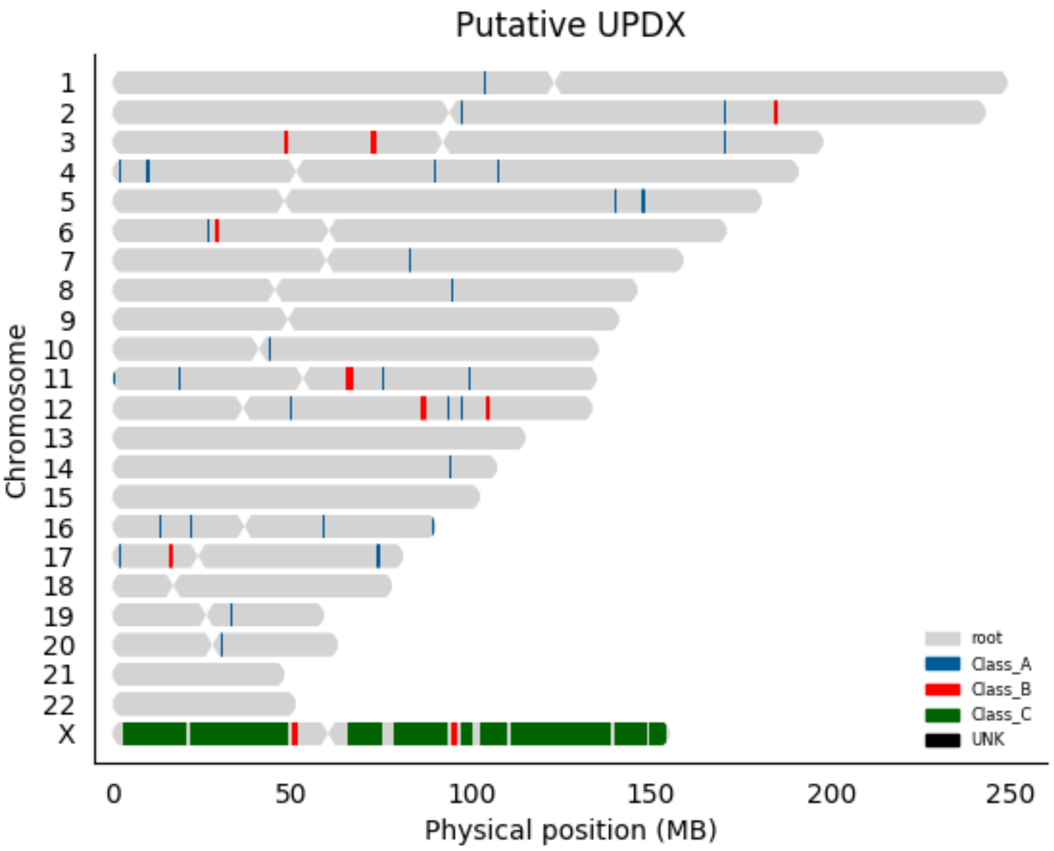
Putative UPDX
Chromosome
X
0
50
100
150
200
250
Physical position (MB)
root
Class_A
Class_B
Class_C
UNK

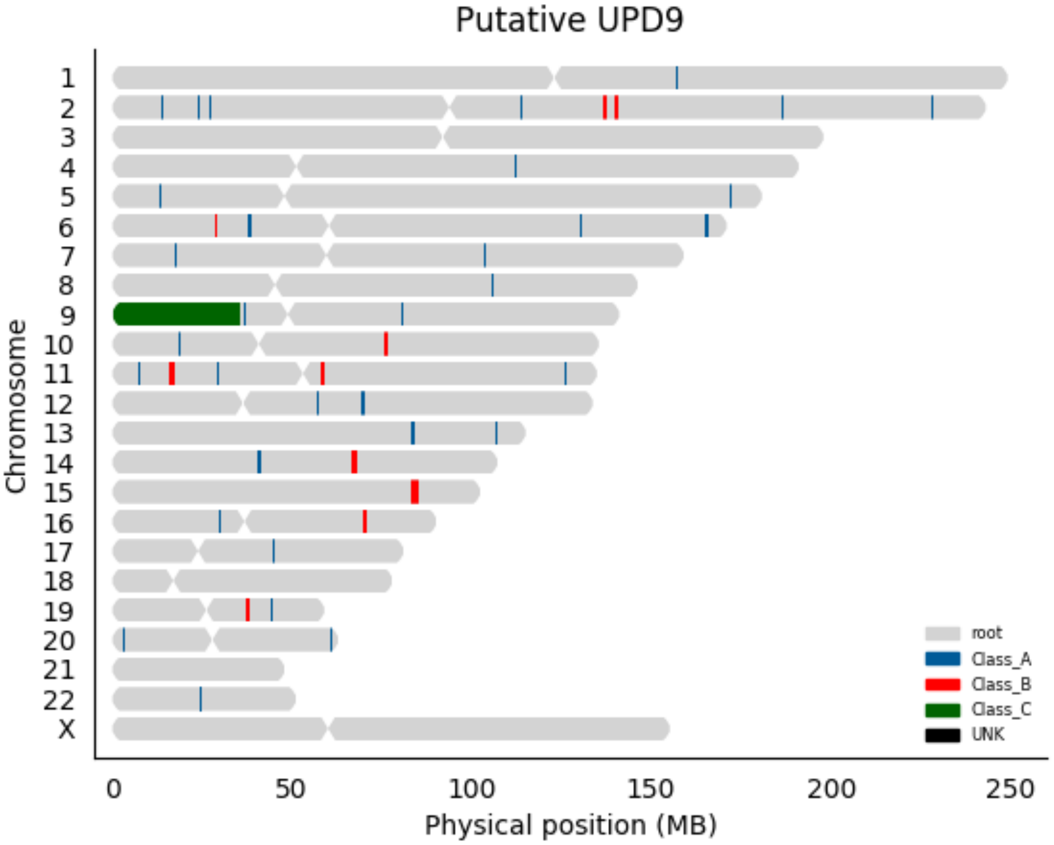

Putative UPD9
Chromosome
X
0
50
100
150
200
250
Physical position (MB)
root
Class_A
Class_B
Class_C
UNK

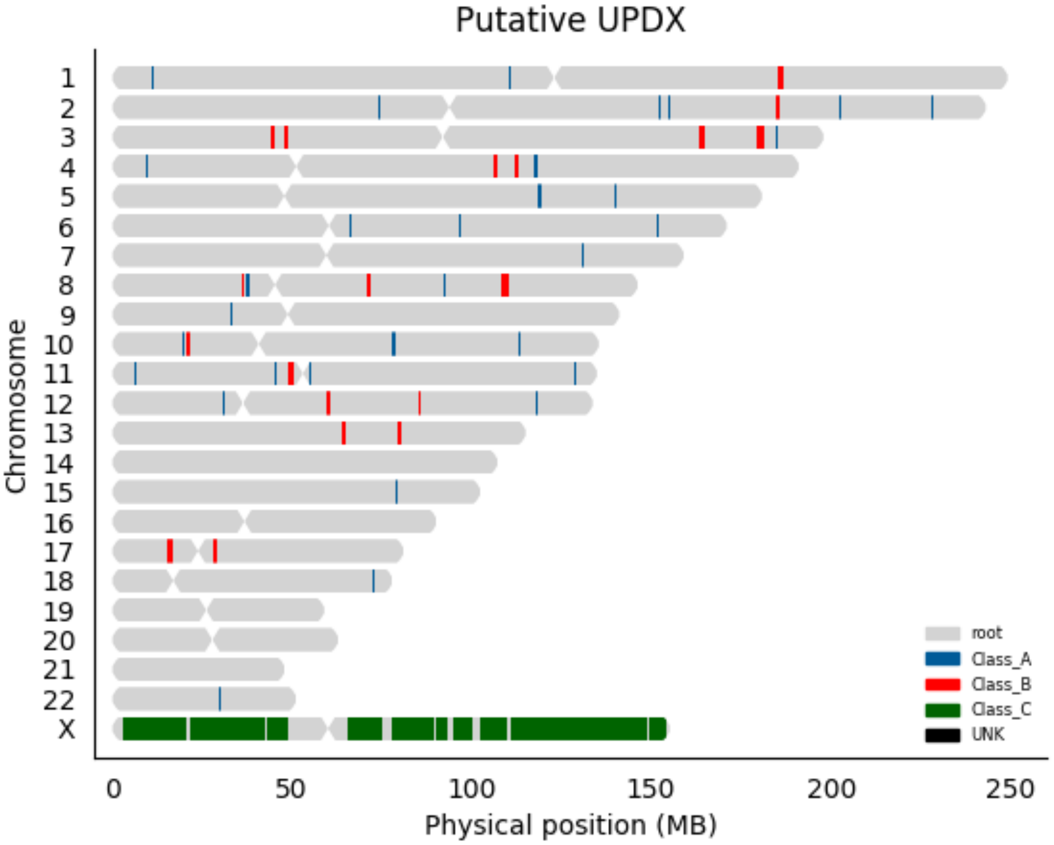

Putative UPDX
Chromosome
X
0
50
100
150
200
250
Physical position (MB)
root
Class_A
Class_B
Class_C
UNK

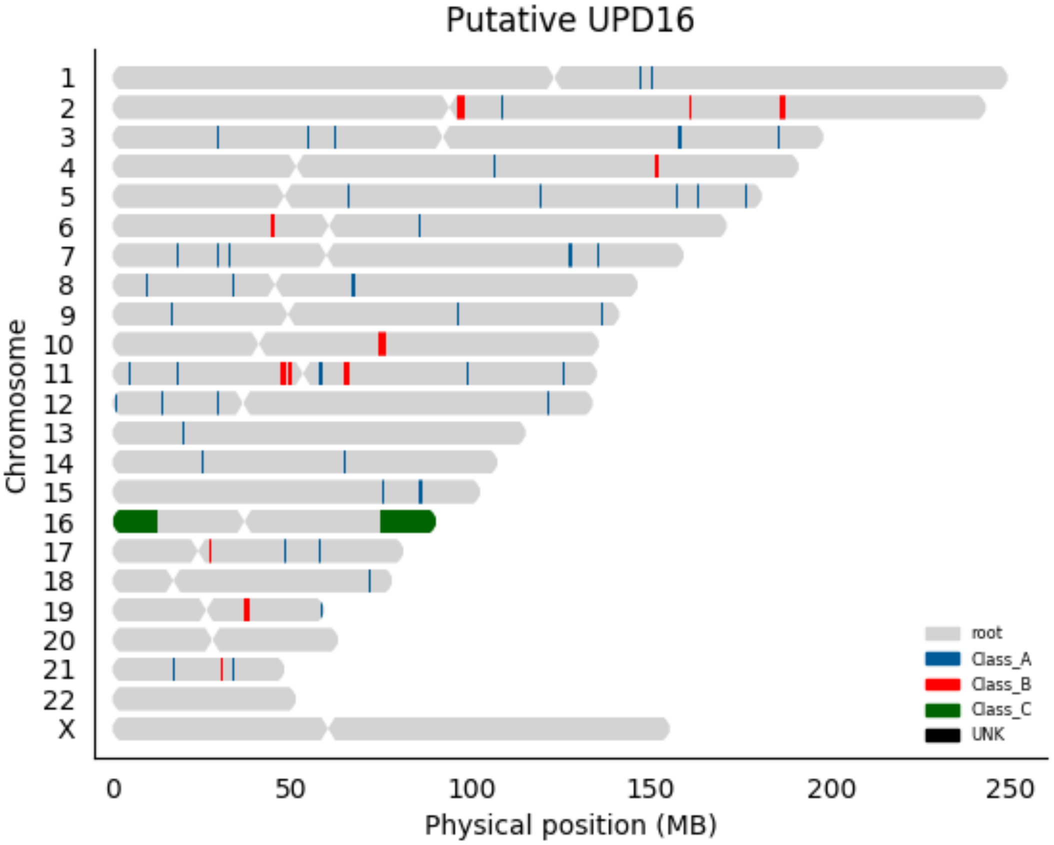

Putative UPD16
Chromosome
X
0
50
100
150
200
250
Physical position (MB)
root
Class_A
Class_B
Class_C
UNK

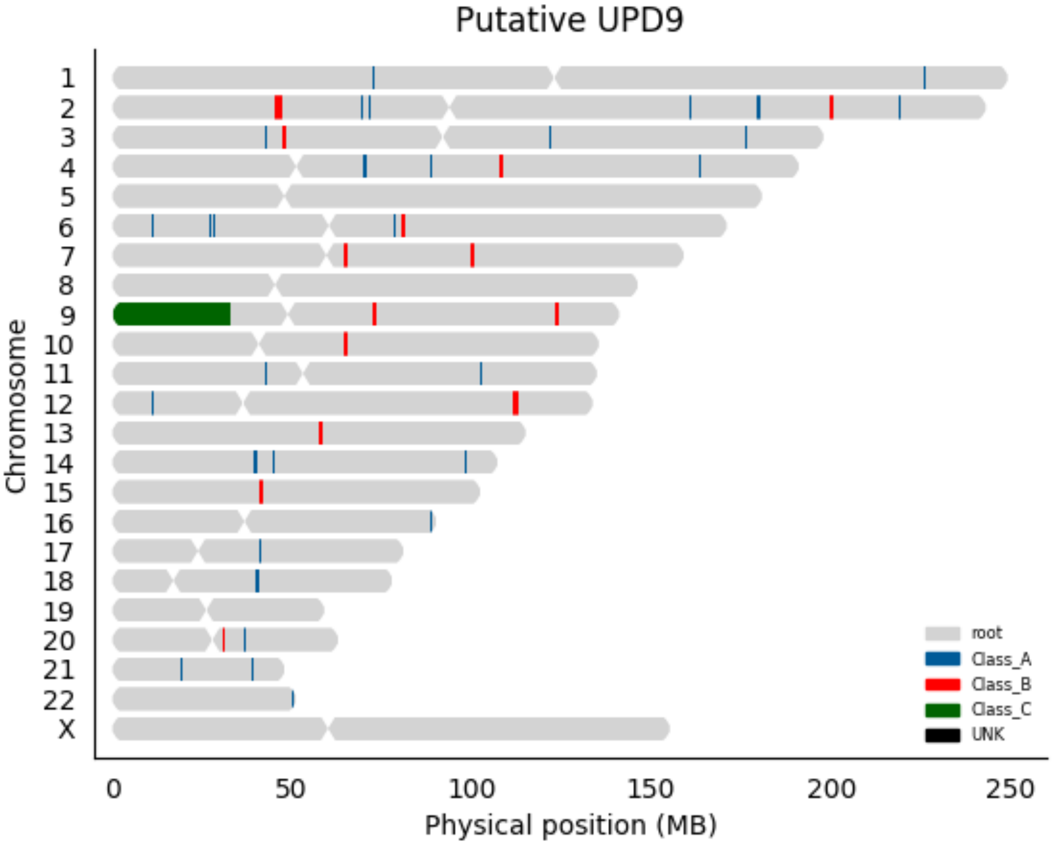

Putative UPD9
Chromosome
X
0
50
100
150
200
250
Physical position (MB)
root
Class_A
Class_B
Class_C
UNK

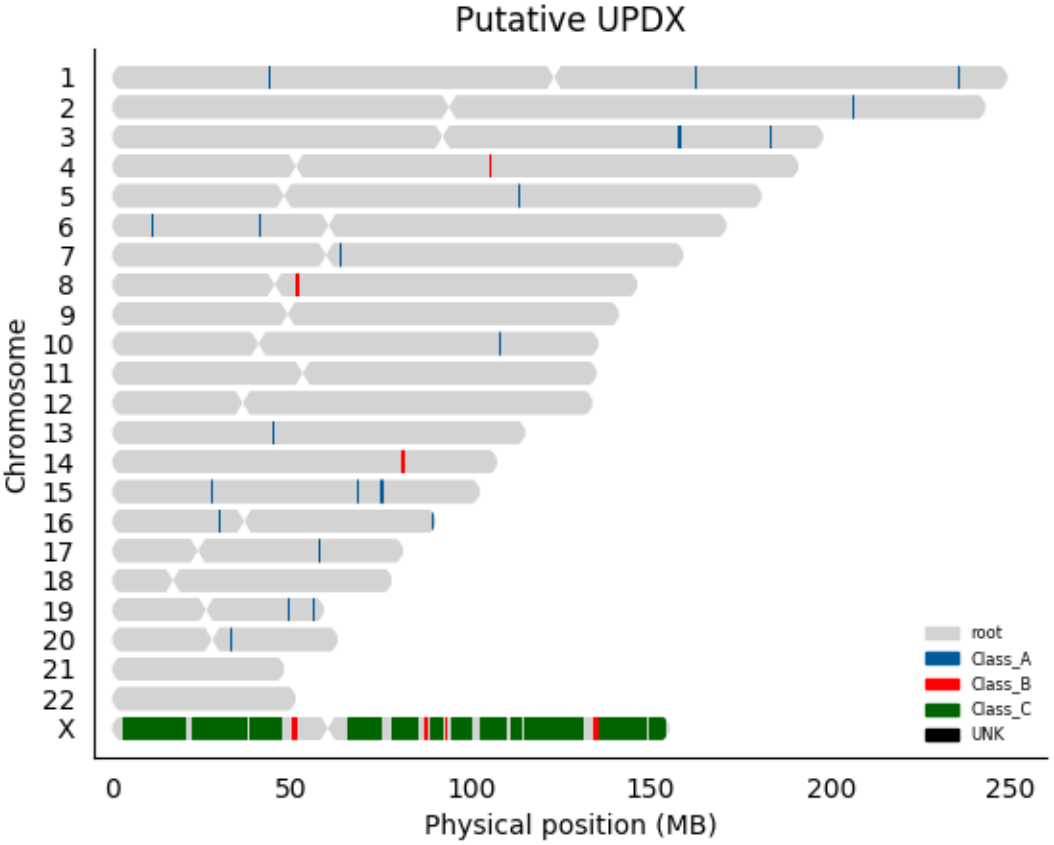

Putative UPDX
Chromosome
X
0
50
100
150
200
250
Physical position (MB)
root
Class_A
Class_B
Class_C
UNK

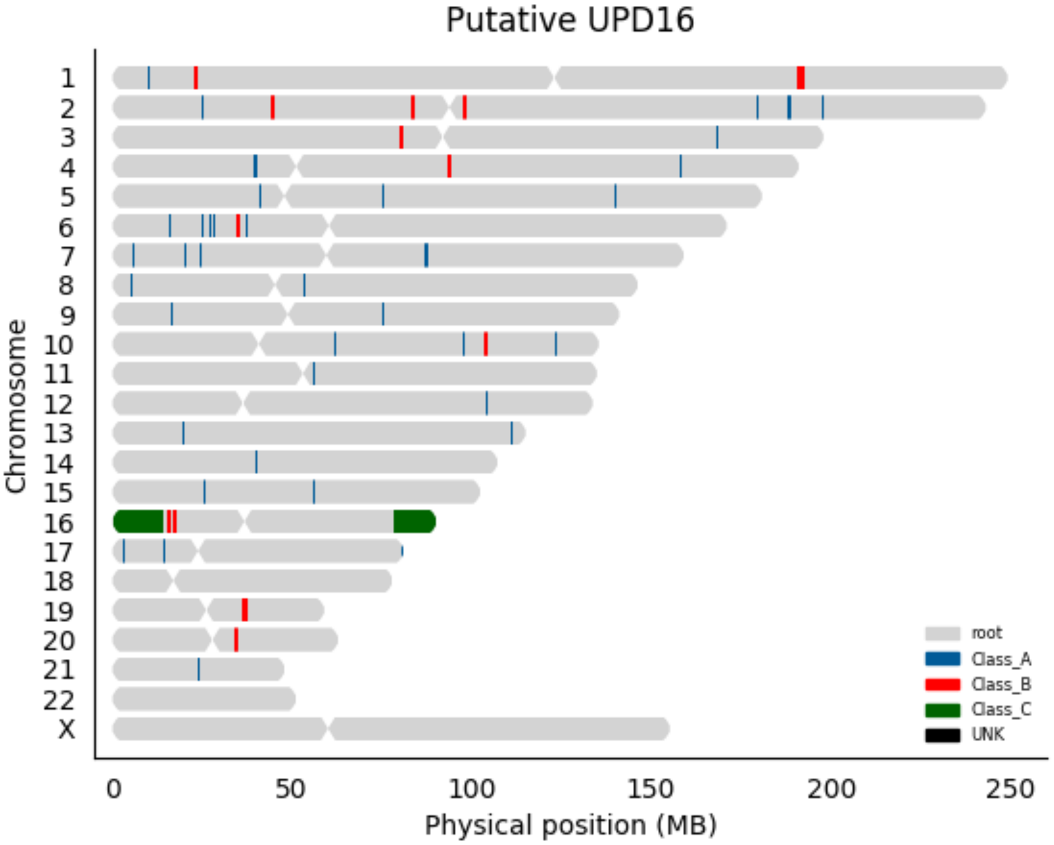

Putative UPD16
Chromosome
X
0
50
100
150
200
250
Physical position (MB)
root
Class_A
Class_B
Class_C
UNK

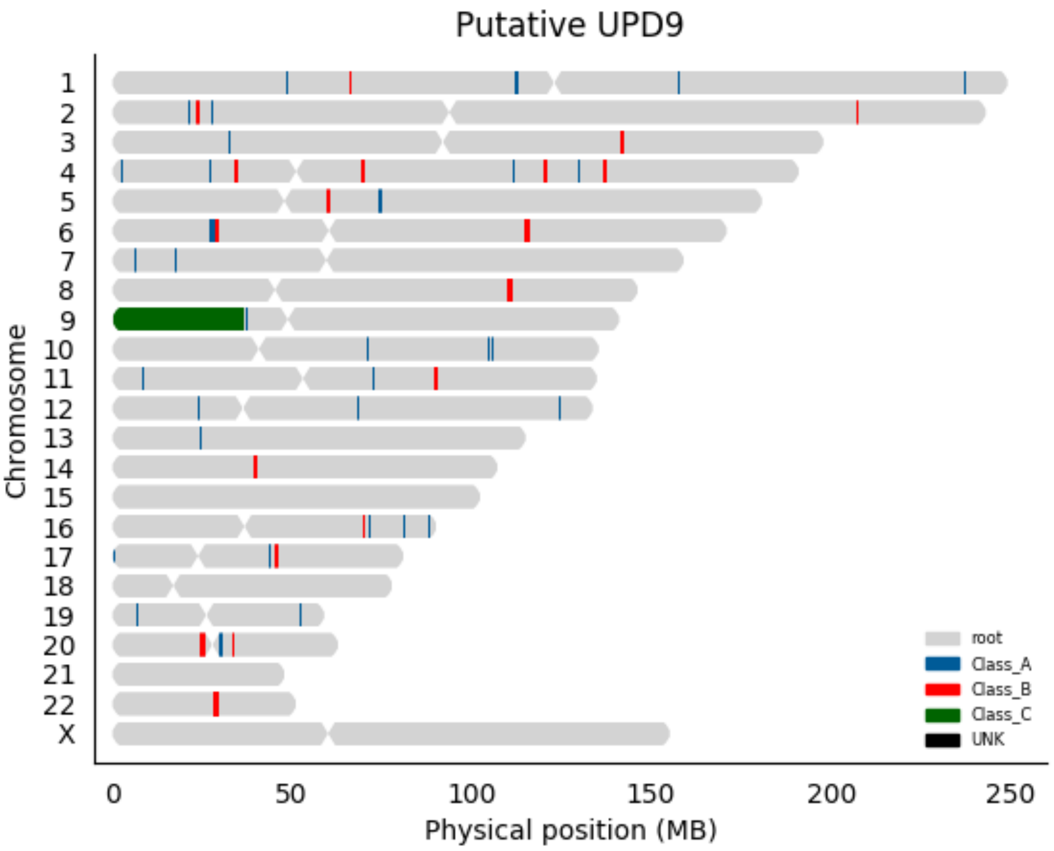

Putative UPD9
Chromosome
X
0
50
100
150
200
250
Physical position (MB)
root
Class_A
Class_B
Class_C
UNK

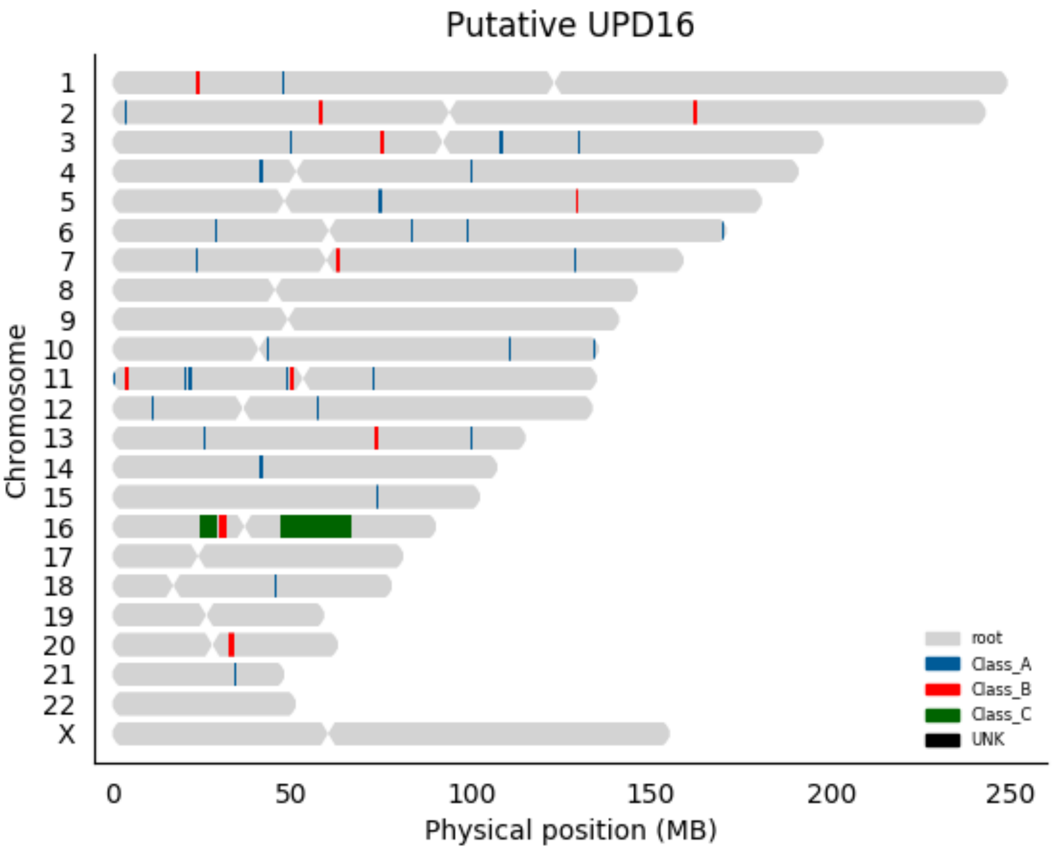

Putative UPD16
Chromosome
X
0
50
100
150
200
250
Physical position (MB)
root
Class_A
Class_B
Class_C
UNK

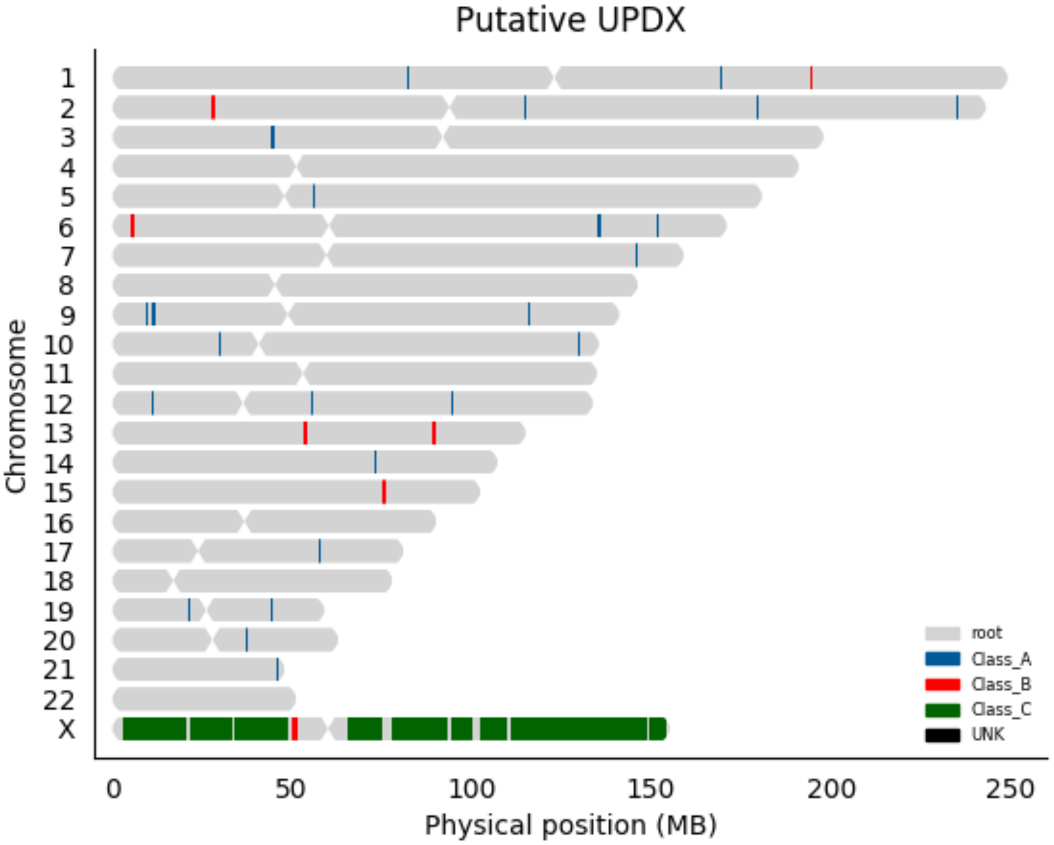

Putative UPDX
Chromosome
X
0
50
100
150
200
250
Physical position (MB)
root
Class_A
Class_B
Class_C
UNK

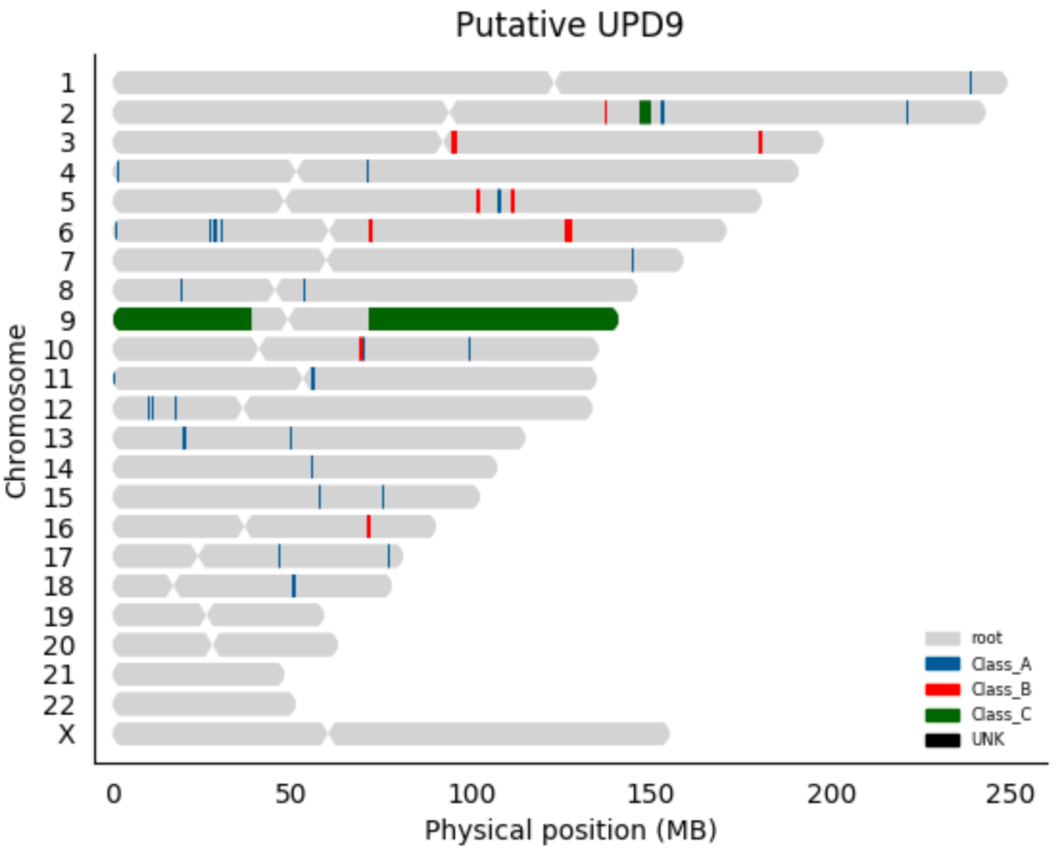
Putative UPD9
Chromosome
X
0
50
100
150
200
250
Physical position (MB)
root
Class_A
Class_B
Class_C
UNK

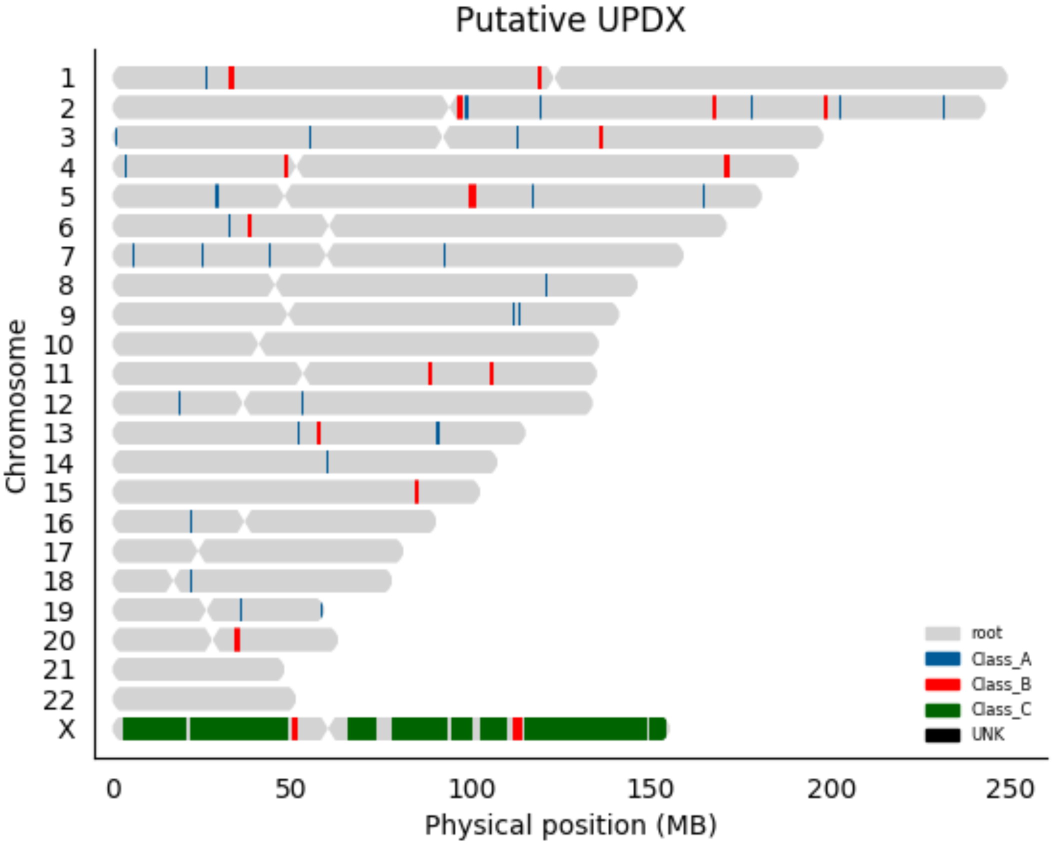
Putative UPDX
Chromosome
X
0
50
100
150
200
250
Physical position (MB)
root
Class_A
Class_B
Class_C
UNK

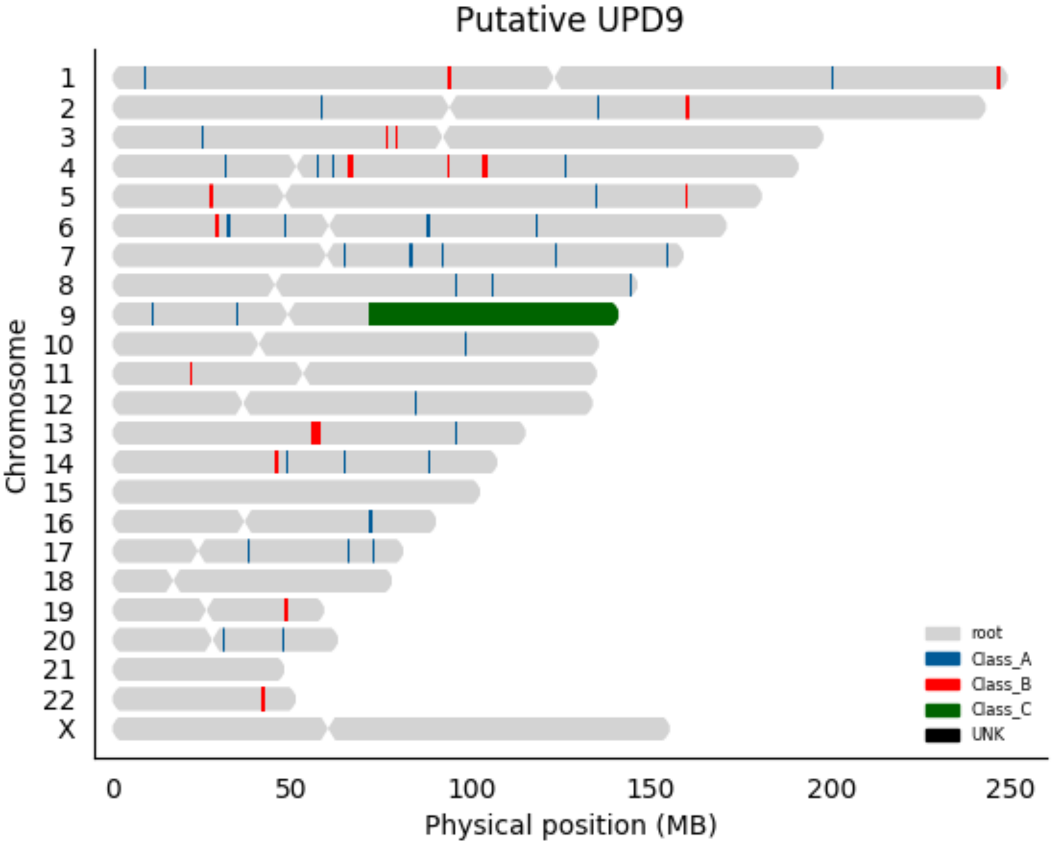

Putative UPD9
Chromosome
X
0
50
100
150
200
250
Physical position (MB)
root
Class_A
Class_B
Class_C
UNK

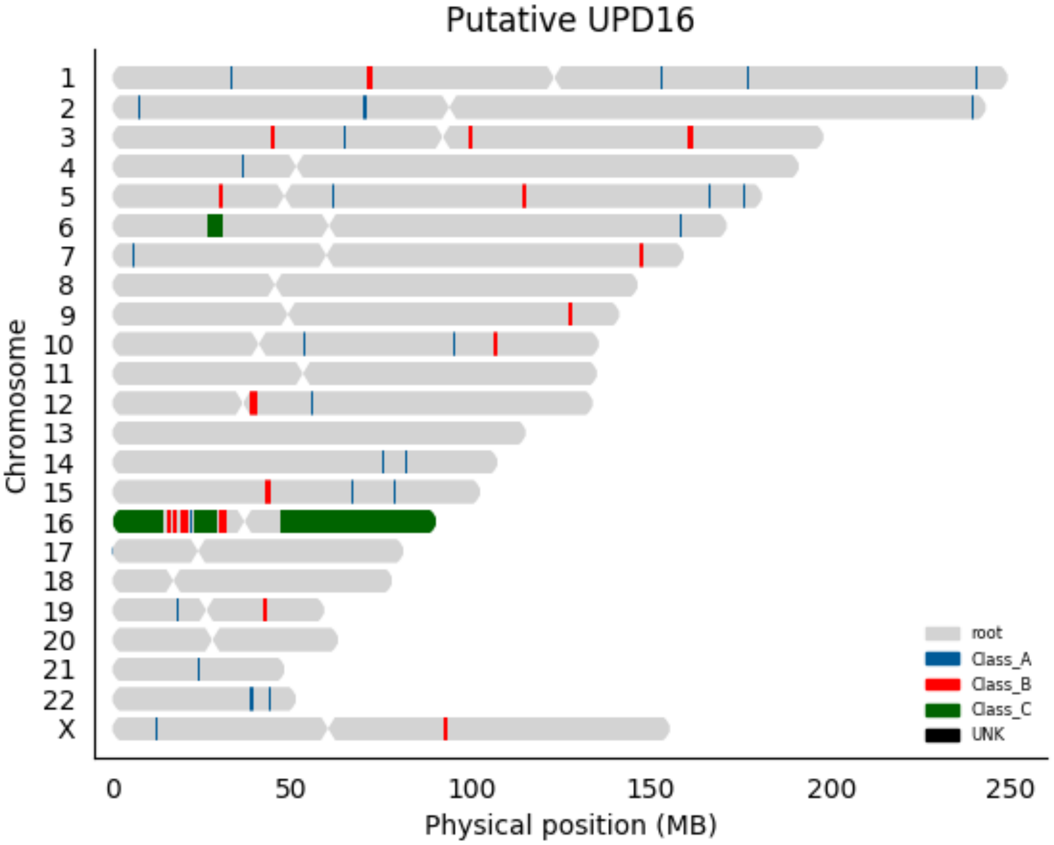

Putative UPD16
Chromosome
X
0
50
100
150
200
250
Physical position (MB)
root
Class_A
Class_B
Class_C
UNK

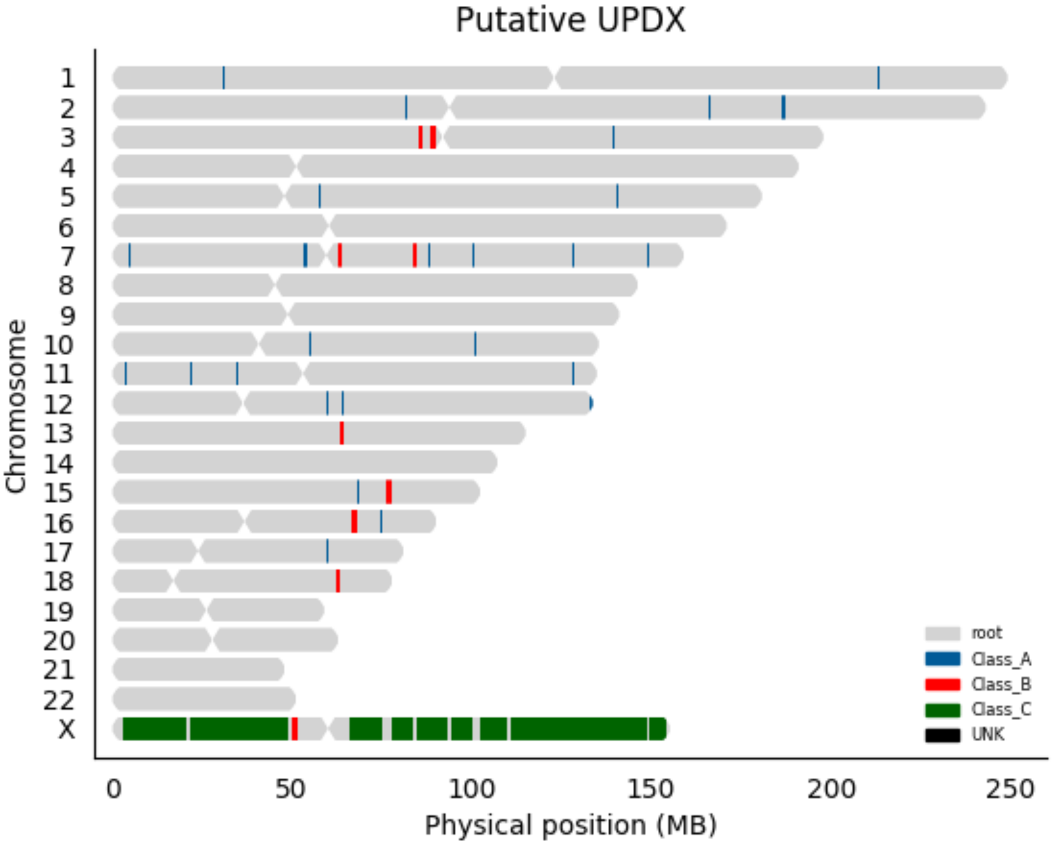
Putative UPDX
Chromosome
X
0
50
100
150
200
250
Physical position (MB)
root
Class_A
Class_B
Class_C
UNK

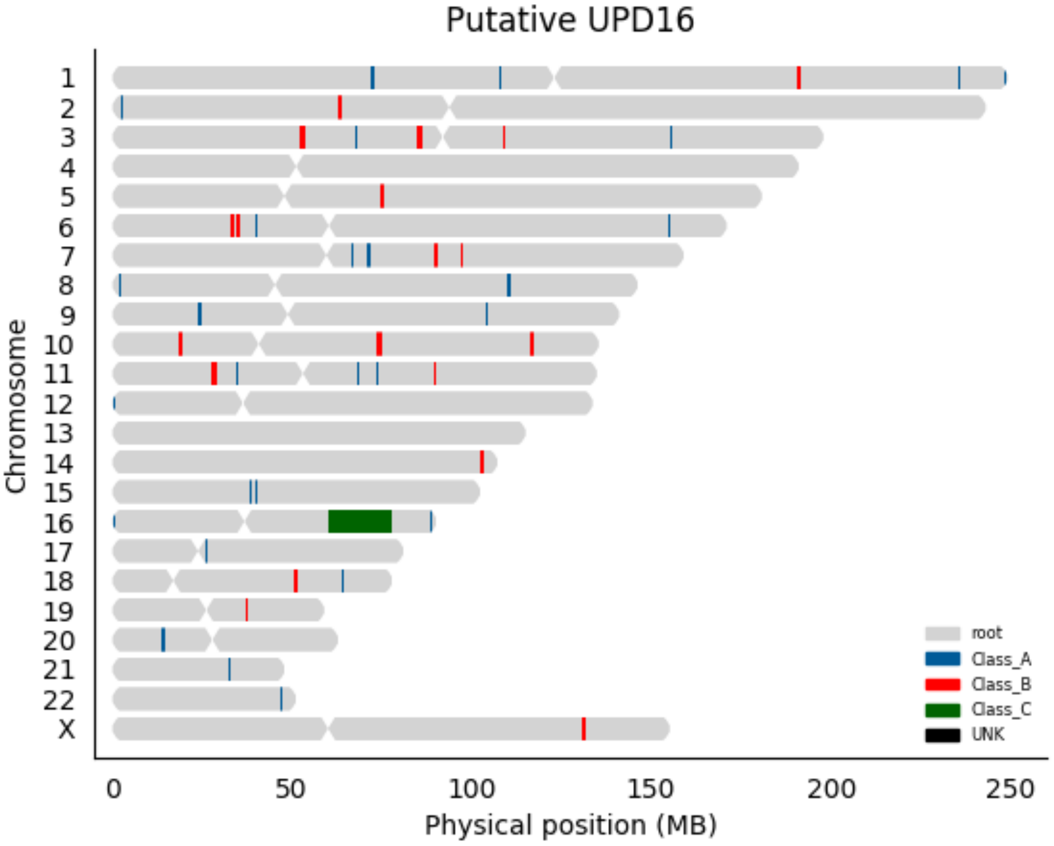

Putative UPD16
Chromosome
X
0
50
100
150
200
250
Physical position (MB)
root
Class_A
Class_B
Class_C
UNK

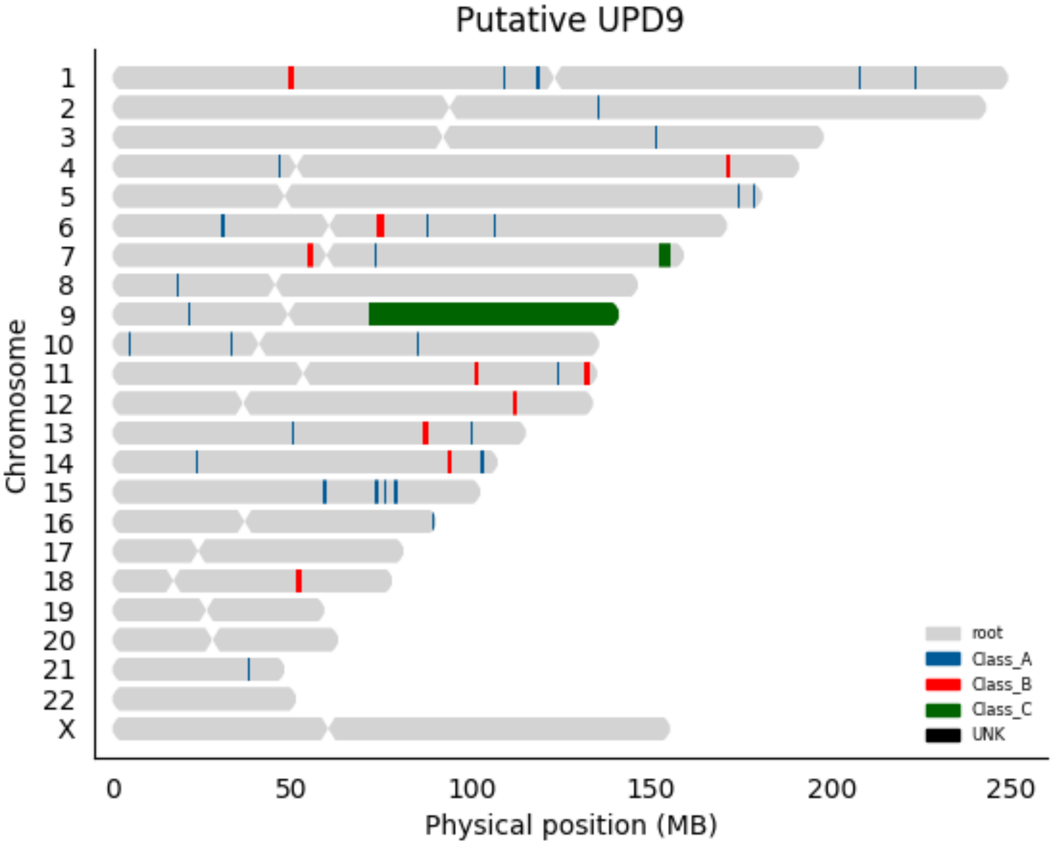
Putative UPD9
Chromosome
X
0
50
100
150
200
250
Physical position (MB)
root
Class_A
Class_B
Class_C
UNK

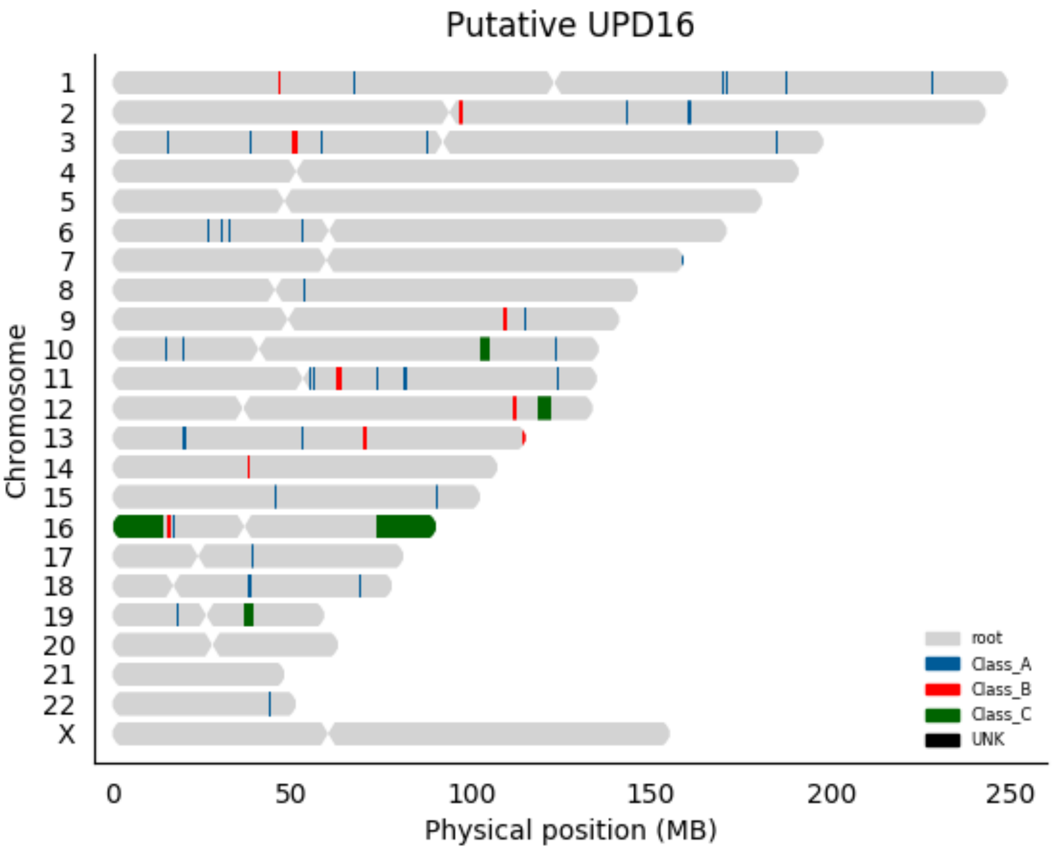

Putative UPD16
Chromosome
X
0
50
100
150
200
250
Physical position (MB)
root
Class_A
Class_B
Class_C
UNK

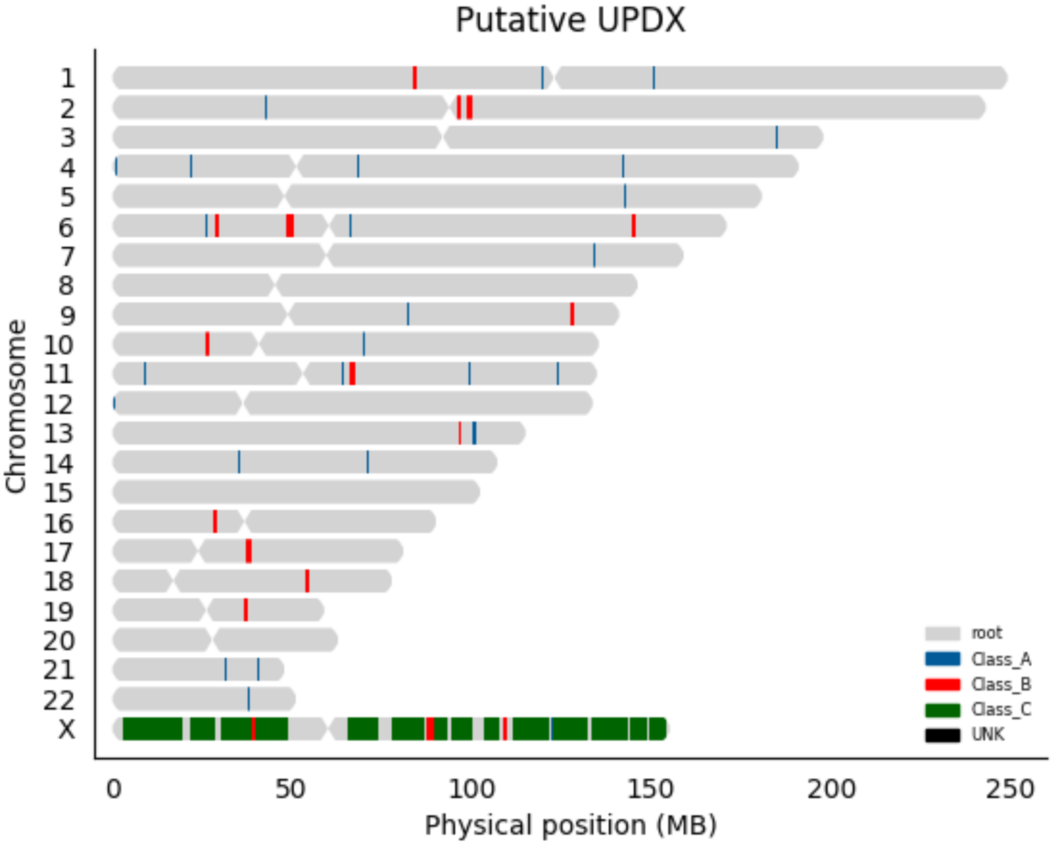

Putative UPDX
Chromosome
X
0
50
100
150
200
250
Physical position (MB)
root
Class_A
Class_B
Class_C
UNK

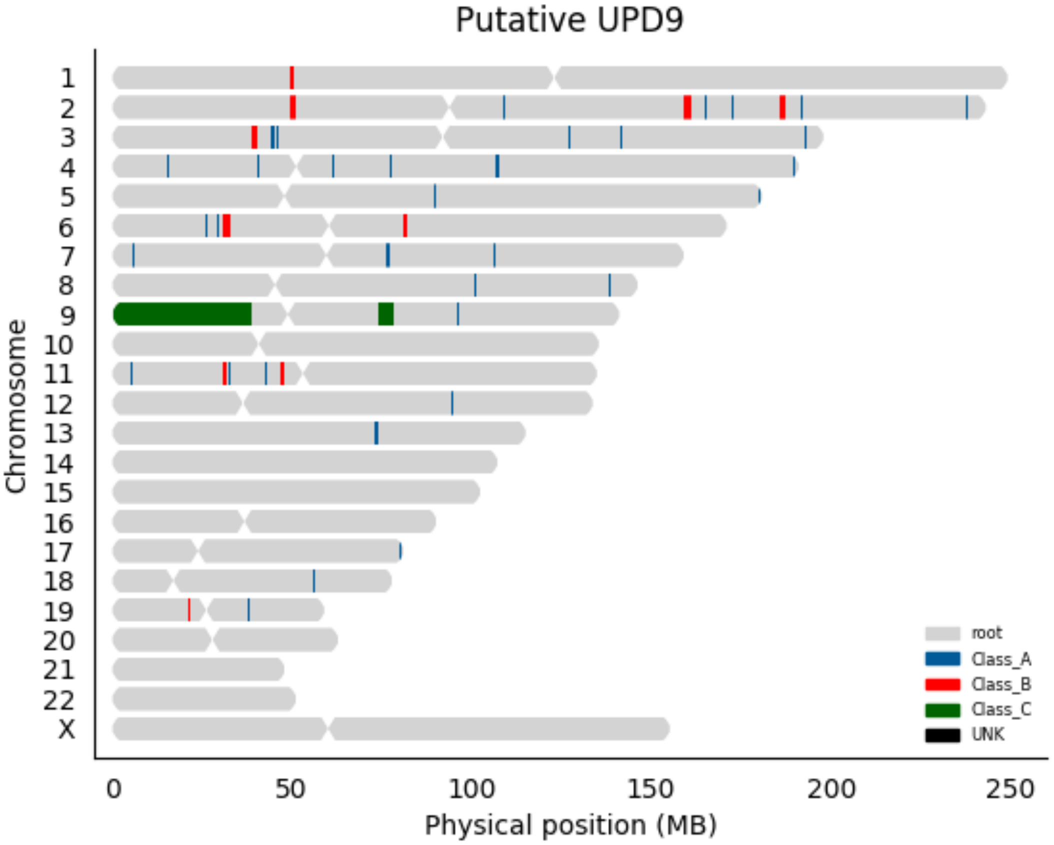
Putative UPD9
Chromosome
X
0
50
100
150
200
250
Physical position (MB)
root
Class_A
Class_B
Class_C
UNK

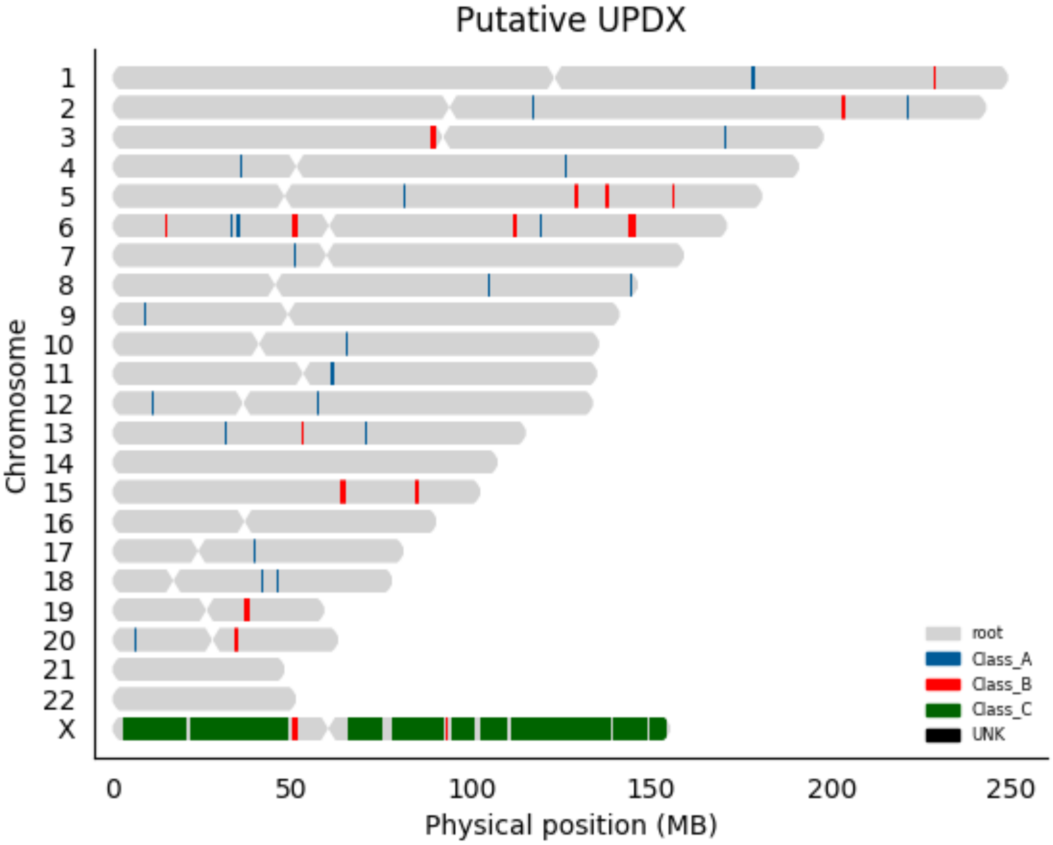
Putative UPDX
Chromosome
X
0
50
100
150
200
250
Physical position (MB)
root
Class_A
Class_B
Class_C
UNK

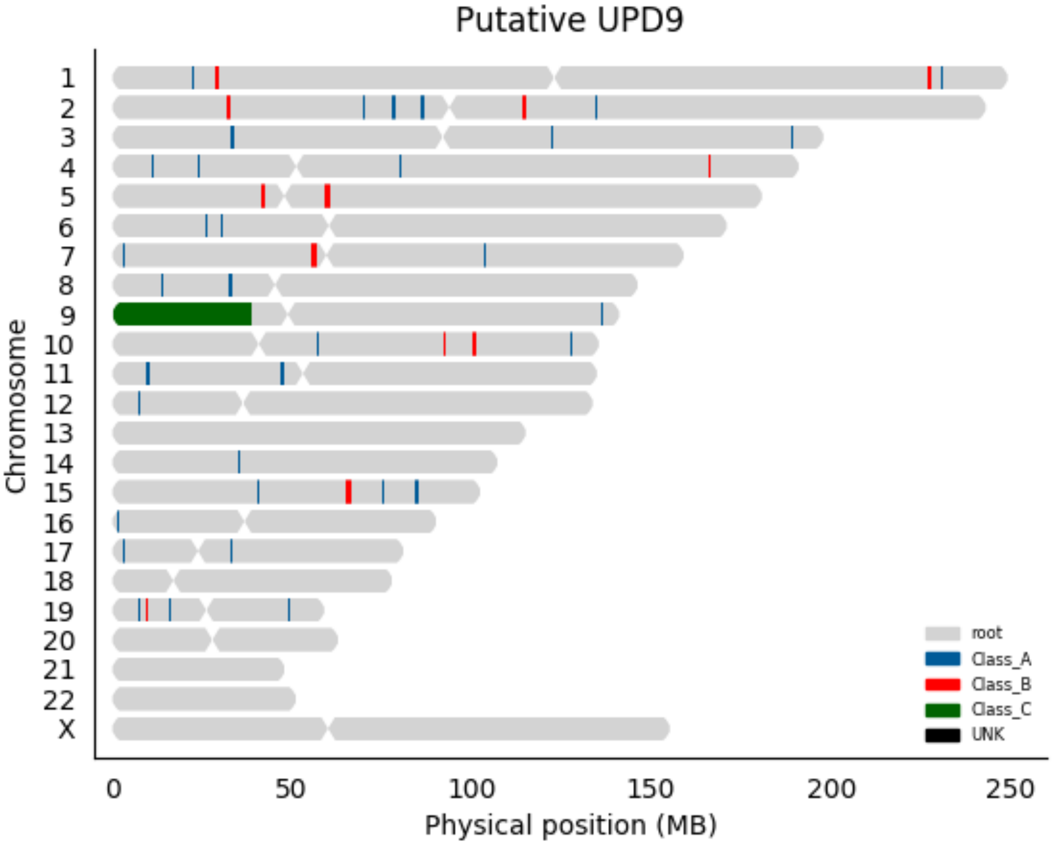

Putative UPD9
Chromosome
X
0
50
100
150
200
250
Physical position (MB)
root
Class_A
Class_B
Class_C
UNK

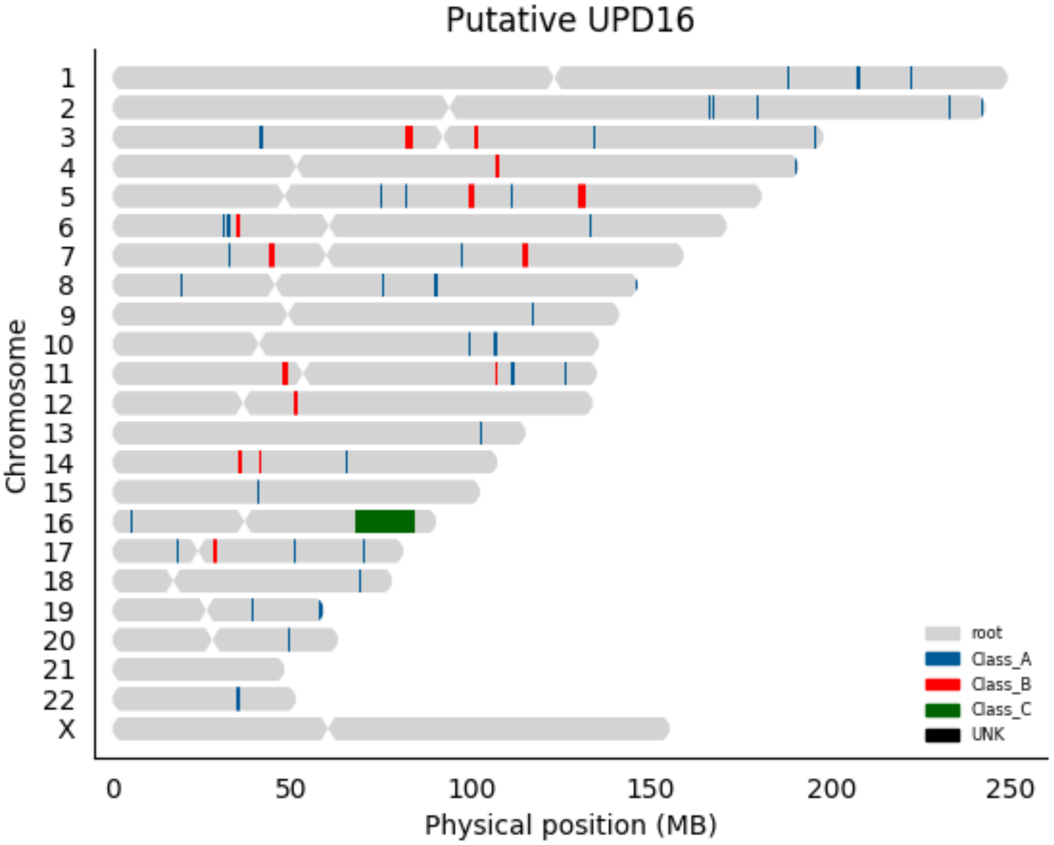

Putative UPD16
Chromosome
X
0
50
100
150
200
250
Physical position (MB)
root
Class_A
Class_B
Class_C
UNK

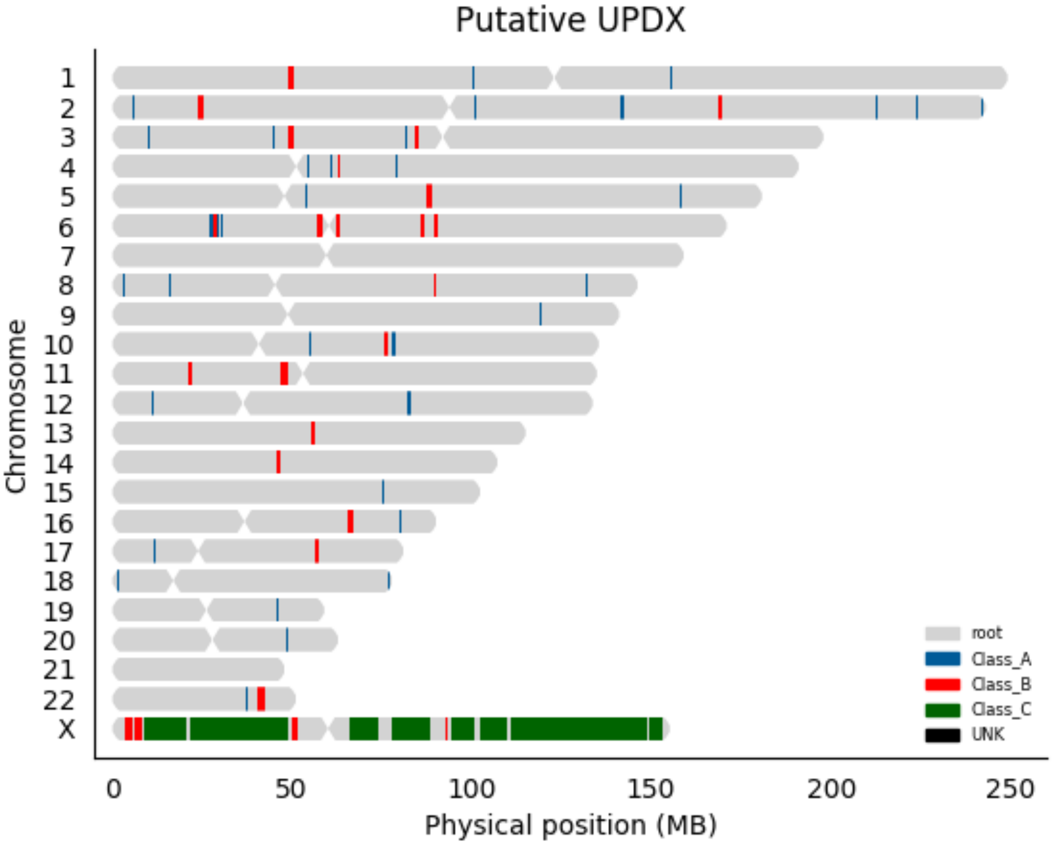

Putative UPDX
Chromosome
X
0
50
100
150
200
250
Physical position (MB)
root
Class_A
Class_B
Class_C
UNK

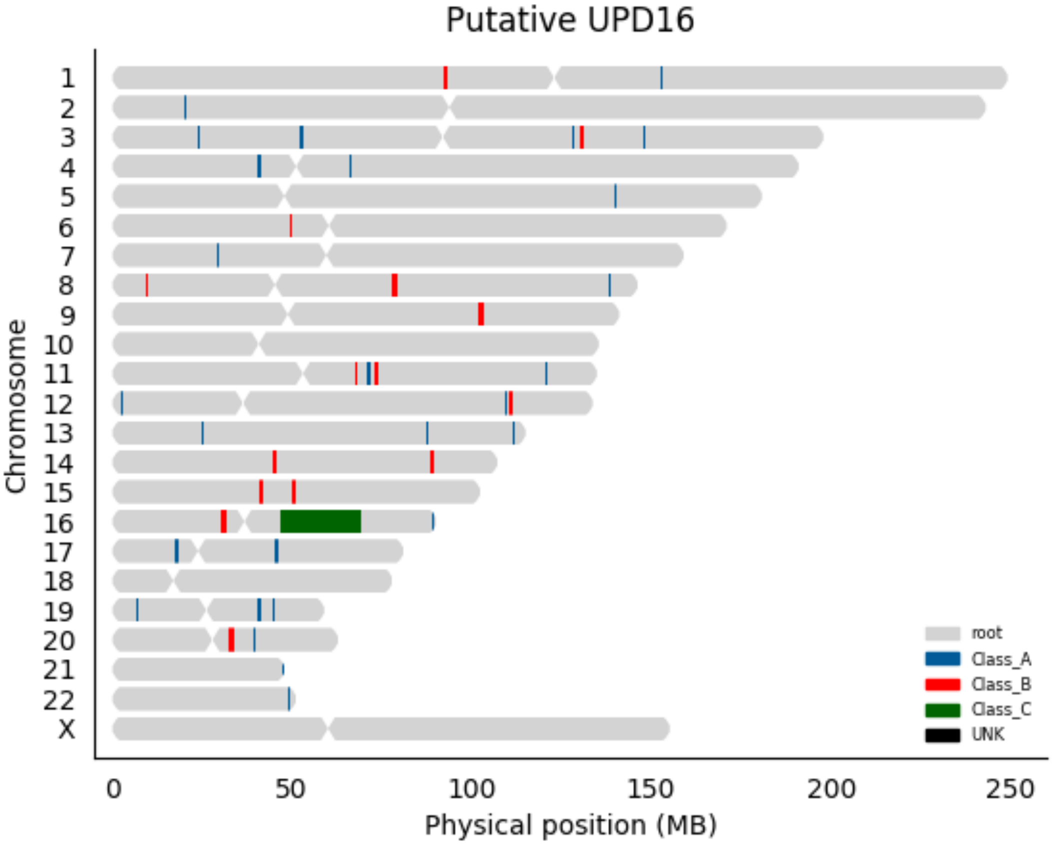

Putative UPD16
Chromosome
X
0
50
100
150
200
250
Physical position (MB)
root
Class_A
Class_B
Class_C
UNK

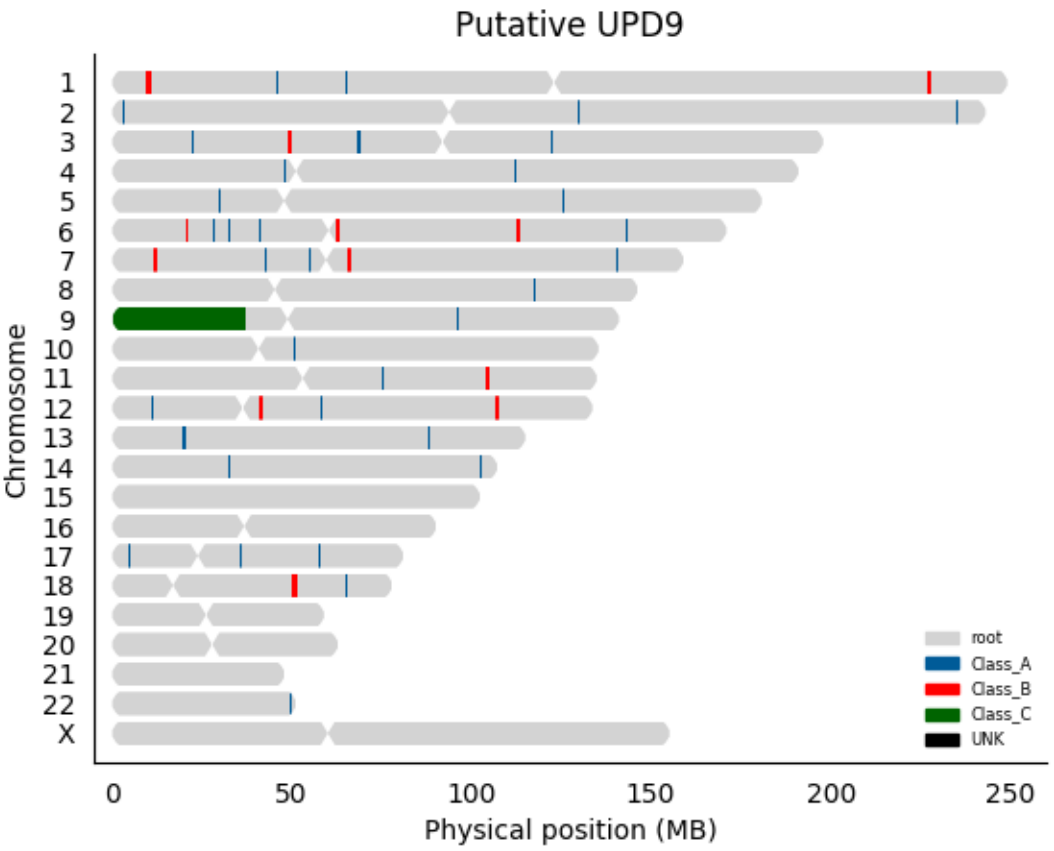

Putative UPD9
Chromosome
X
0
50
100
150
200
250
Physical position (MB)
root
Class_A
Class_B
Class_C
UNK

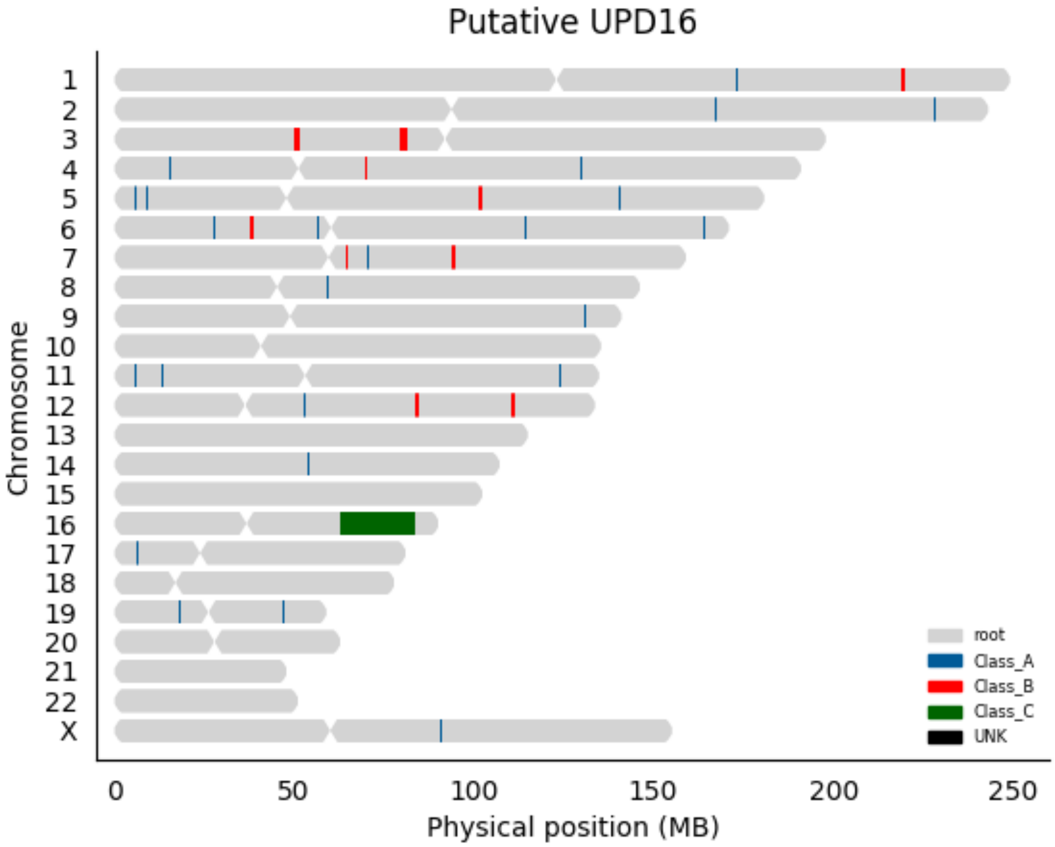

Putative UPD16
Chromosome
X
0
50
100
150
200
250
Physical position (MB)
root
Class_A
Class_B
Class_C
UNK

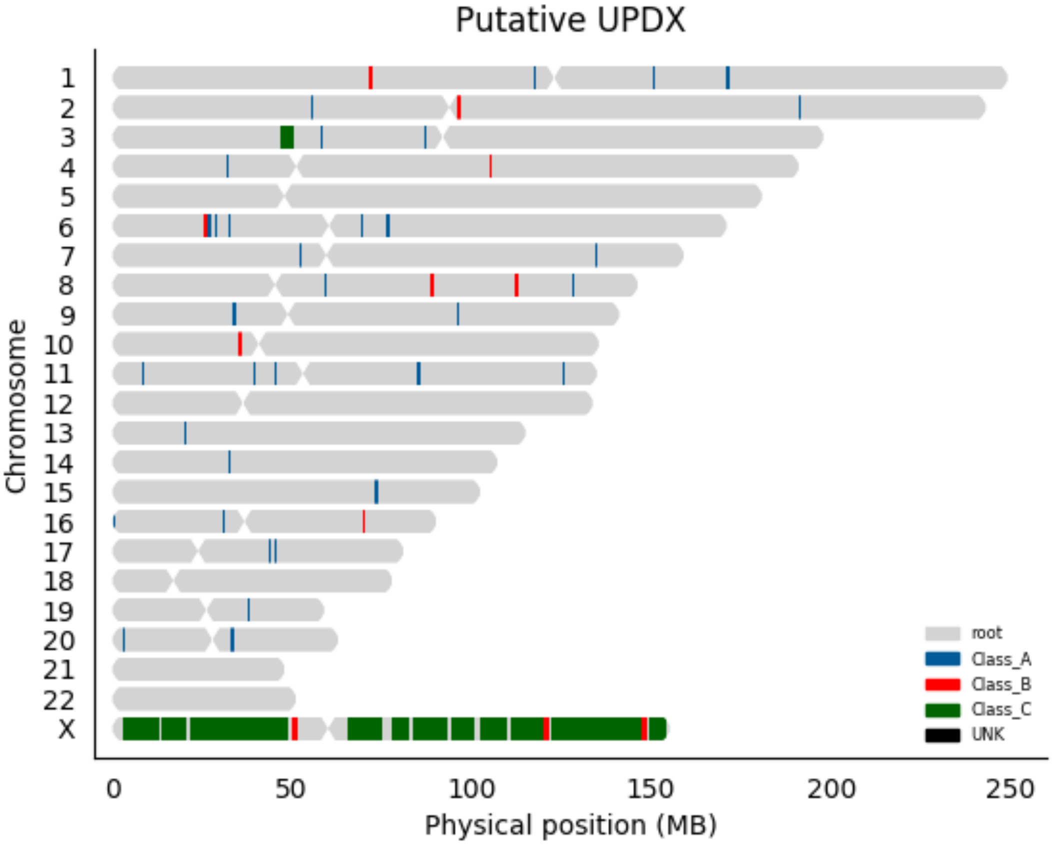

Putative UPDX
Chromosome
X
0
50
100
150
200
250
Physical position (MB)
root
Class_A
Class_B
Class_C
UNK

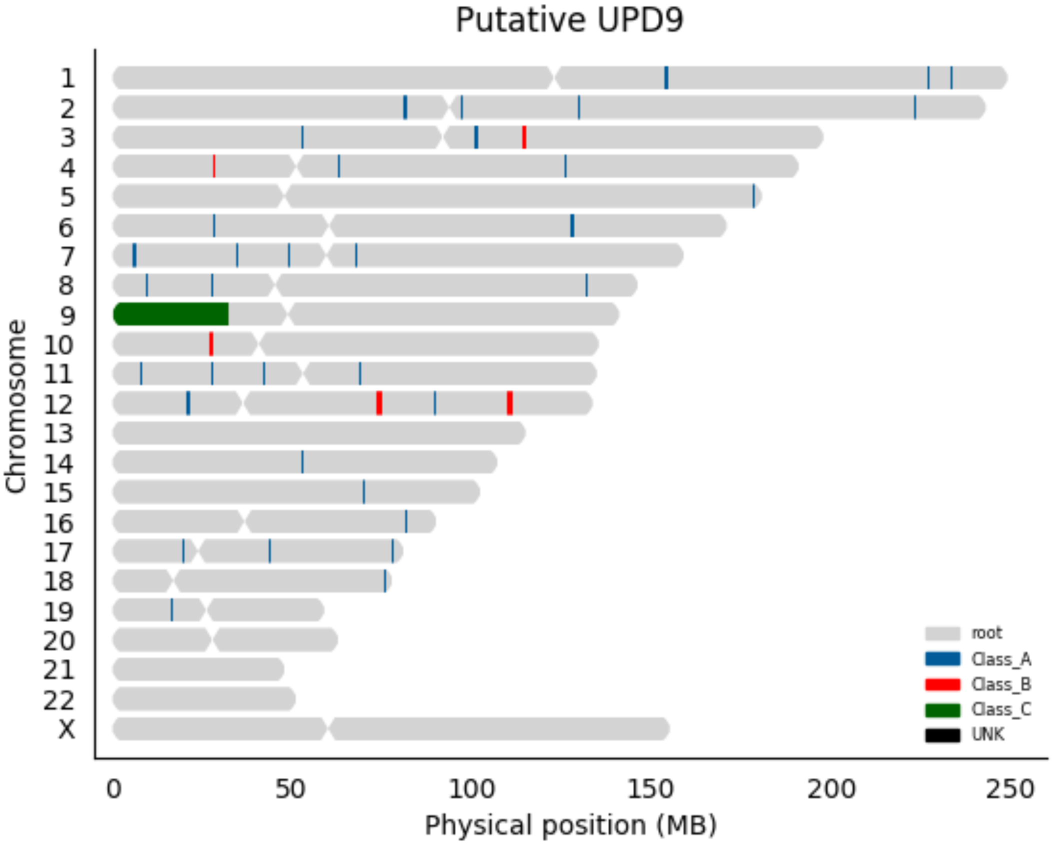
Putative UPD9
Chromosome
X
0
50
100
150
200
250
Physical position (MB)
root
Class_A
Class_B
Class_C
UNK

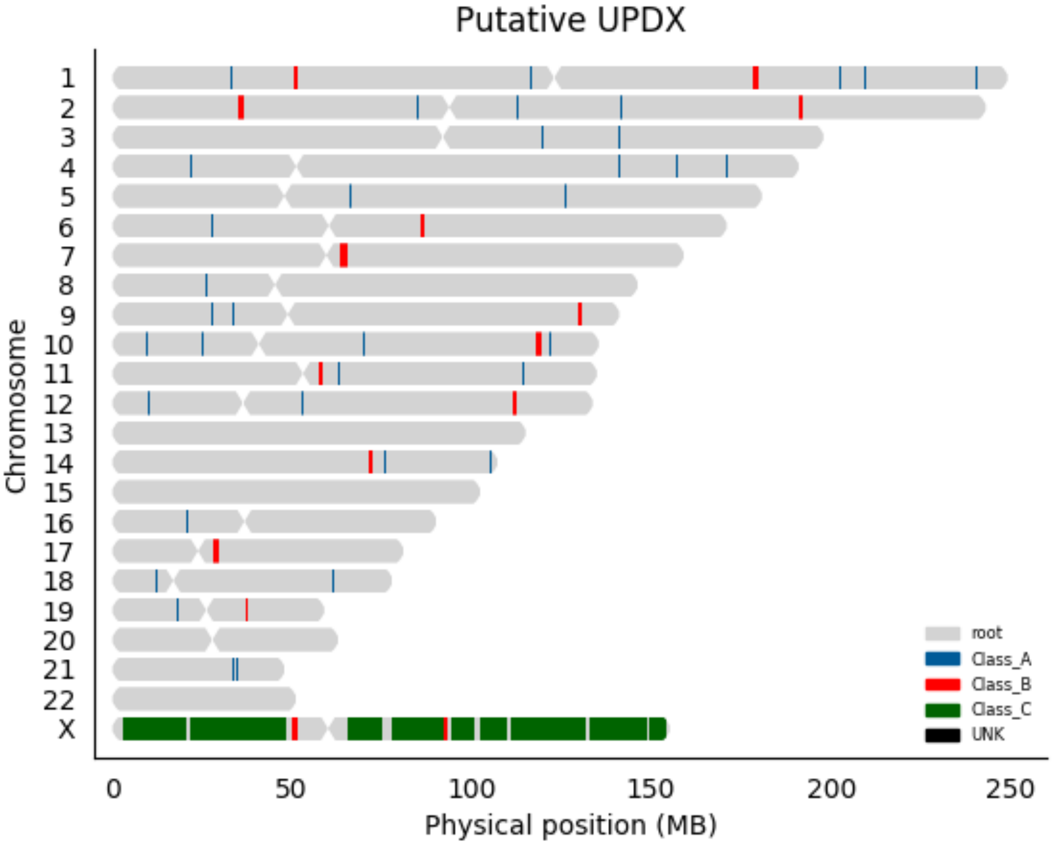

Putative UPDX
Chromosome
X
root
Class_A
Class_B
Class_C
UNK
0
50
100
150
200
250
Physical position (MB)

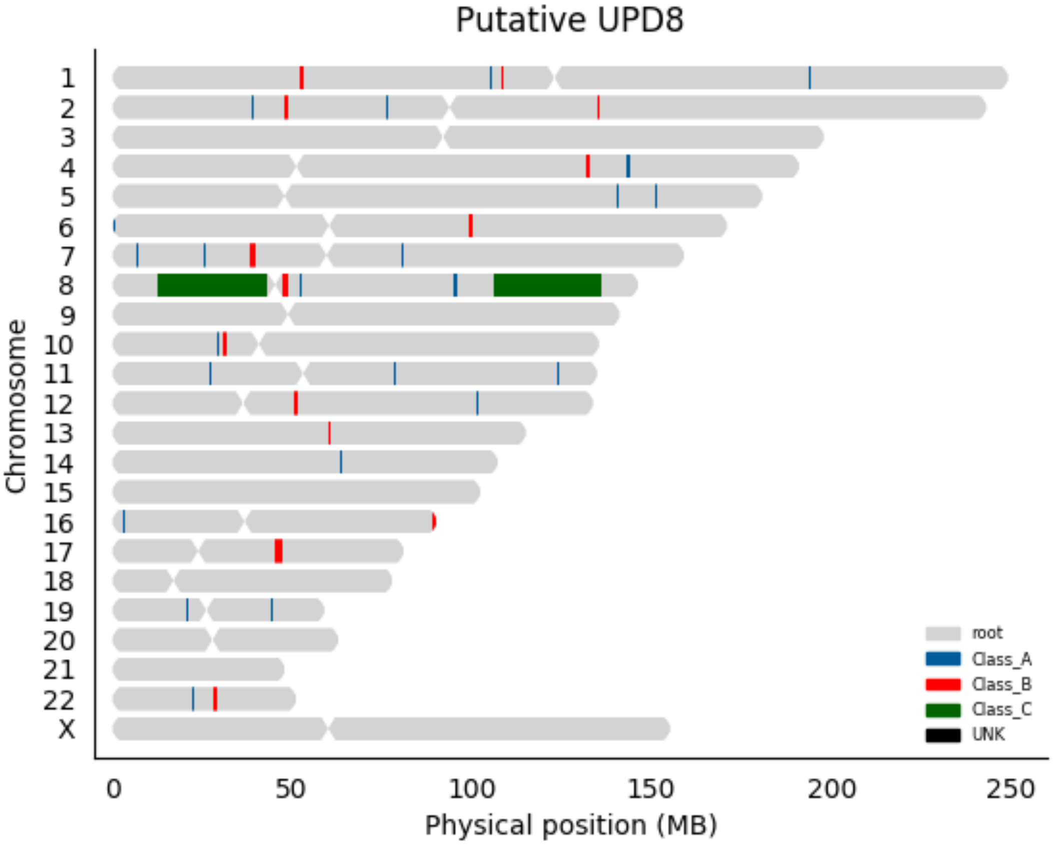

Putative UPD8
Chromosome
X
0
50
100
150
200
250
Physical position (MB)
root
Class_A
Class_B
Class_C
UNK

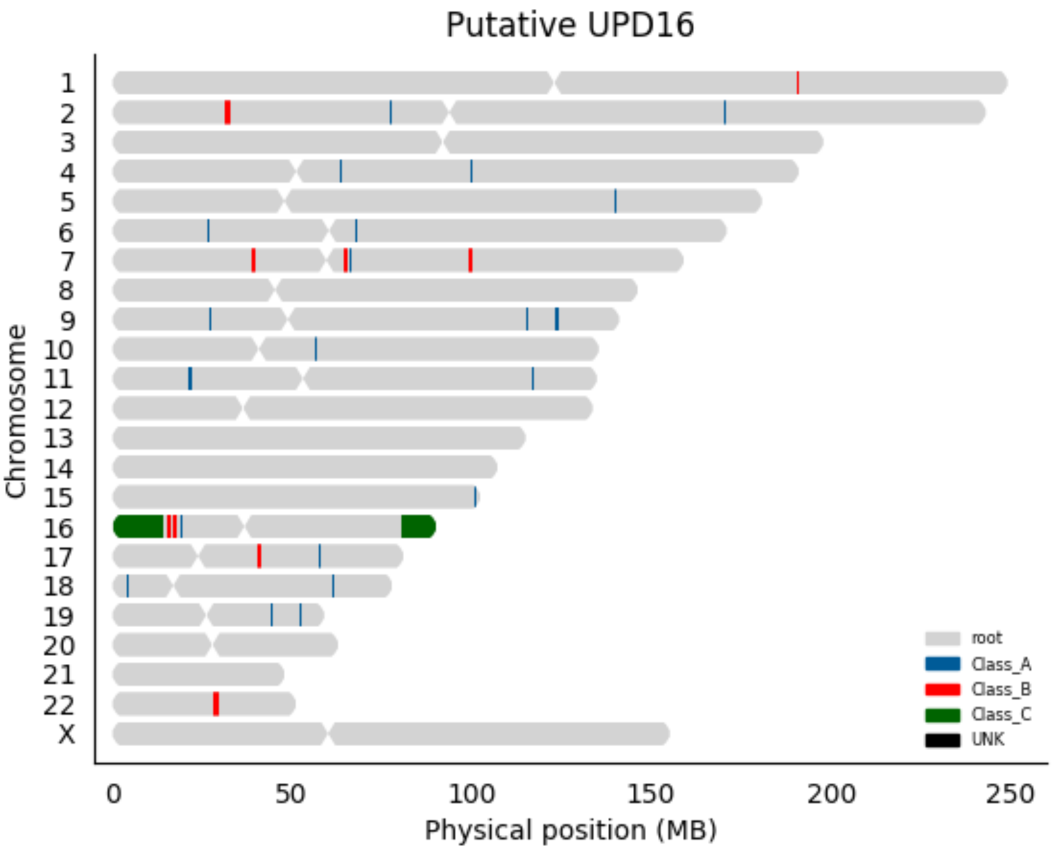

Putative UPD16
Chromosome
X
0
50
100
150
200
250
Physical position (MB)
root
Class_A
Class_B
Class_C
UNK

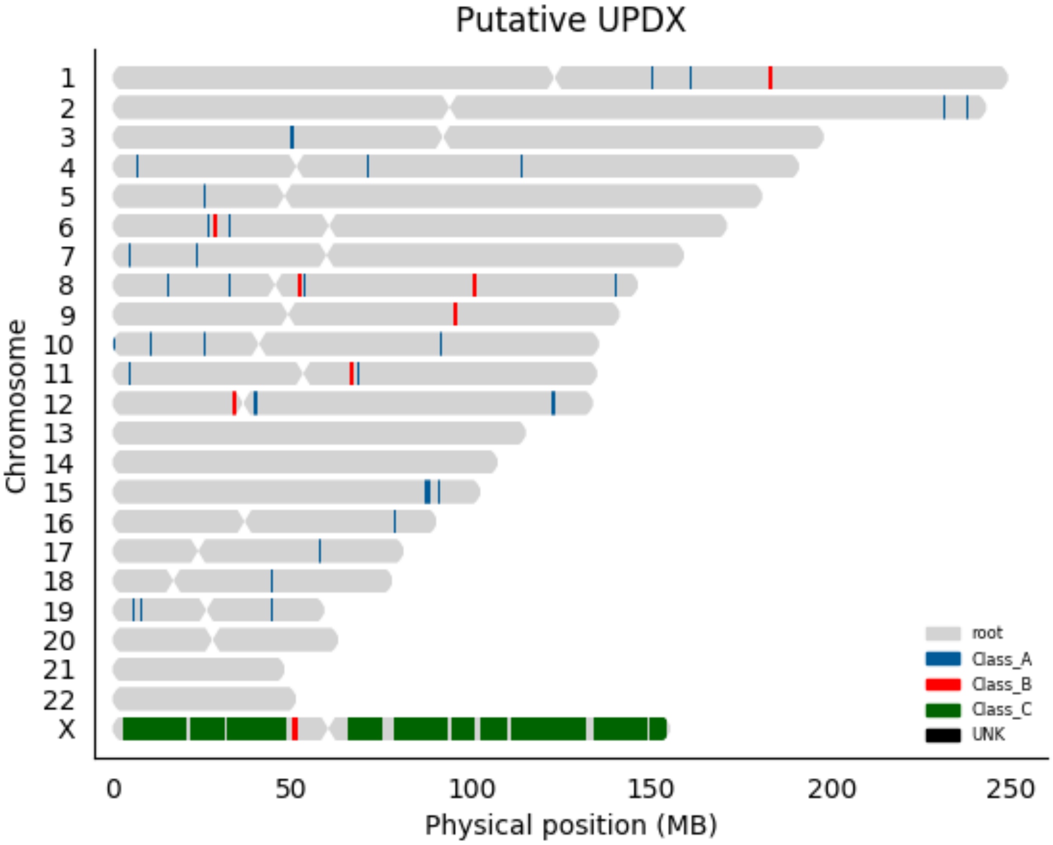

Putative UPDX
Chromosome
X
0
50
100
150
200
250
Physical position (MB)
root
Class_A
Class_B
Class_C
UNK

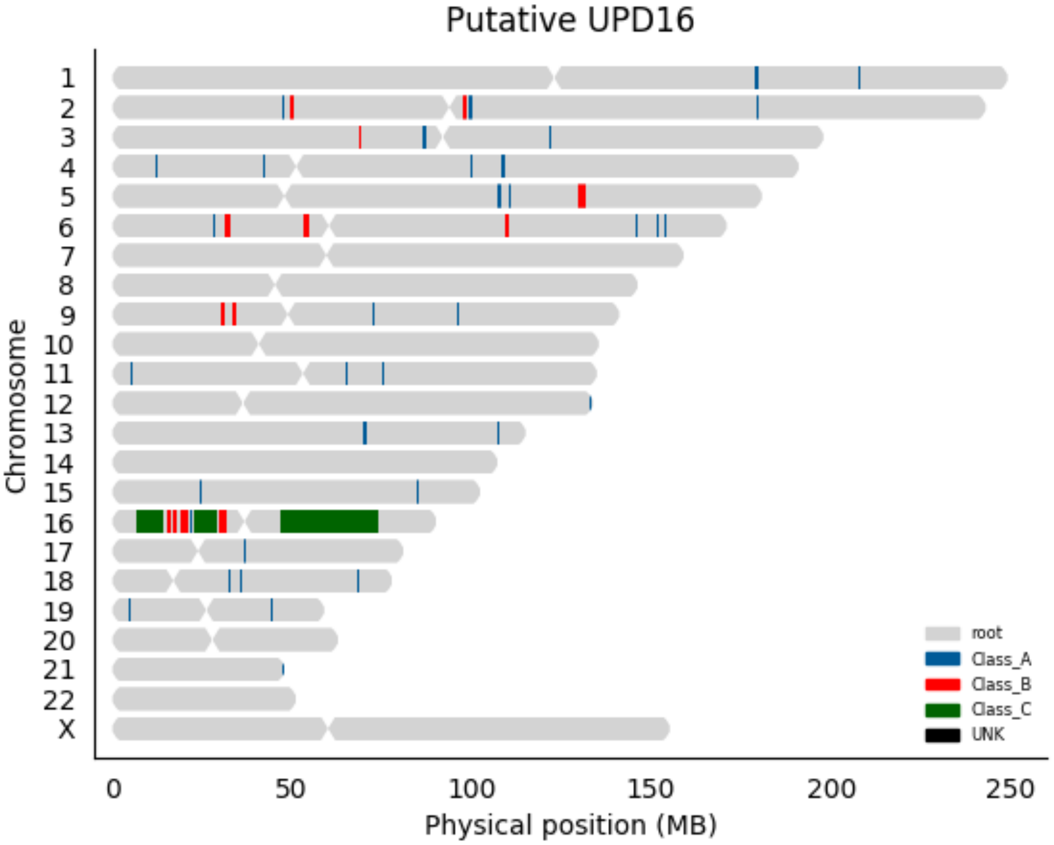

Putative UPD16
Chromosome
X
0
50
100
150
200
250
Physical position (MB)
root
Class_A
Class_B
Class_C
UNK

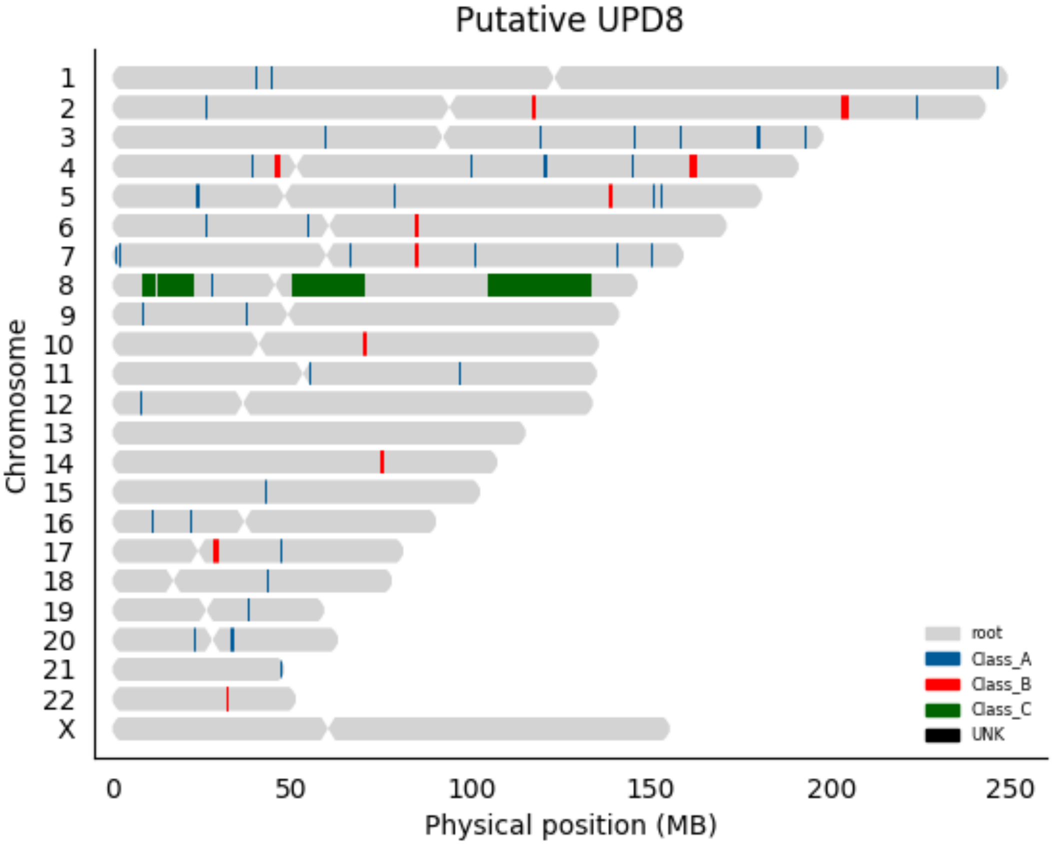
Putative UPD8
Chromosome
X
0
50
100
150
200
250
Physical position (MB)
root
Class_A
Class_B
Class_C
UNK

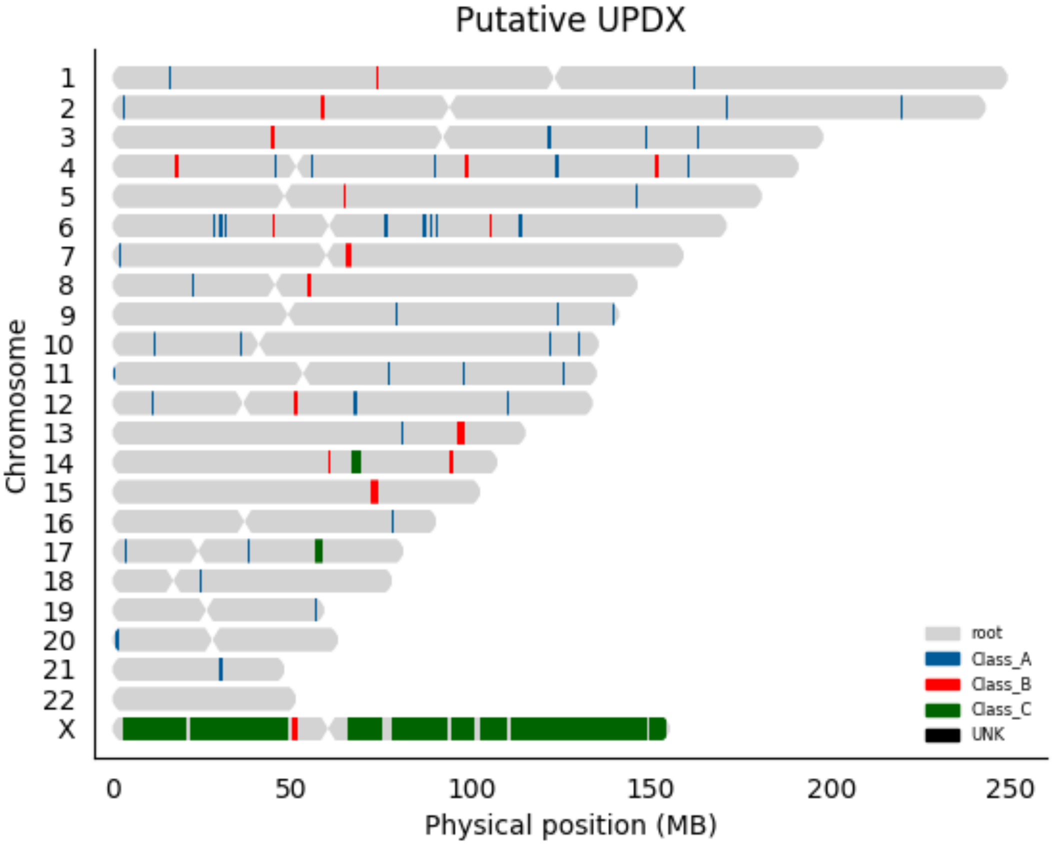

Putative UPDX
Chromosome
X
0
50
100
150
200
250
Physical position (MB)
root
Class_A
Class_B
Class_C
UNK

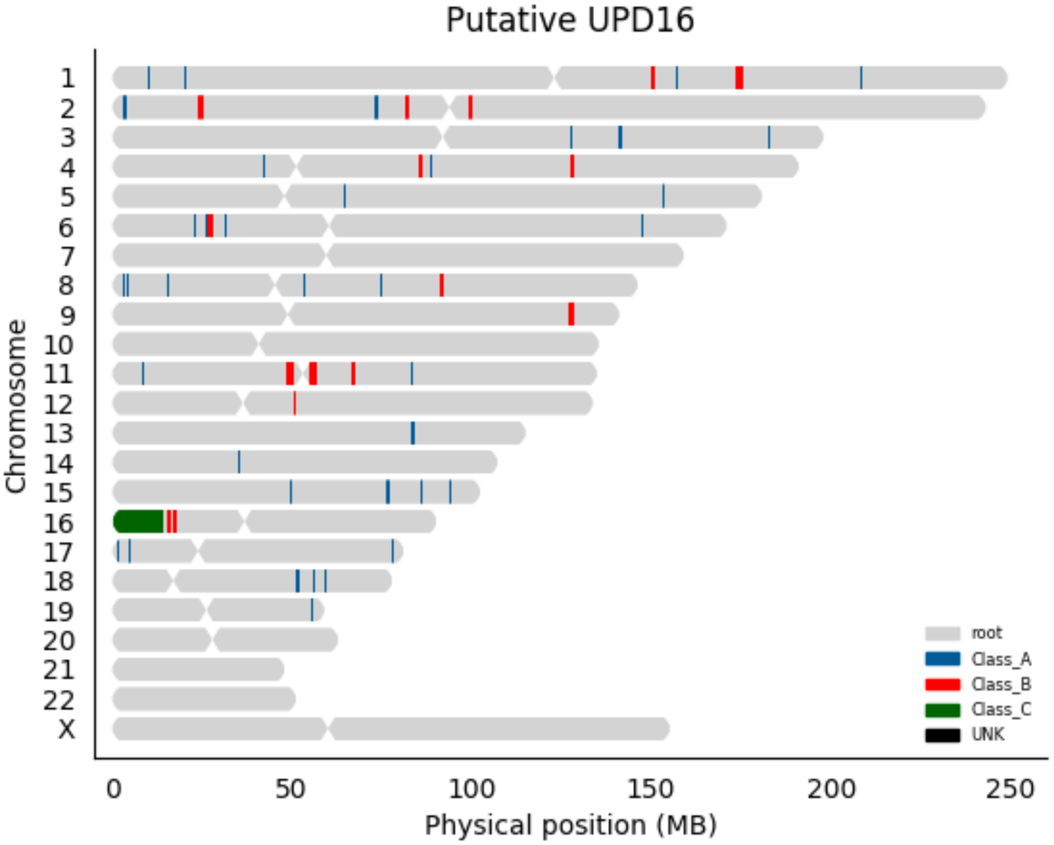

Putative UPD16
Chromosome
X
0
50
100
150
200
250
Physical position (MB)
root
Class_A
Class_B
Class_C
UNK

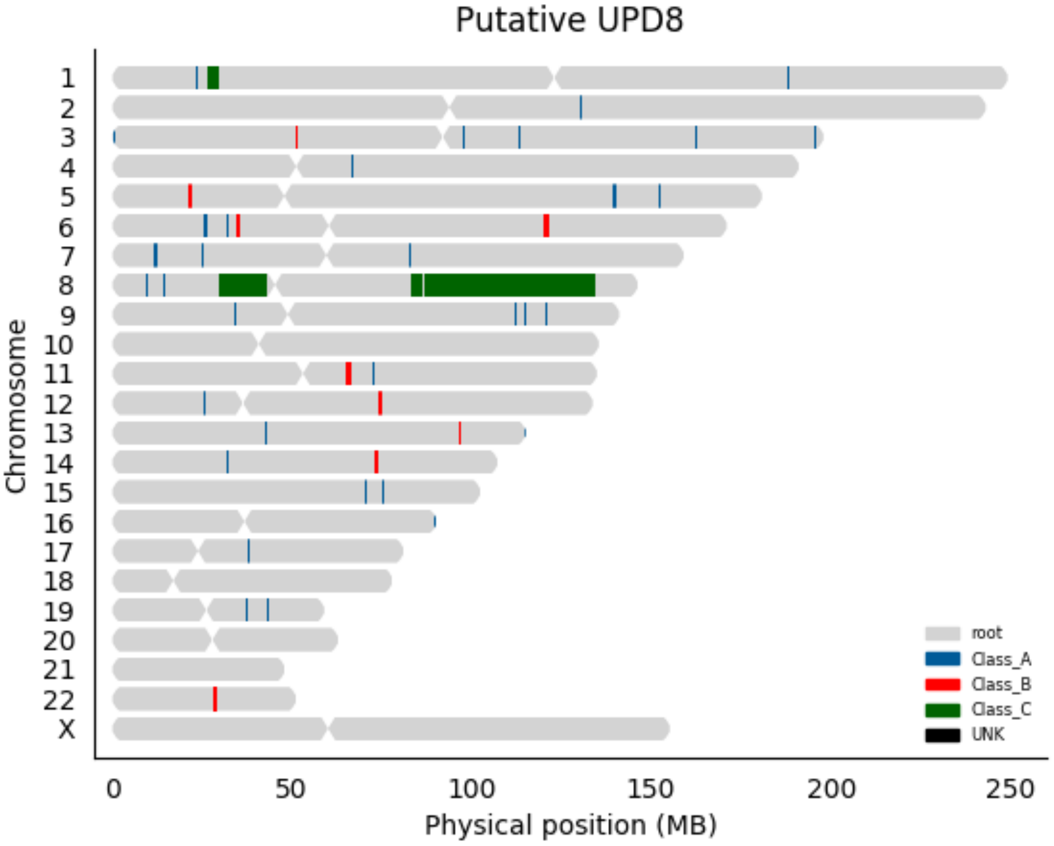
Putative UPD8
Chromosome
X
0
50
100
150
200
250
Physical position (MB)
root
Class_A
Class_B
Class_C
UNK

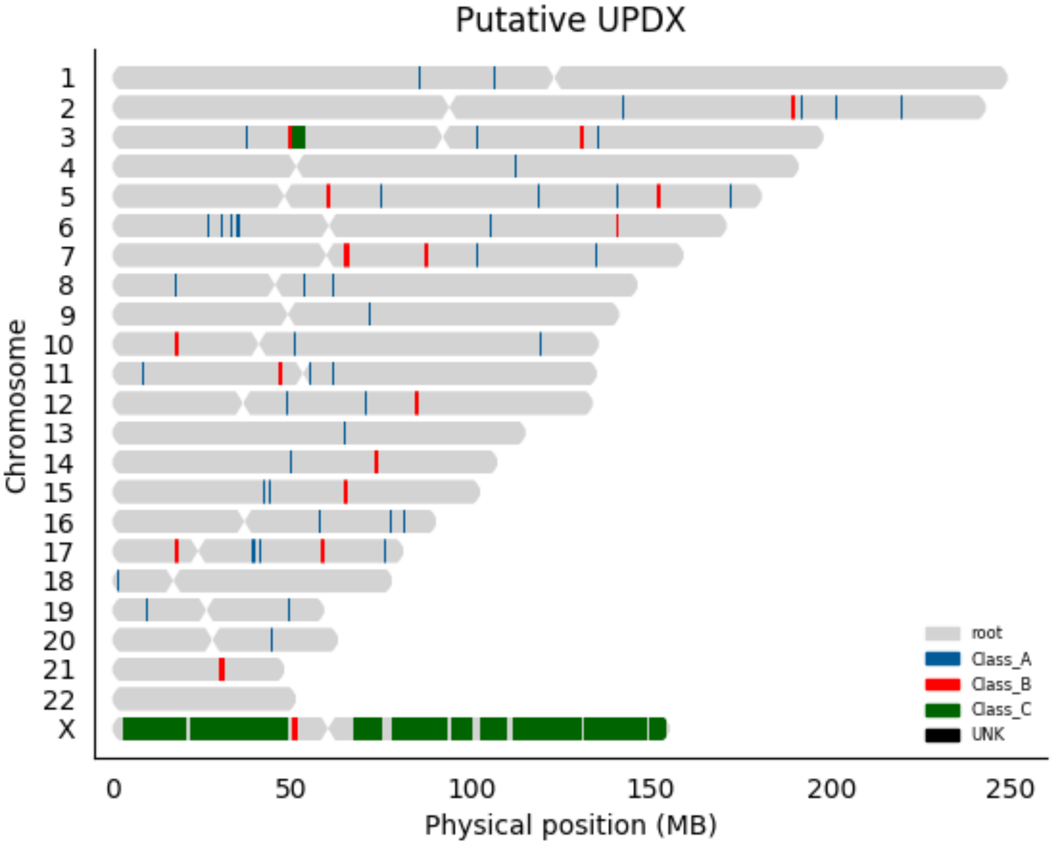

Putative UPDX
Chromosome
X
0
50
100
150
200
250
Physical position (MB)
root
Class_A
Class_B
Class_C
UNK

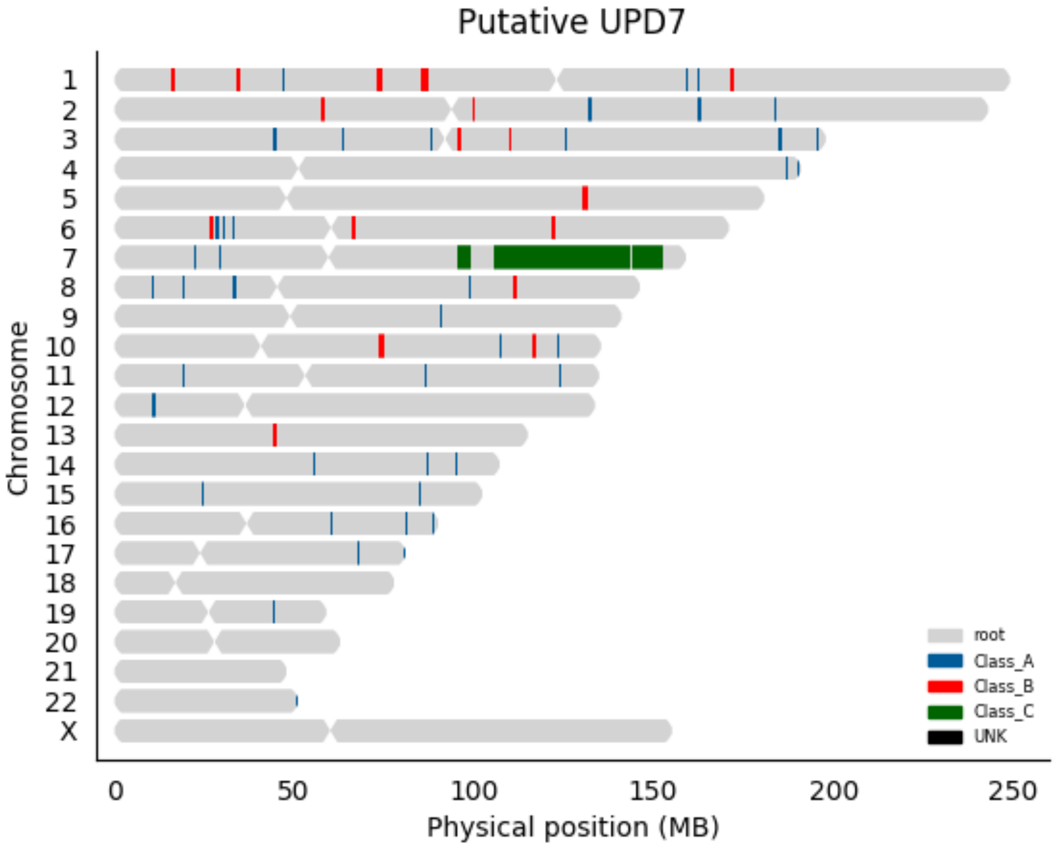

Putative UPD7
Chromosome
X
0
50
100
150
200
250
Physical position (MB)
root
Class_A
Class_B
Class_C
UNK

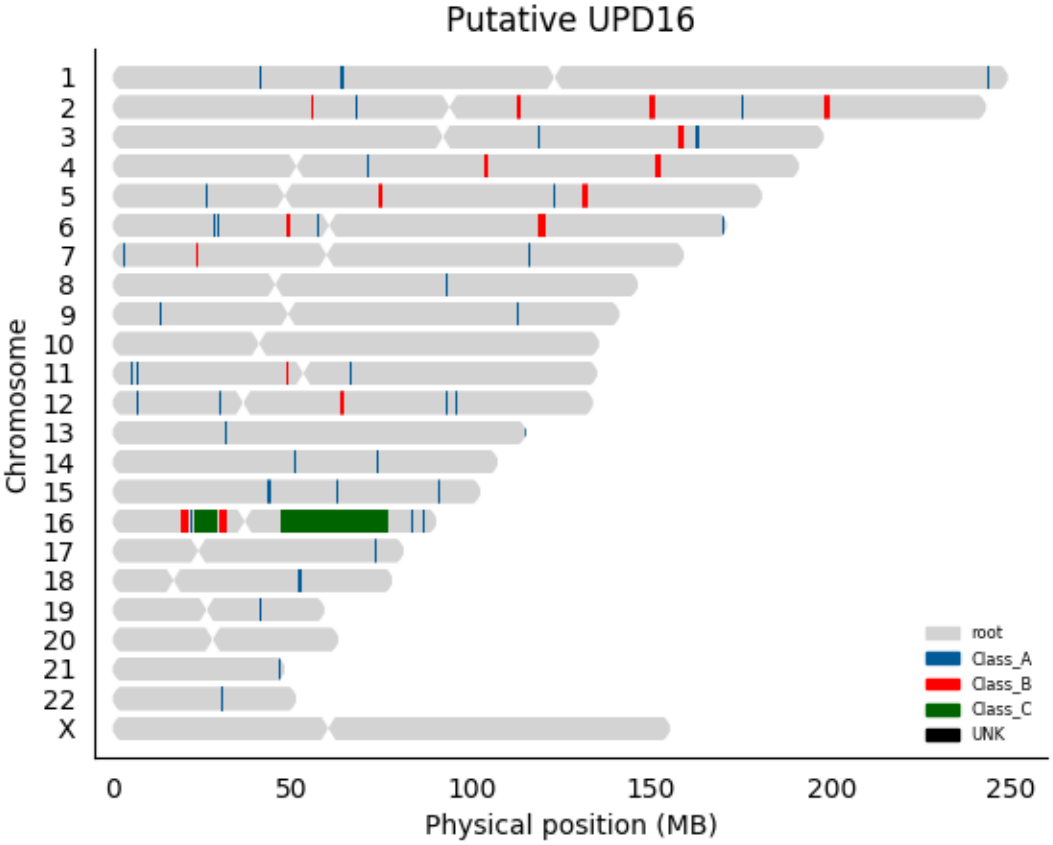

Putative UPD16
Chromosome
X
0
50
100
150
200
250
Physical position (MB)
root
Class_A
Class_B
Class_C
UNK

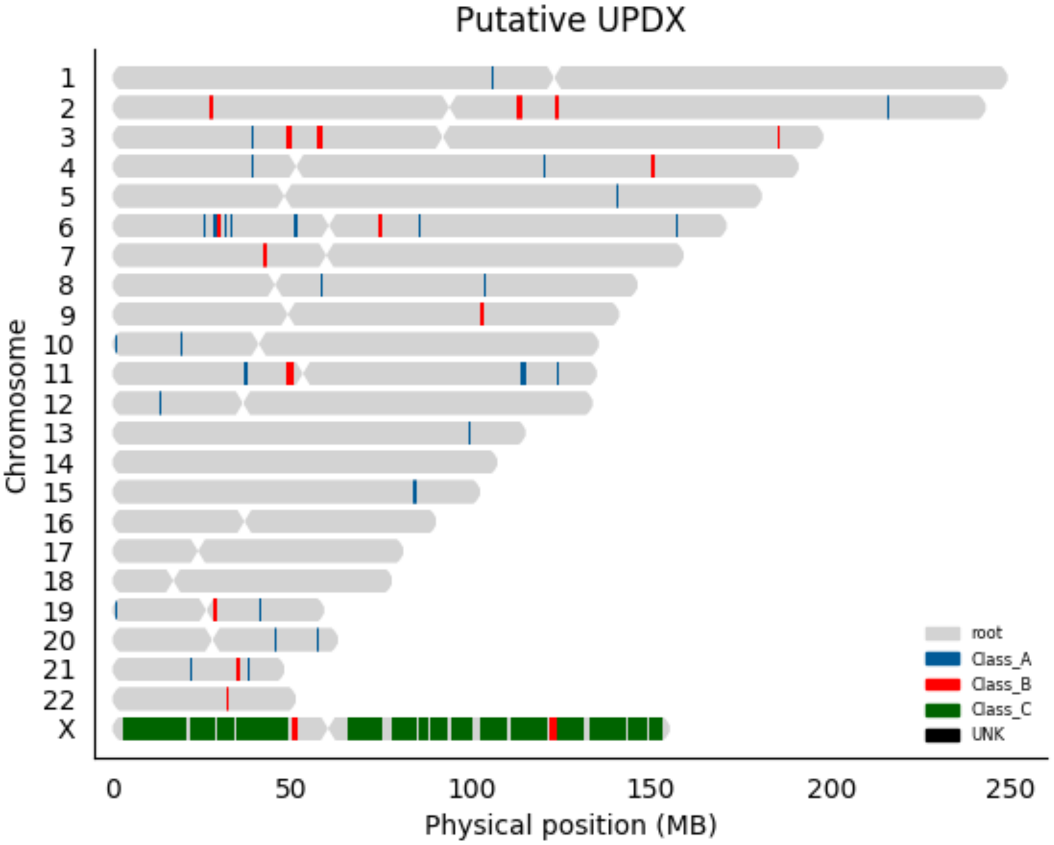

Putative UPDX
Chromosome
X
0
50
100
150
200
250
Physical position (MB)
root
Class_A
Class_B
Class_C
UNK

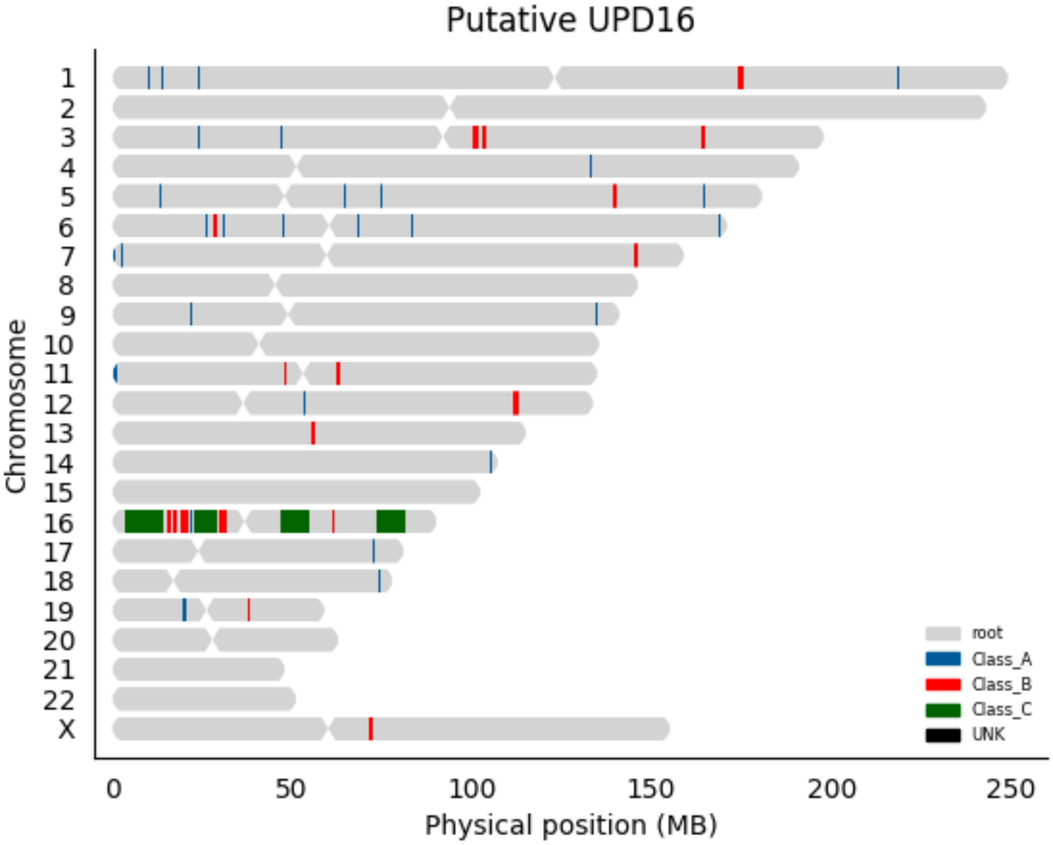

Putative UPD16
Chromosome
X
root
Class_A
Class_B
Class_C
UNK
0
50
100
150
200
250
Physical position (MB)

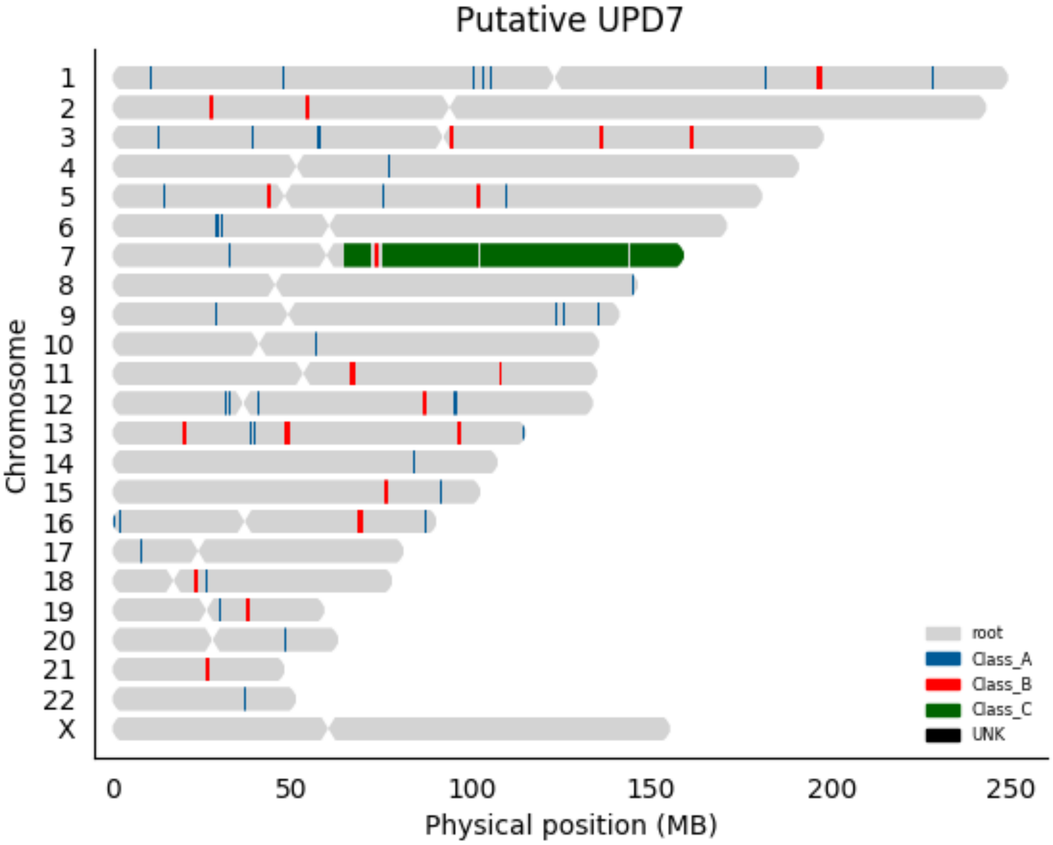
Putative UPD7
Chromosome
X
0
50
100
150
200
250
Physical position (MB)
root
Class_A
Class_B
Class_C
UNK

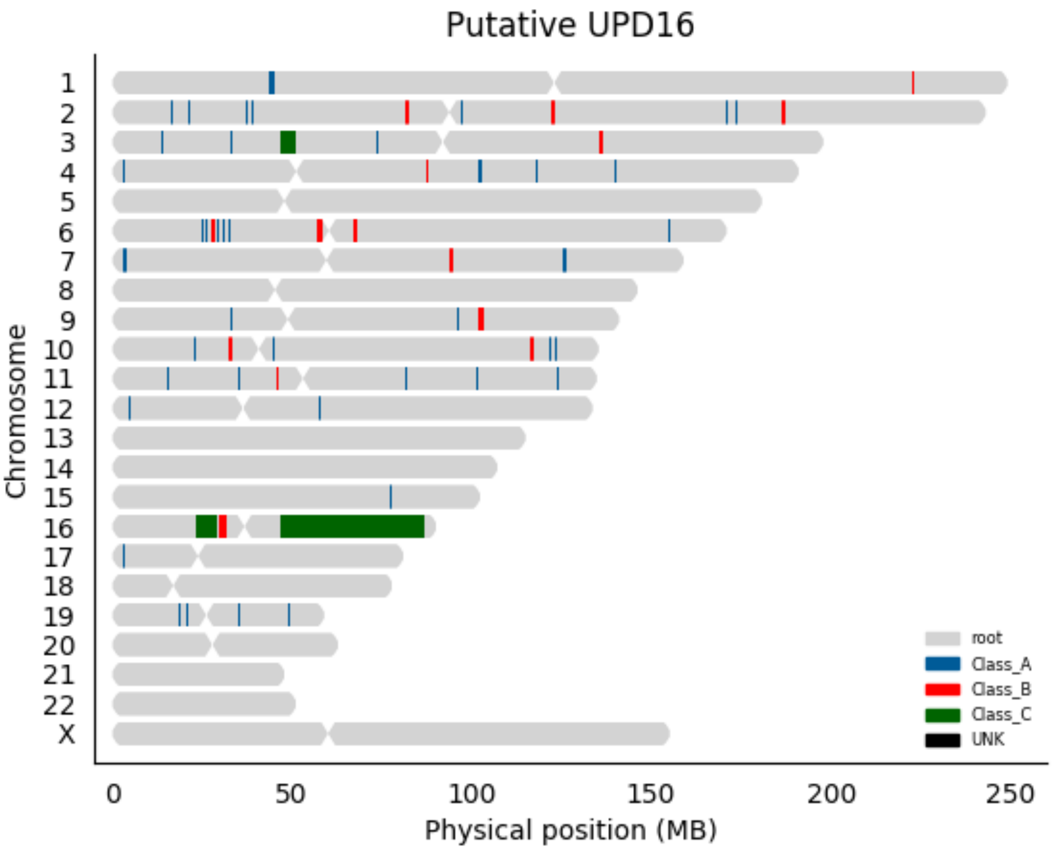

Putative UPD16
Chromosome
X
0
50
100
150
200
250
Physical position (MB)
root
Class_A
Class_B
Class_C
UNK

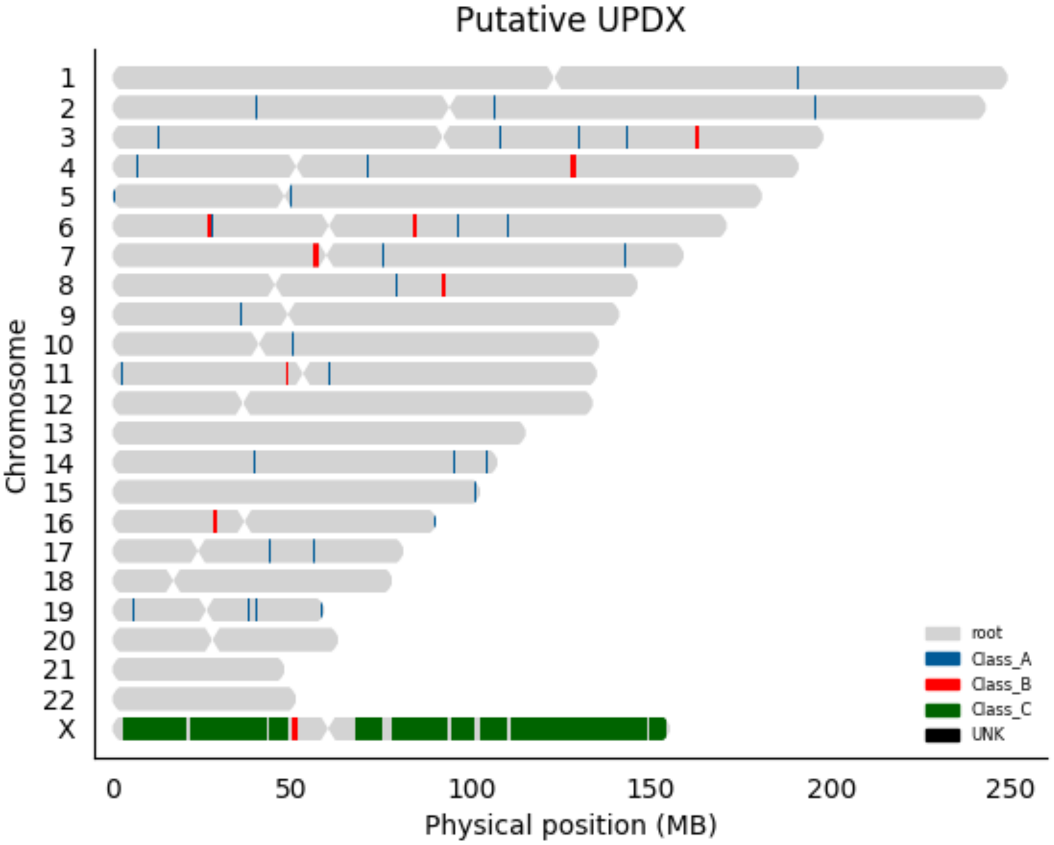

Putative UPDX
Chromosome
X
0
50
100
150
200
250
Physical position (MB)
root
Class_A
Class_B
Class_C
UNK

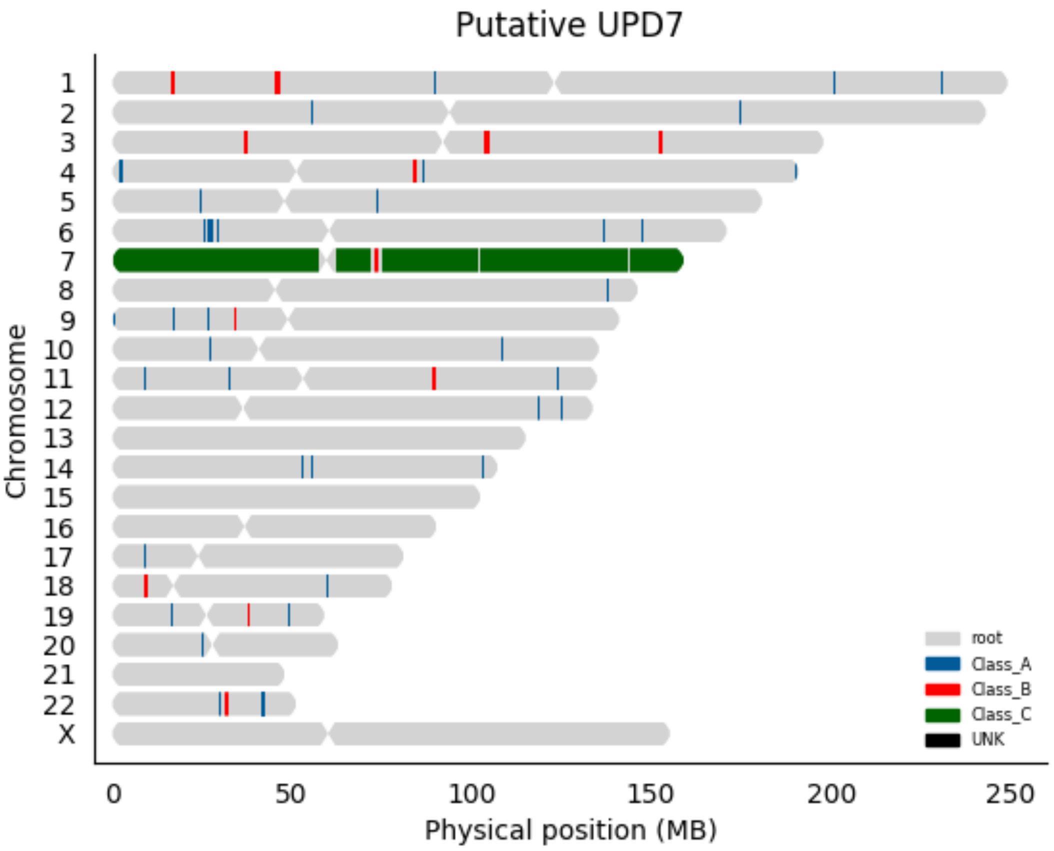

Putative UPD7
Chromosome
X
0
50
100
150
200
250
Physical position (MB)
root
Class_A
Class_B
Class_C
UNK

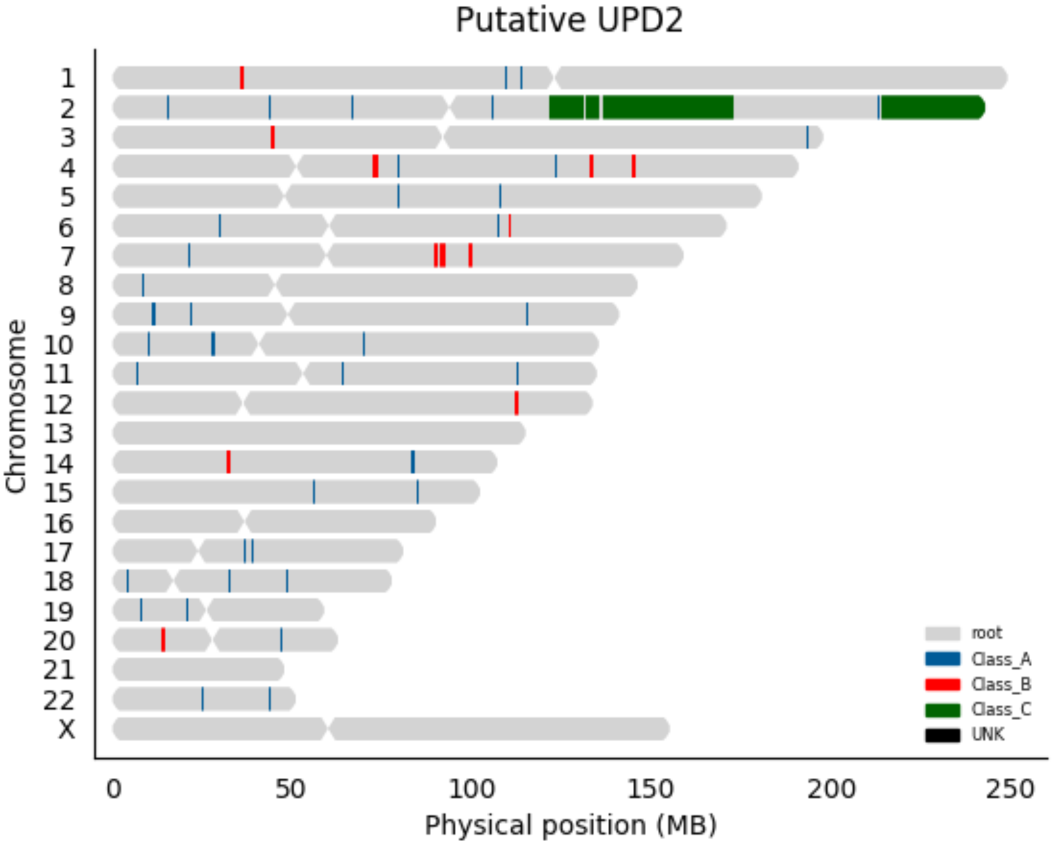

Putative UPD2
Chromosome
X
0
50
100
150
200
250
Physical position (MB)
root
Class_A
Class_B
Class_C
UNK

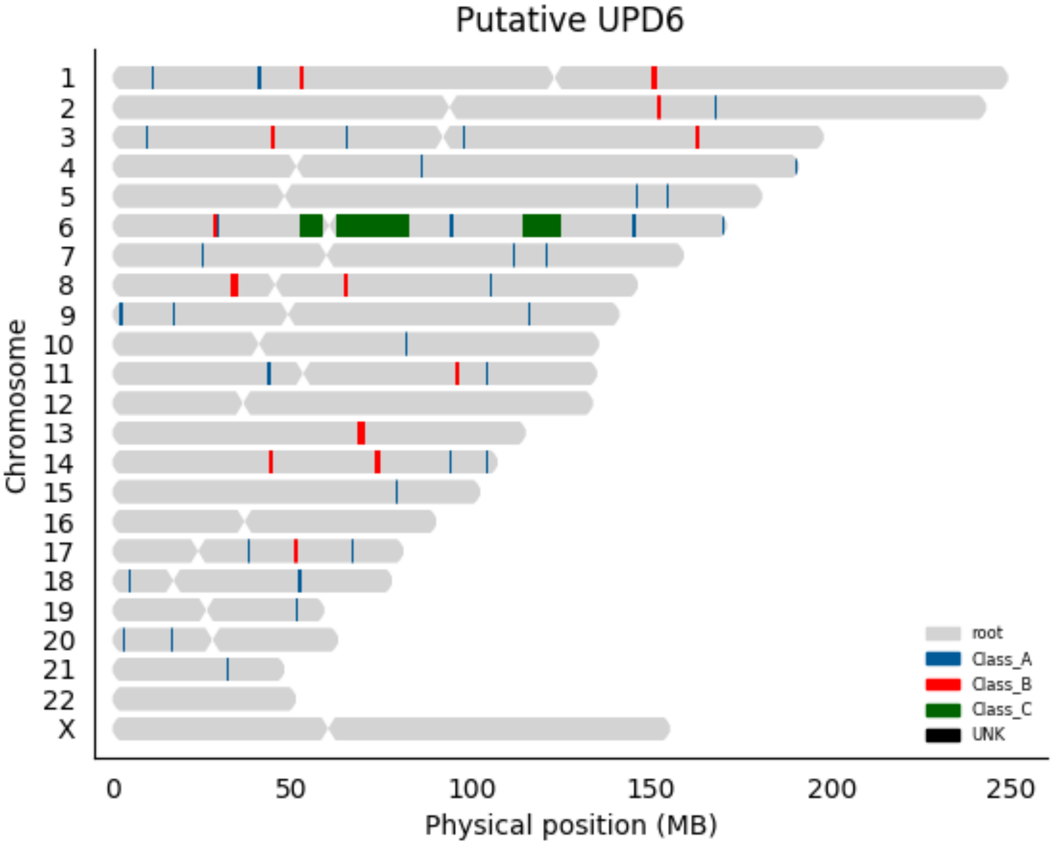

Putative UPD6
Chromosome
X
0
50
100
150
200
250
Physical position (MB)
root
Class_A
Class_B
Class_C
UNK

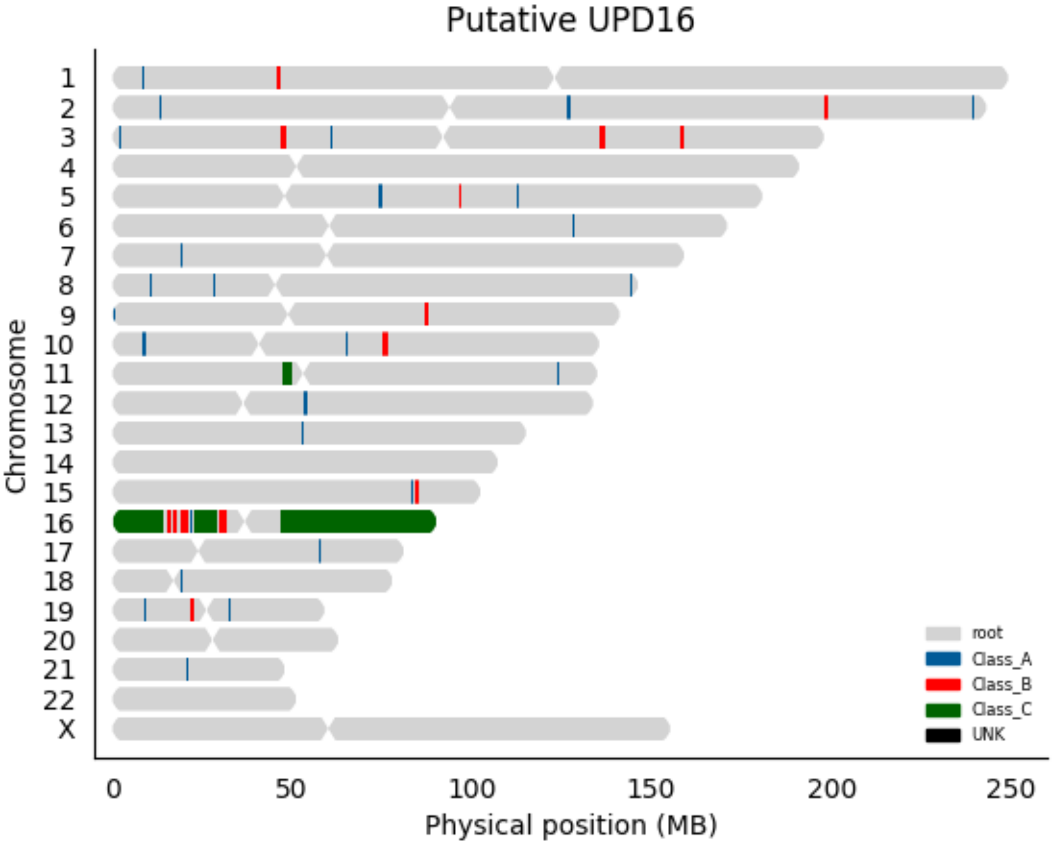

Putative UPD16
Chromosome
X
0
50
100
150
200
250
Physical position (MB)
root
Class_A
Class_B
Class_C
UNK

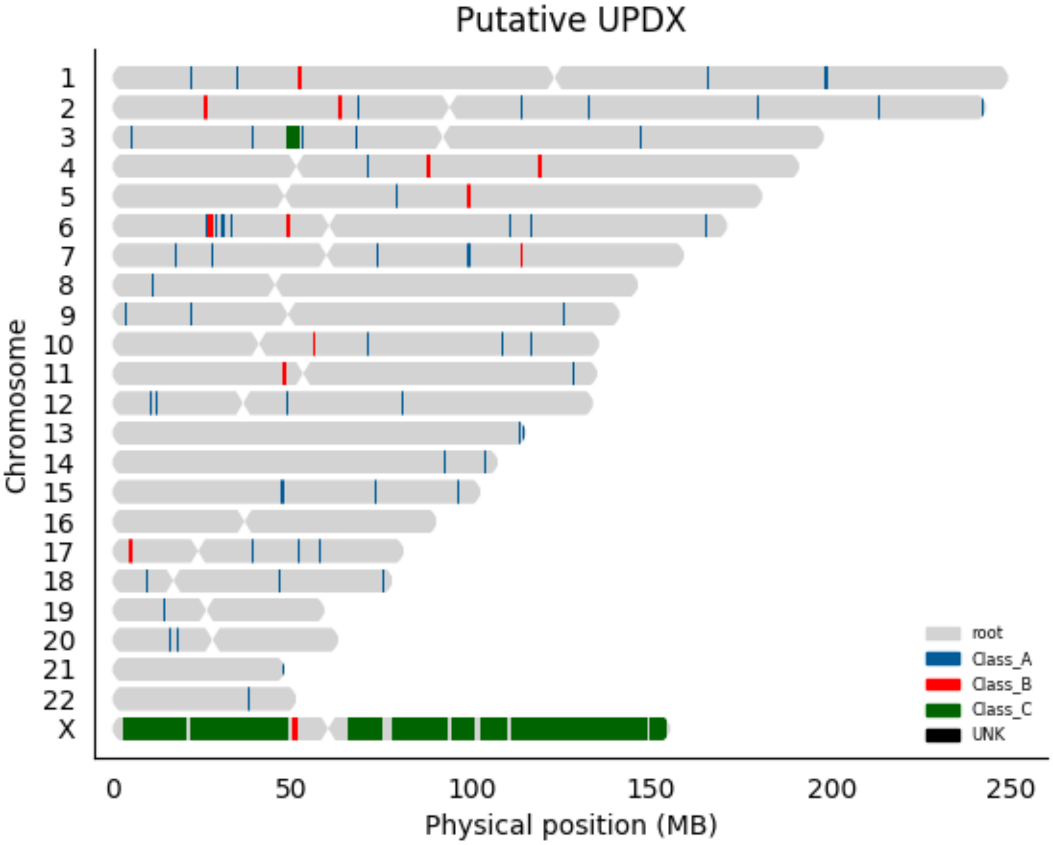

Putative UPDX
Chromosome
X
0
50
100
150
200
250
Physical position (MB)
root
Class_A
Class_B
Class_C
UNK

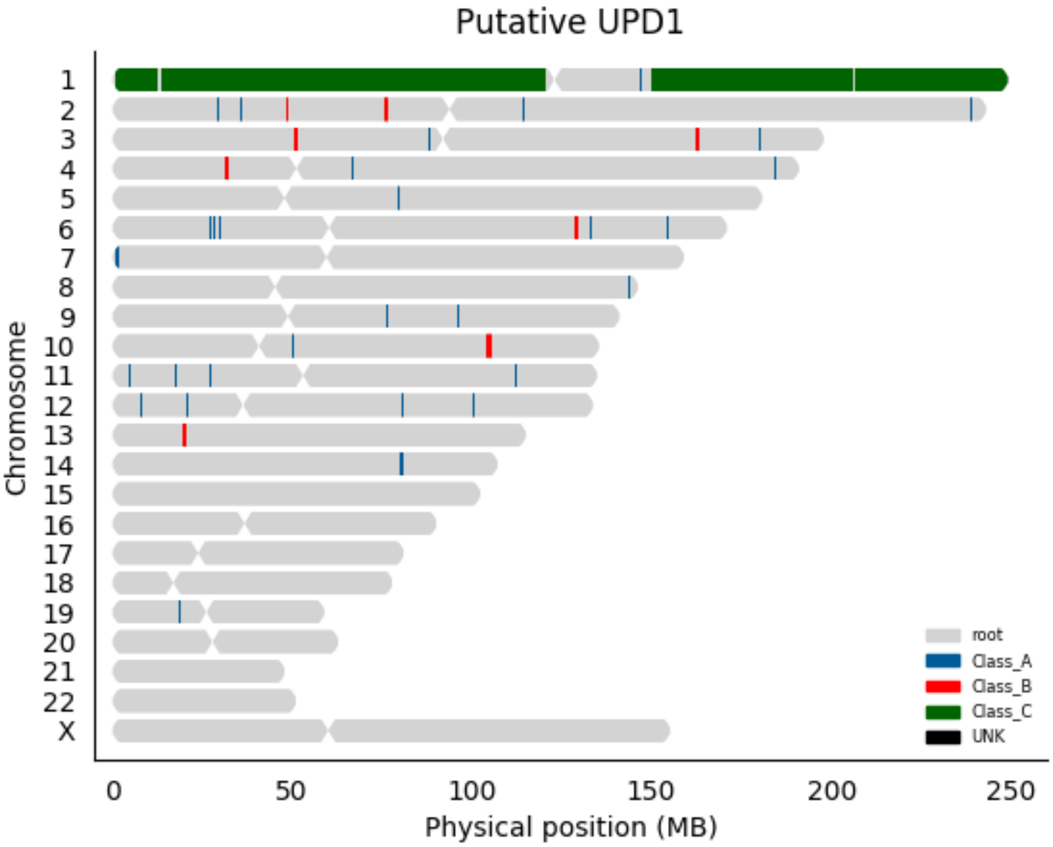

Putative UPD1
Chromosome
X
0
50
100
150
200
250
Physical position (MB)
root
Class_A
Class_B
Class_C
UNK

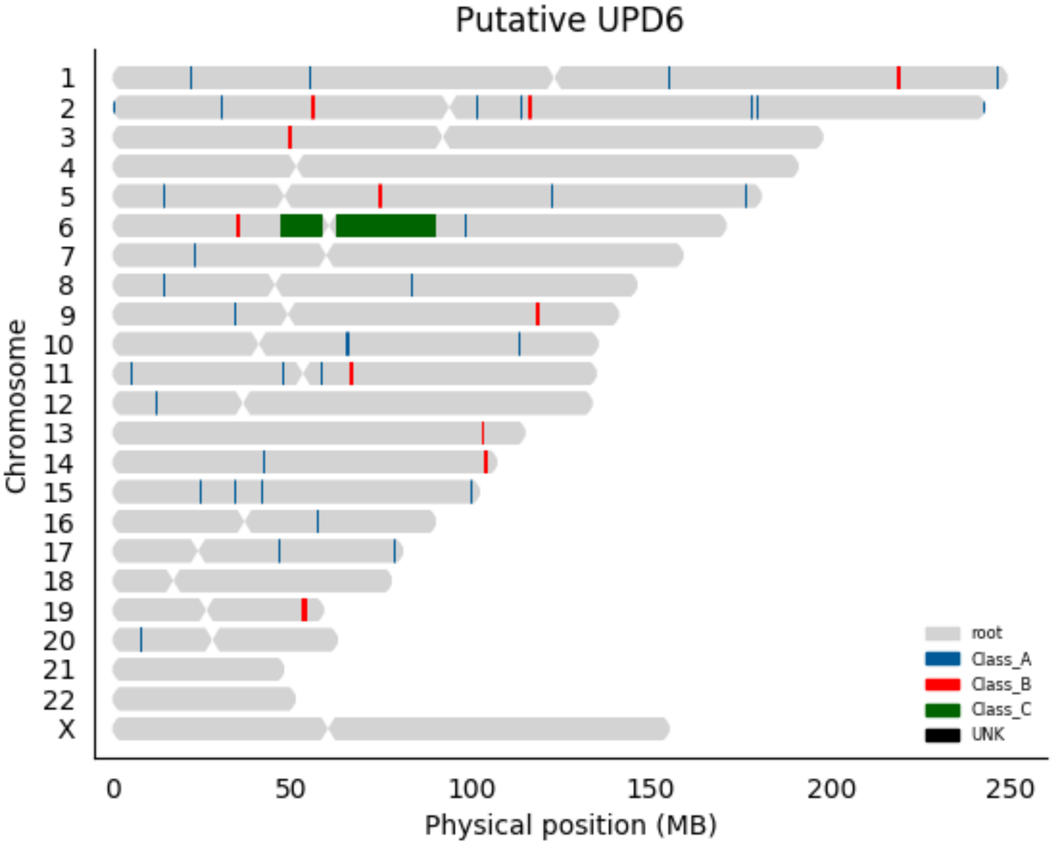

Putative UPD6
Chromosome
X
0
50
100
150
200
250
Physical position (MB)
root
Class_A
Class_B
Class_C
UNK

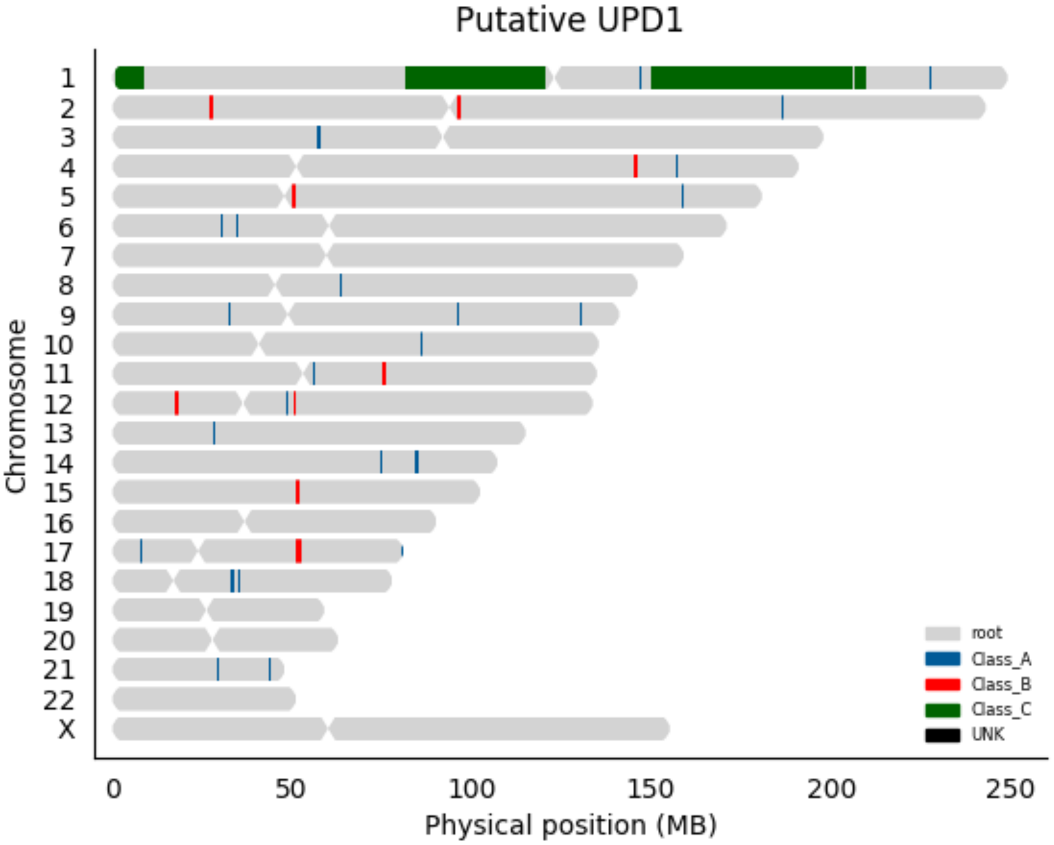

Putative UPD1
Chromosome
X
0
50
100
150
200
250
Physical position (MB)
root
Class_A
Class_B
Class_C
UNK

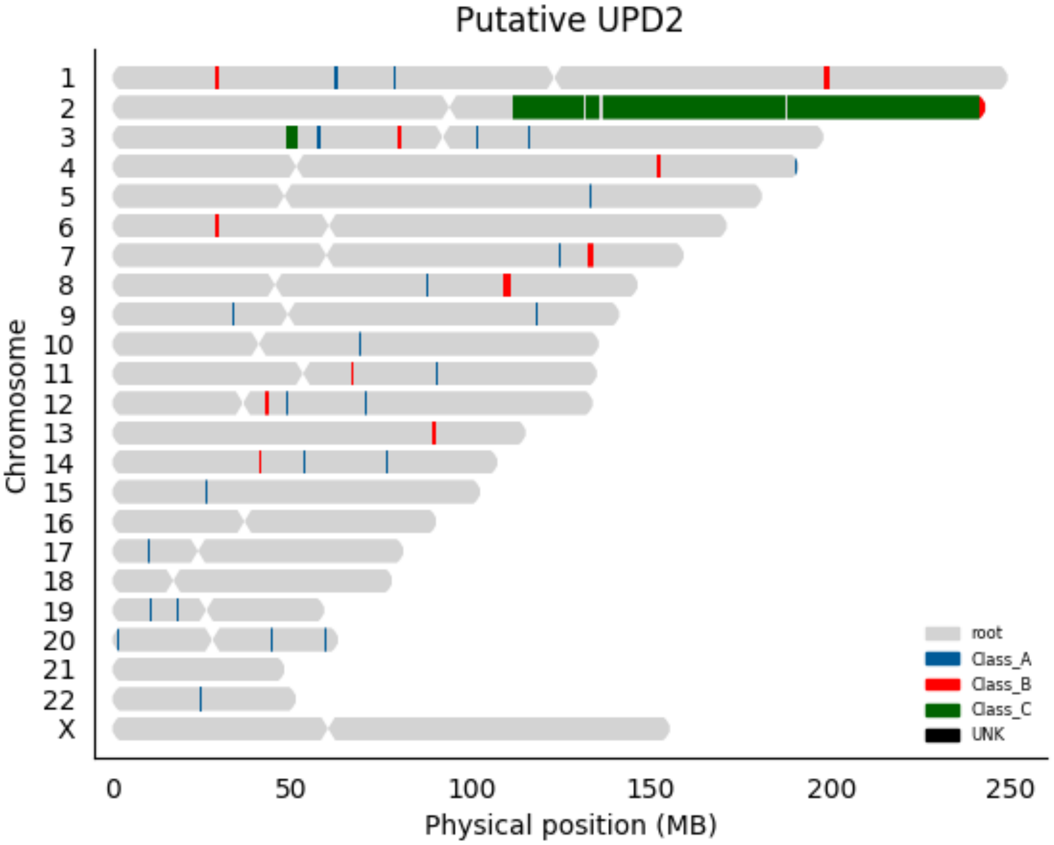

Putative UPD2
Chromosome
X
0
50
100
150
200
250
Physical position (MB)
root
Class_A
Class_B
Class_C
UNK

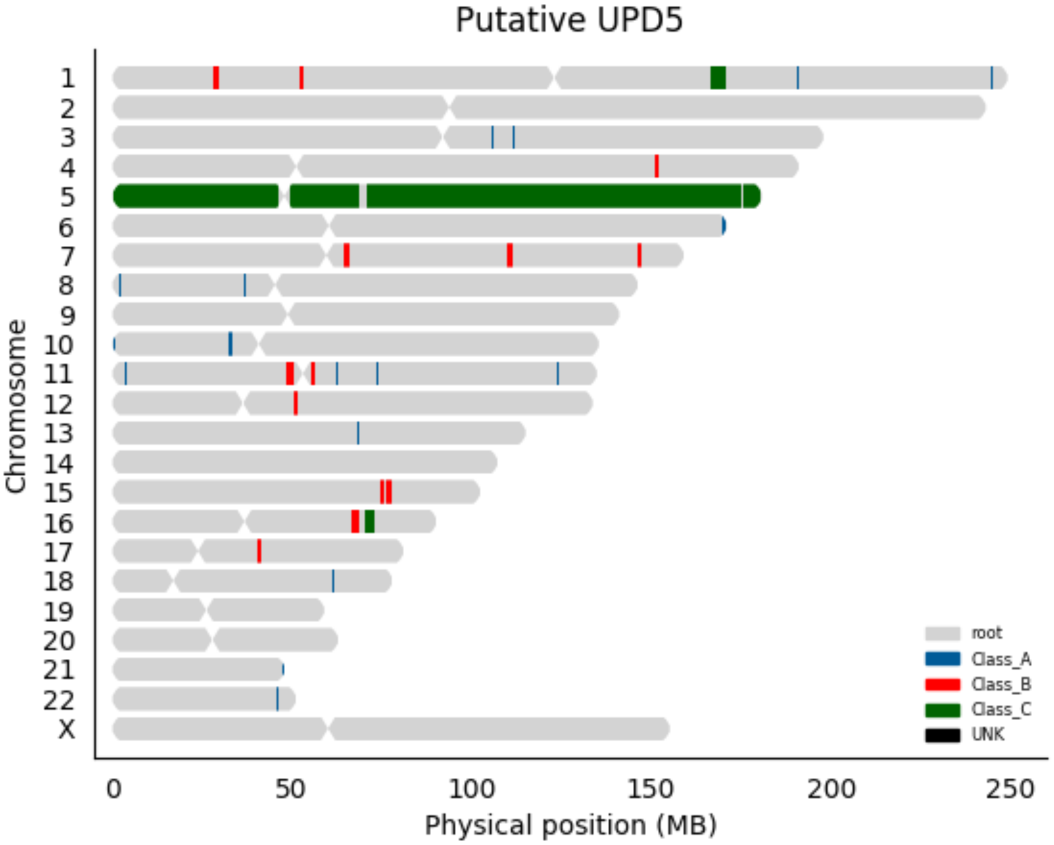
Putative UPD5
Chromosome
X
0
50
100
150
200
250
Physical position (MB)
root
Class_A
Class_B
Class_C
UNK

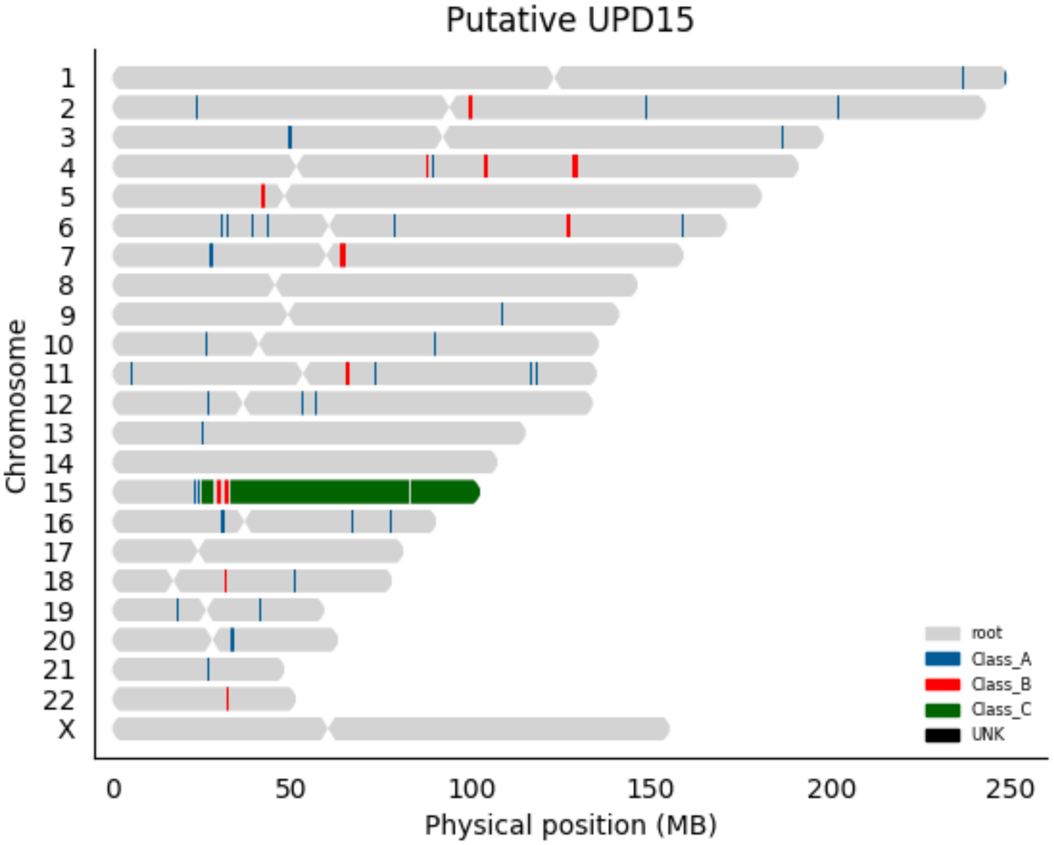
Putative UPD15
Chromosome
X
0
50
100
150
200
250
Physical position (MB)
root
Class_A
Class_B
Class_C
UNK

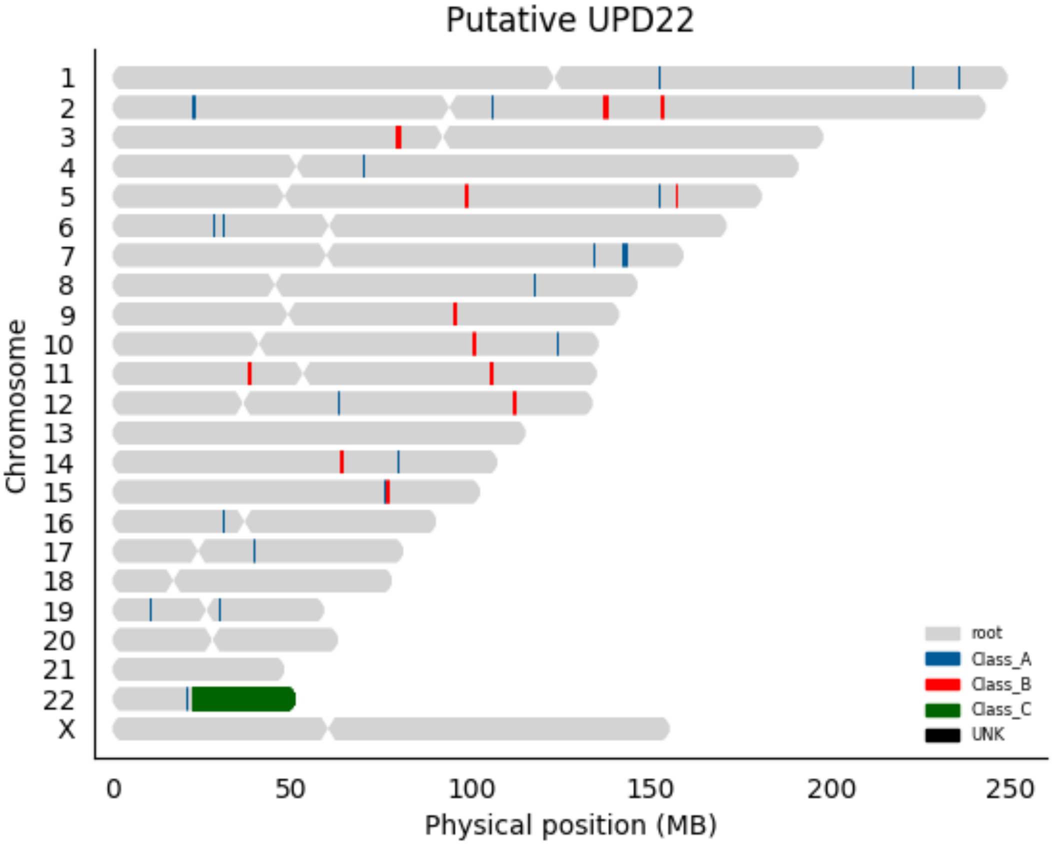
Putative UPD22
Chromosome
X
0
50
100
150
200
250
Physical position (MB)
root
Class_A
Class_B
Class_C
UNK

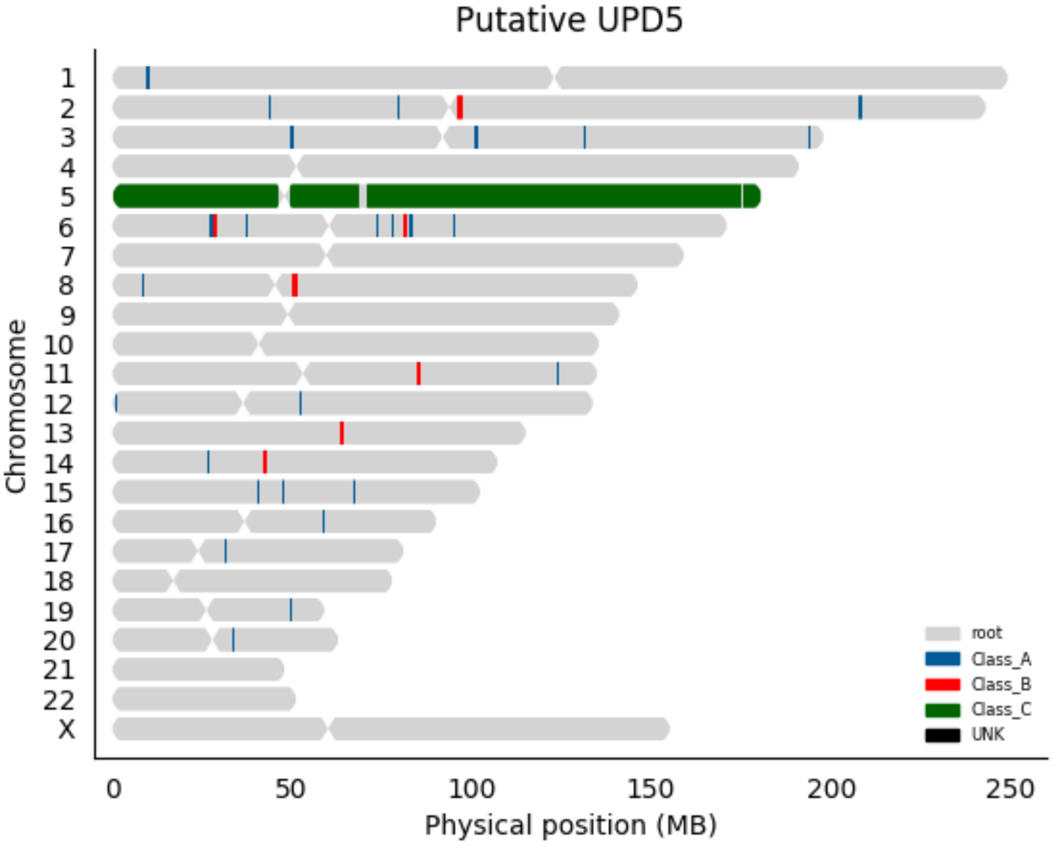
Putative UPD5
Chromosome
X
0
50
100
150
200
250
Physical position (MB)
root
Class_A
Class_B
Class_C
UNK

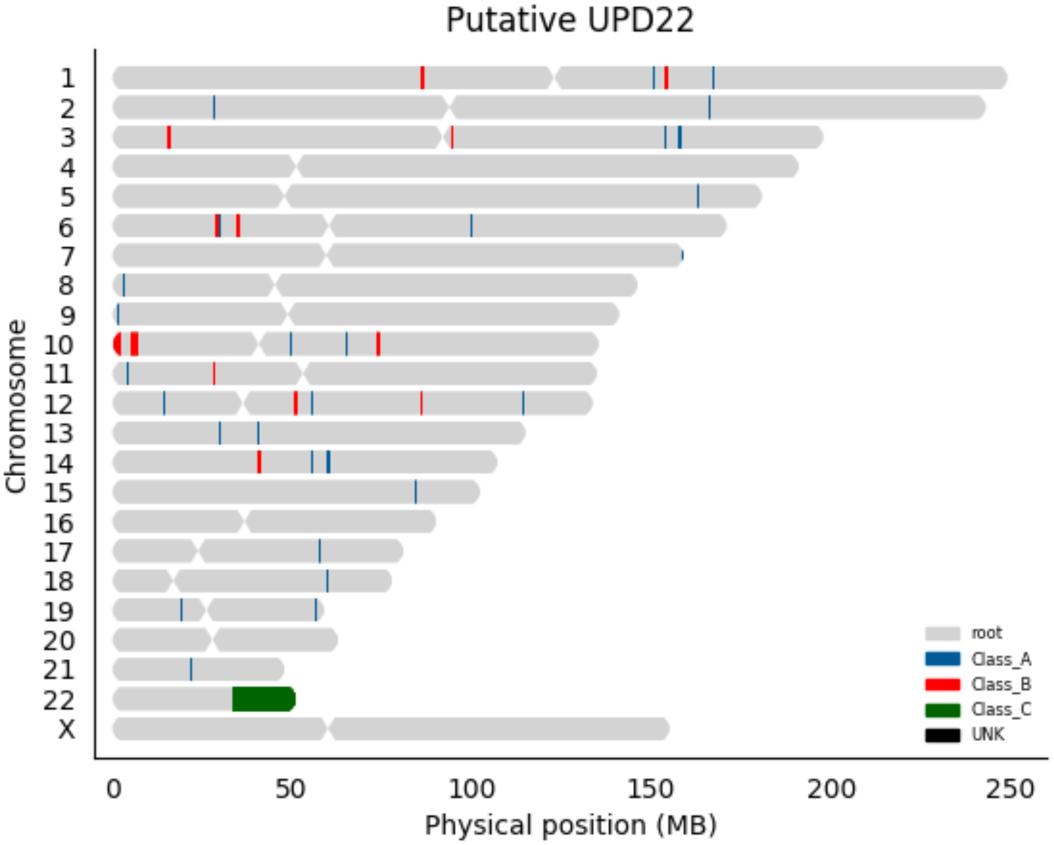

Putative UPD22
Chromosome
X
0
50
100
150
200
250
Physical position (MB)
root
Class_A
Class_B
Class_C
UNK

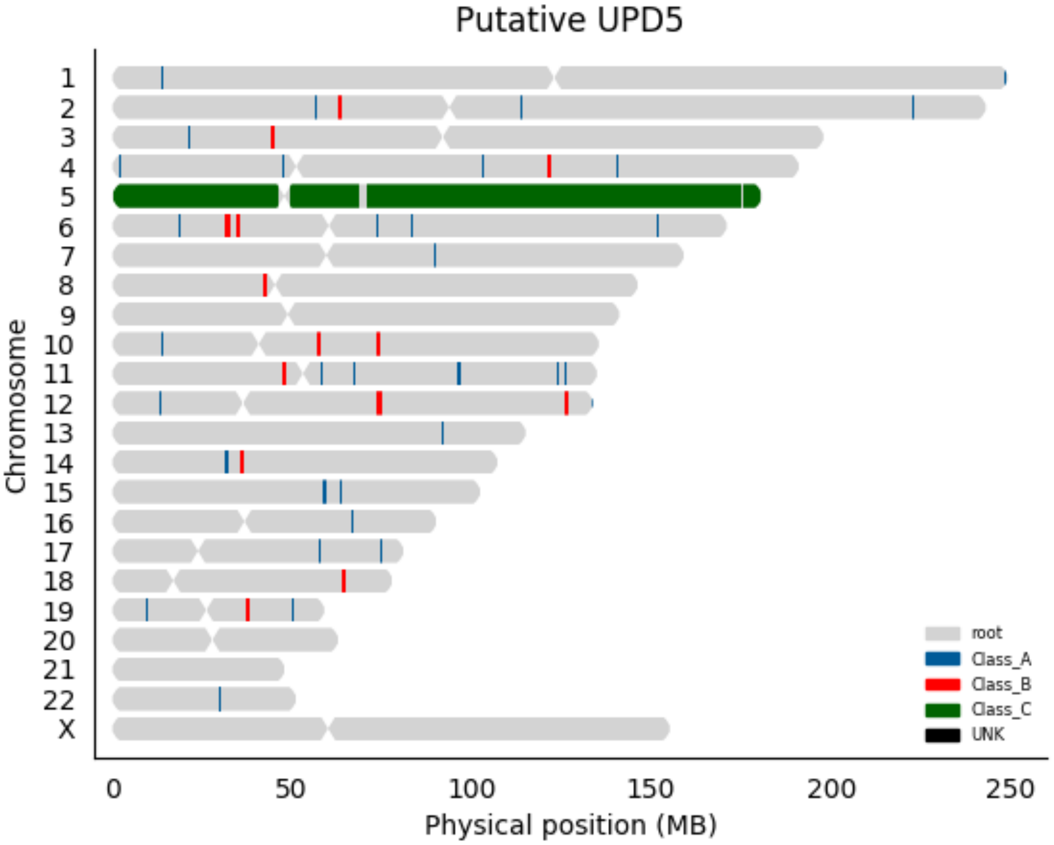

Putative UPD5
Chromosome
X
0
50
100
150
200
250
Physical position (MB)
root
Class_A
Class_B
Class_C
UNK

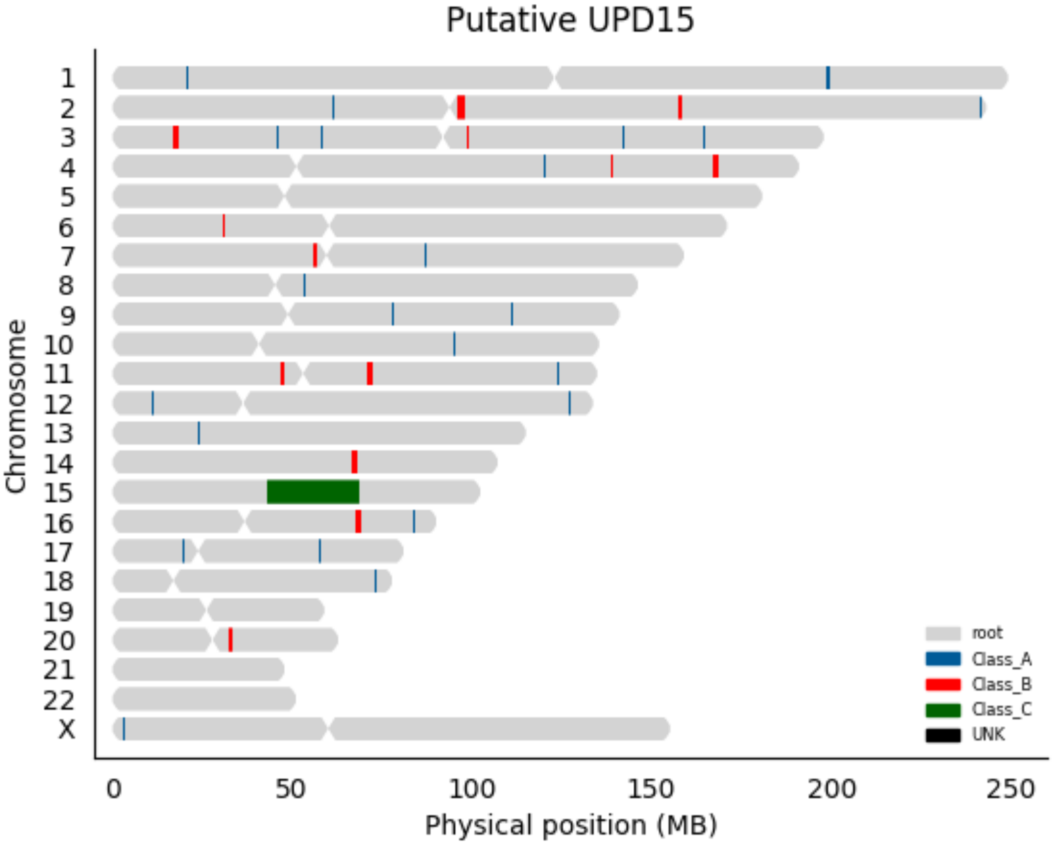

Putative UPD15
Chromosome
X
0
50
100
150
200
250
Physical position (MB)
root
Class_A
Class_B
Class_C
UNK

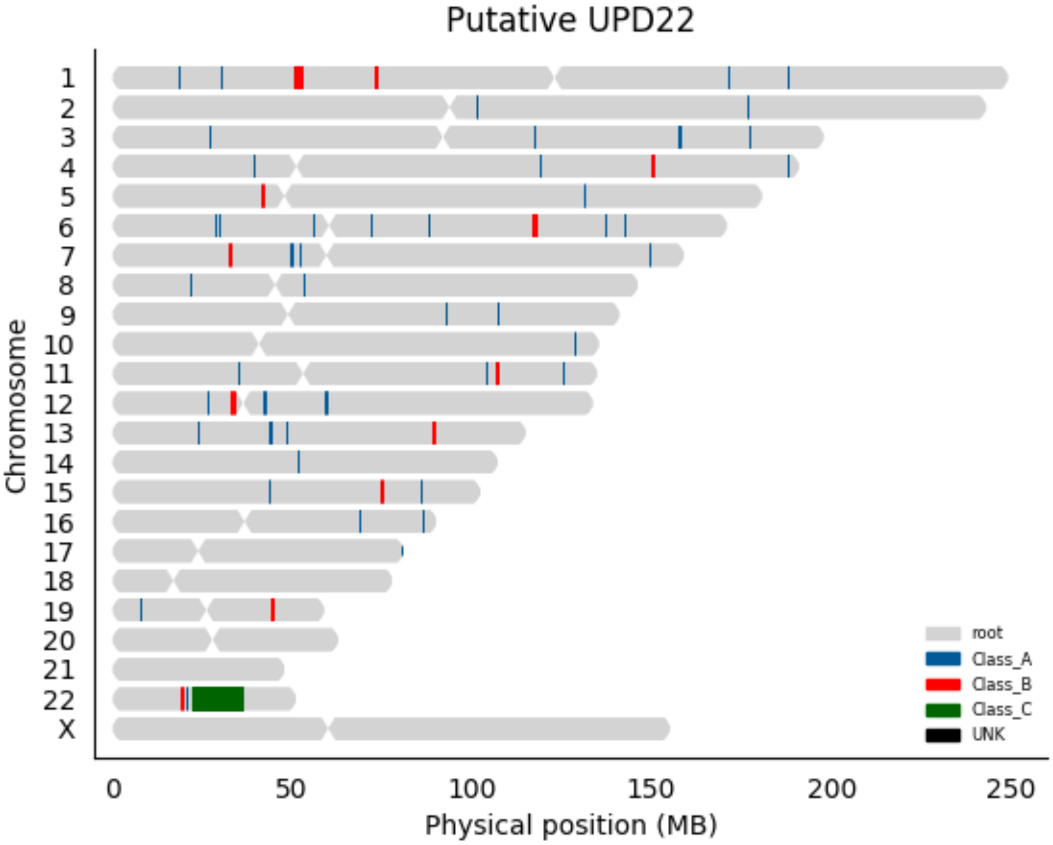

Putative UPD22
Chromosome
X
0
50
100
150
200
250
Physical position (MB)
root
Class_A
Class_B
Class_C
UNK

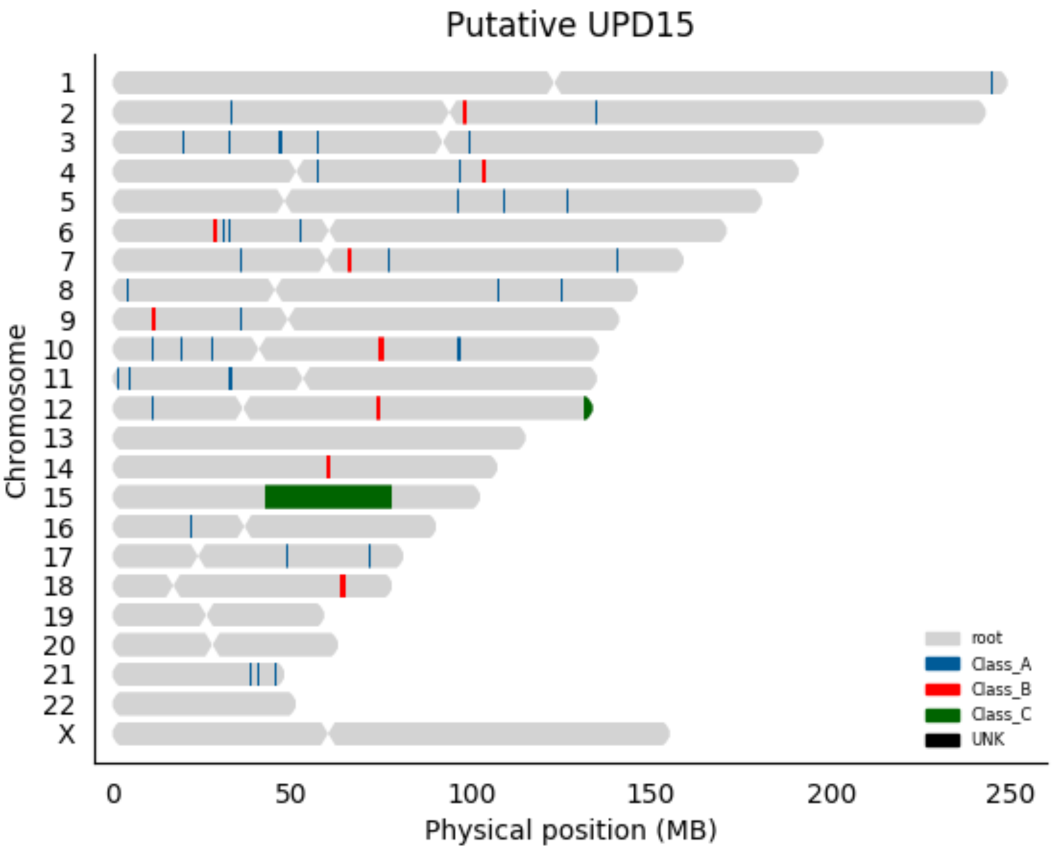

Putative UPD15
Chromosome
X
0
50
100
150
200
250
Physical position (MB)
root
Class_A
Class_B
Class_C
UNK

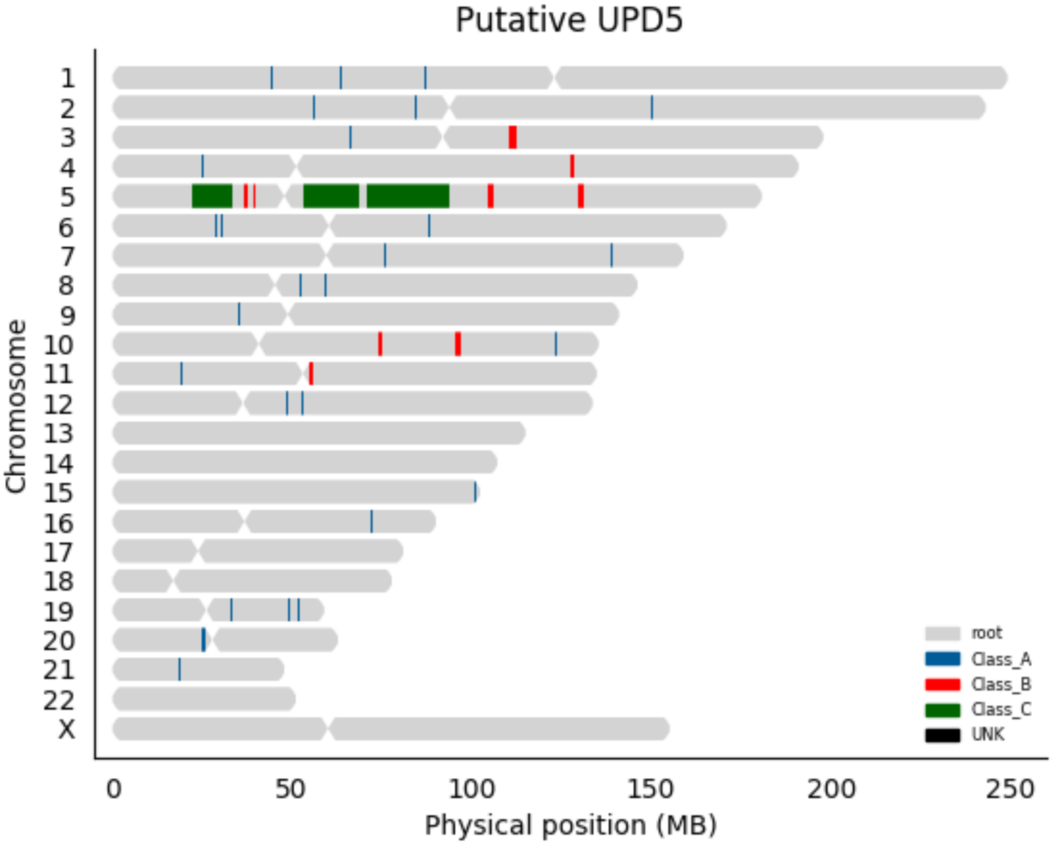

Putative UPD5
Chromosome
X
0
50
100
150
200
250
Physical position (MB)
root
Class_A
Class_B
Class_C
UNK

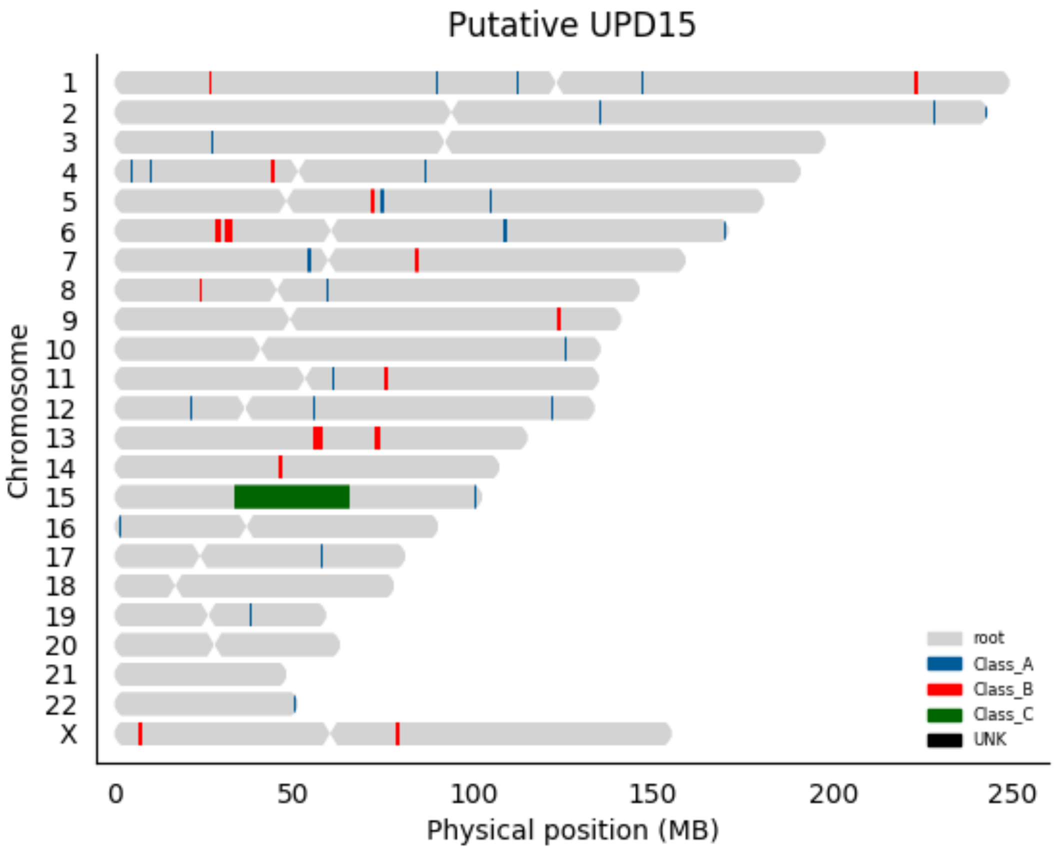

Putative UPD15
Chromosome
X
0
50
100
150
200
250
Physical position (MB)
root
Class_A
Class_B
Class_C
UNK

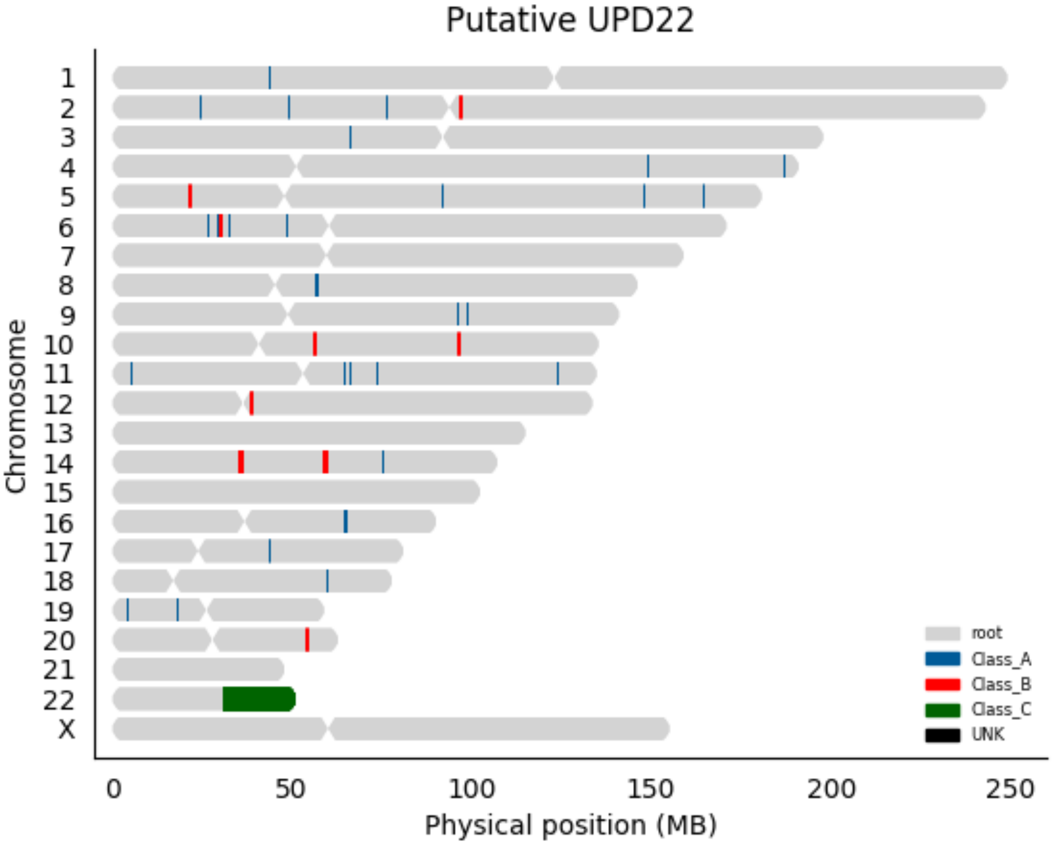

Putative UPD22
Chromosome
X
0
50
100
150
200
250
Physical position (MB)
root
Class_A
Class_B
Class_C
UNK

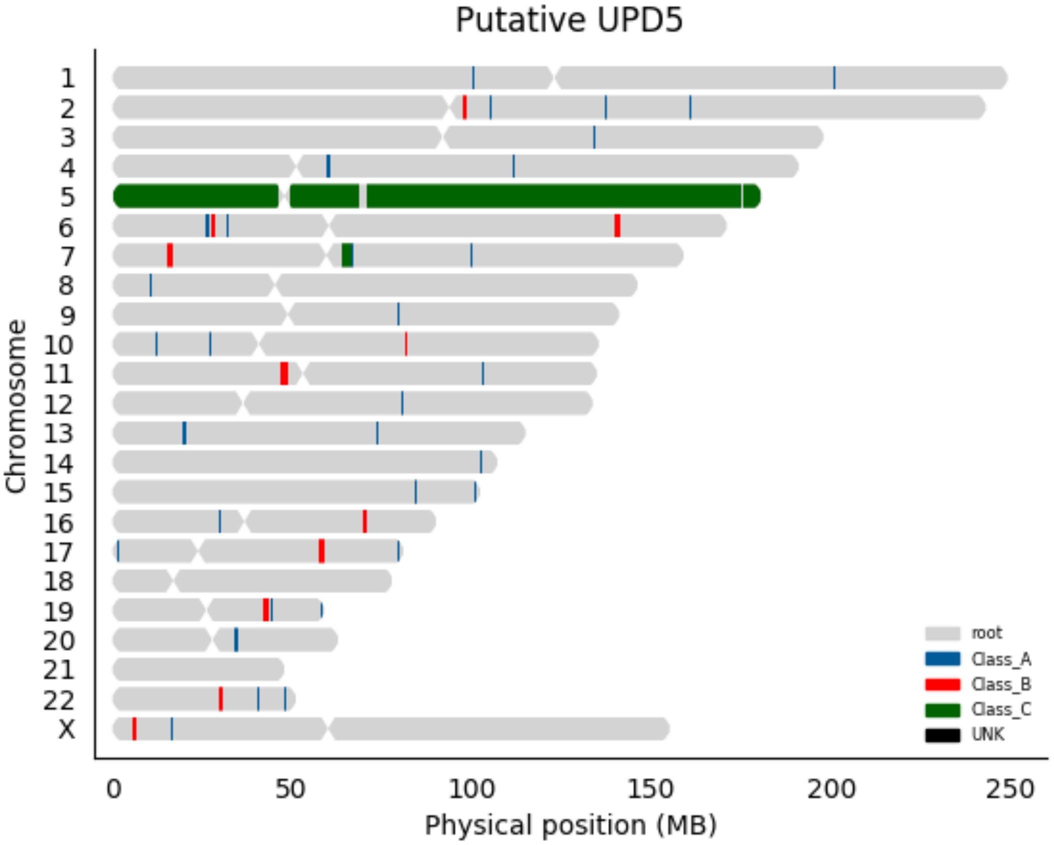
Putative UPD5
Chromosome
X
0
50
100
150
200
250
Physical position (MB)
root
Class_A
Class_B
Class_C
UNK

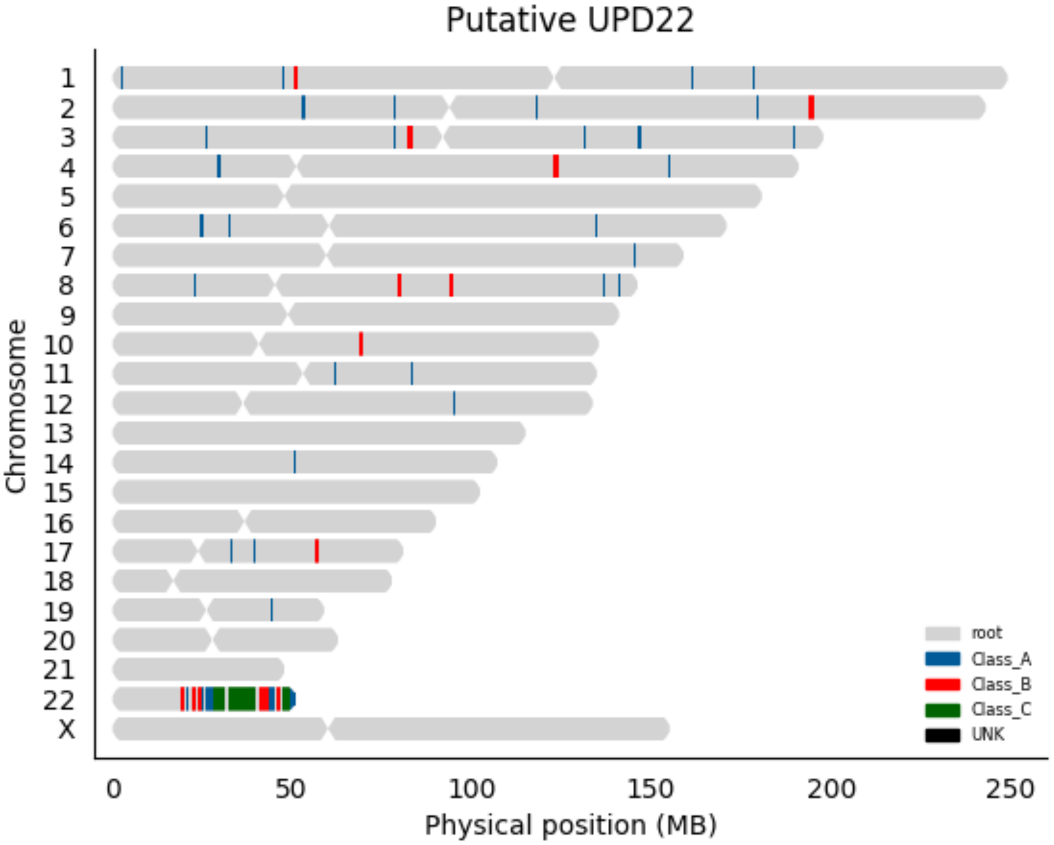

Putative UPD22
Chromosome
X
0
50
100
150
200
250
Physical position (MB)
root
Class_A
Class_B
Class_C
UNK

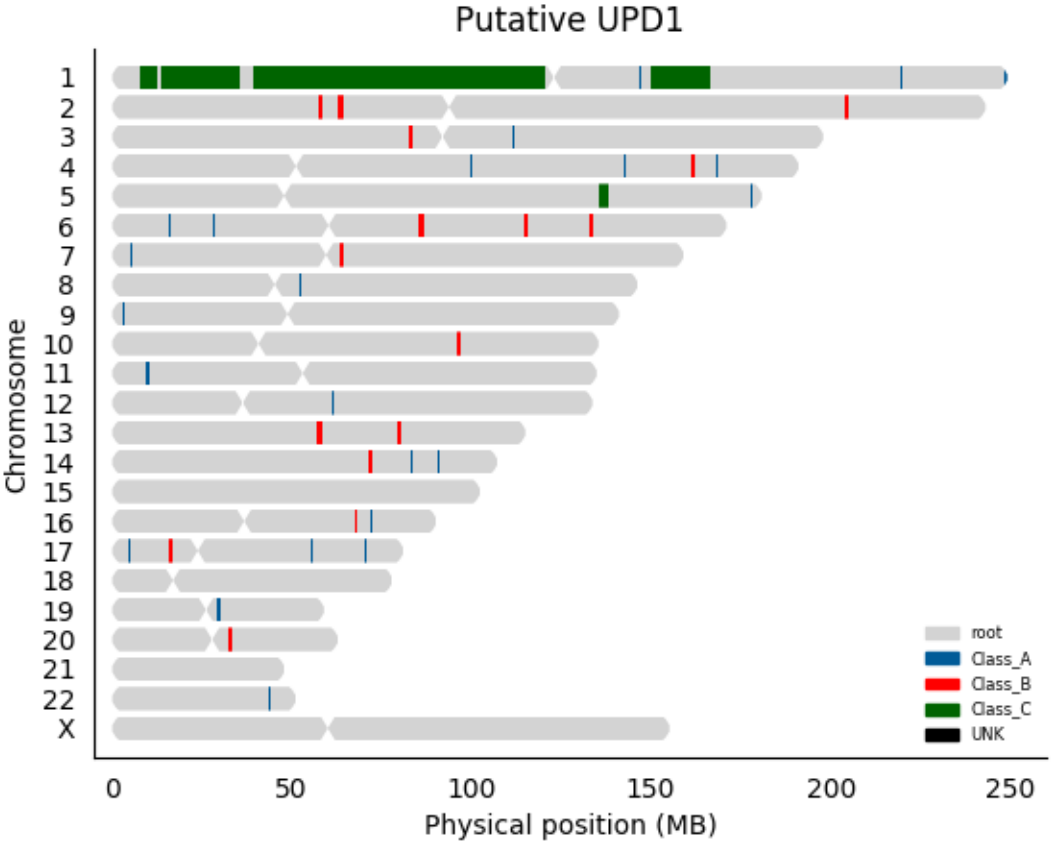

Putative UPD1
Chromosome
X
0
50
100
150
200
250
Physical position (MB)
root
Class_A
Class_B
Class_C
UNK

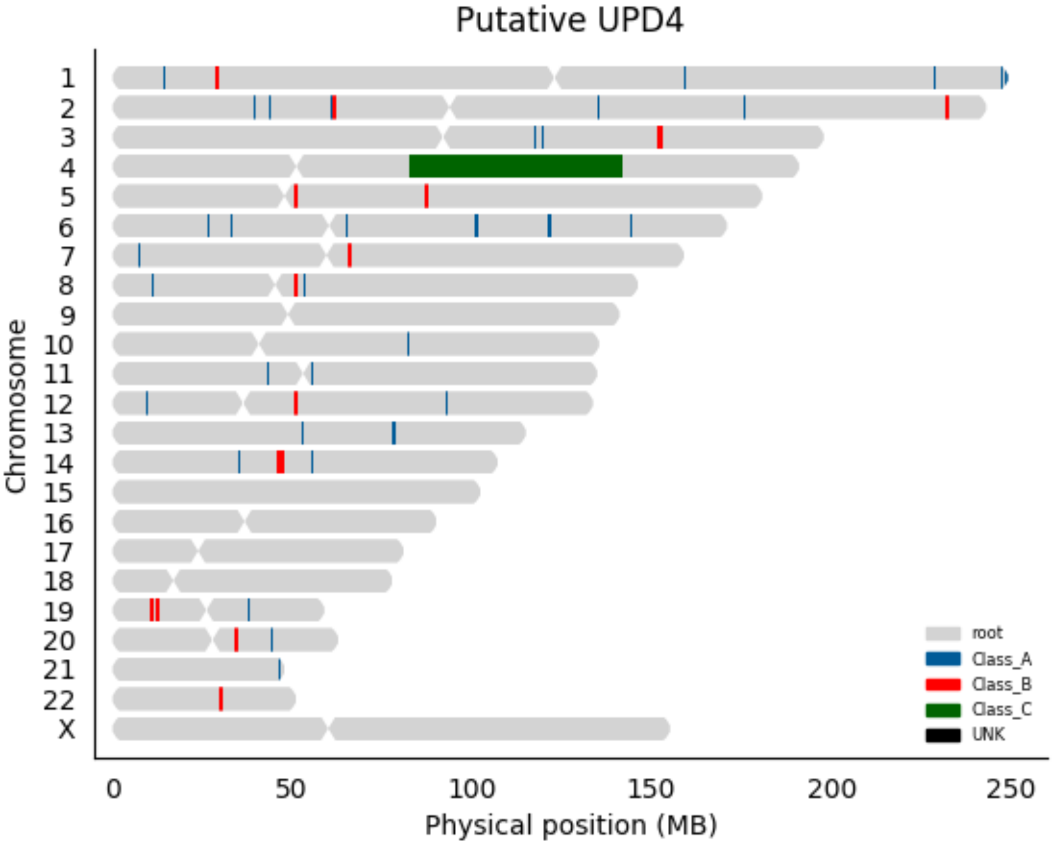
Putative UPD4
Chromosome
X
0
50
100
150
200
250
Physical position (MB)
root
Class_A
Class_B
Class_C
UNK

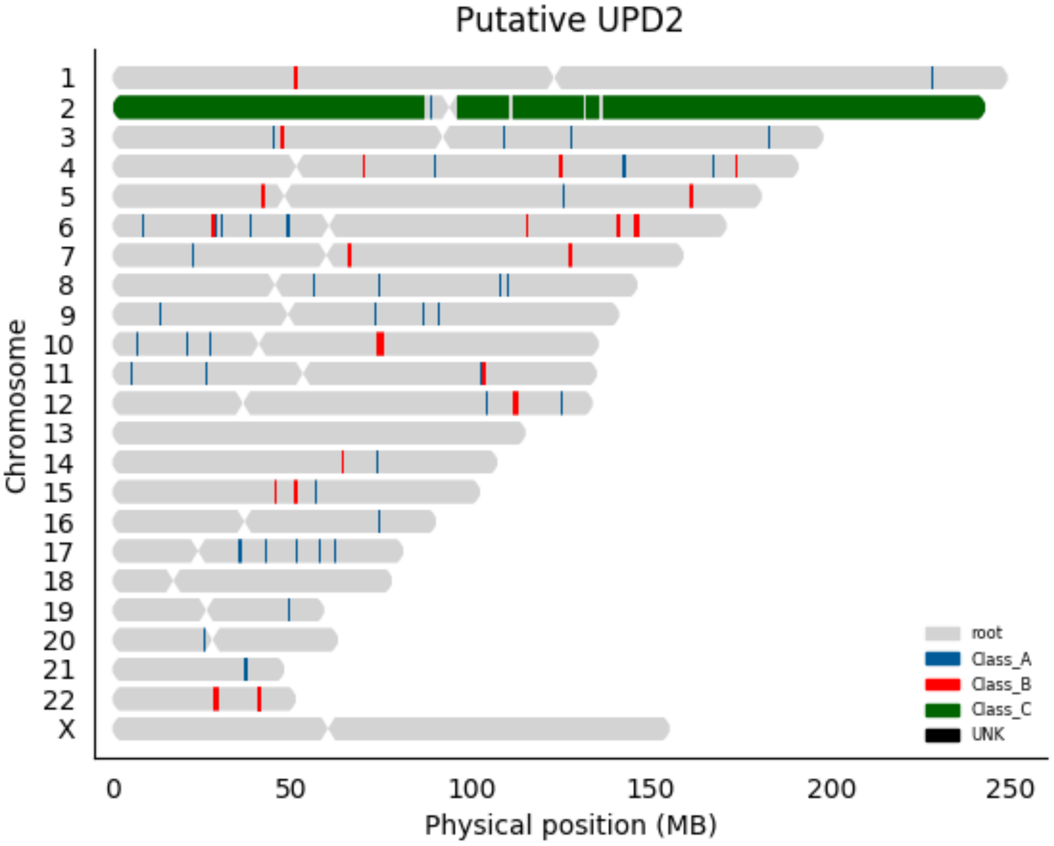

Putative UPD2
Chromosome
X
0
50
100
150
200
250
Physical position (MB)
root
Class_A
Class_B
Class_C
UNK

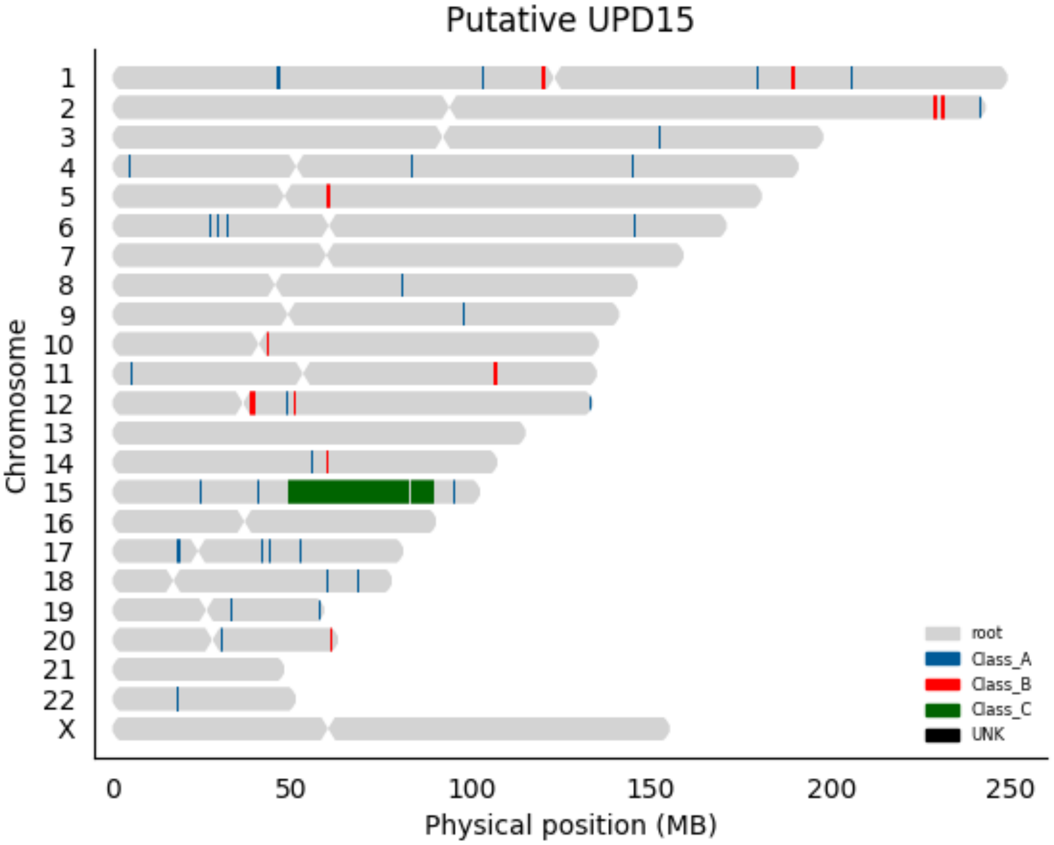

Putative UPD15
Chromosome
X
0
50
100
150
200
250
Physical position (MB)
root
Class_A
Class_B
Class_C
UNK

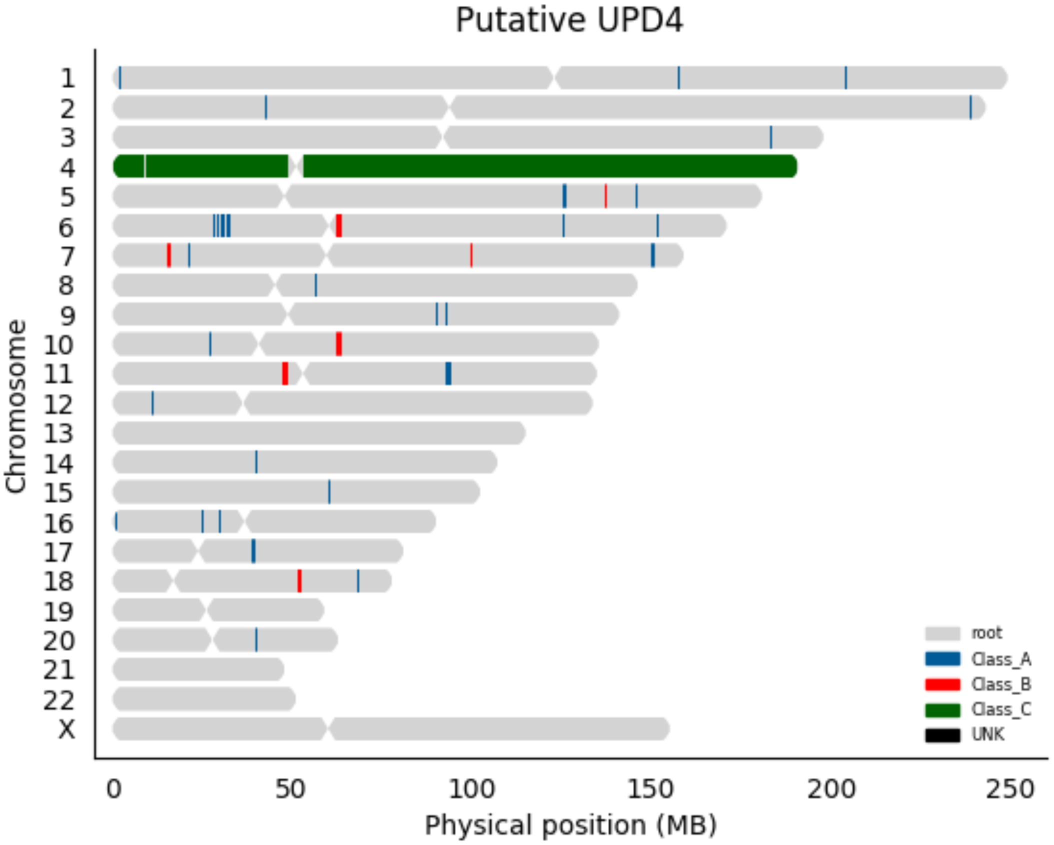
Putative UPD4
Chromosome
X
root
Class_A
Class_B
Class_C
UNK
0
50
100
150
200
250
Physical position (MB)

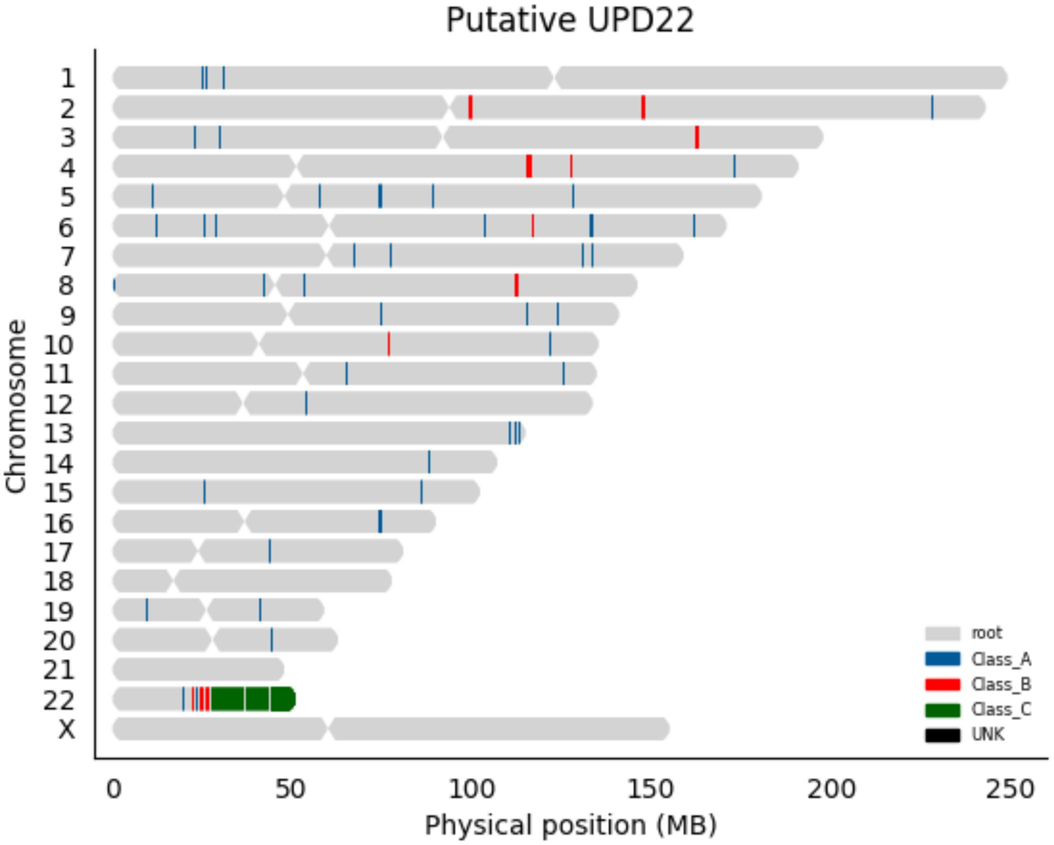

Putative UPD22
Chromosome
X
0
50
100
150
200
250
Physical position (MB)
root
Class_A
Class_B
Class_C
UNK

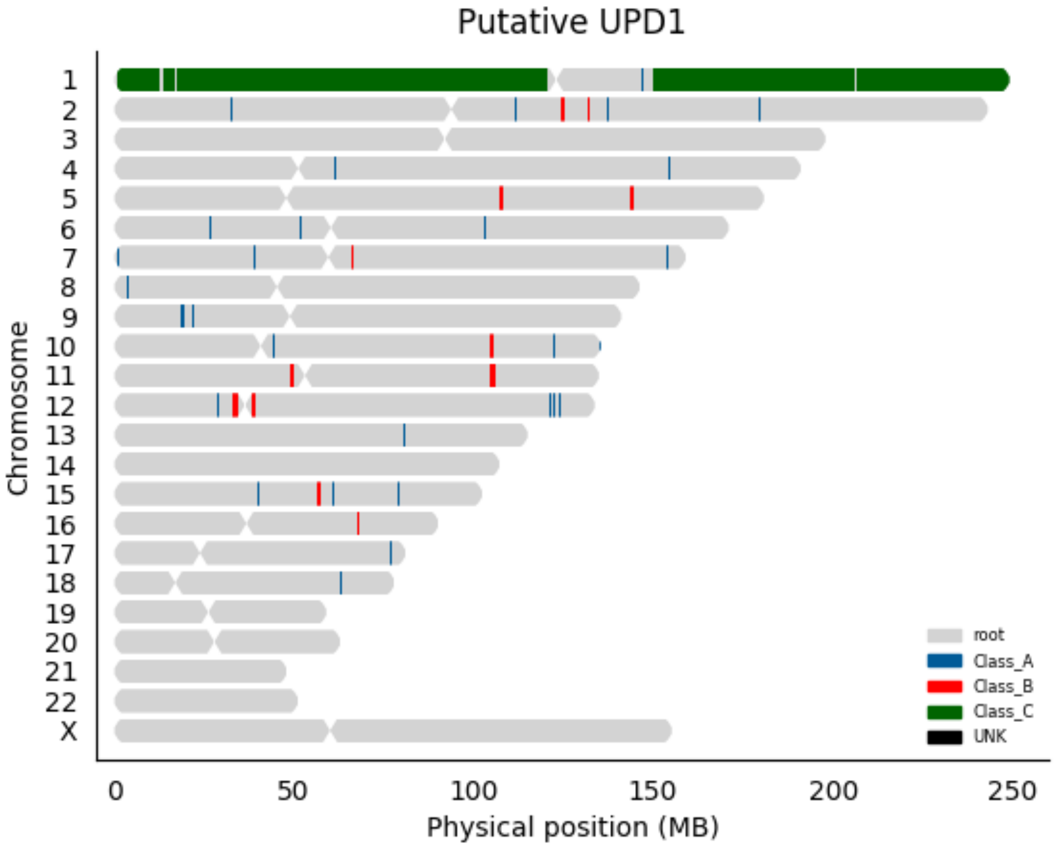
Putative UPD1
Chromosome
X
0
50
100
150
200
250
Physical position (MB)
root
Class_A
Class_B
Class_C
UNK

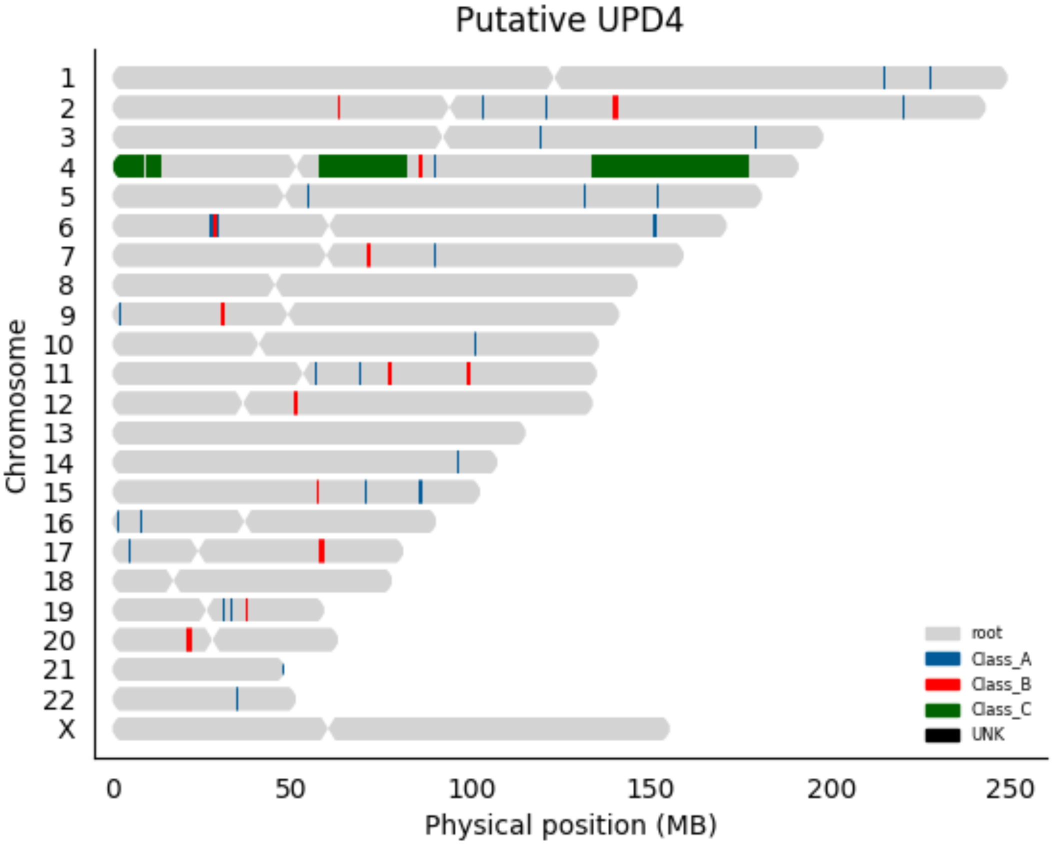
Putative UPD4
Chromosome
X
0
50
100
150
200
250
Physical position (MB)
root
Class_A
Class_B
Class_C
UNK

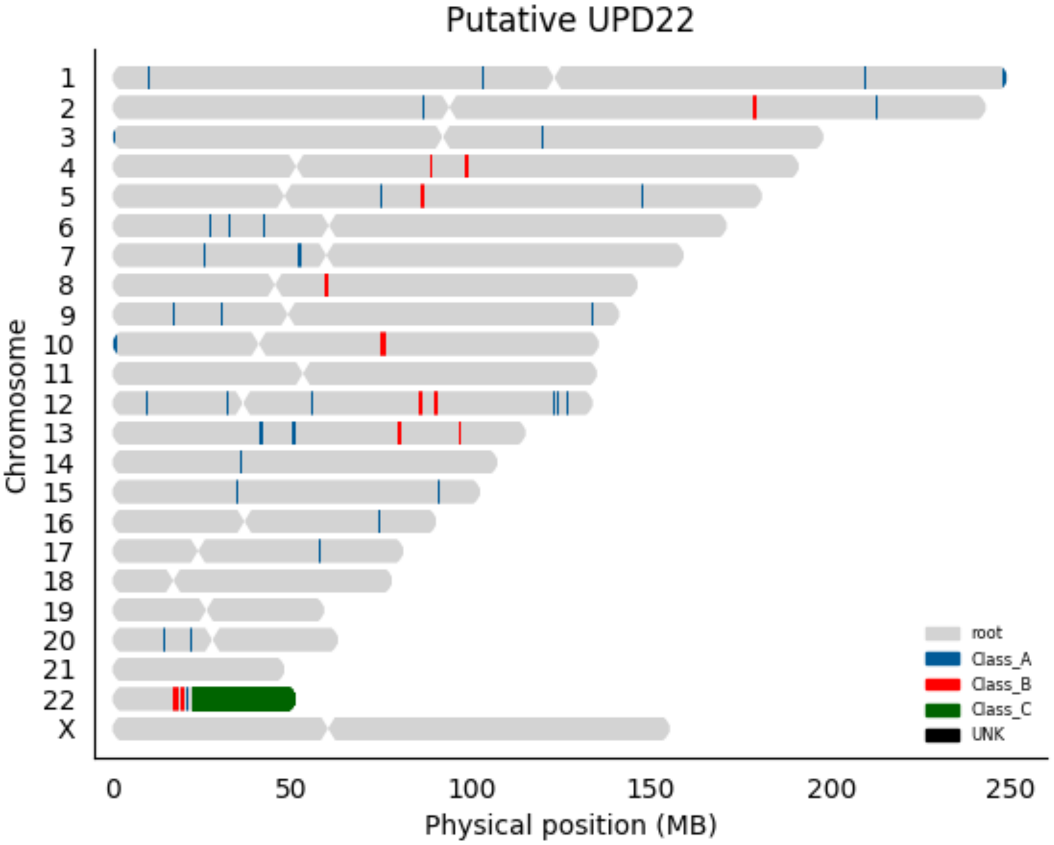

Putative UPD22
Chromosome
X
0
50
100
150
200
250
Physical position (MB)
root
Class_A
Class_B
Class_C
UNK

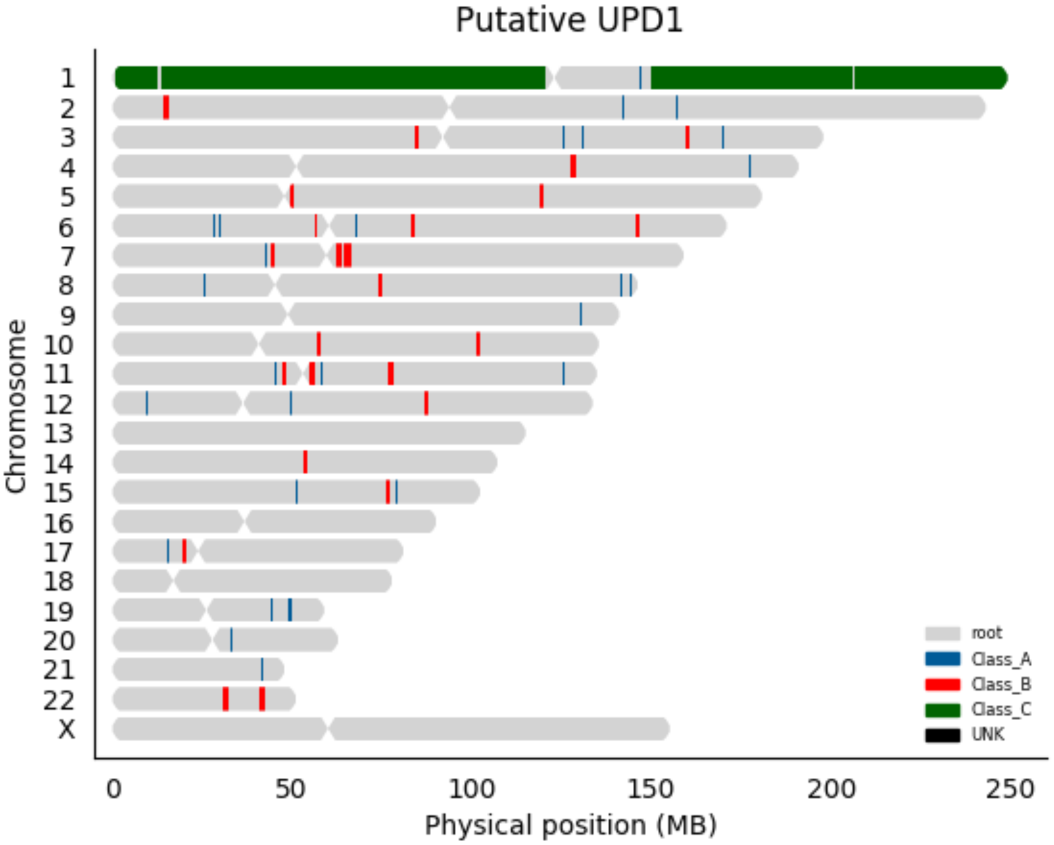

Putative UPD1
Chromosome
X
0
50
100
150
200
250
Physical position (MB)
root
Class_A
Class_B
Class_C
UNK

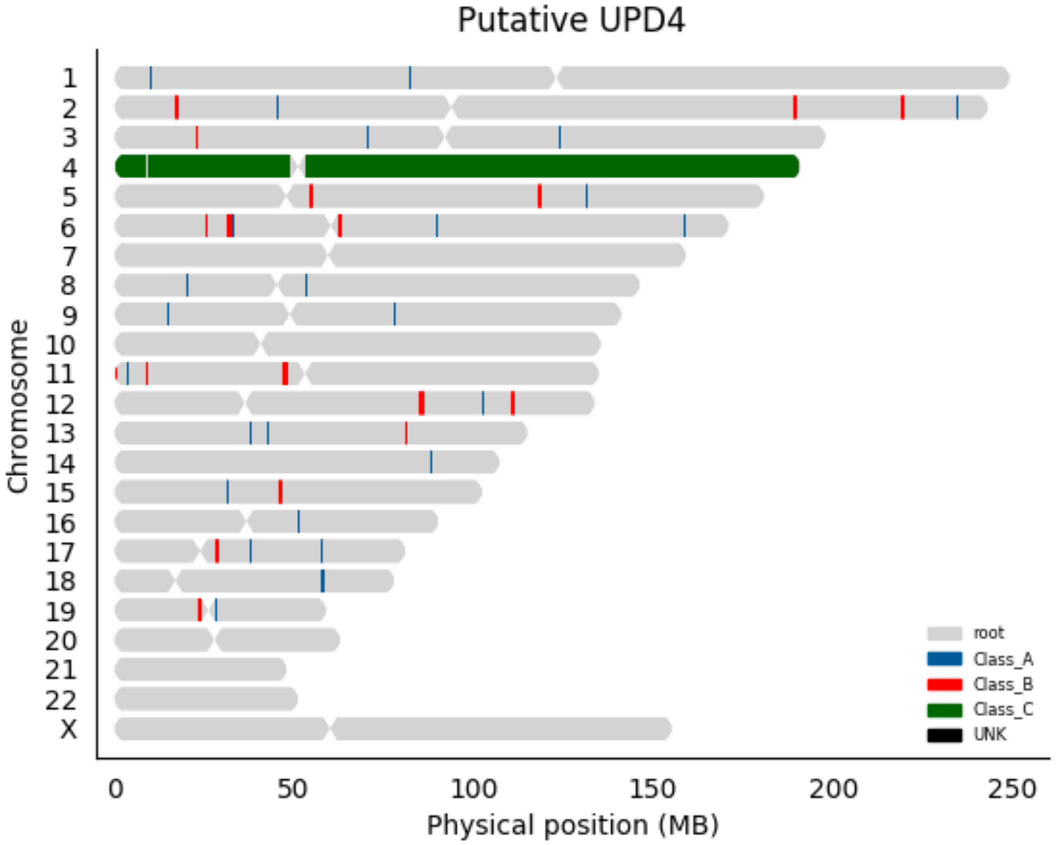

Putative UPD4
Chromosome
X
0
50
100
150
200
250
Physical position (MB)
root
Class_A
Class_B
Class_C
UNK

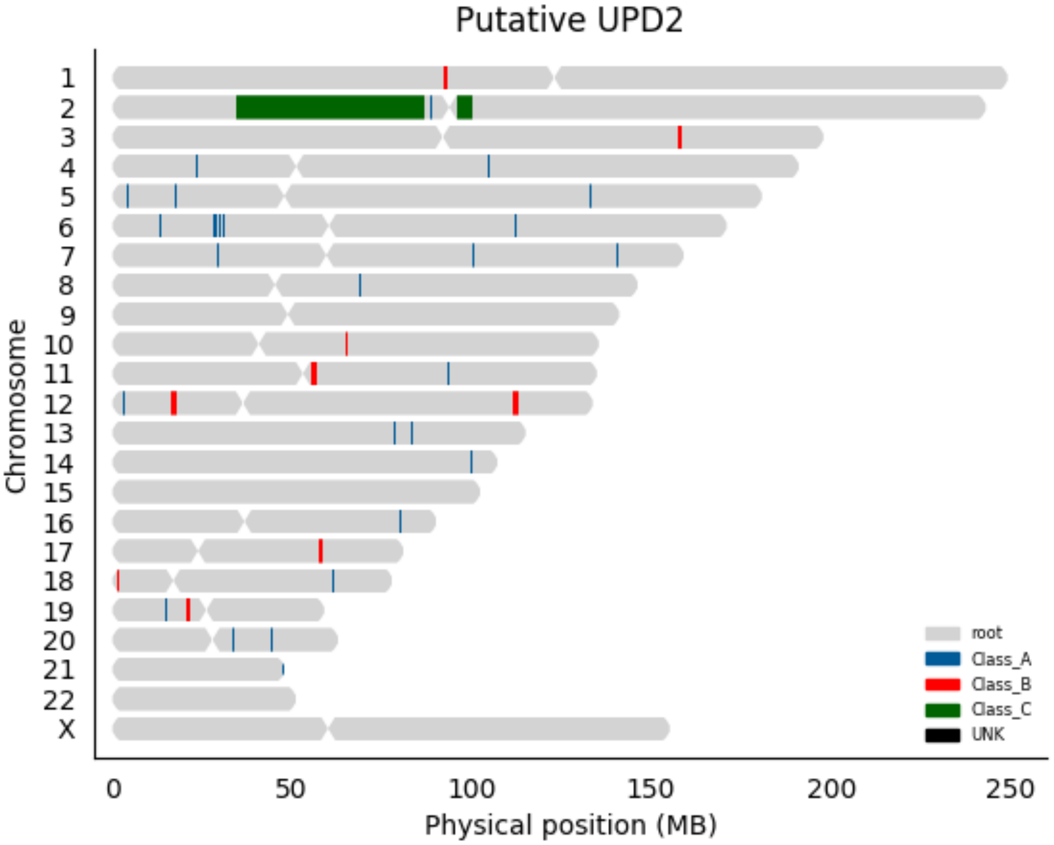

Putative UPD2
Chromosome
X
0
50
100
150
200
250
Physical position (MB)
root
Class_A
Class_B
Class_C
UNK

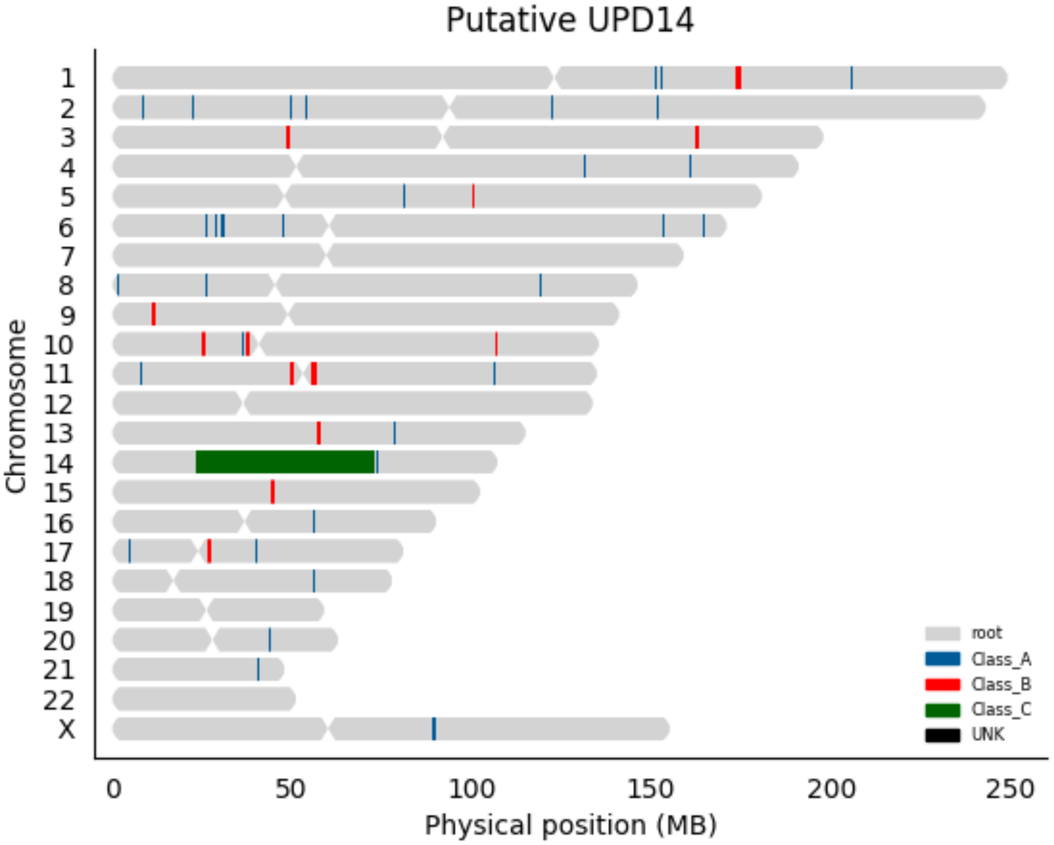

Putative UPD14
Chromosome
X
0
50
100
150
200
250
Physical position (MB)
root
Class_A
Class_B
Class_C
UNK

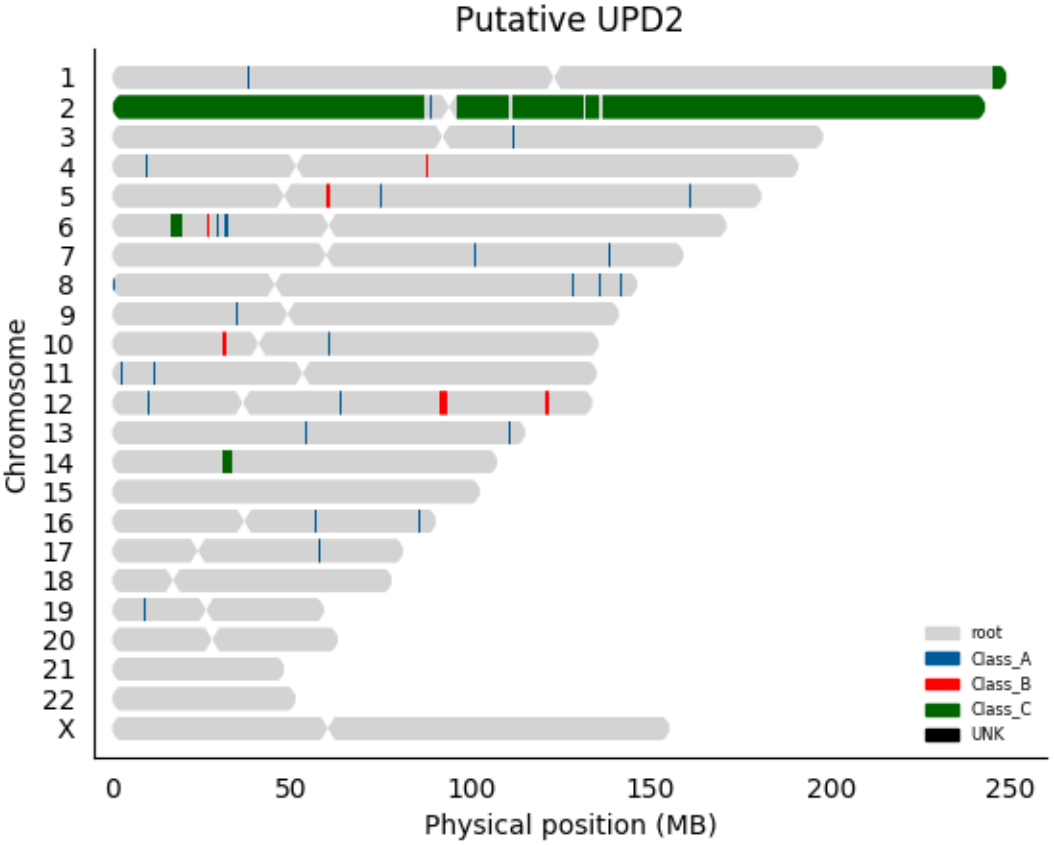

Putative UPD2
Chromosome
X
0
50
100
150
200
250
Physical position (MB)
root
Class_A
Class_B
Class_C
UNK

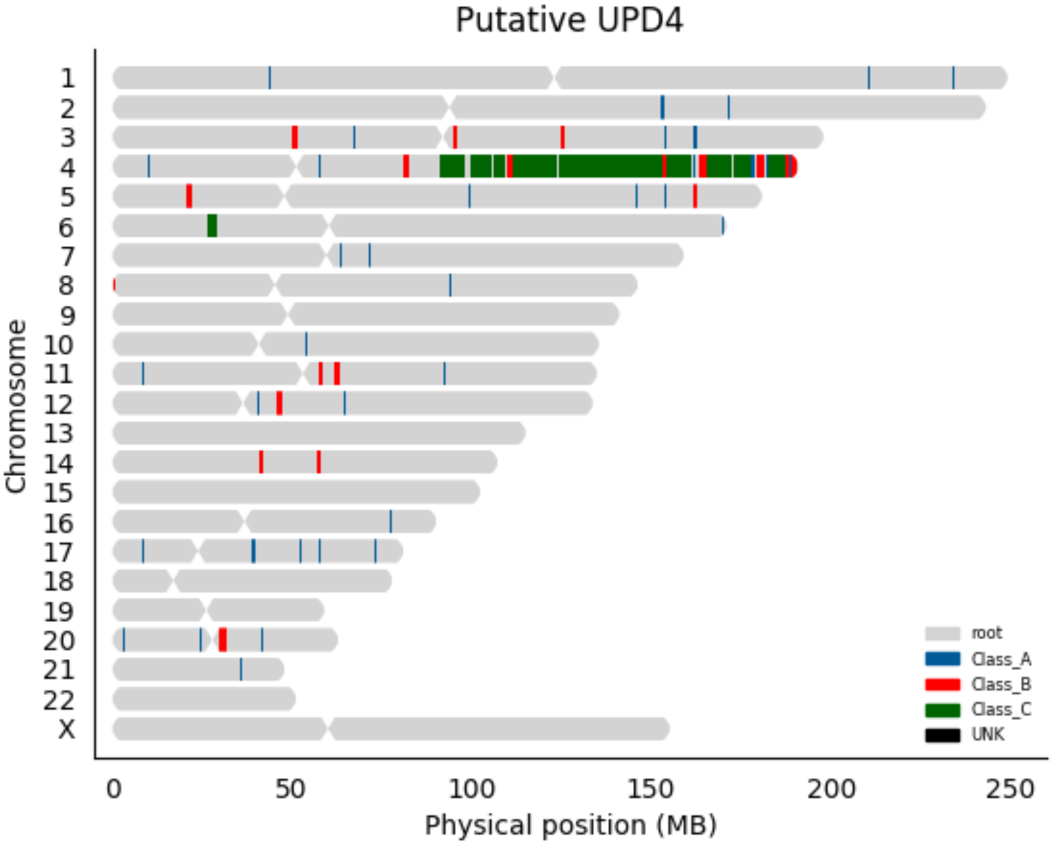

Putative UPD4
Chromosome
X
0
50
100
150
200
250
Physical position (MB)
root
Class_A
Class_B
Class_C
UNK

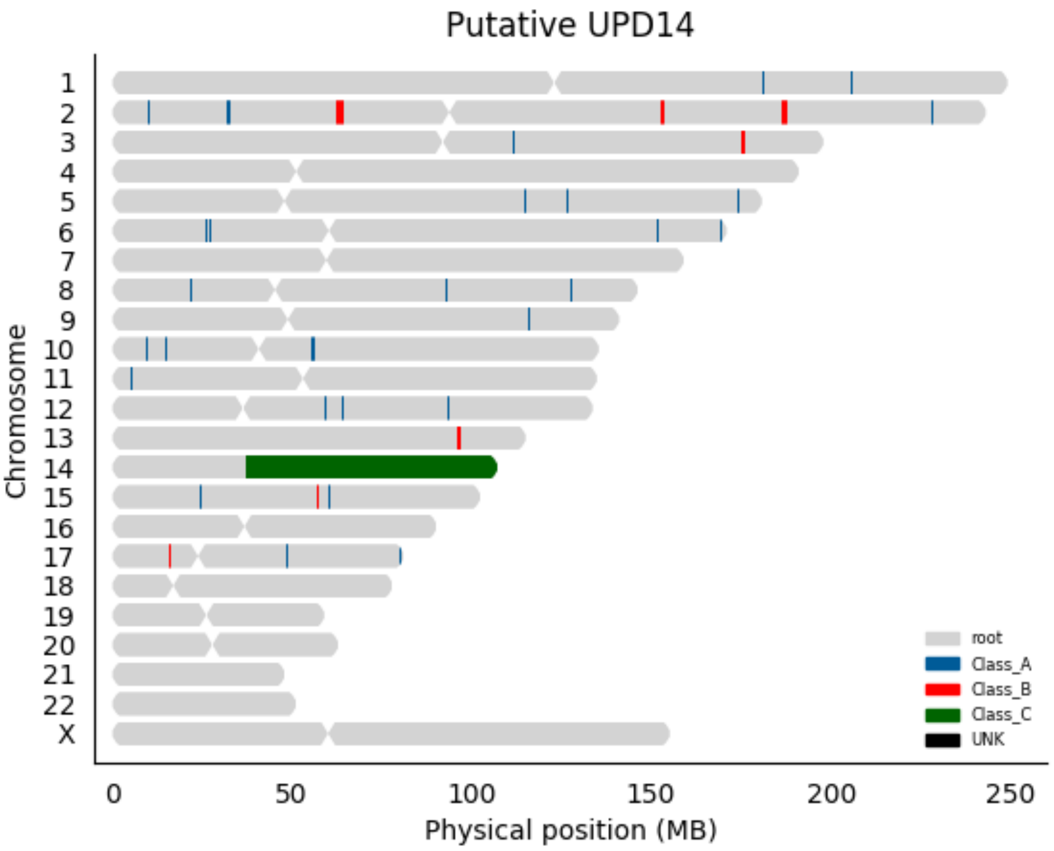

Putative UPD14
X
Chromosome
0
50
100
150
200
250
Physical position (MB)
root
Class_A
Class_B
Class_C
UNK

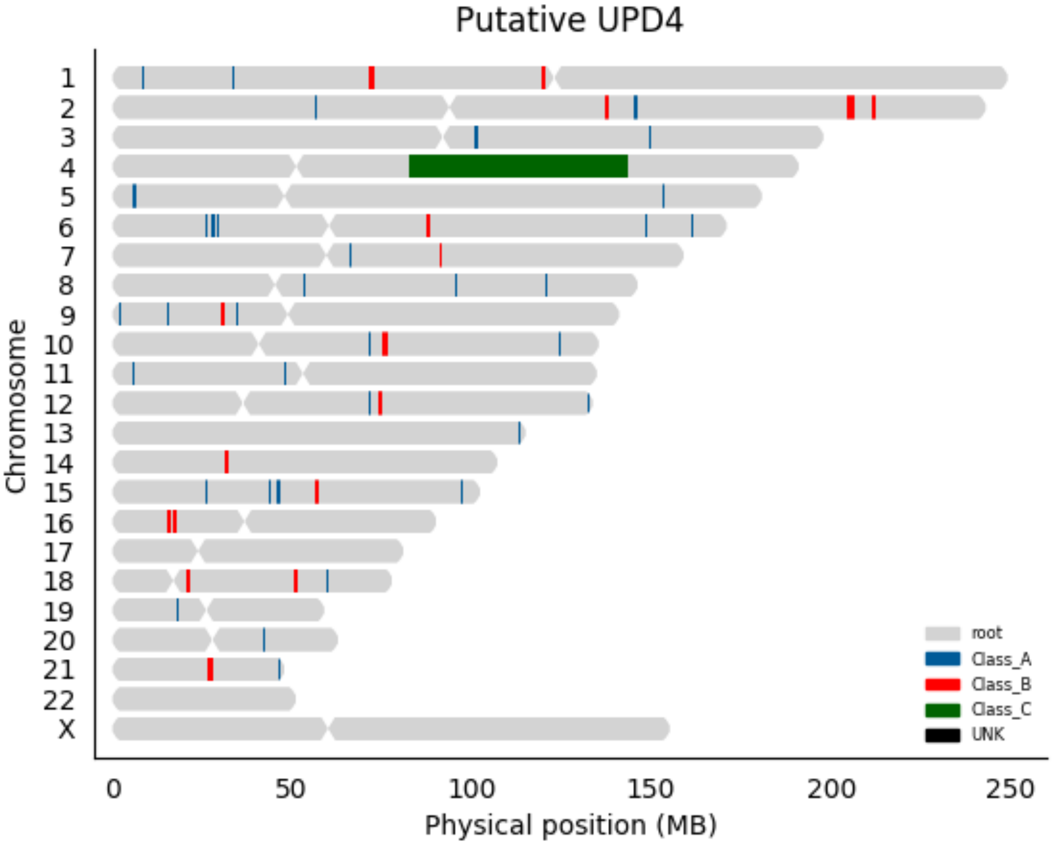

Putative UPD4
Chromosome
X
0
50
100
150
200
250
Physical position (MB)
root
Class_A
Class_B
Class_C
UNK

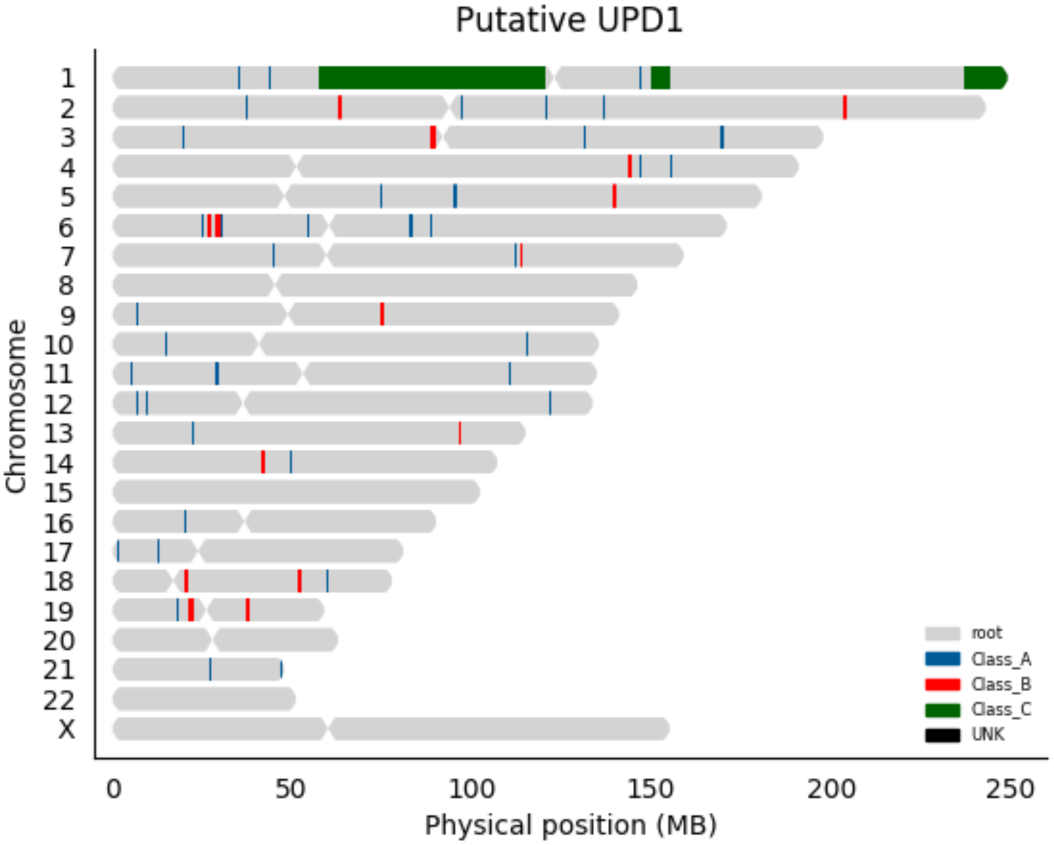

Putative UPD1
Chromosome
X
0
50
100
150
200
250
Physical position (MB)
root
Class_A
Class_B
Class_C
UNK

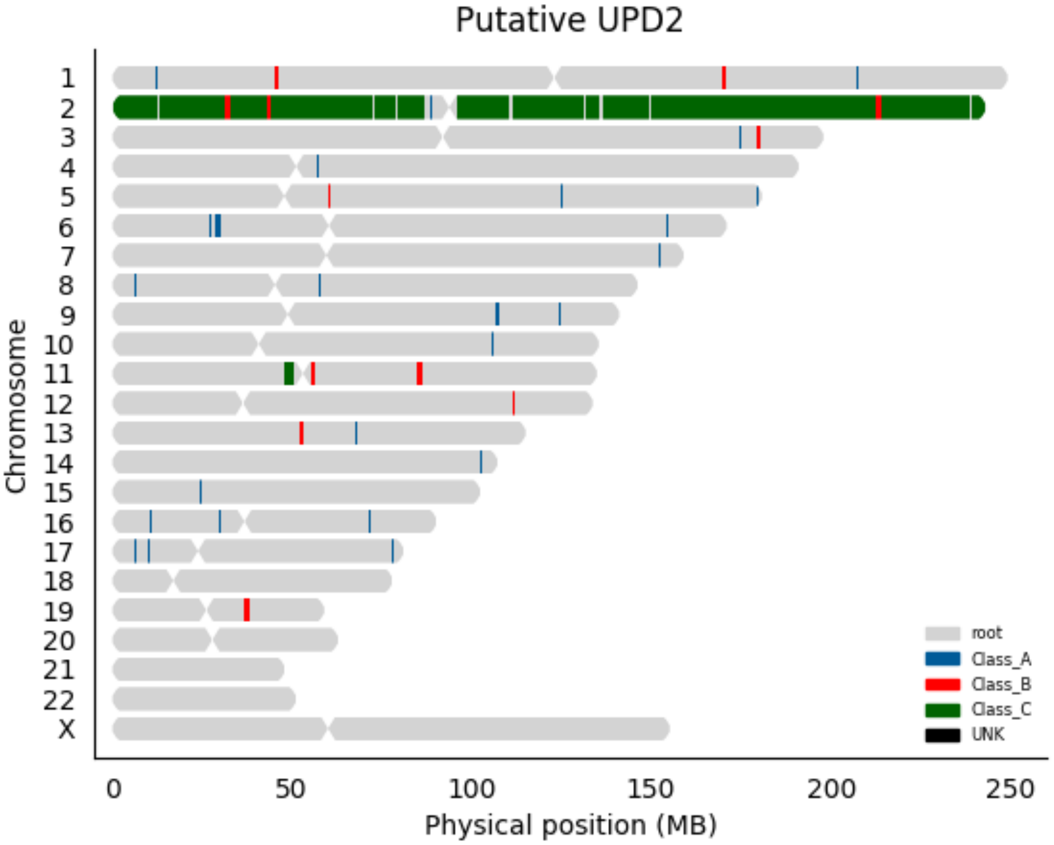

Putative UPD2
Chromosome
X
0
50
100
150
200
250
Physical position (MB)
root
Class_A
Class_B
Class_C
UNK

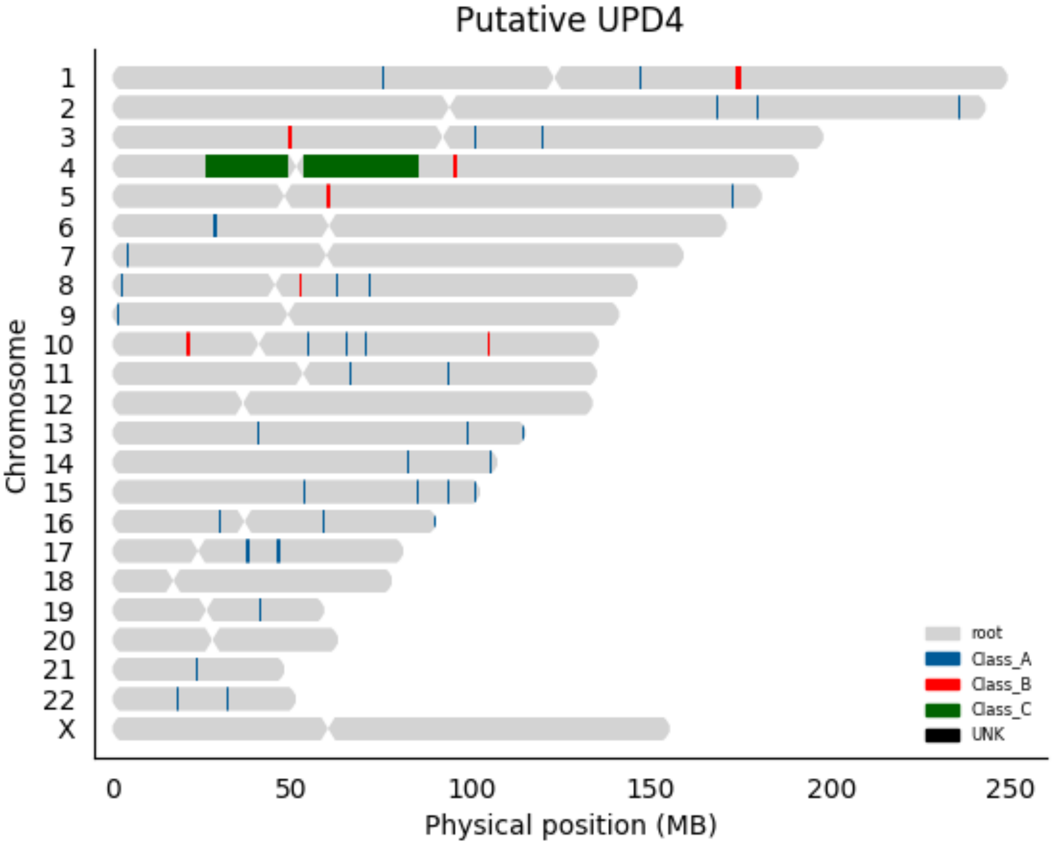

Putative UPD4
Chromosome
X
0
50
100
150
200
250
Physical position (MB)
root
Class_A
Class_B
Class_C
UNK

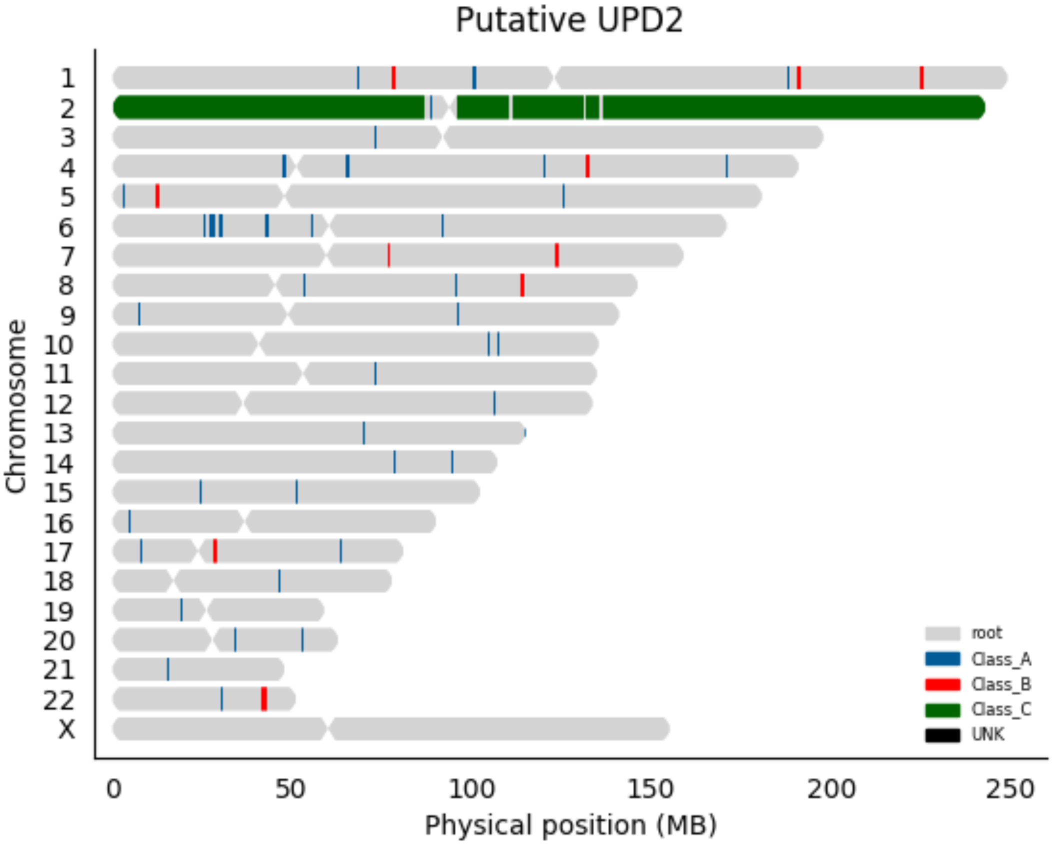

Putative UPD2
Chromosome
X
0
50
100
150
200
250
Physical position (MB)
root
Class_A
Class_B
Class_C
UNK

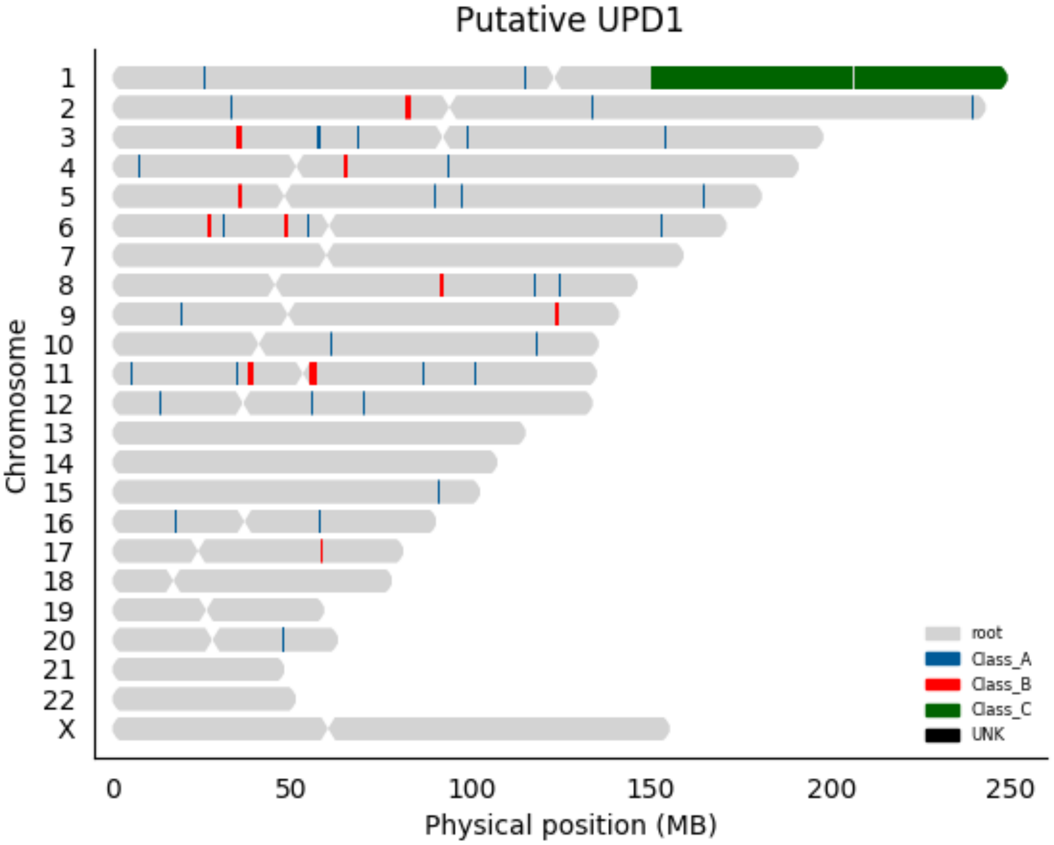

Putative UPD1
Chromosome
X
0
50
100
150
200
250
Physical position (MB)
root
Class_A
Class_B
Class_C
UNK

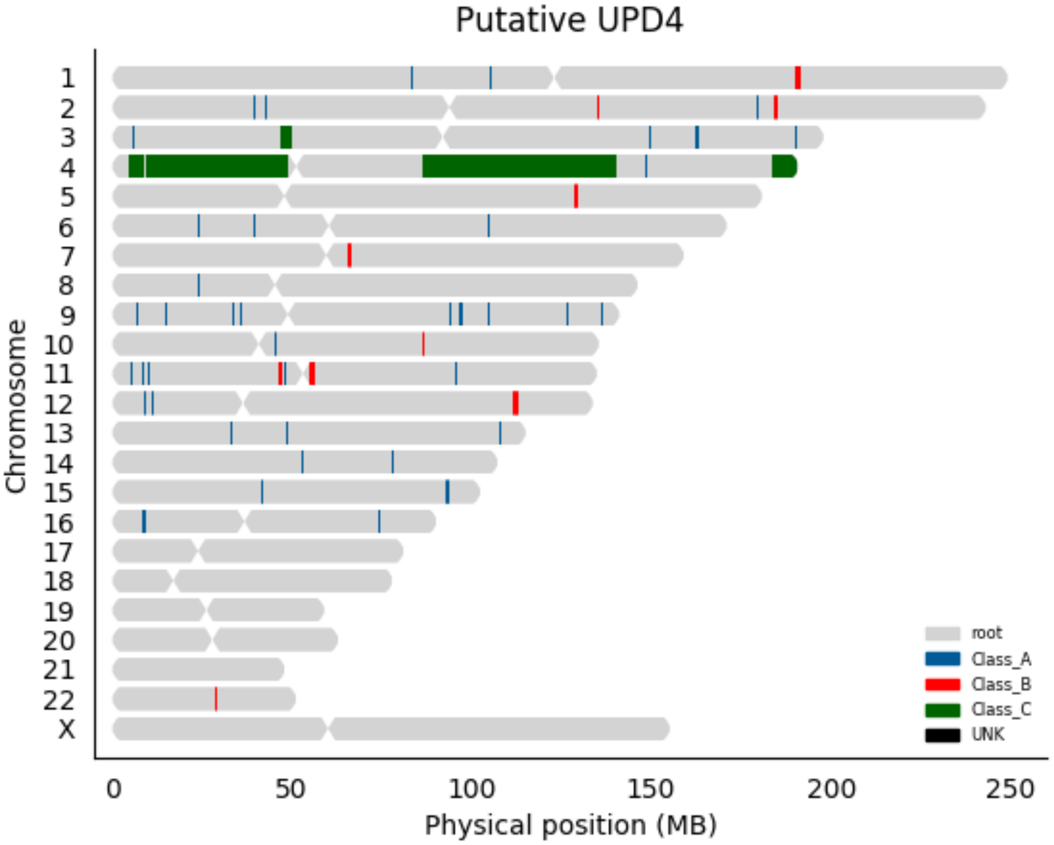

Putative UPD4
Chromosome
X
0
50
100
150
200
250
Physical position (MB)
root
Class_A
Class_B
Class_C
UNK

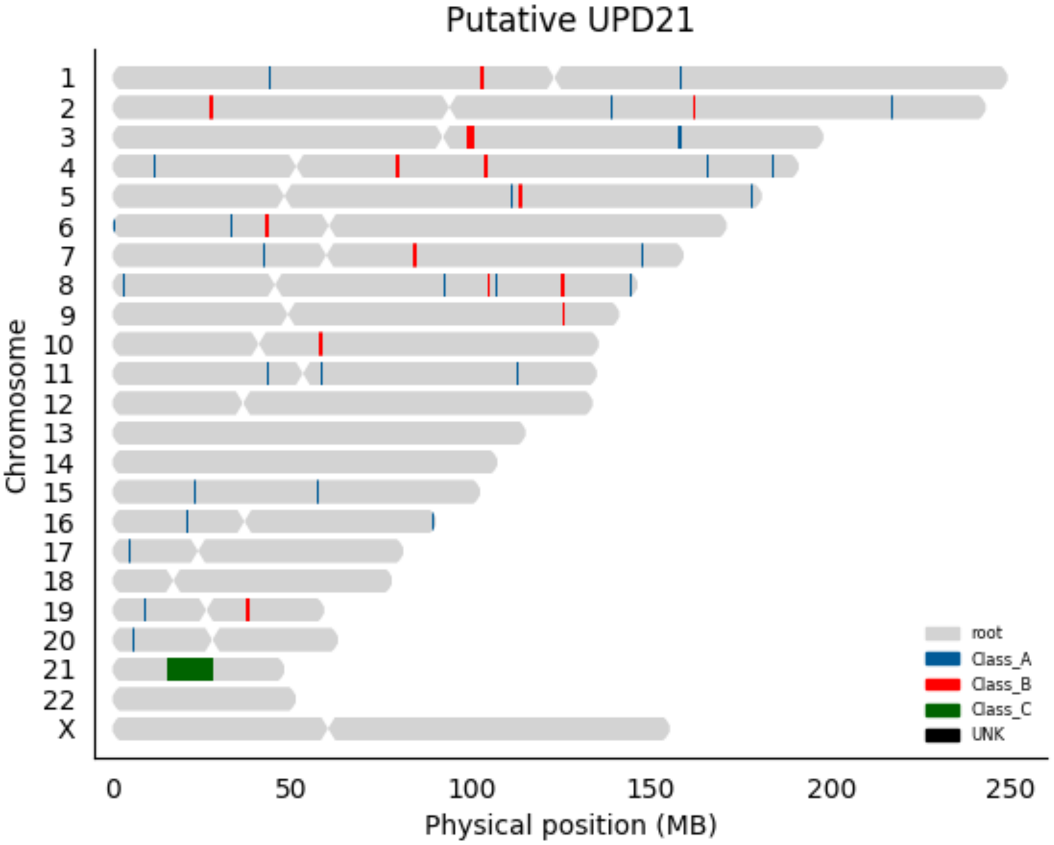

Putative UPD21
Chromosome
X
0
50
100
150
200
250
Physical position (MB)
root
Class_A
Class_B
Class_C
UNK

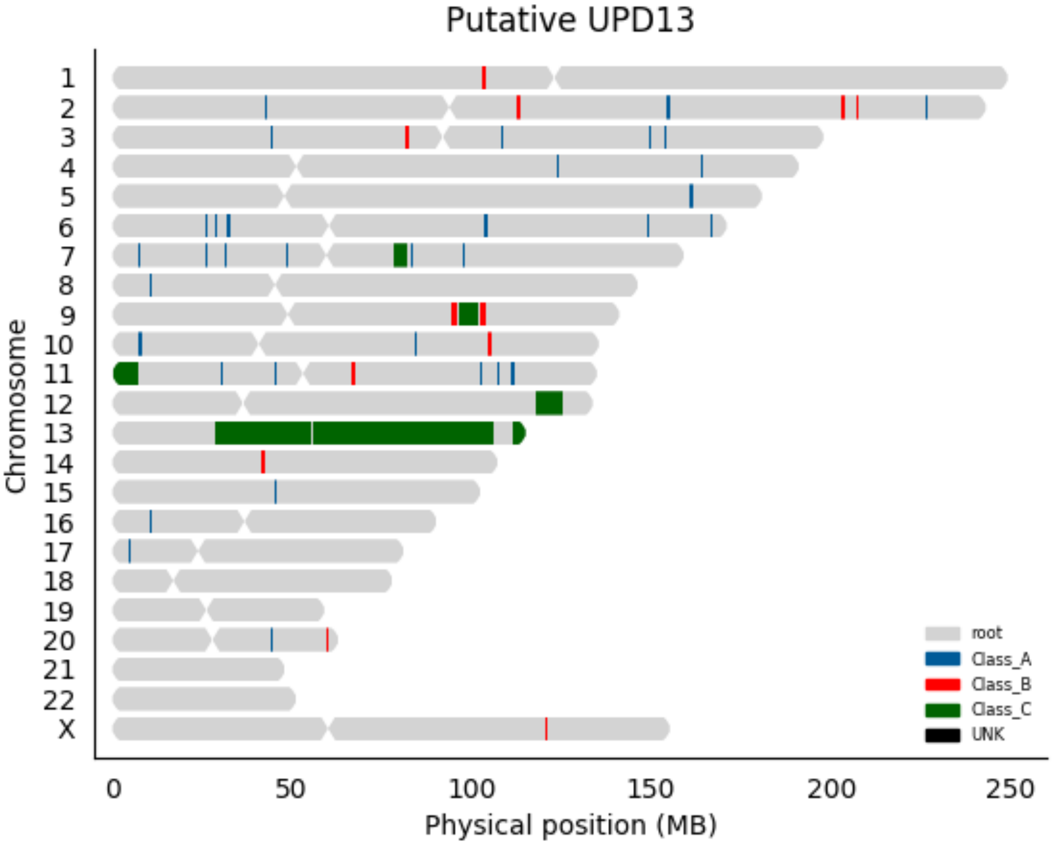
Putative UPD13
Chromosome
X
0
50
100
150
200
250
Physical position (MB)
root
Class_A
Class_B
Class_C
UNK

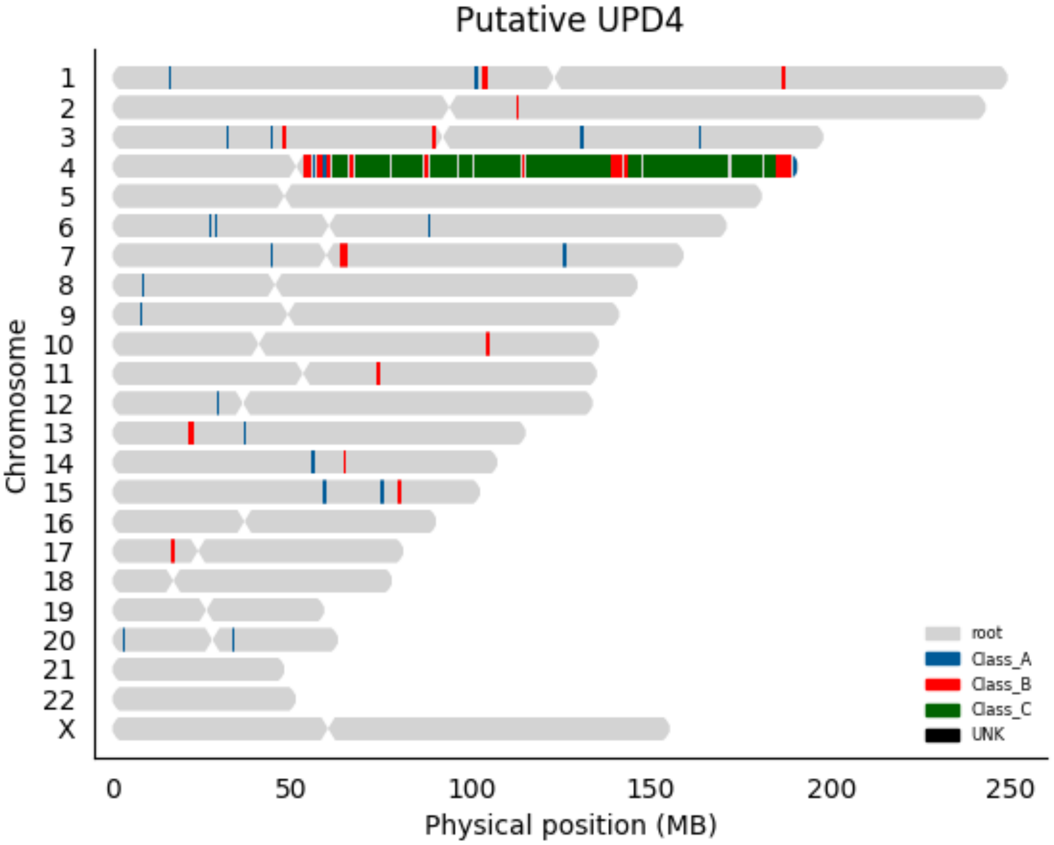

Putative UPD4
Chromosome
X
0
50
100
150
200
250
Physical position (MB)
root
Class_A
Class_B
Class_C
UNK

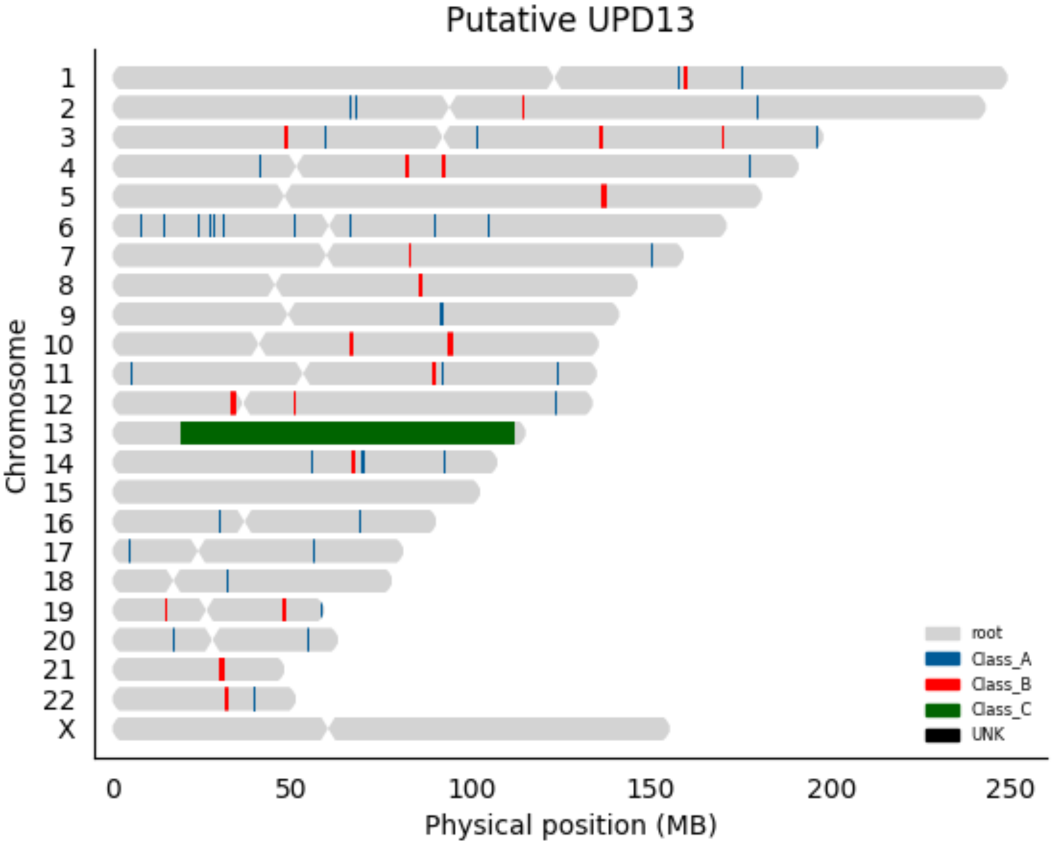

Putative UPD13
Chromosome
X
0
50
100
150
200
250
Physical position (MB)
root
Class_A
Class_B
Class_C
UNK

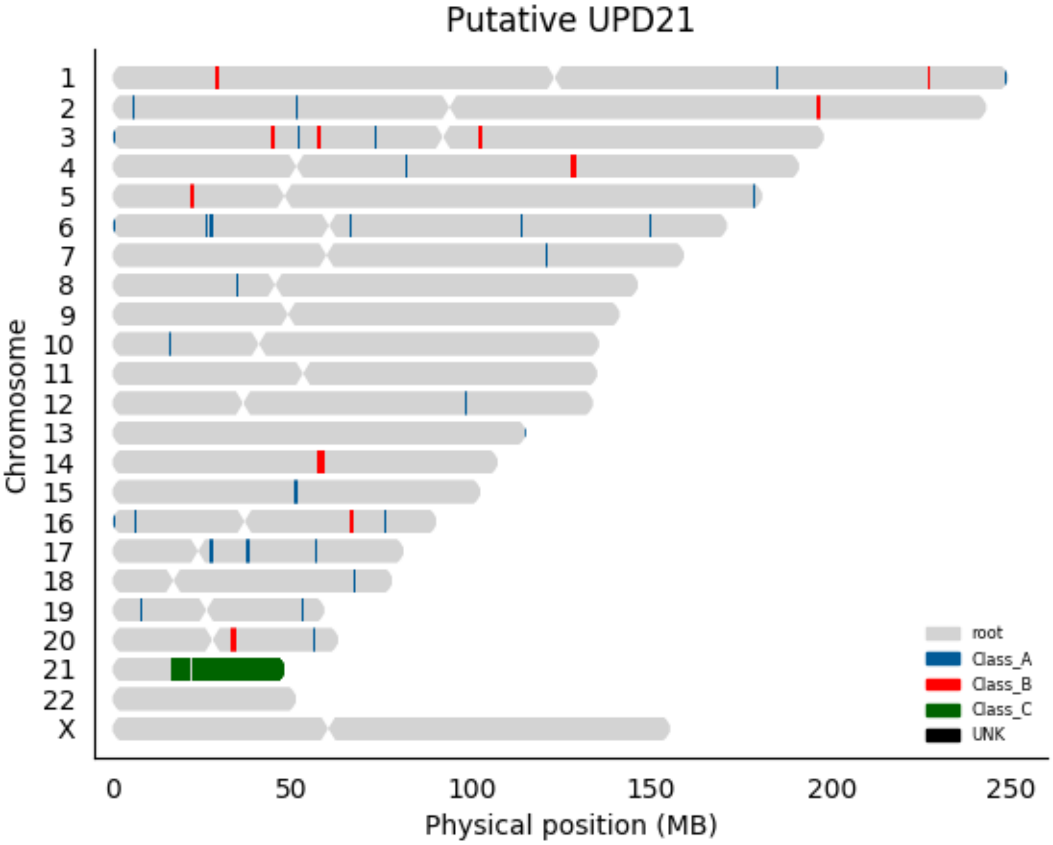

Putative UPD21
Chromosome
X
0
50
100
150
200
250
Physical position (MB)
root
Class_A
Class_B
Class_C
UNK

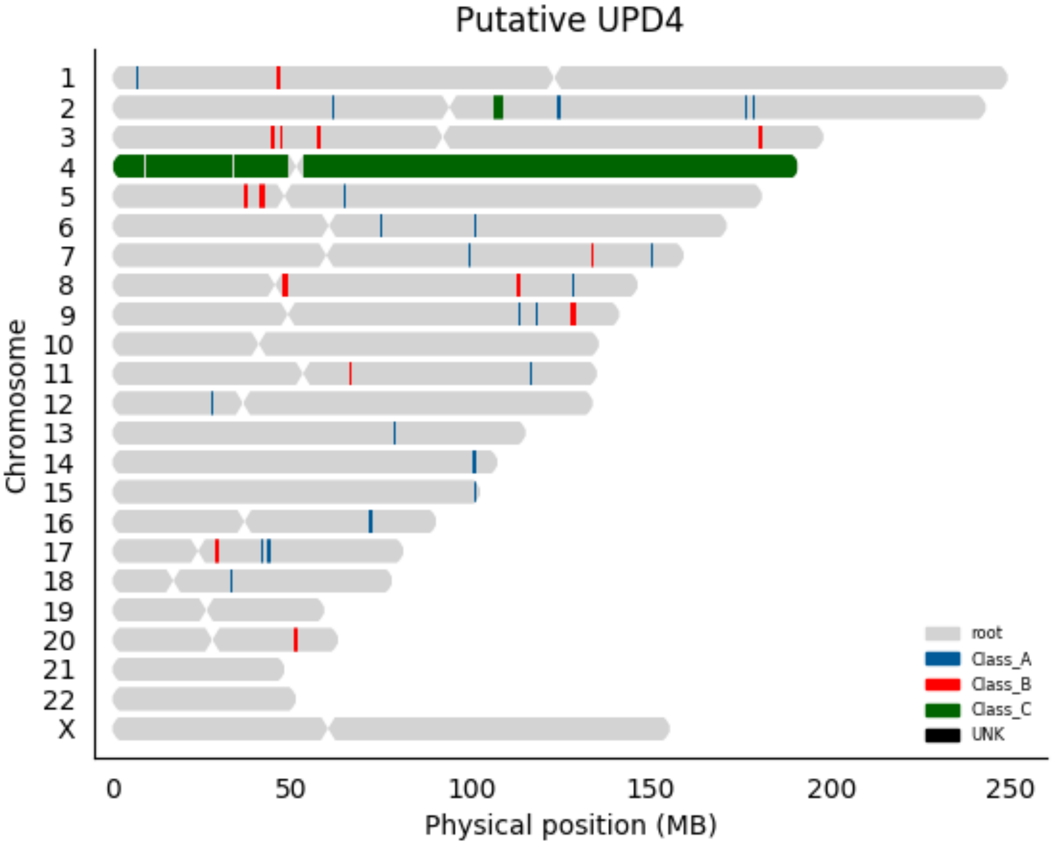

Putative UPD4
Chromosome
X
0
50
100
150
200
250
Physical position (MB)
root
Class_A
Class_B
Class_C
UNK

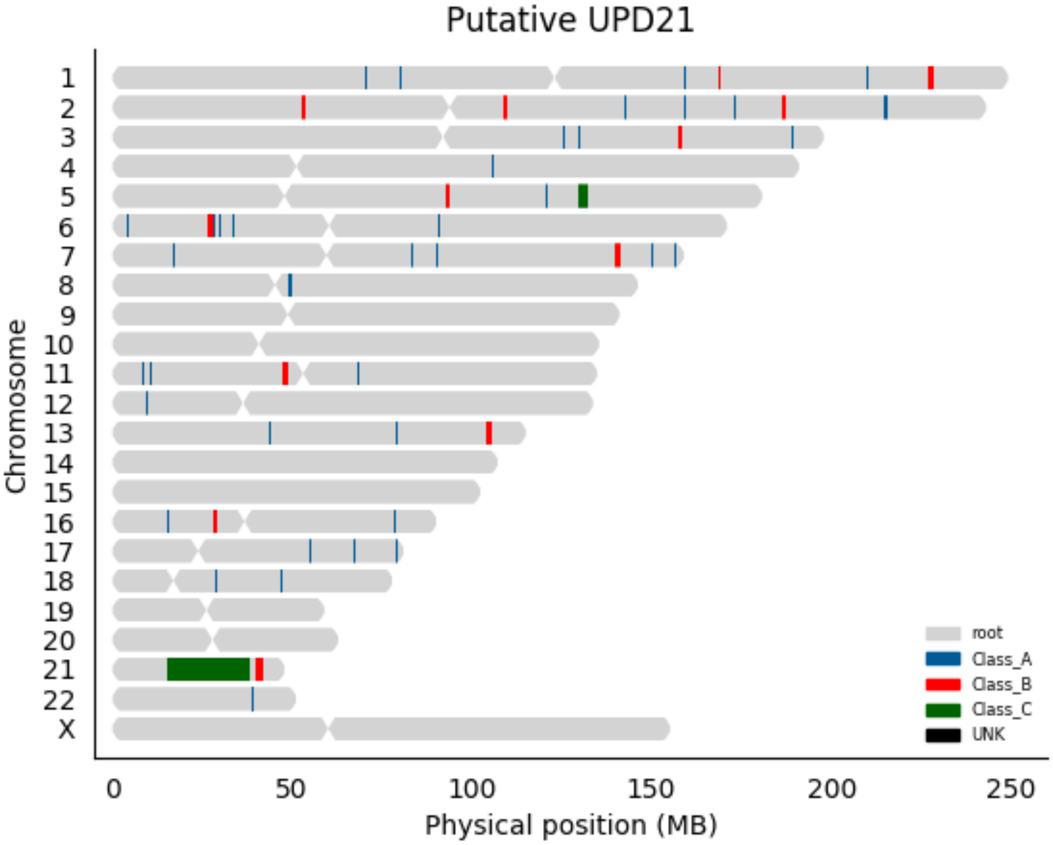
Putative UPD21
Chromosome
X
root
Class_A
Class_B
Class_C
UNK
0
50
100
150
200
250
Physical position (MB)

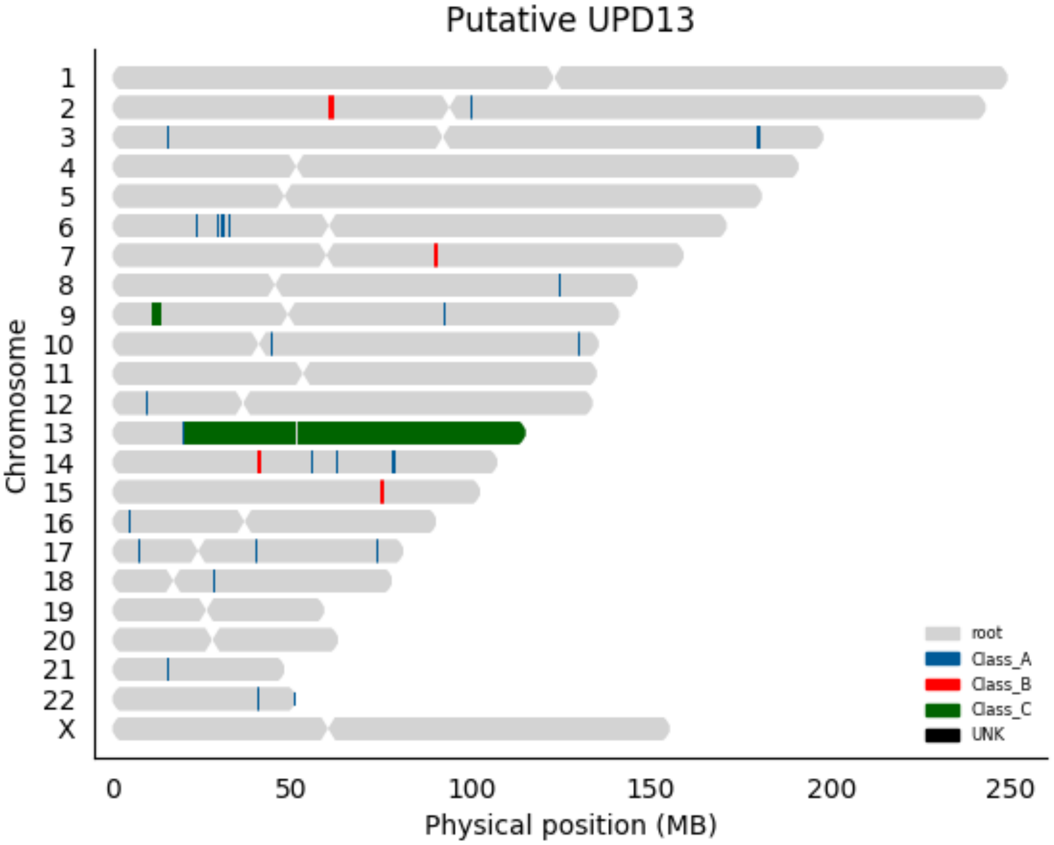
Putative UPD13
Chromosome
X
0
50
100
150
200
250
Physical position (MB)
root
Class_A
Class_B
Class_C
UNK

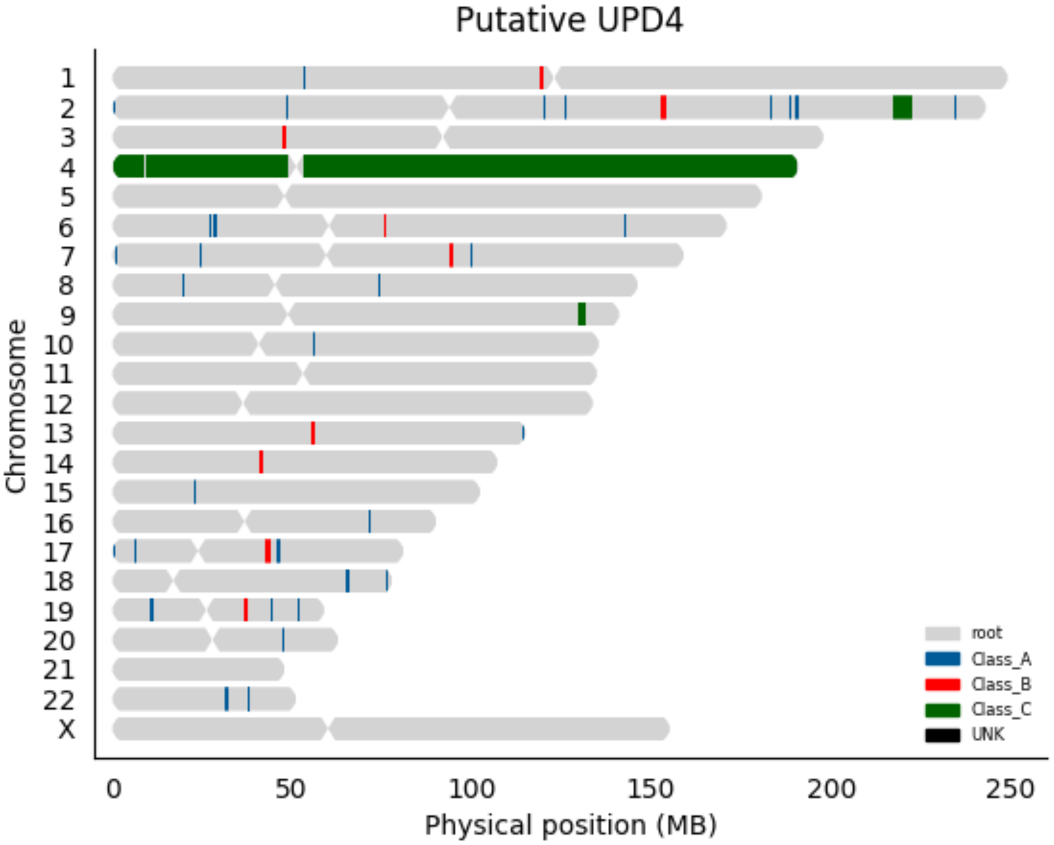
Putative UPD4
Chromosome
X
0
50
100
150
200
250
Physical position (MB)
root
Class_A
Class_B
Class_C
UNK

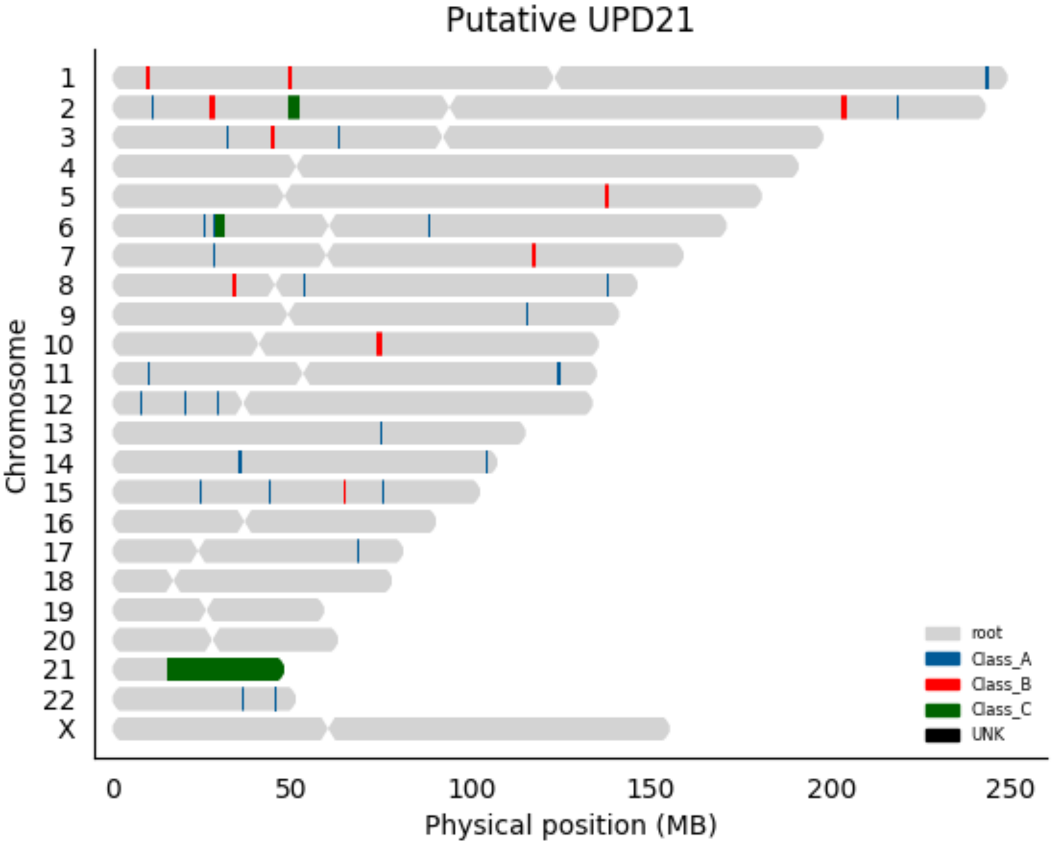
Putative UPD21
Chromosome
X
0
50
100
150
200
250
Physical position (MB)
root
Class_A
Class_B
Class_C
UNK

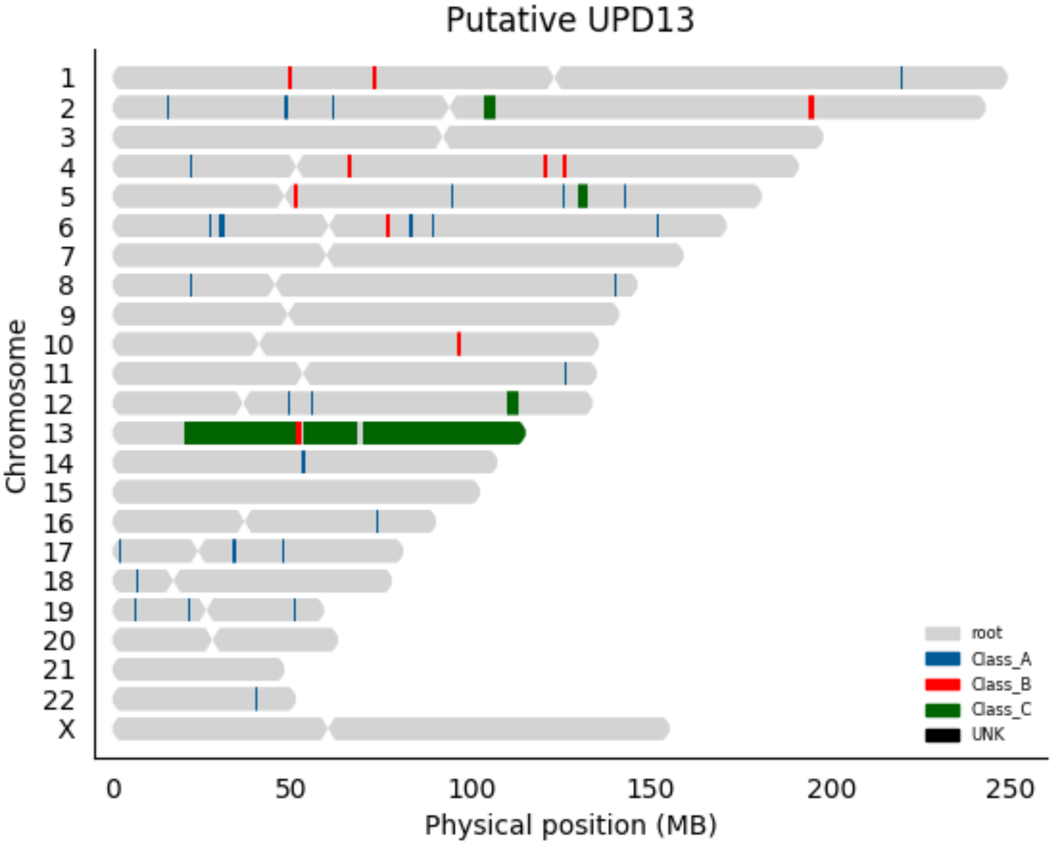

Putative UPD13
Chromosome
X
0
50
100
150
200
250
Physical position (MB)
root
Class_A
Class_B
Class_C
UNK

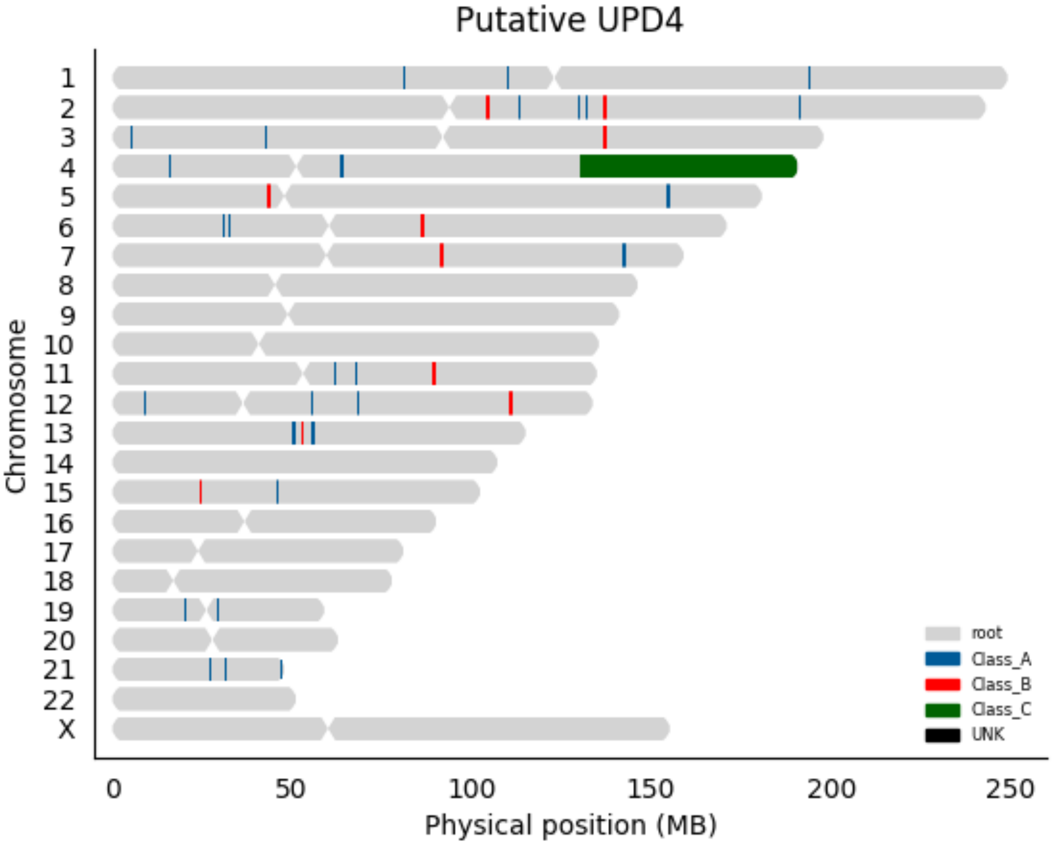

Putative UPD4
Chromosome
X
root
Class_A
Class_B
Class_C
UNK
0
50
100
150
200
250
Physical position (MB)

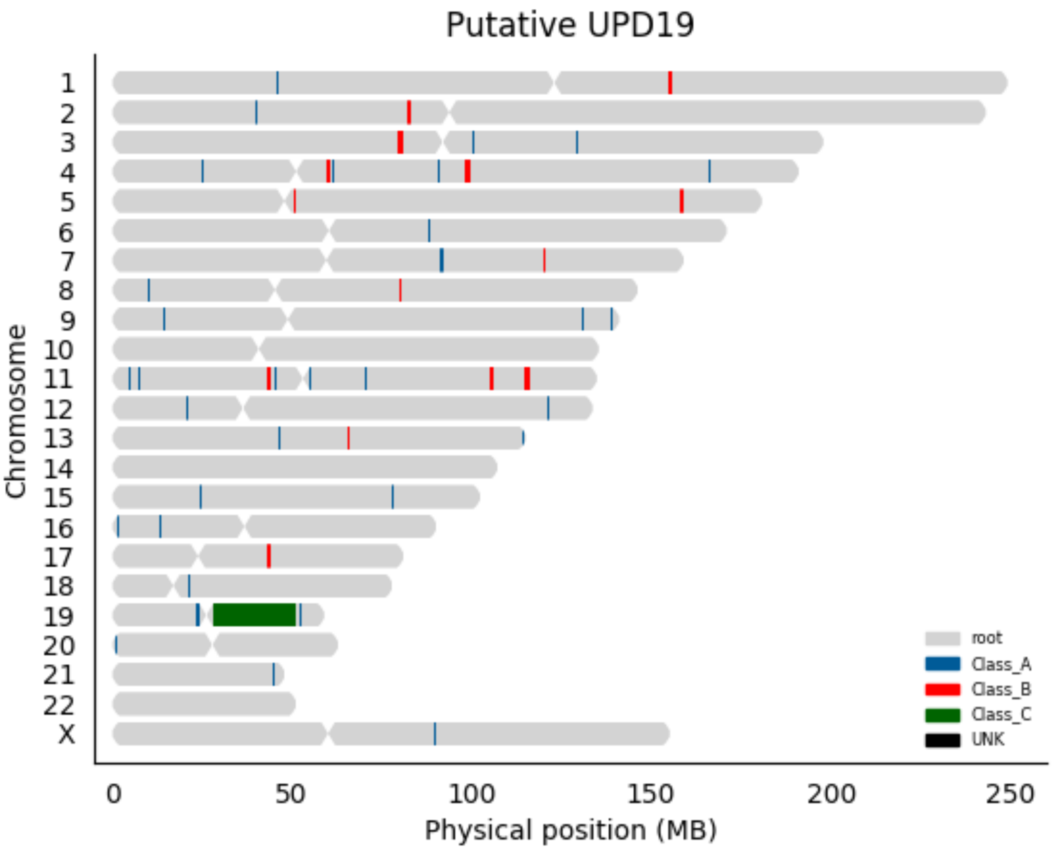
Putative UPD19
Chromosome
X
0
50
100
150
200
250
Physical position (MB)
root
Class_A
Class_B
Class_C
UNK

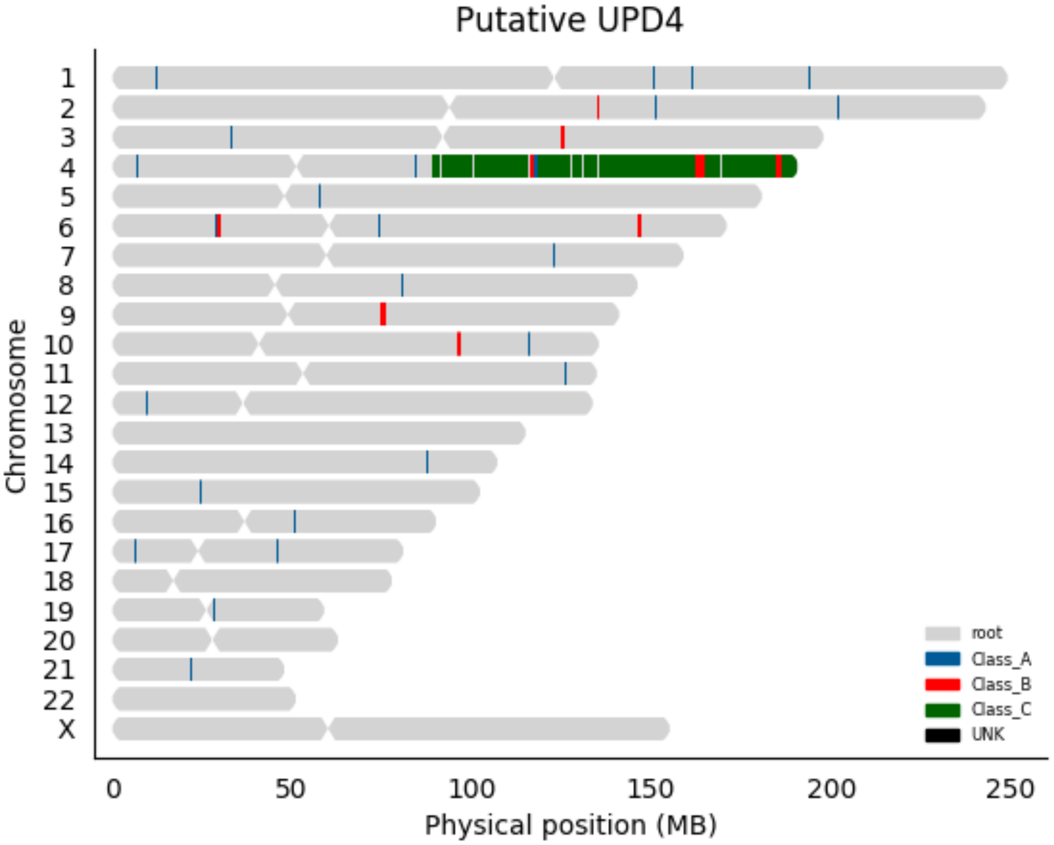

Putative UPD4
X
Chromosome
0
50
100
150
200
250
Physical position (MB)
root
Class_A
Class_B
Class_C
UNK

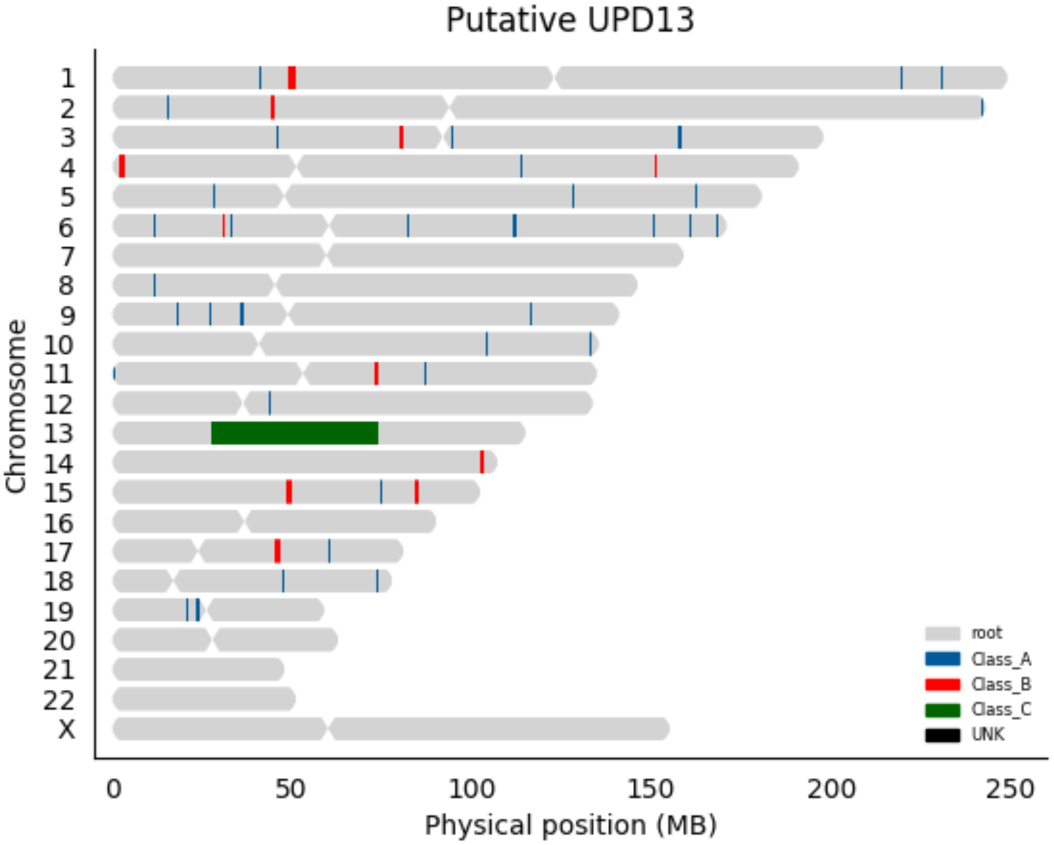
Putative UPD13
Chromosome
X
0
50
100
150
200
250
Physical position (MB)
root
Class_A
Class_B
Class_C
UNK

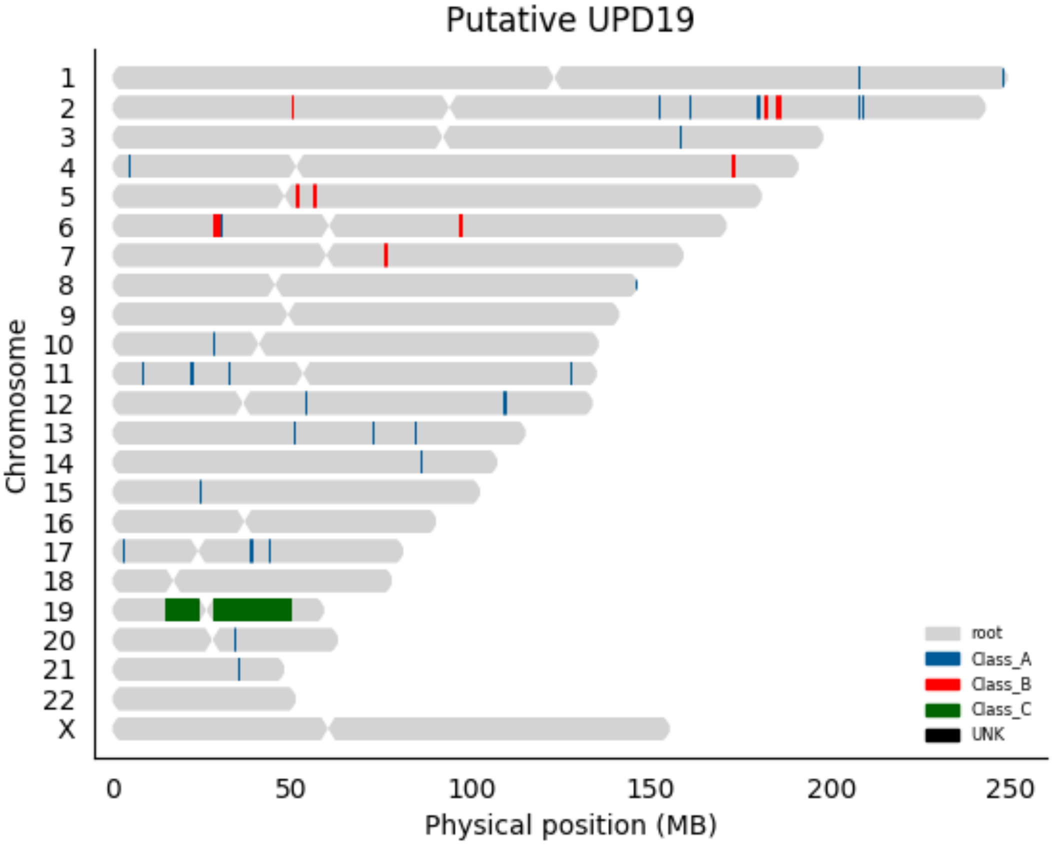

Putative UPD19
Chromosome
X
0
50
100
150
200
250
Physical position (MB)
root
Class_A
Class_B
Class_C
UNK

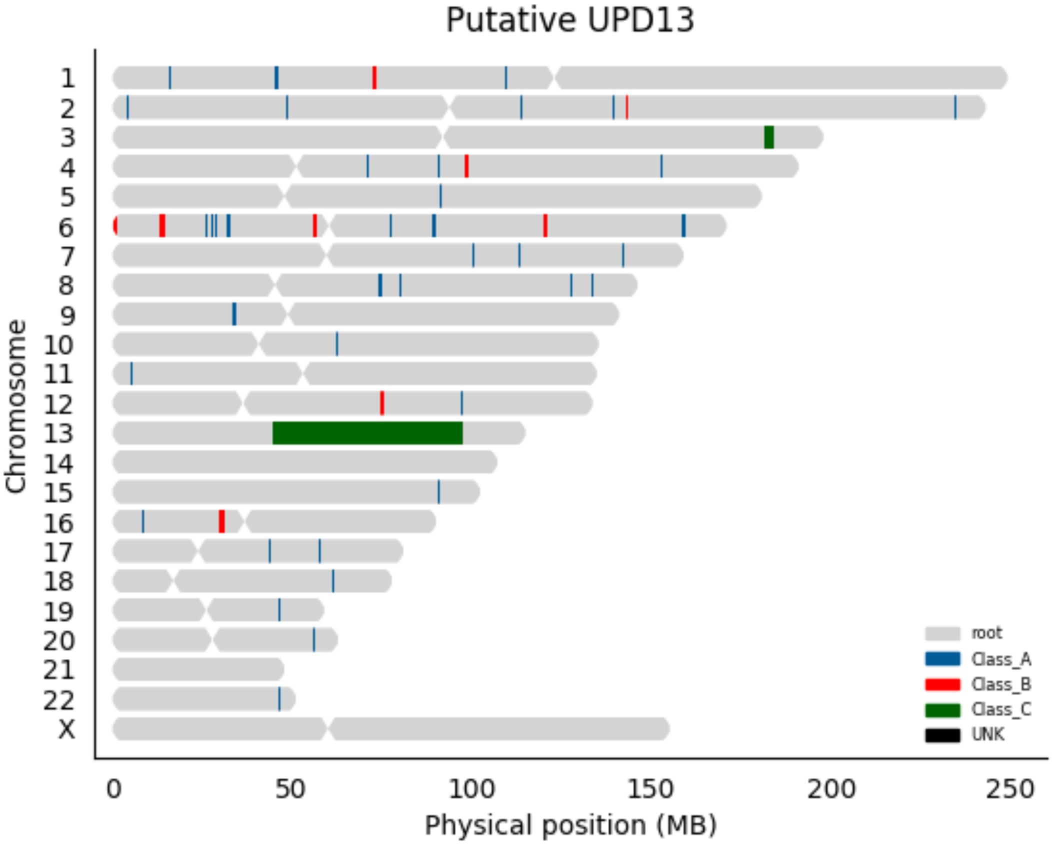

Putative UPD13
Chromosome
X
0
50
100
150
200
250
Physical position (MB)
root
Class_A
Class_B
Class_C
UNK

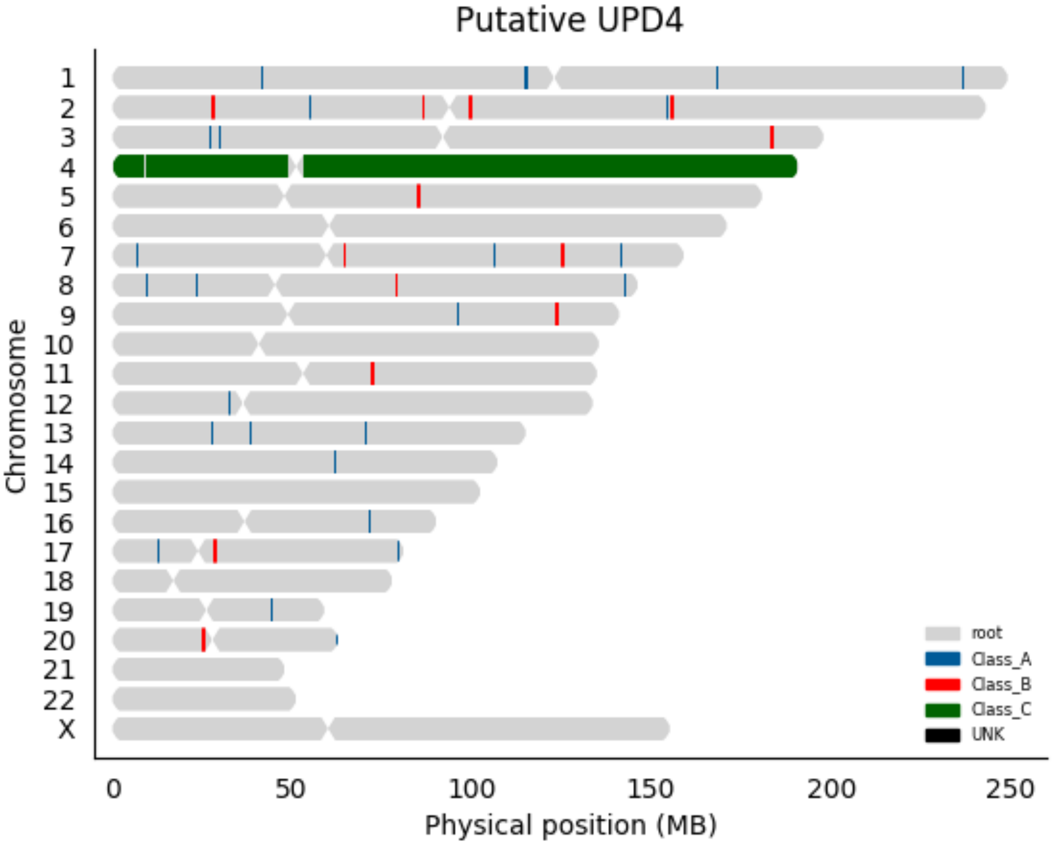

Putative UPD4
Chromosome
X
0
50
100
150
200
250
Physical position (MB)
root
Class_A
Class_B
Class_C
UNK

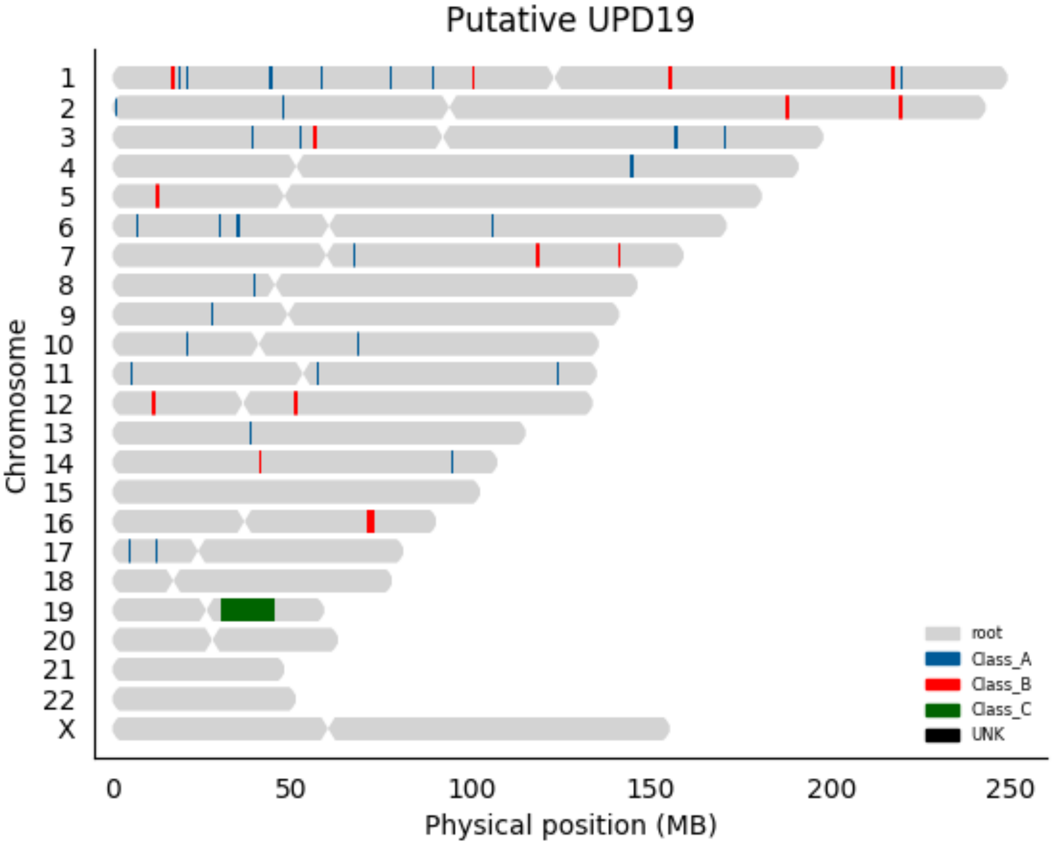

Putative UPD19
Chromosome
X
0
50
100
150
200
250
Physical position (MB)
root
Class_A
Class_B
Class_C
UNK

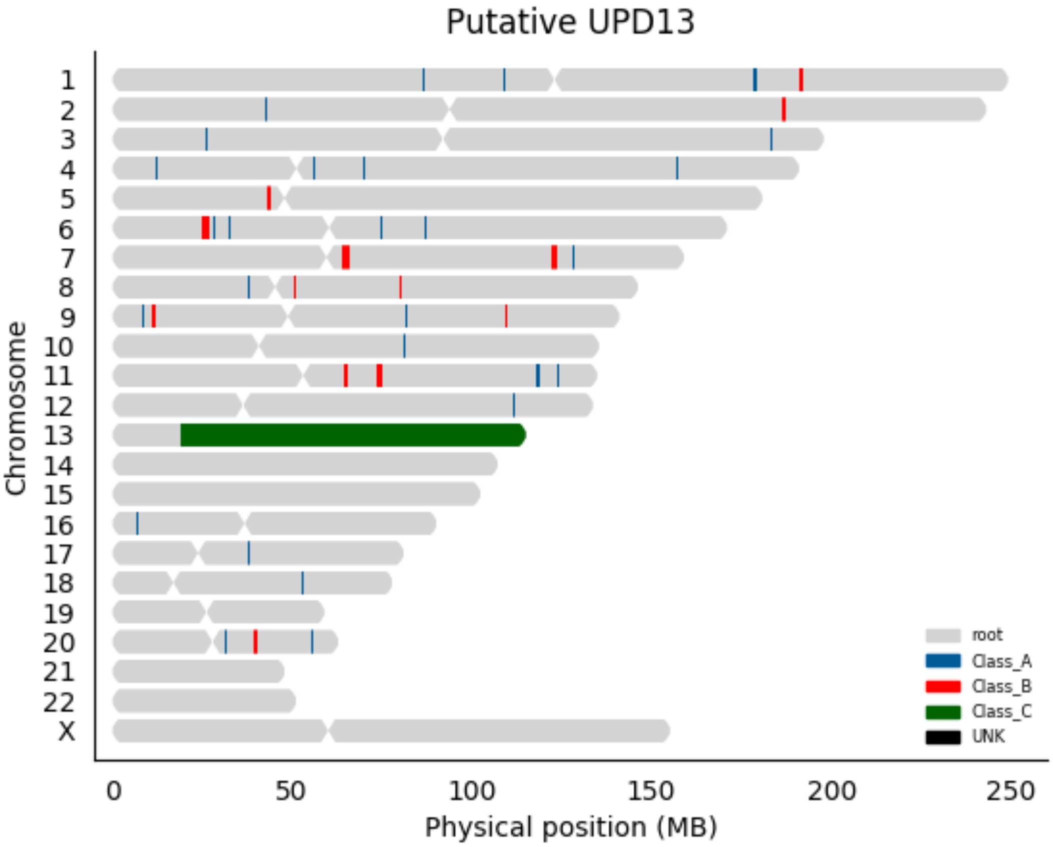

Putative UPD13
X
Chromosome
0
50
100
150
200
250
Physical position (MB)
root
Class_A
Class_B
Class_C
UNK

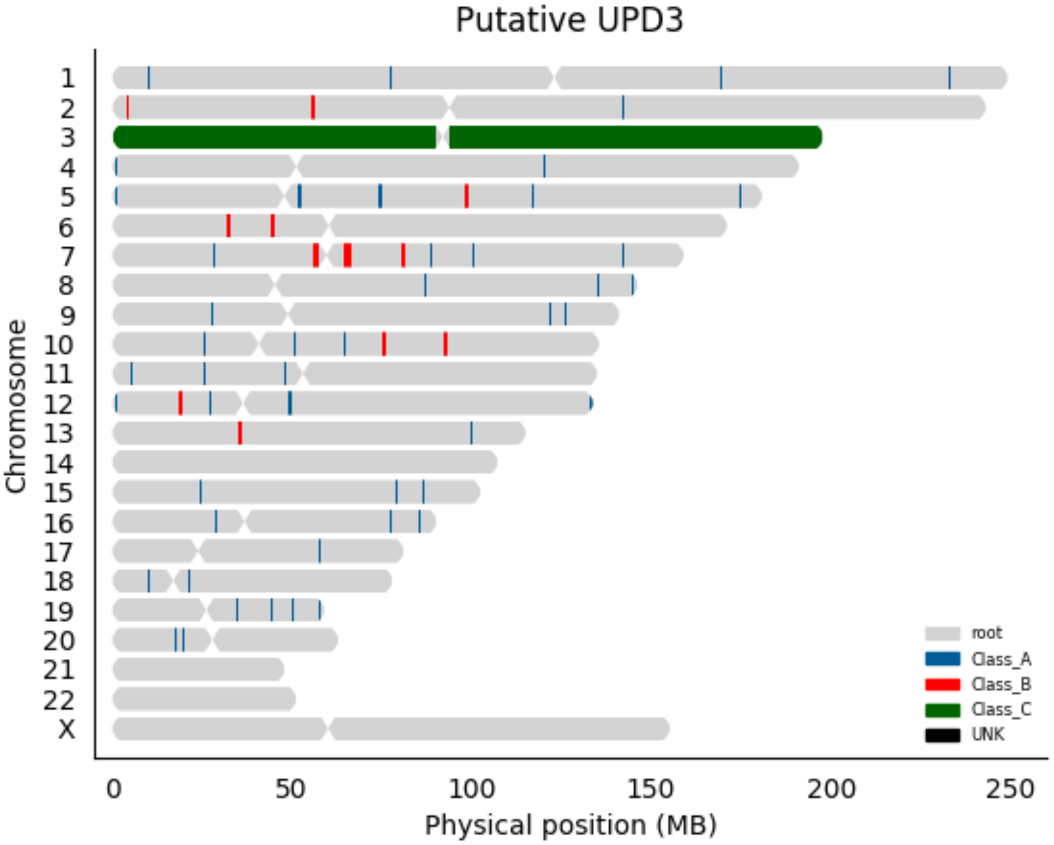
Putative UPD3
Chromosome
X
0
50
100
150
200
250
Physical position (MB)
root
Class_A
Class_B
Class_C
UNK

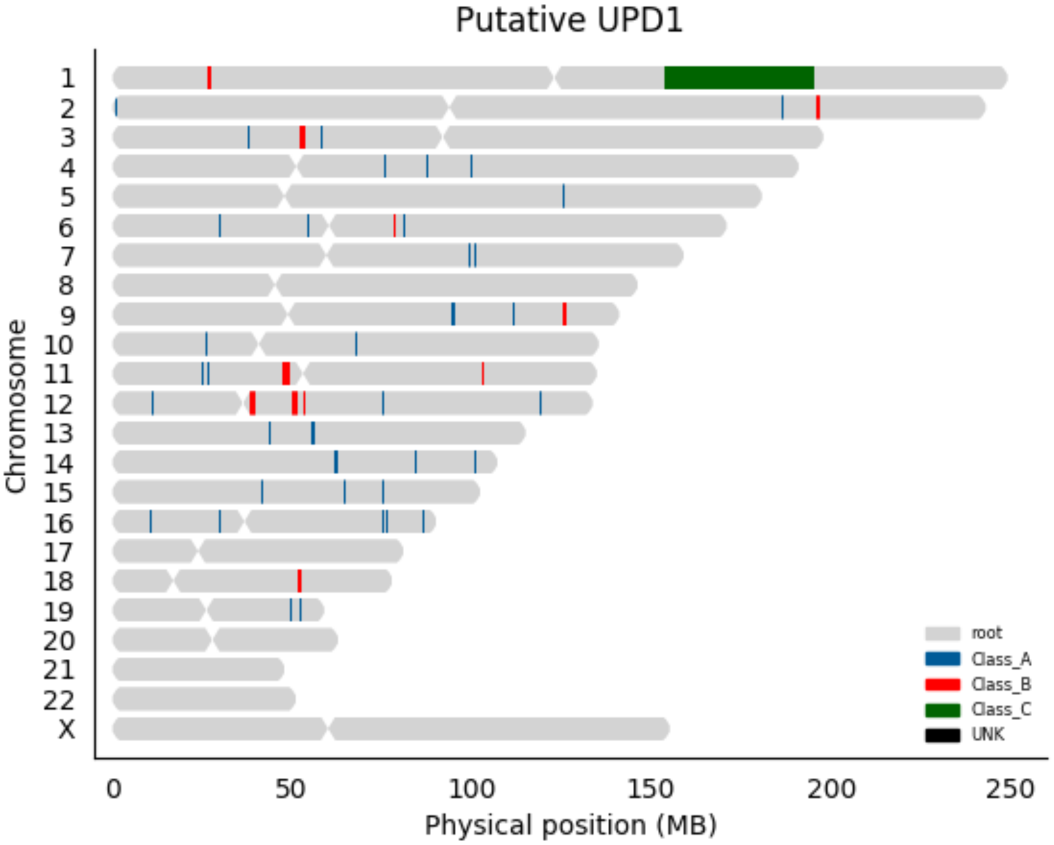

Putative UPD1
Chromosome
X
0
50
100
150
200
250
Physical position (MB)
root
Class_A
Class_B
Class_C
UNK

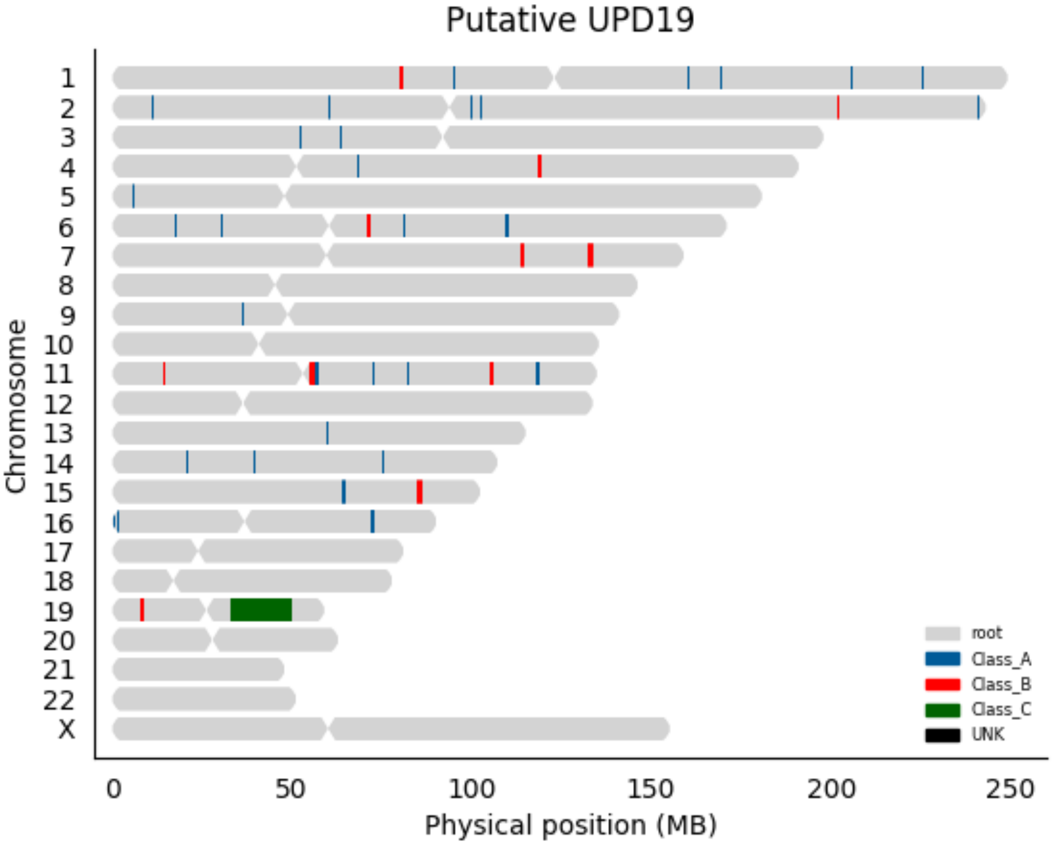

Putative UPD19
Chromosome
X
0
50
100
150
200
250
Physical position (MB)
root
Class_A
Class_B
Class_C
UNK

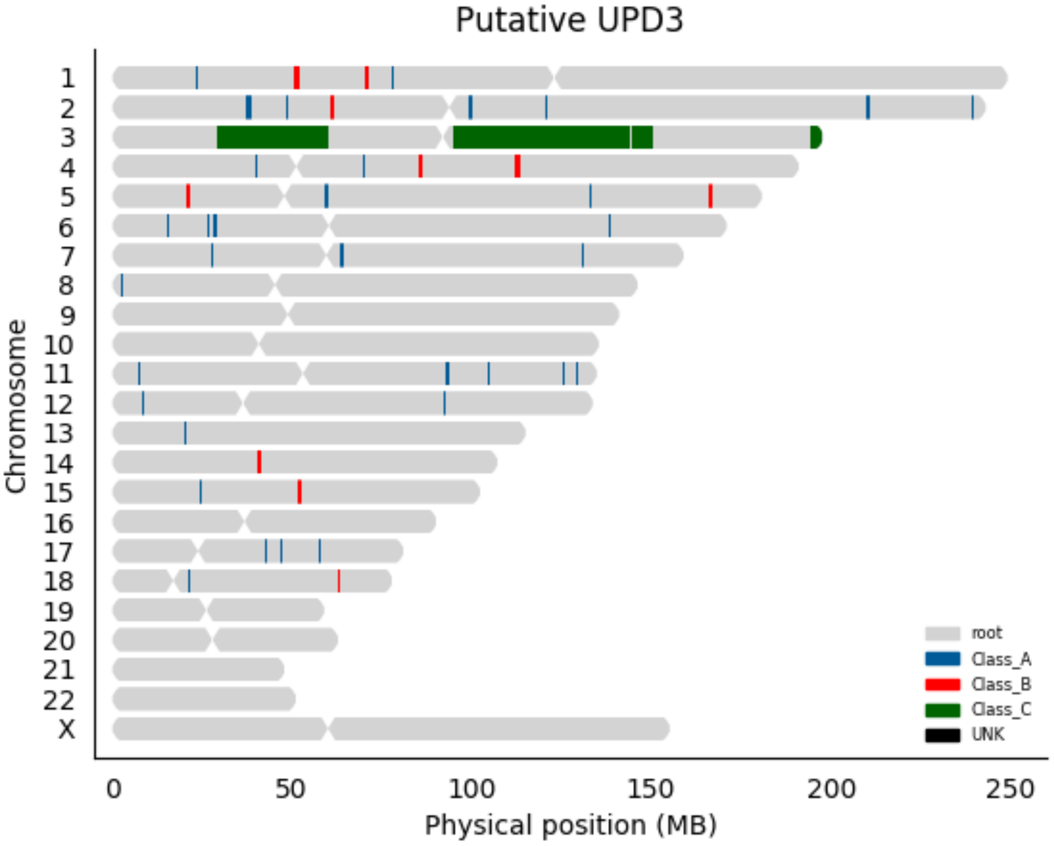

Putative UPD3
Chromosome
X
0
50
100
150
200
250
Physical position (MB)
root
Class_A
Class_B
Class_C
UNK

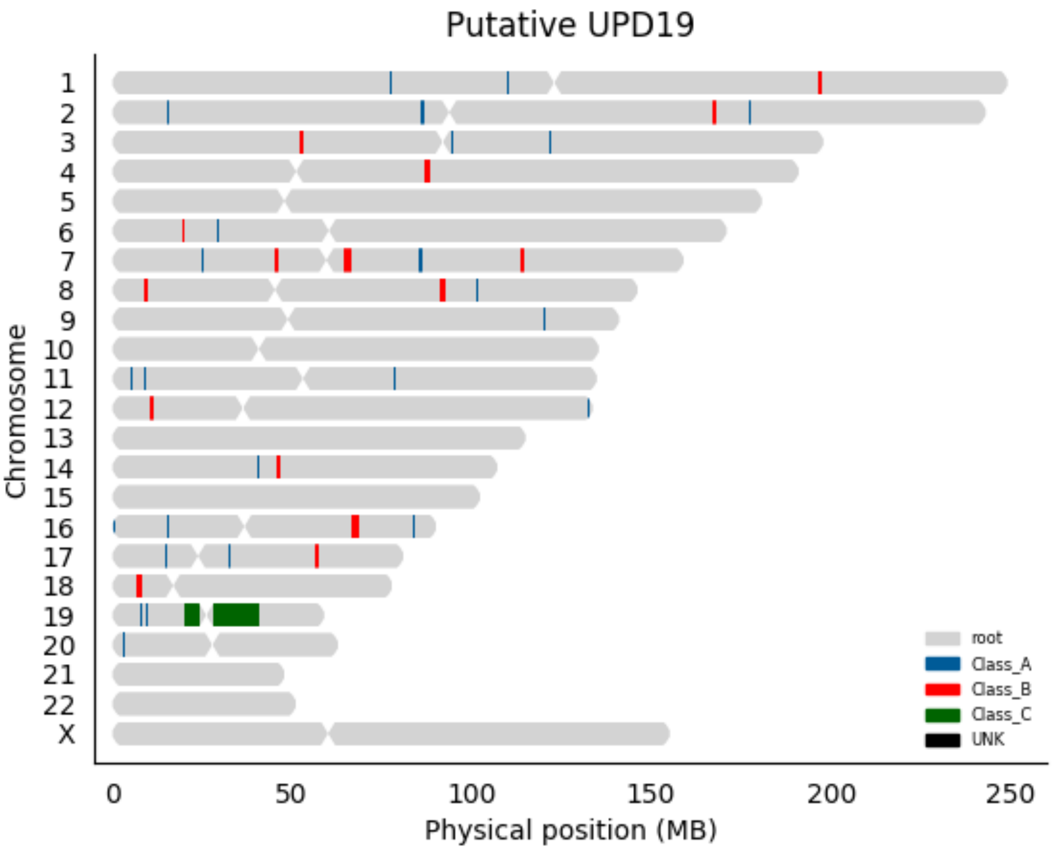
Putative UPD19
Chromosome
X
0
50
100
150
200
250
Physical position (MB)
root
Class_A
Class_B
Class_C
UNK

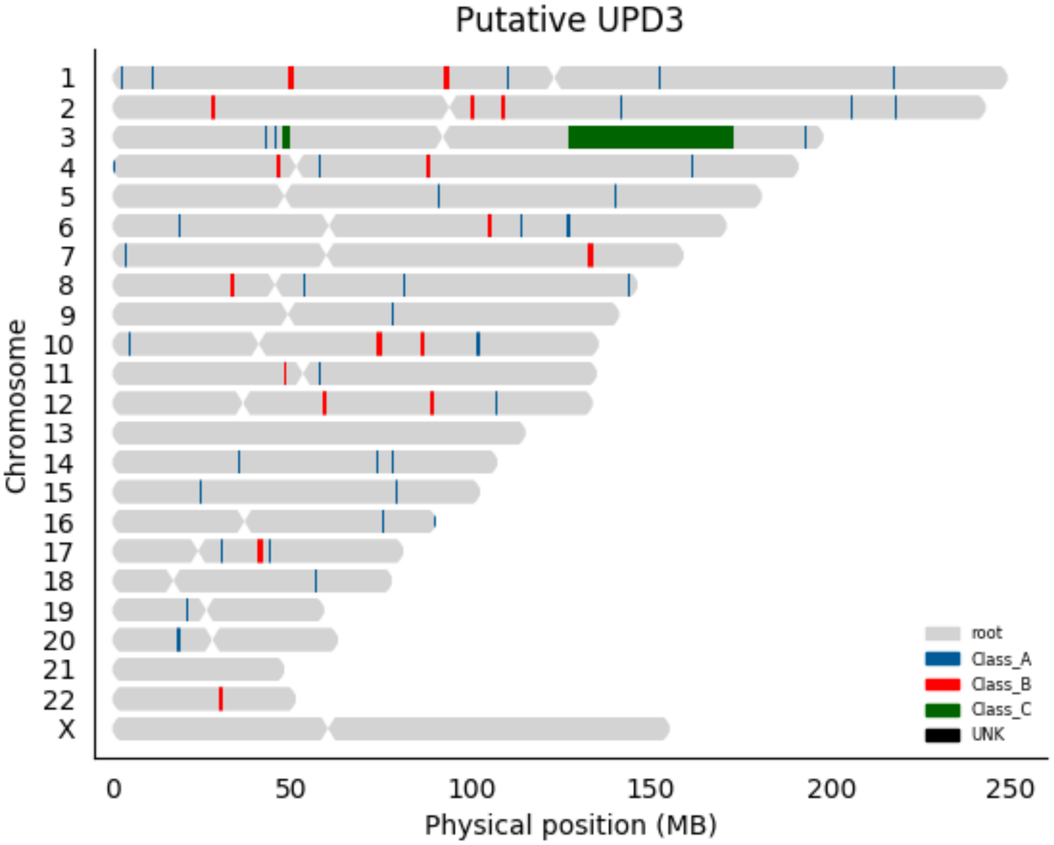
Putative UPD3
Chromosome
X
0
50
100
150
200
250
Physical position (MB)
root
Class_A
Class_B
Class_C
UNK

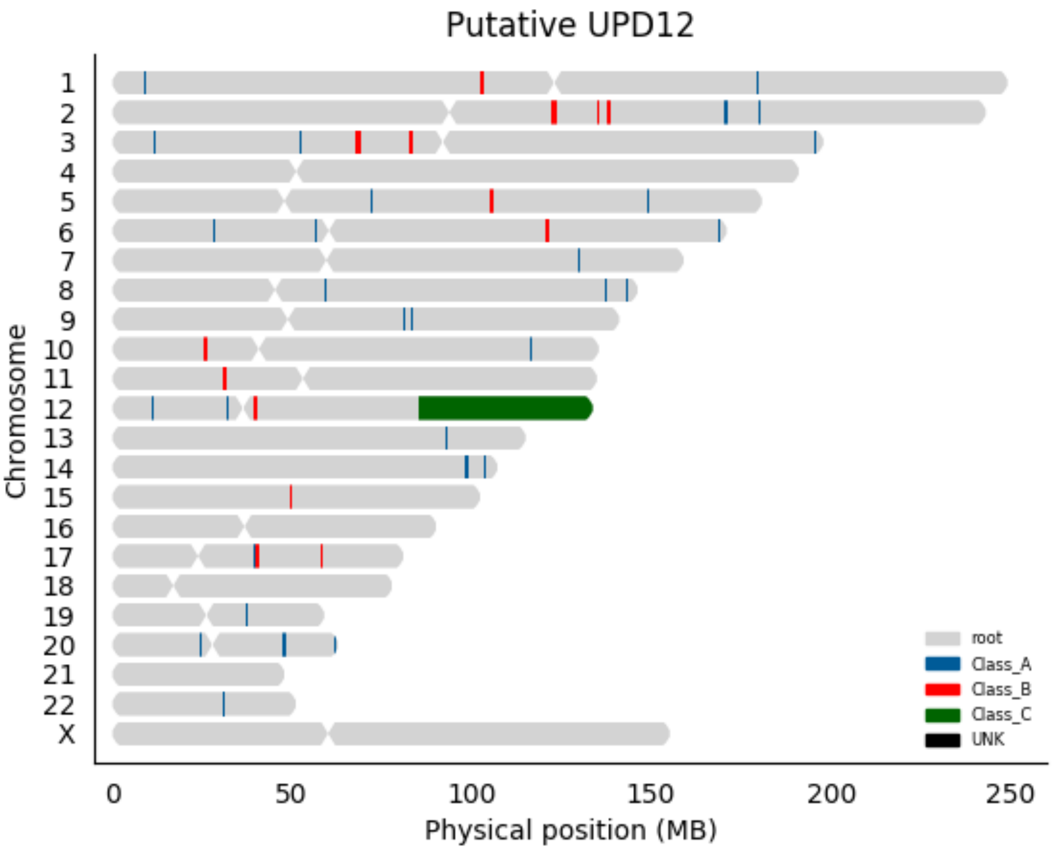

Putative UPD12
Chromosome
X
0
50
100
150
200
250
Physical position (MB)
root
Class_A
Class_B
Class_C
UNK

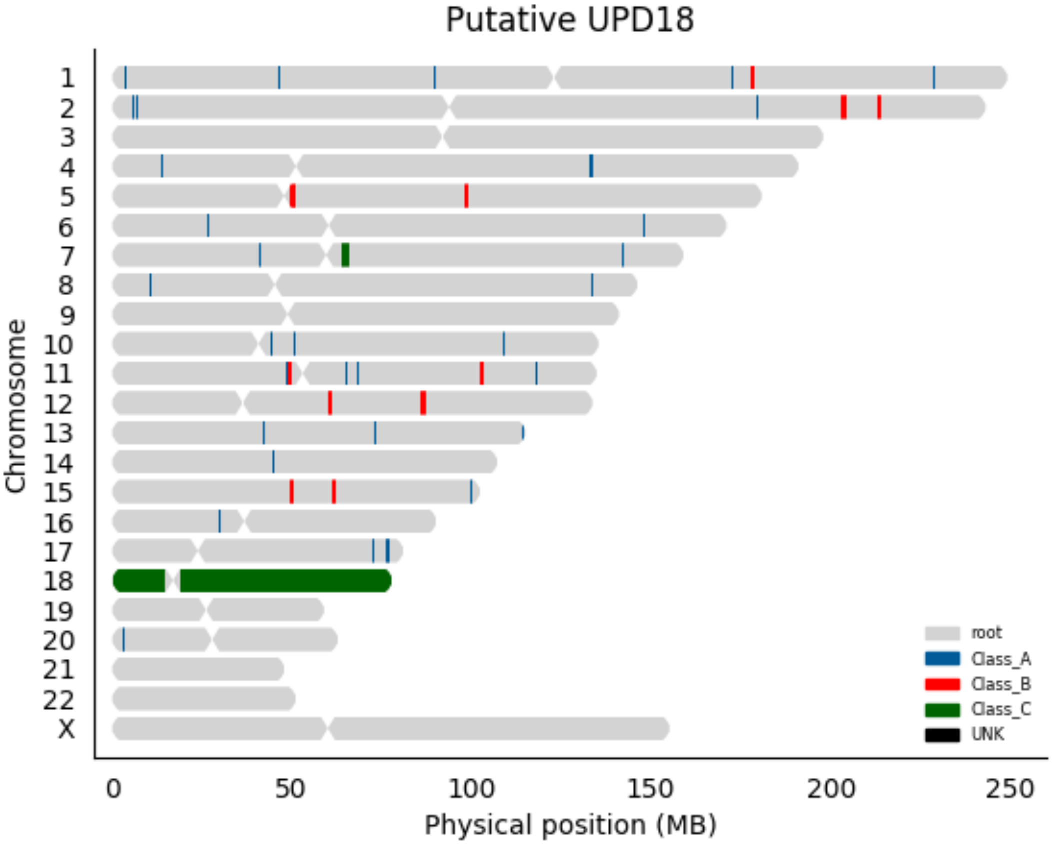
Putative UPD18
Chromosome
X
0
50
100
150
200
250
Physical position (MB)
root
Class_A
Class_B
Class_C
UNK

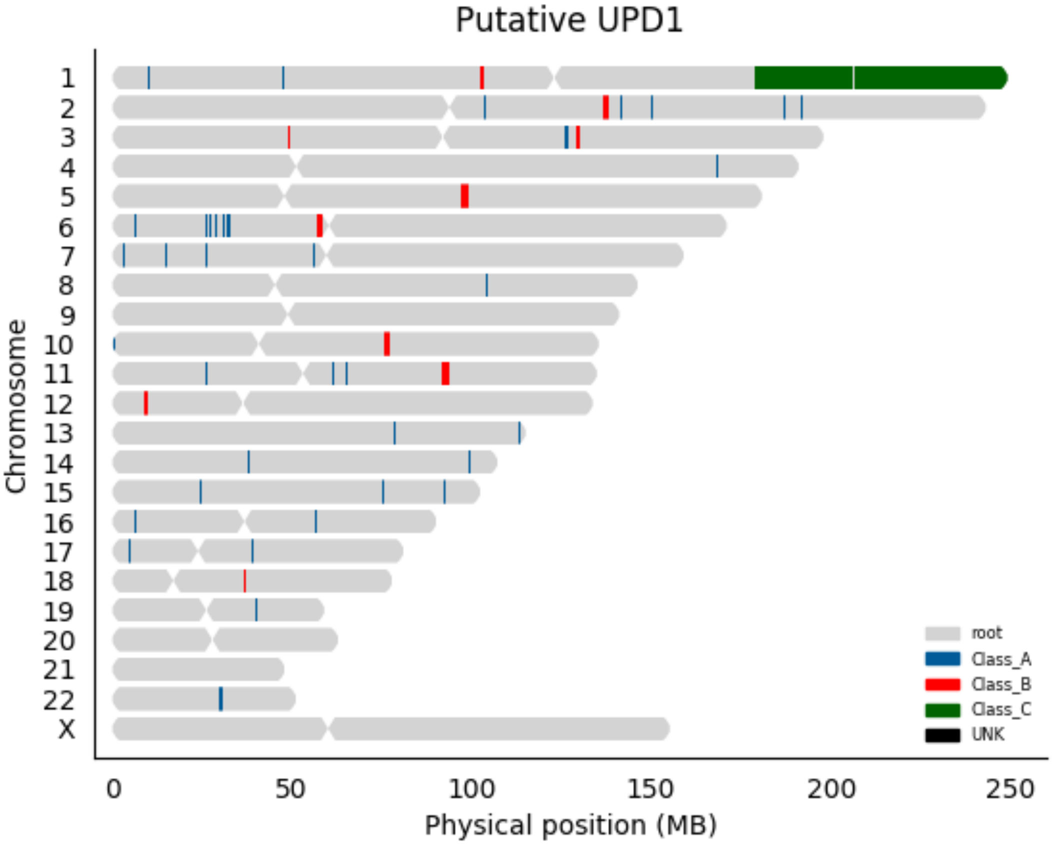

Putative UPD1
X
Chromosome
0
50
100
150
200
250
Physical position (MB)
root
Class_A
Class_B
Class_C
UNK

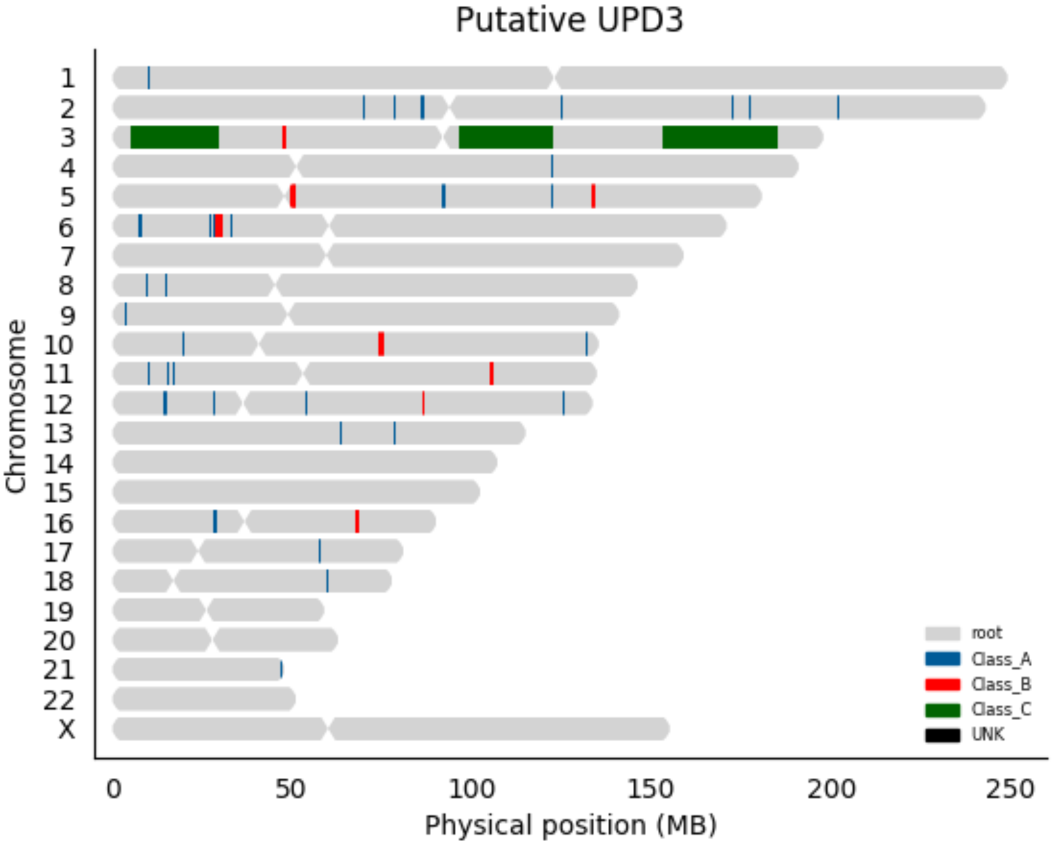

Putative UPD3
X
Chromosome
0
50
100
150
200
250
Physical position (MB)
root
Class_A
Class_B
Class_C
UNK

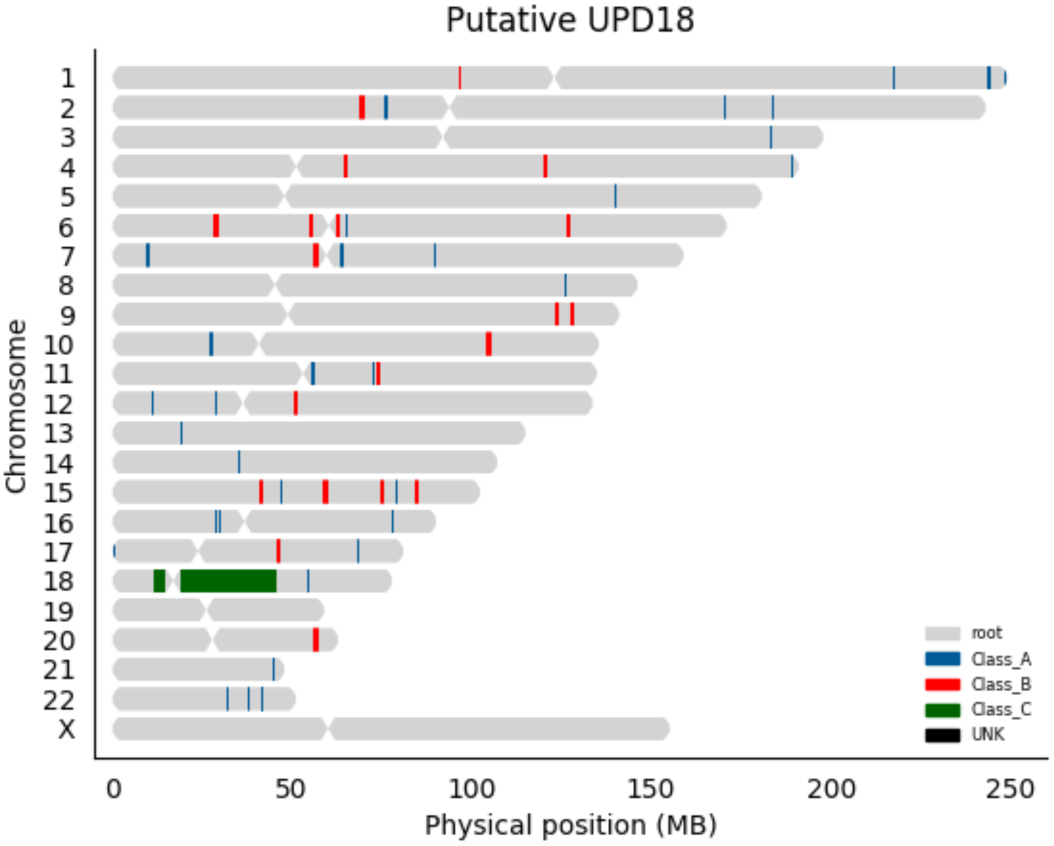
Putative UPD18
Chromosome
X
0
50
100
150
200
250
Physical position (MB)
root
Class_A
Class_B
Class_C
UNK

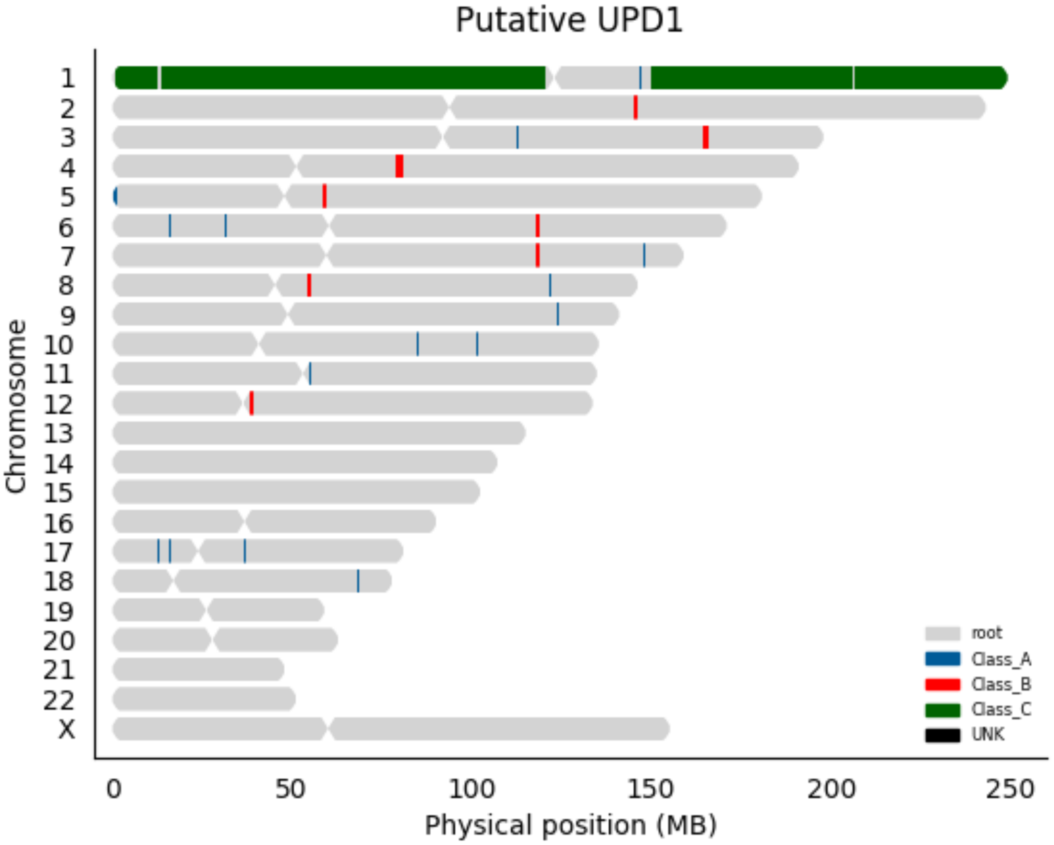

Putative UPD1
Chromosome
X
0
50
100
150
200
250
Physical position (MB)
root
Class_A
Class_B
Class_C
UNK

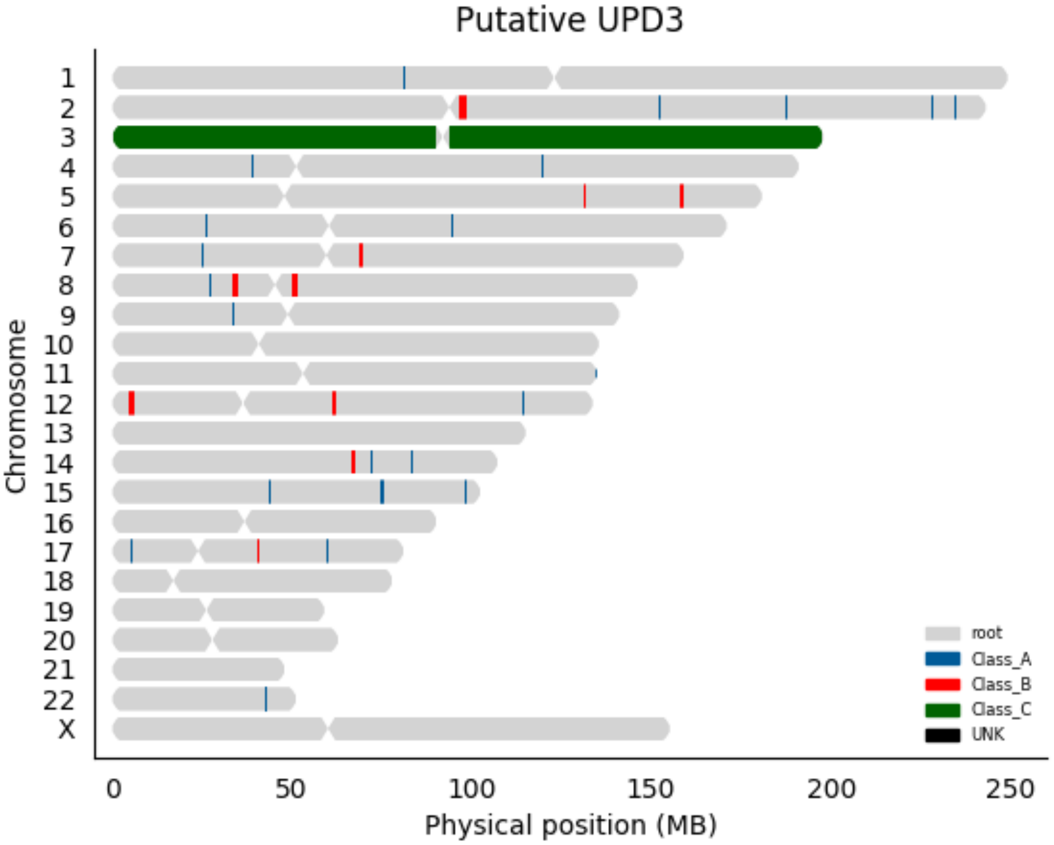
Putative UPD3
Chromosome
X
root
Class_A
Class_B
Class_C
UNK
0
50
100
150
200
250
Physical position (MB)

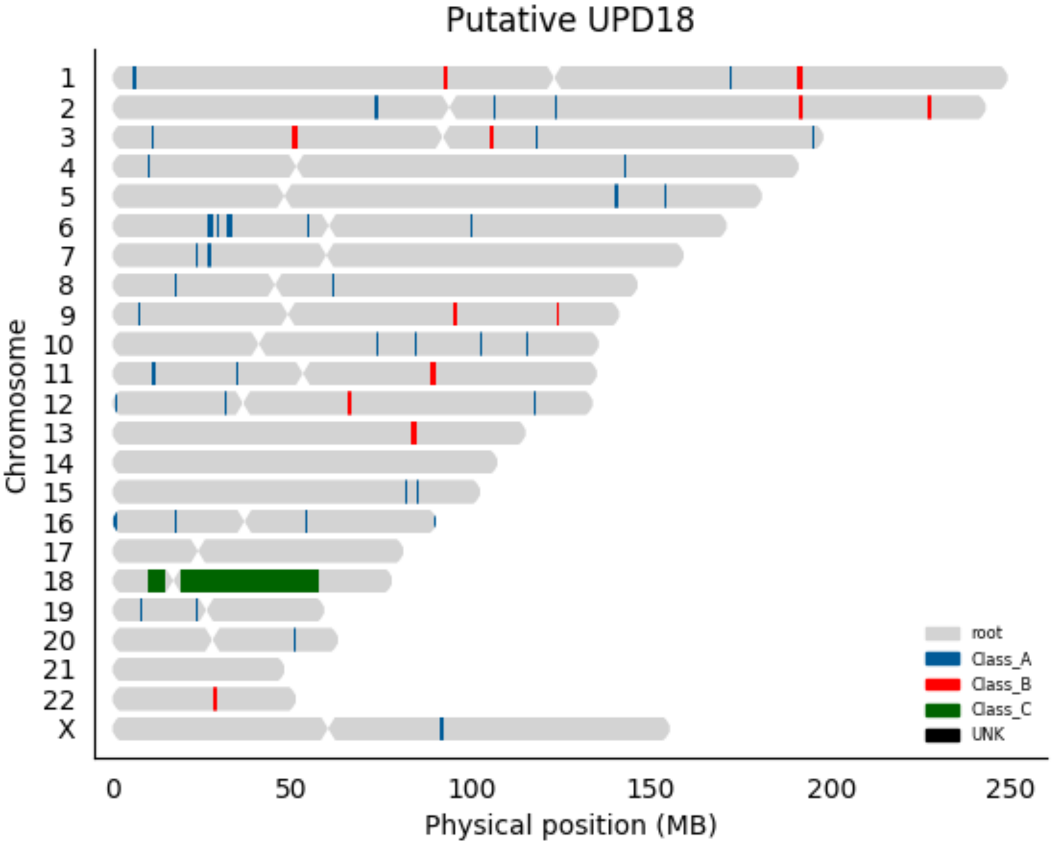

Putative UPD18
Chromosome
X
0
50
100
150
200
250
Physical position (MB)
root
Class_A
Class_B
Class_C
UNK

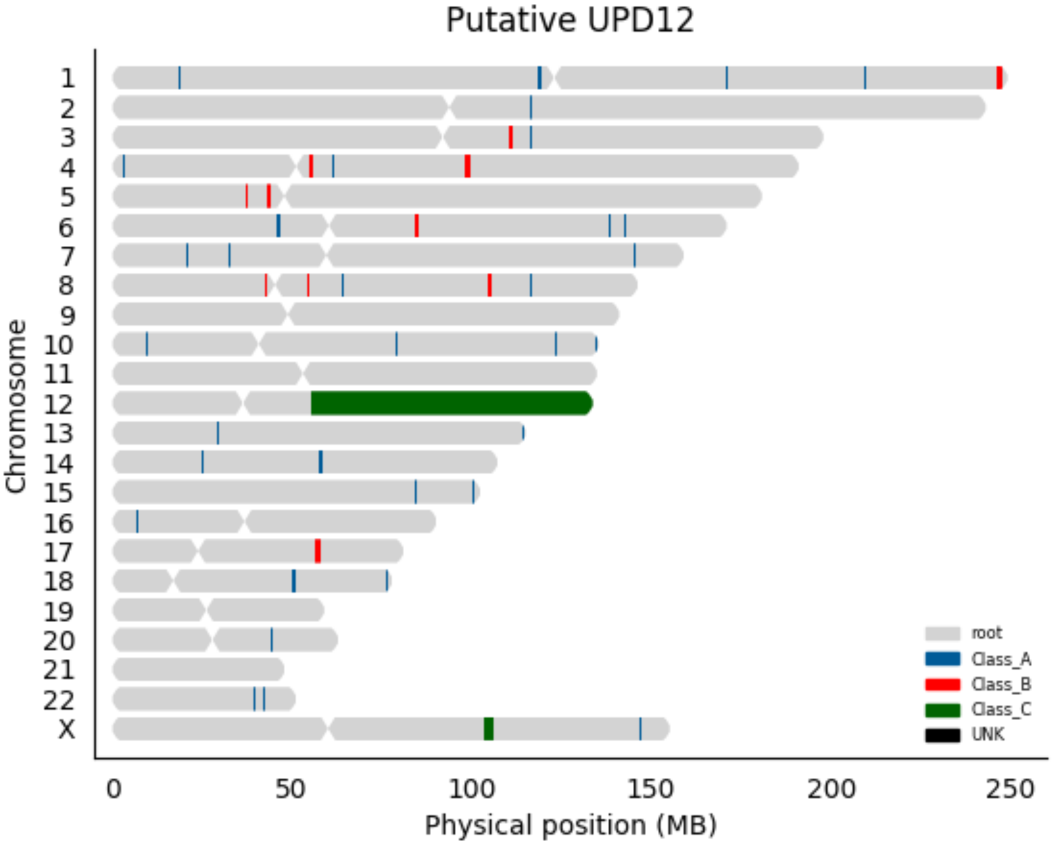

Putative UPD12
Chromosome
X
0
50
100
150
200
250
Physical position (MB)
root
Class_A
Class_B
Class_C
UNK

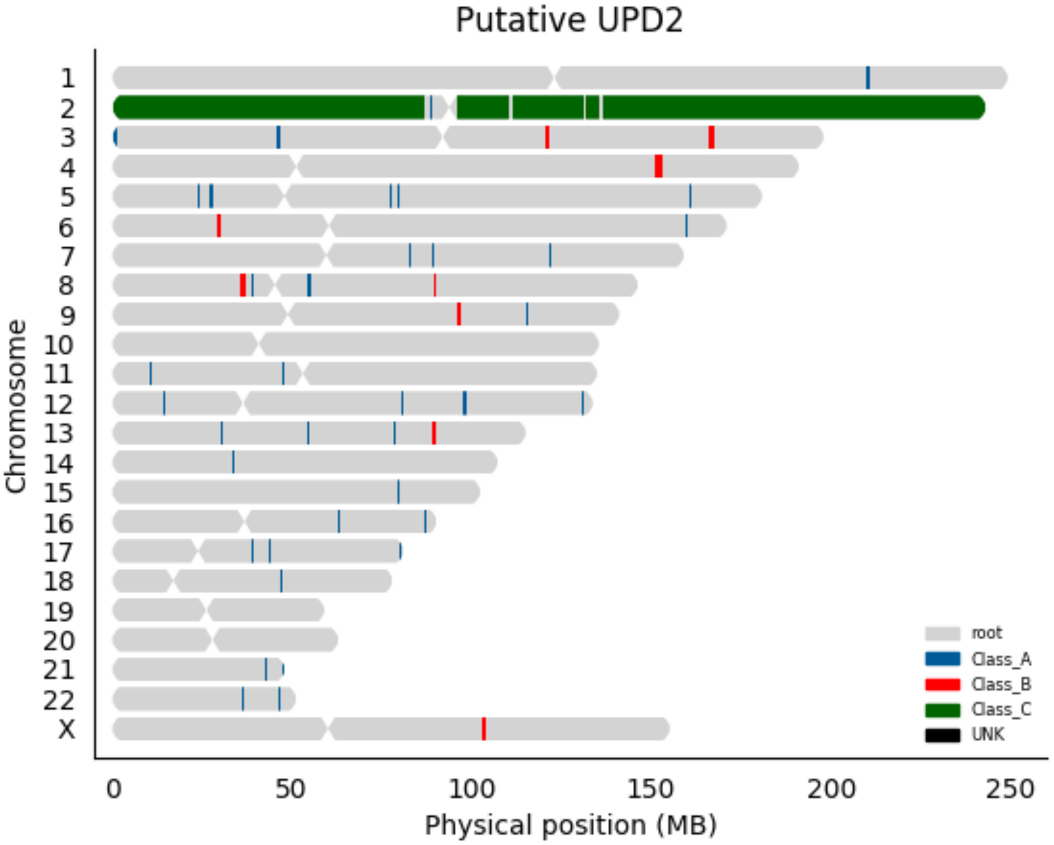

Putative UPD2
Chromosome
X
0
50
100
150
200
250
Physical position (MB)
root
Class_A
Class_B
Class_C
UNK

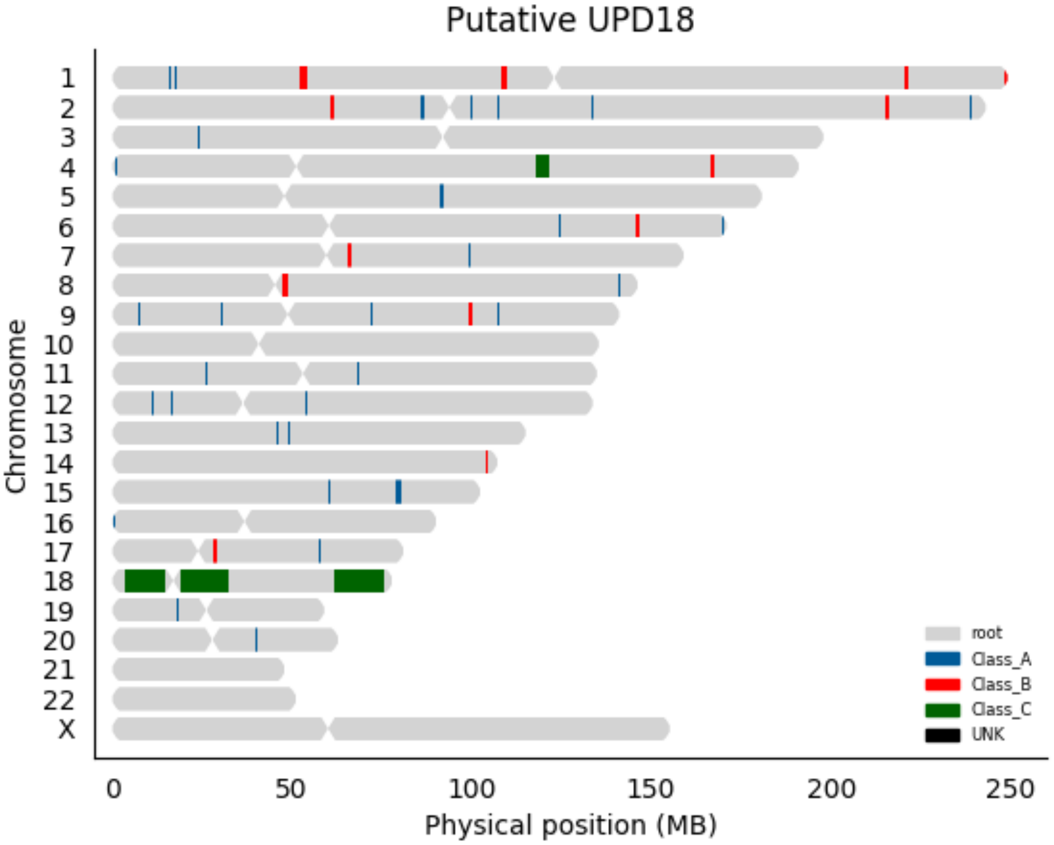

Putative UPD18
Chromosome
X
0
50
100
150
200
250
Physical position (MB)
root
Class_A
Class_B
Class_C
UNK

Putative UPD12

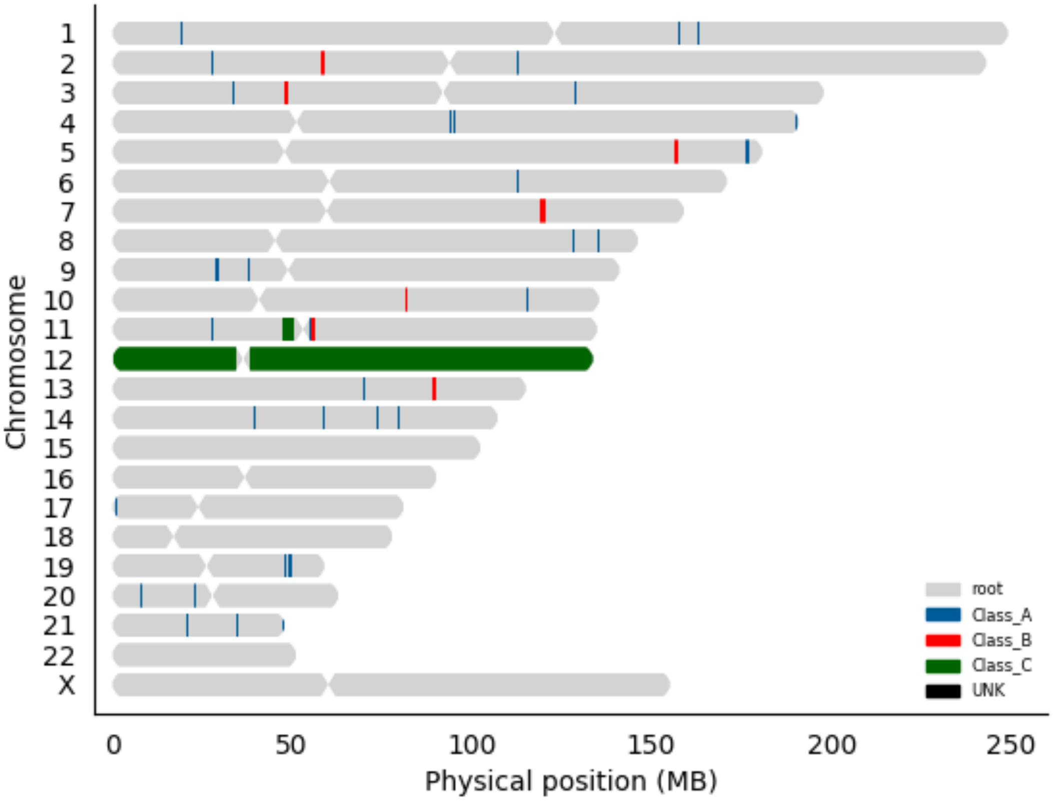

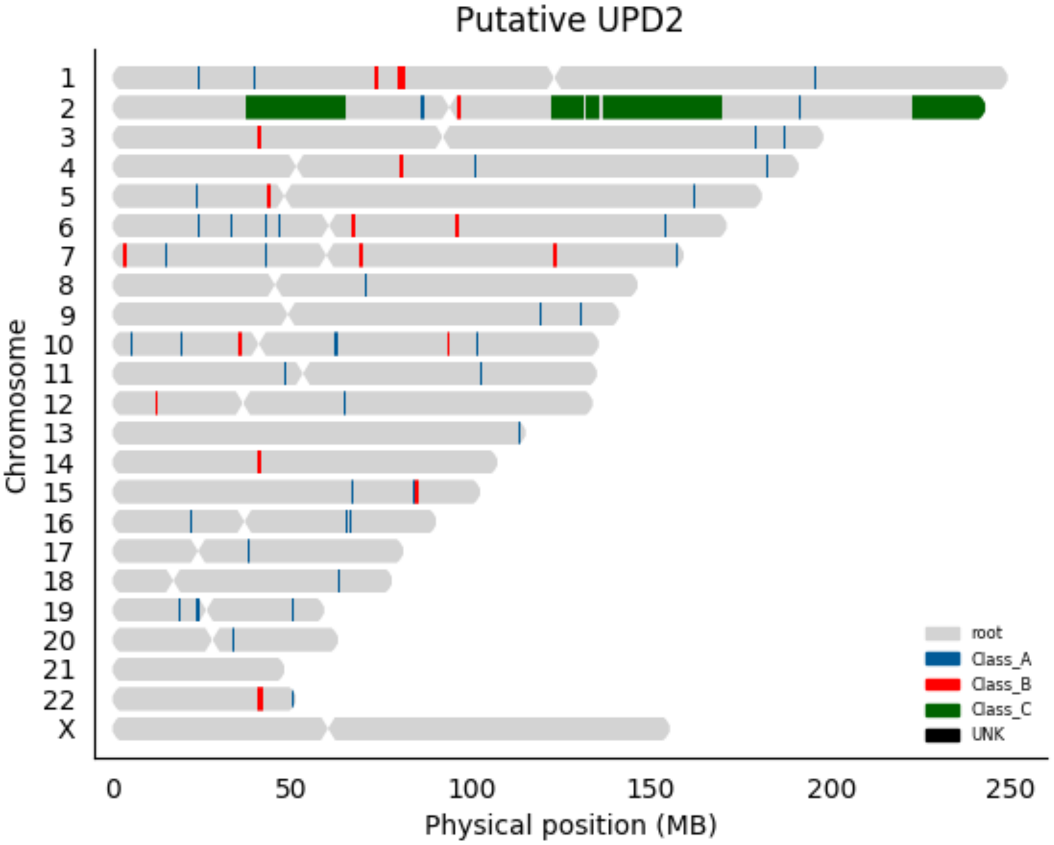

Putative UPD2
Chromosome
X
0
50
100
150
200
250
Physical position (MB)
root
Class_A
Class_B
Class_C
UNK

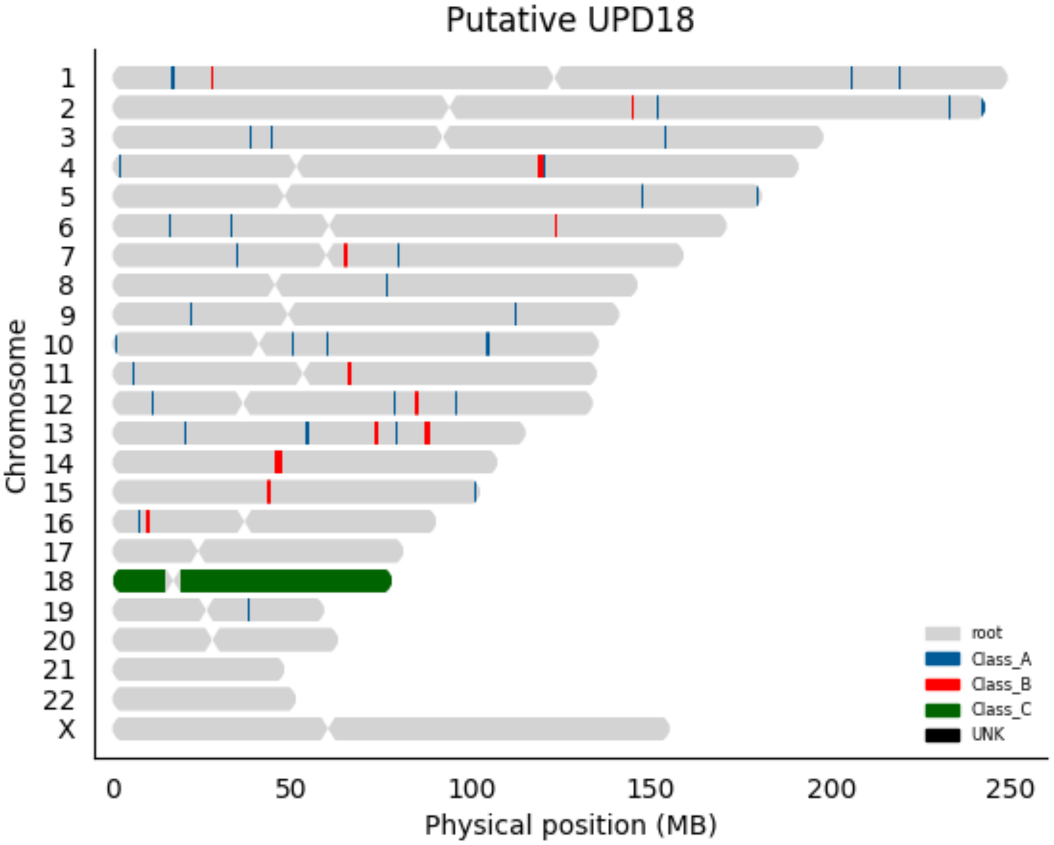

Putative UPD18
Chromosome
X
0
50
100
150
200
250
Physical position (MB)
root
Class_A
Class_B
Class_C
UNK

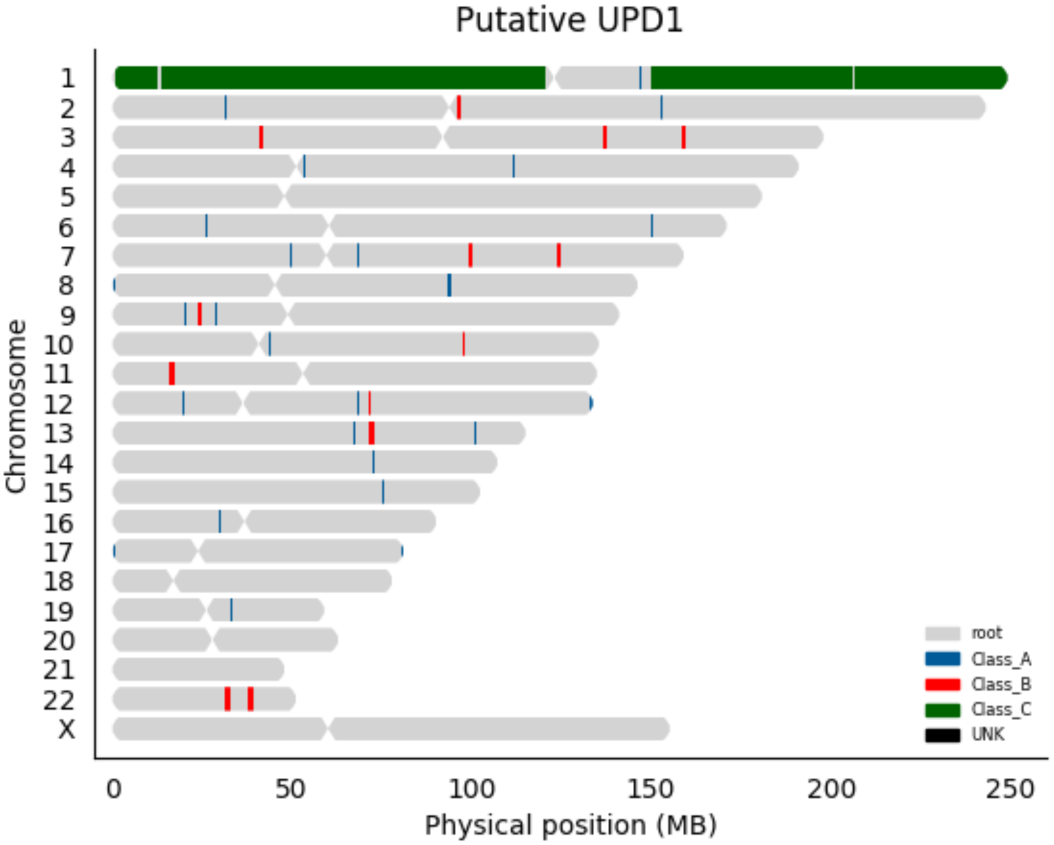

Putative UPD1
Chromosome
X
0
50
100
150
200
250
Physical position (MB)
root
Class_A
Class_B
Class_C
UNK

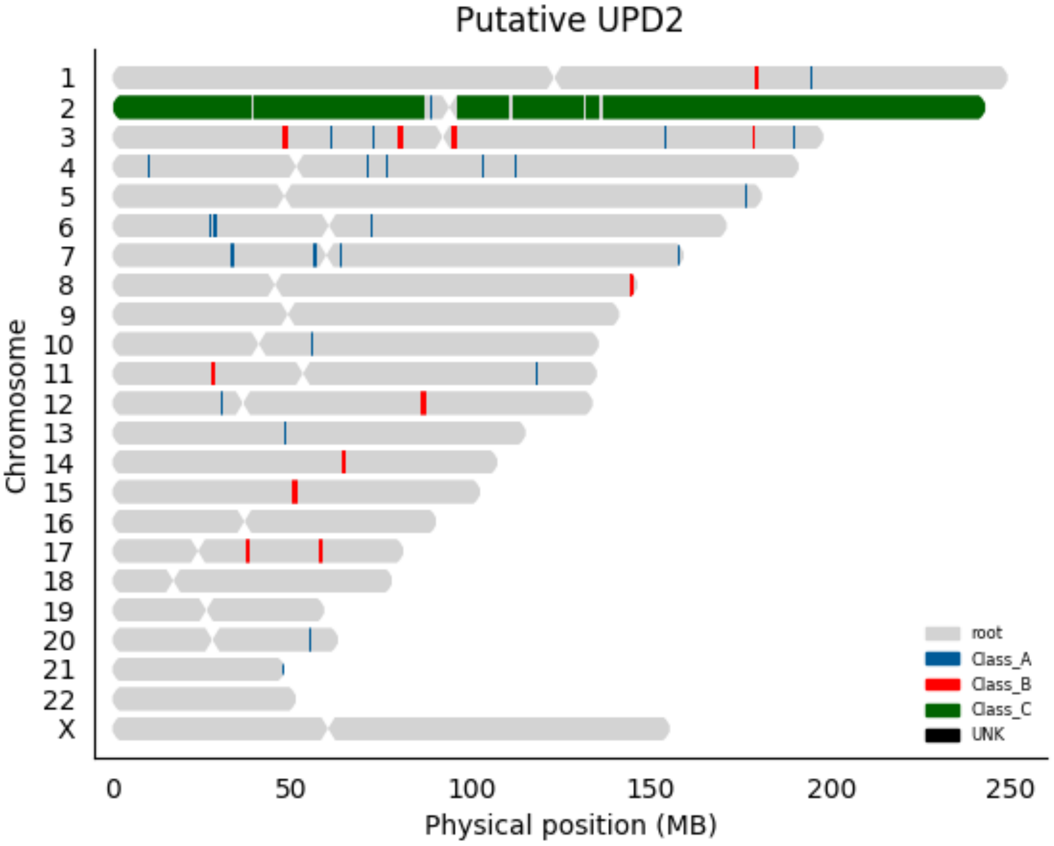

Putative UPD2
Chromosome
X
root
Class_A
Class_B
Class_C
UNK
0
50
100
150
200
250
Physical position (MB)

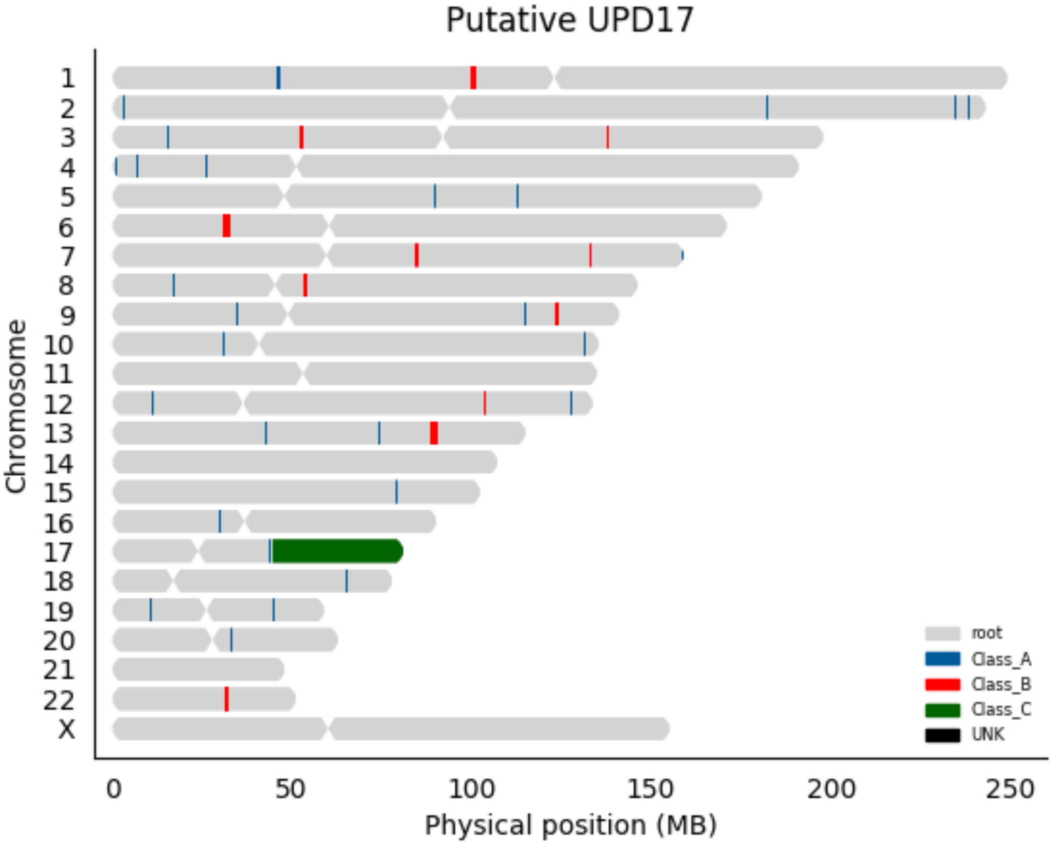
Putative UPD17
Chromosome
X
0
50
100
150
200
250
Physical position (MB)
root
Class_A
Class_B
Class_C
UNK

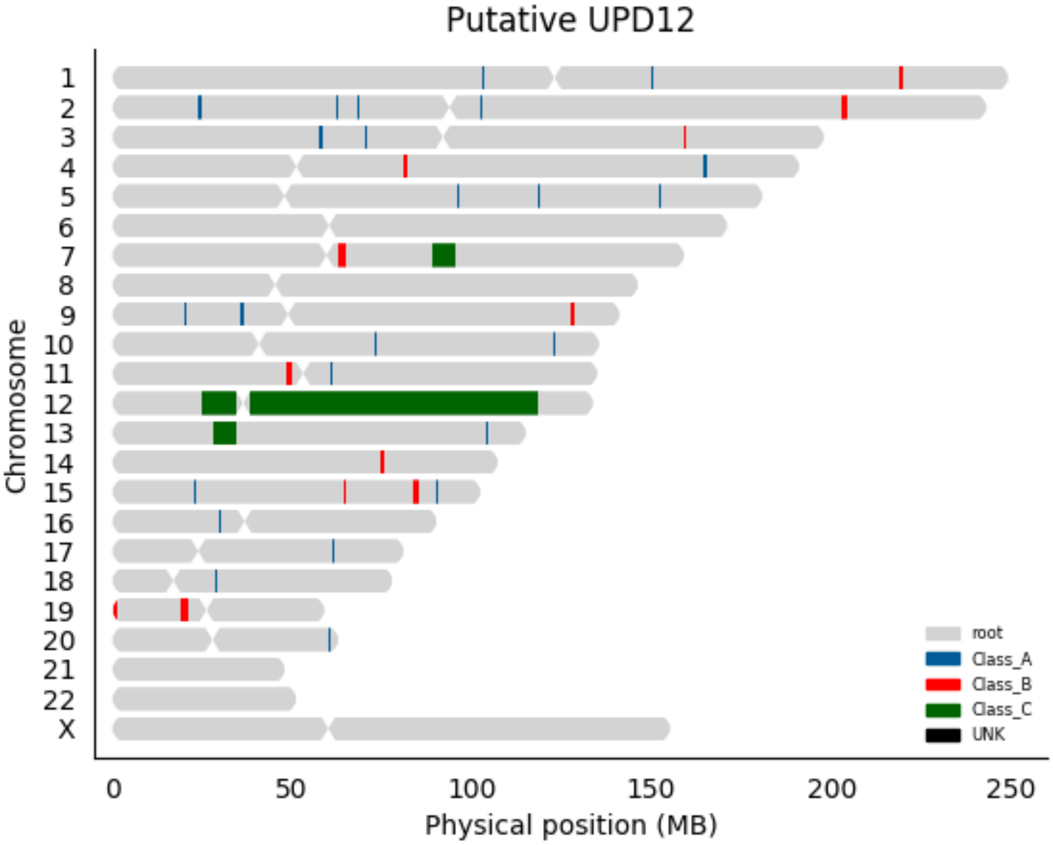

Putative UPD12
Chromosome
X
0
50
100
150
200
250
Physical position (MB)
root
Class_A
Class_B
Class_C
UNK

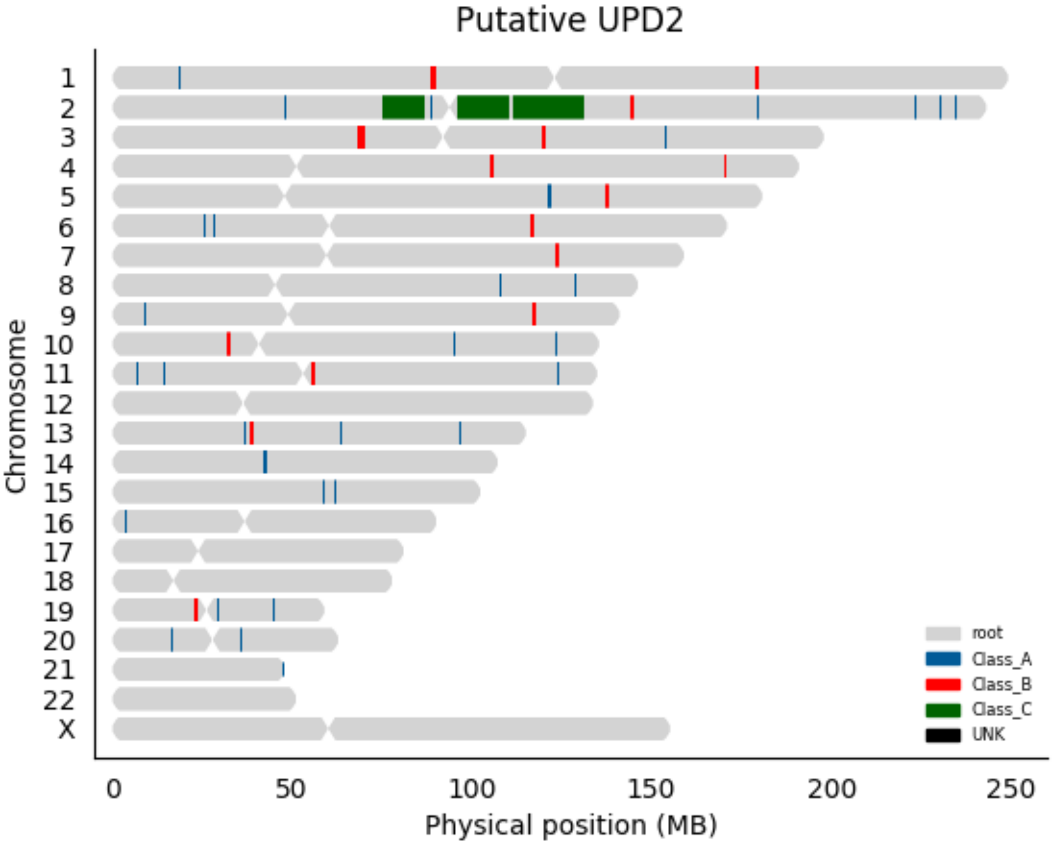

Putative UPD2
Chromosome
X
0
50
100
150
200
250
Physical position (MB)
root
Class_A
Class_B
Class_C
UNK

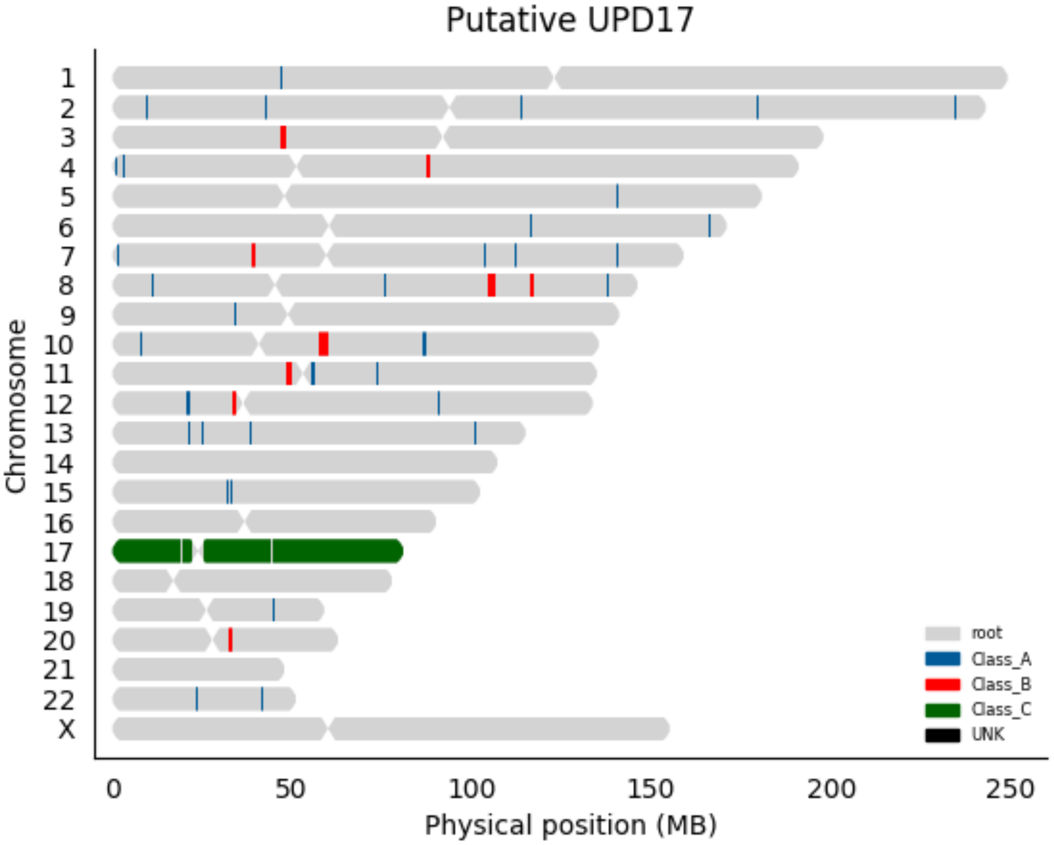

Putative UPD17
Chromosome
X
0
50
100
150
200
250
Physical position (MB)
root
Class_A
Class_B
Class_C
UNK

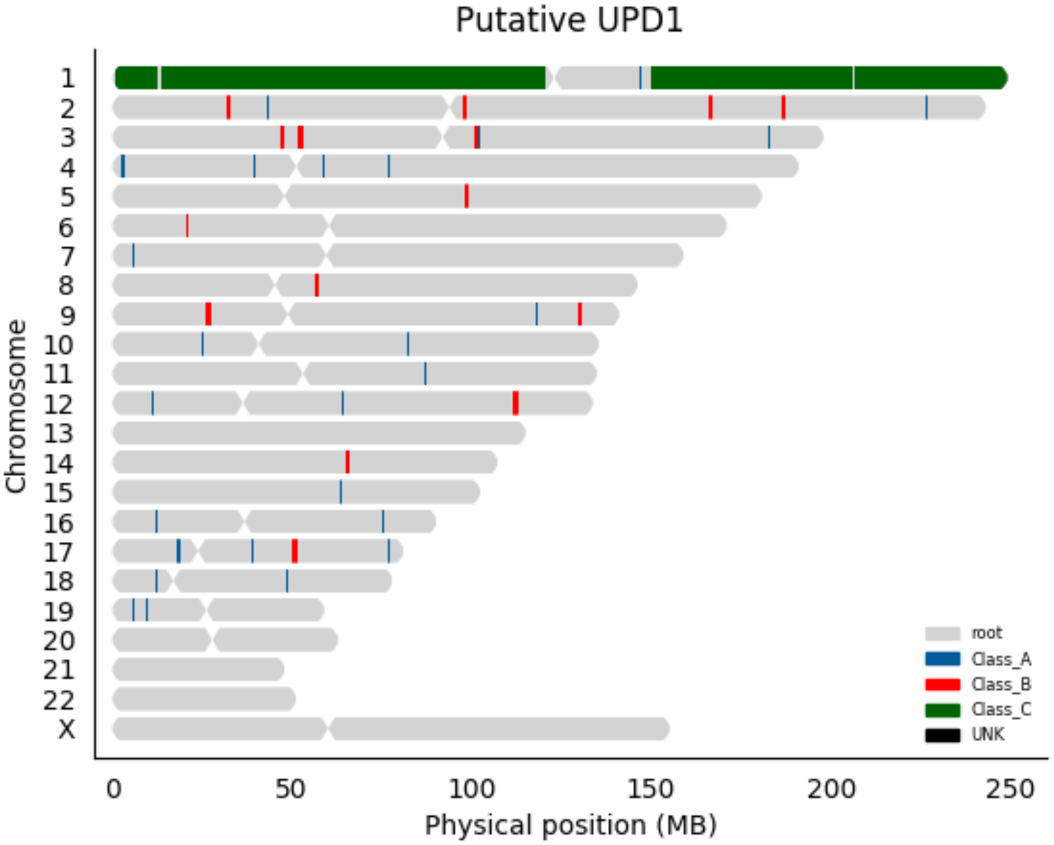

Putative UPD1
Chromosome
X
0
50
100
150
200
250
Physical position (MB)
root
Class_A
Class_B
Class_C
UNK

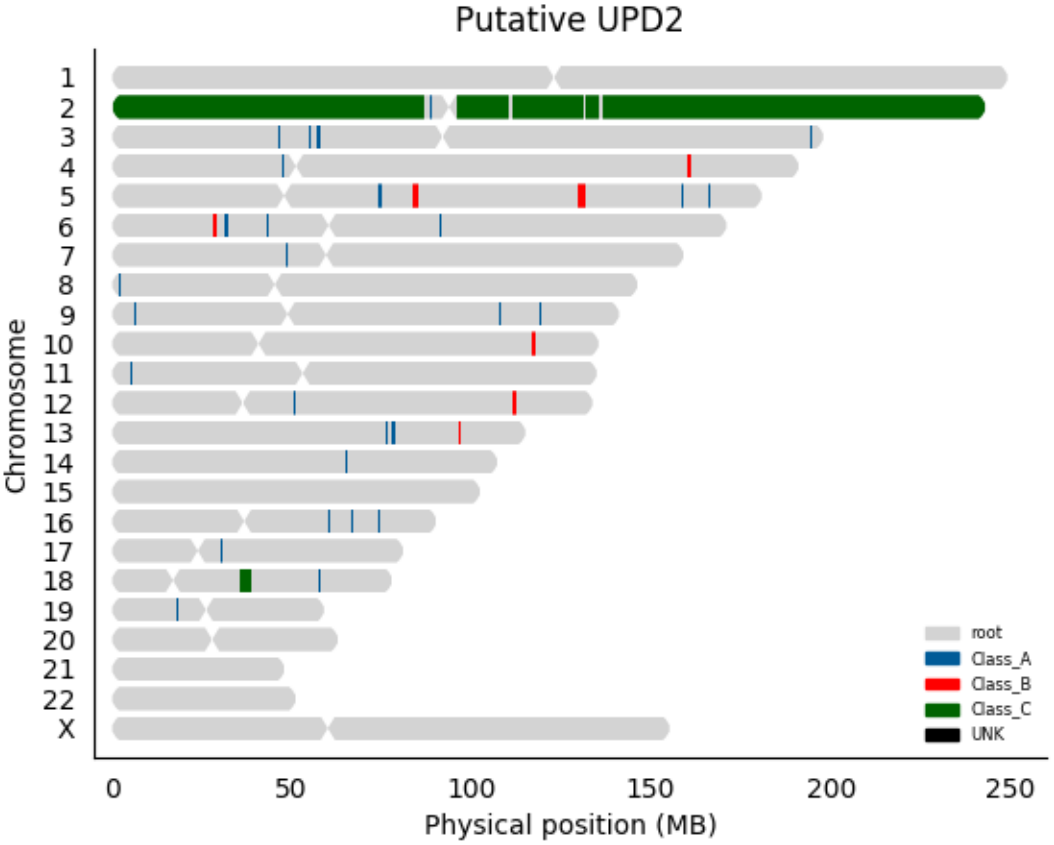
Putative UPD2
Chromosome
X
0
50
100
150
200
250
Physical position (MB)
root
Class_A
Class_B
Class_C
UNK

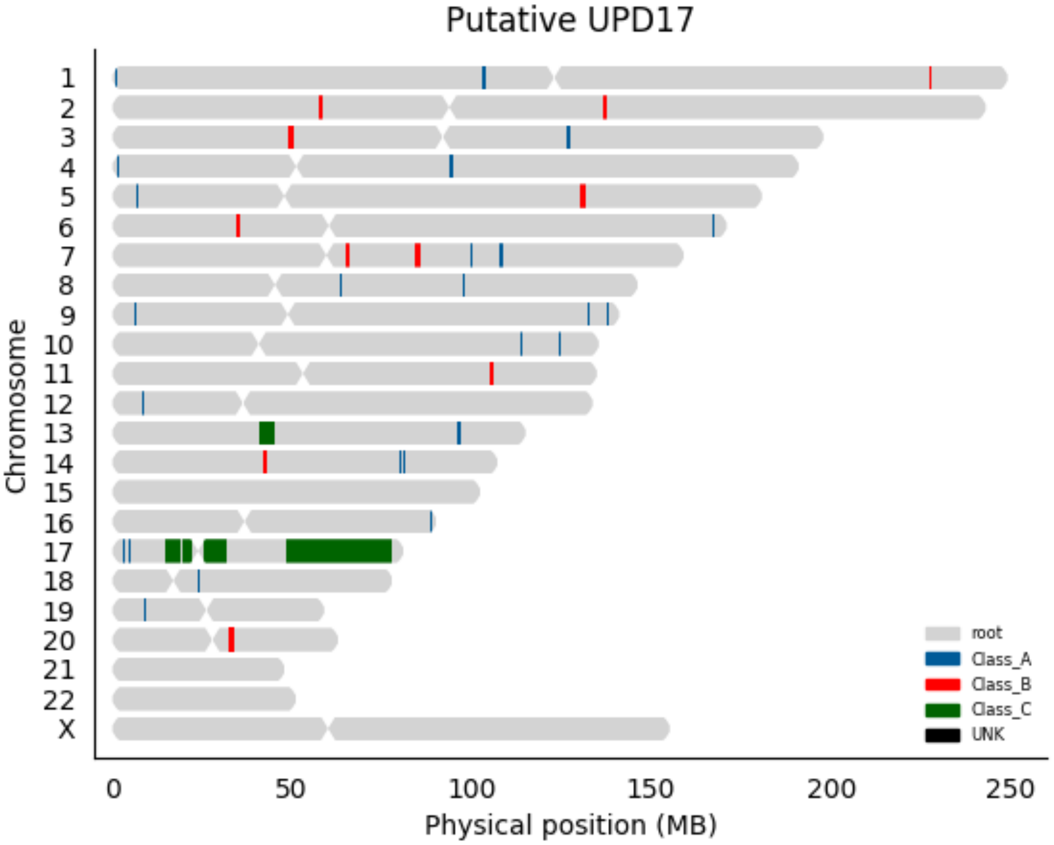

Putative UPD17
Chromosome
X
0
50
100
150
200
250
Physical position (MB)
root
Class_A
Class_B
Class_C
UNK

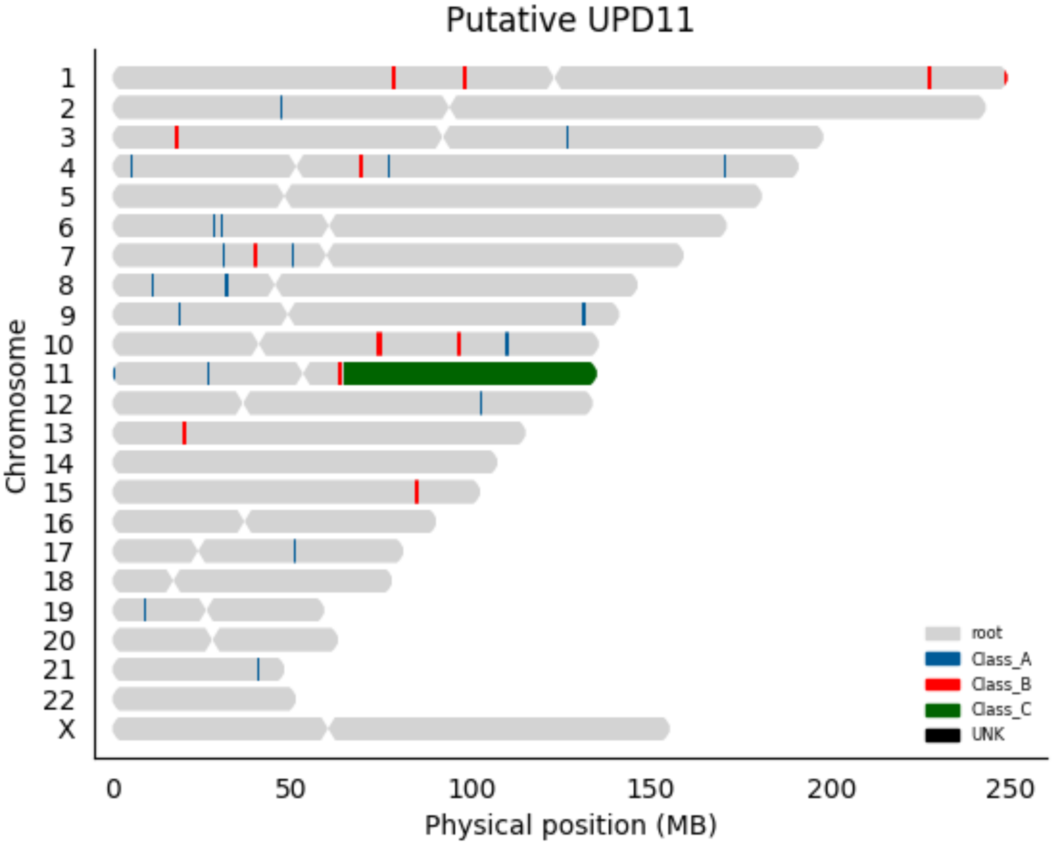

Putative UPD11
Chromosome
X
0
50
100
150
200
250
Physical position (MB)
root
Class_A
Class_B
Class_C
UNK

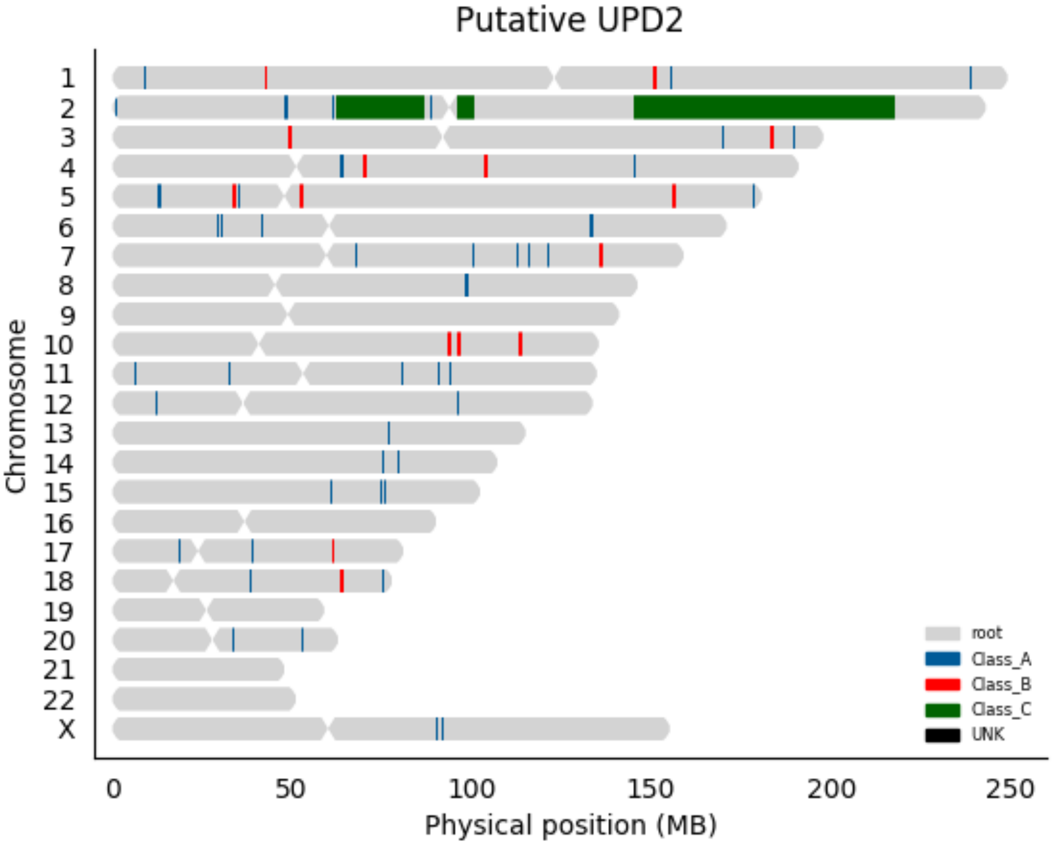
Putative UPD2
Chromosome
X
0
50
100
150
200
250
Physical position (MB)
root
Class_A
Class_B
Class_C
UNK

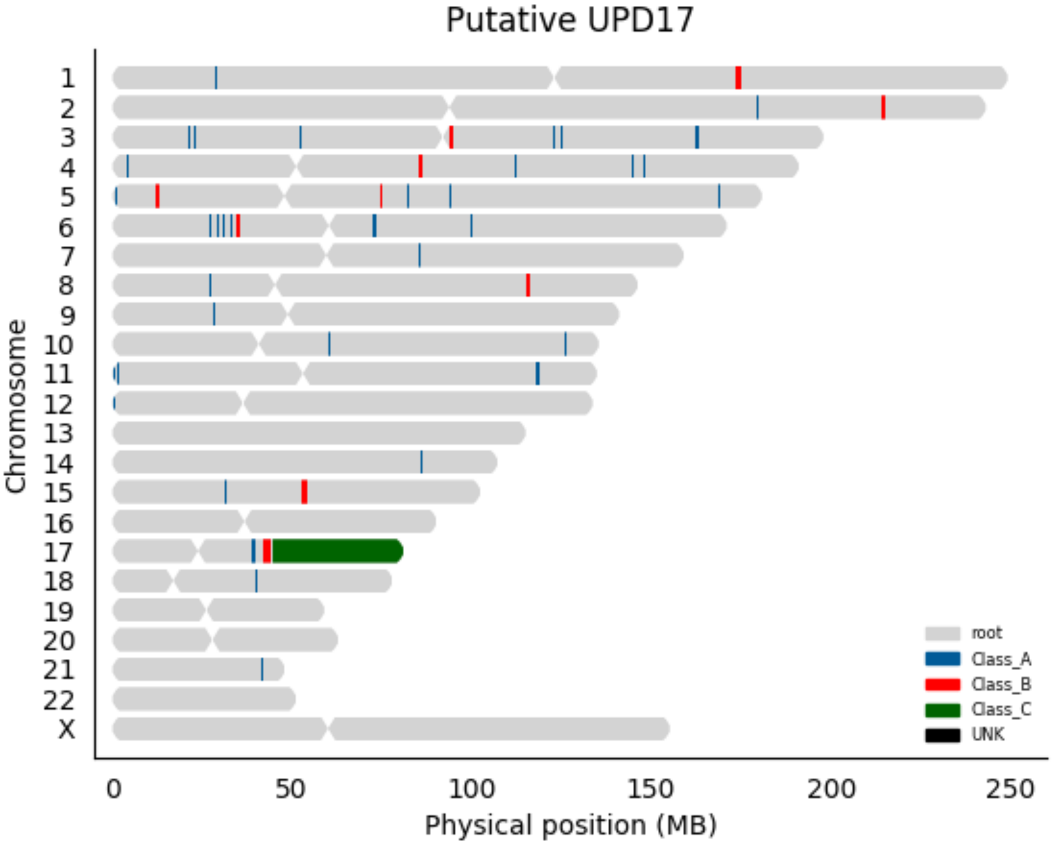
Putative UPD17
Chromosome
X
0
50
100
150
200
250
Physical position (MB)
root
Class_A
Class_B
Class_C
UNK

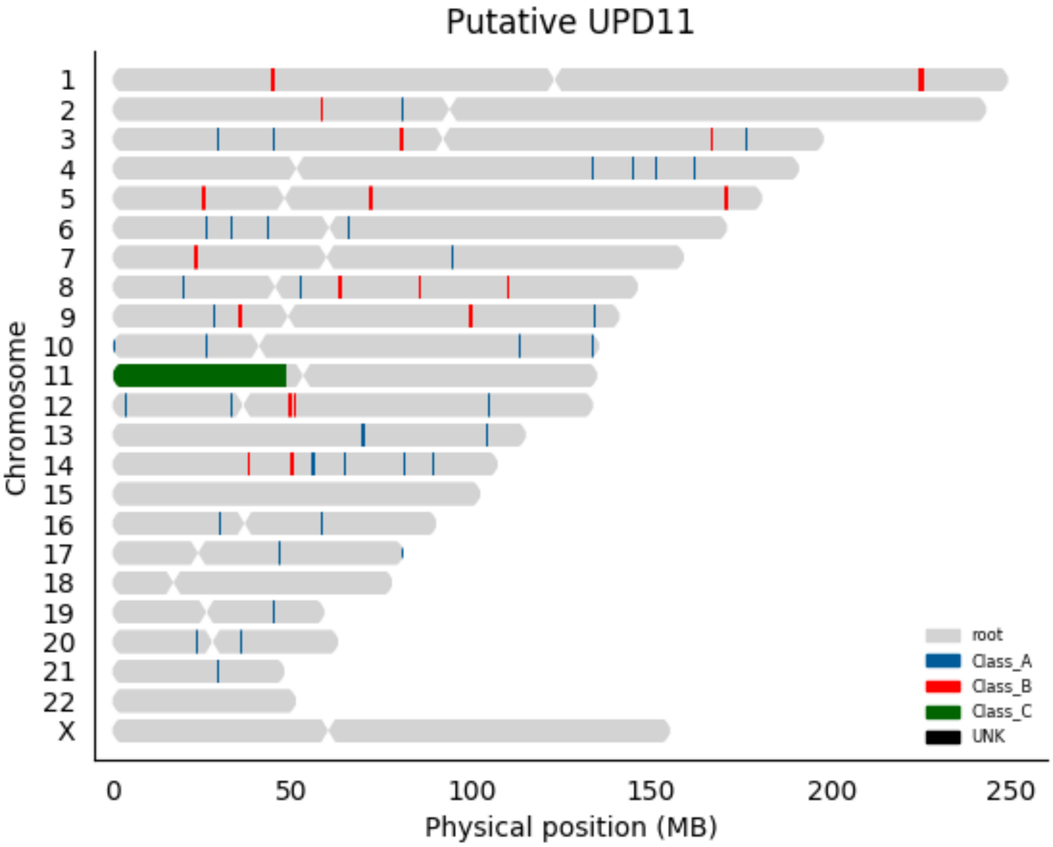

Putative UPD11
Chromosome
X
0
50
100
150
200
250
Physical position (MB)
root
Class_A
Class_B
Class_C
UNK

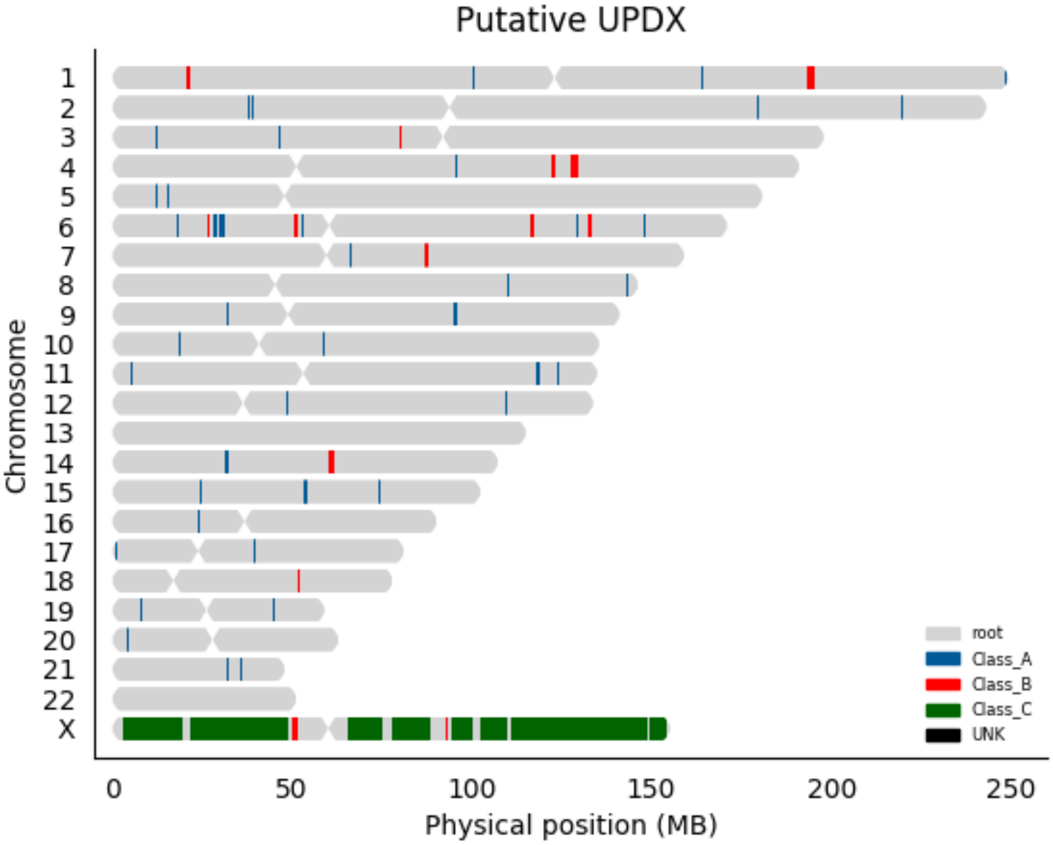

Putative UPDX
Chromosome
X
0
50
100
150
200
250
Physical position (MB)
root
Class_A
Class_B
Class_C
UNK

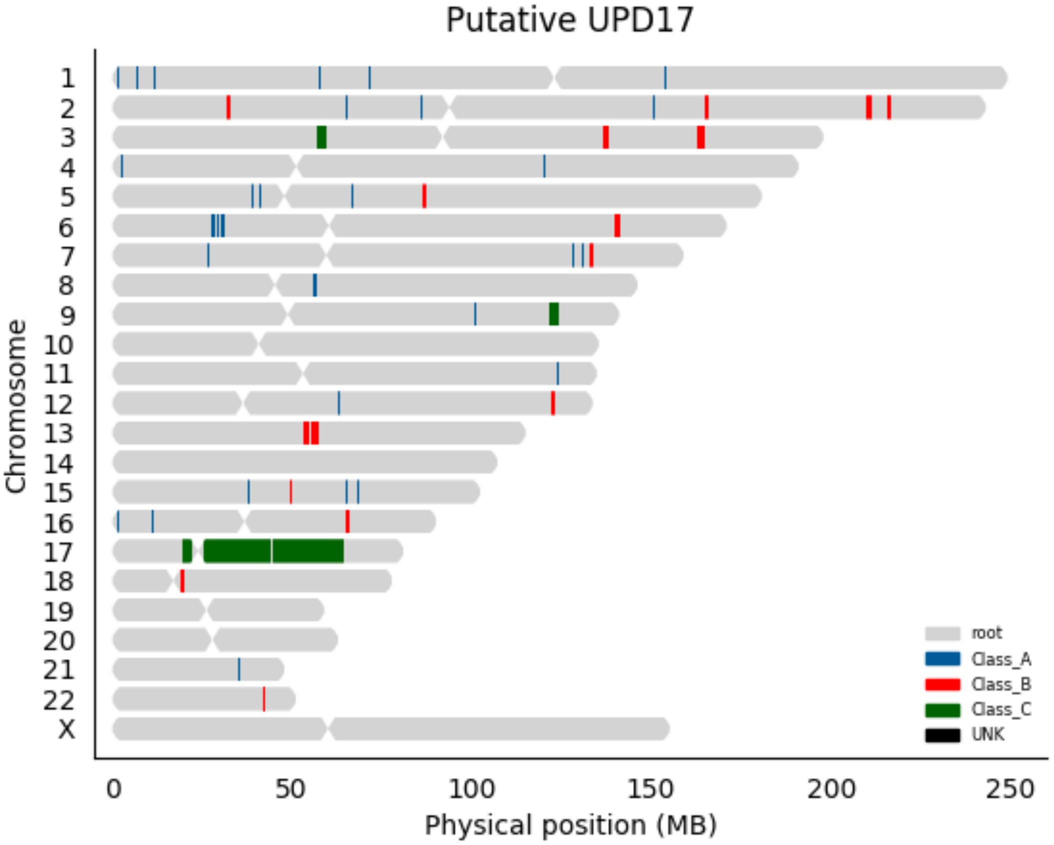

Putative UPD17
X
Chromosome
0
50
100
150
200
250
Physical position (MB)
root
Class_A
Class_B
Class_C
UNK

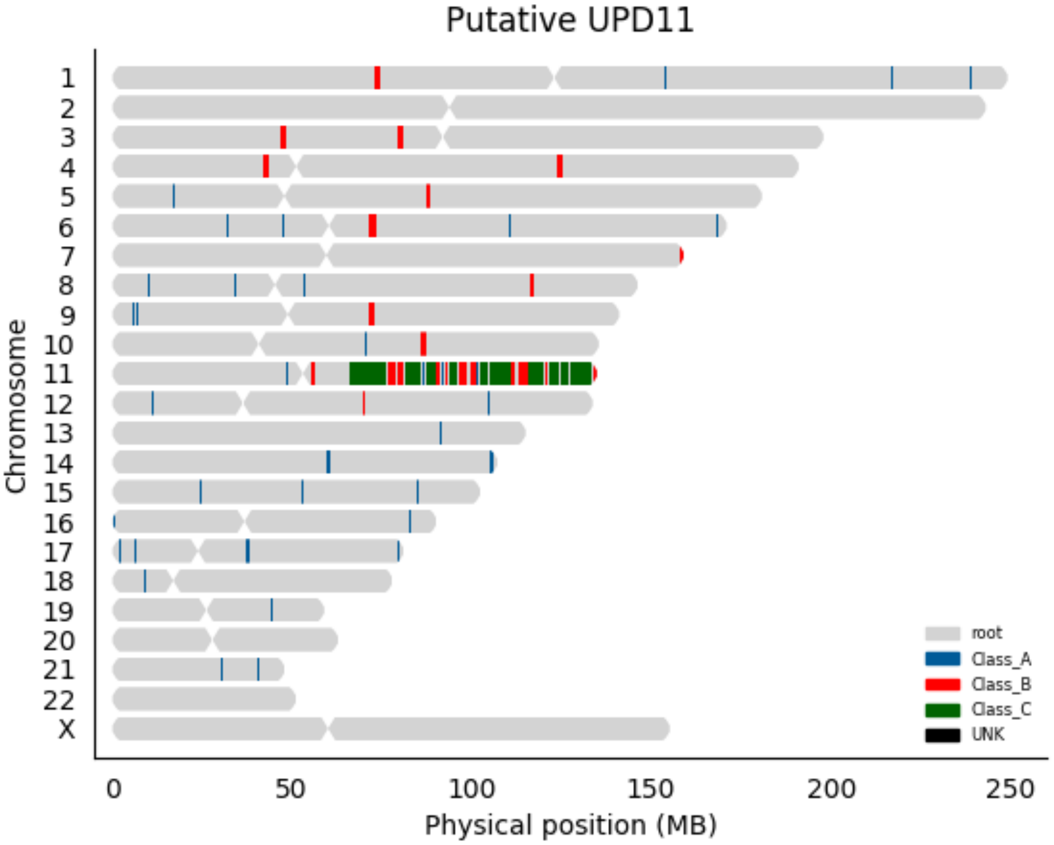

Putative UPD11
Chromosome
X
0
50
100
150
200
250
Physical position (MB)
root
Class_A
Class_B
Class_C
UNK

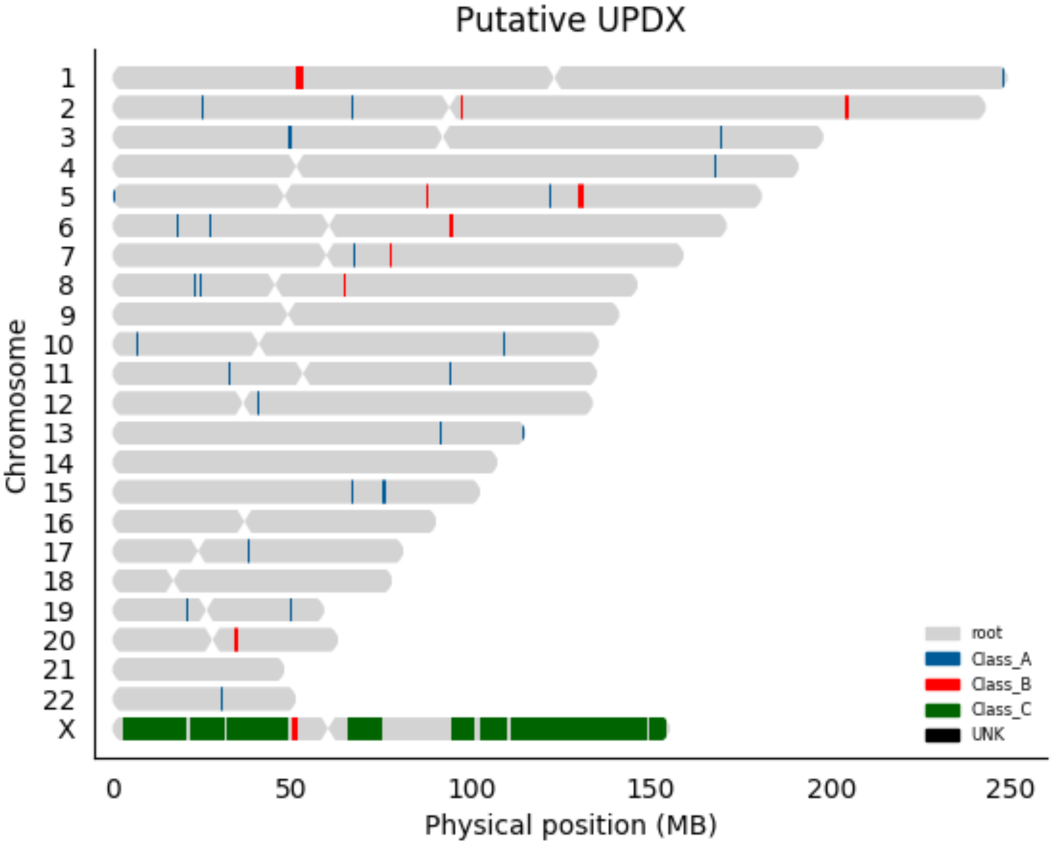

Putative UPDX
Chromosome
X
0
50
100
150
200
250
Physical position (MB)
root
Class_A
Class_B
Class_C
UNK

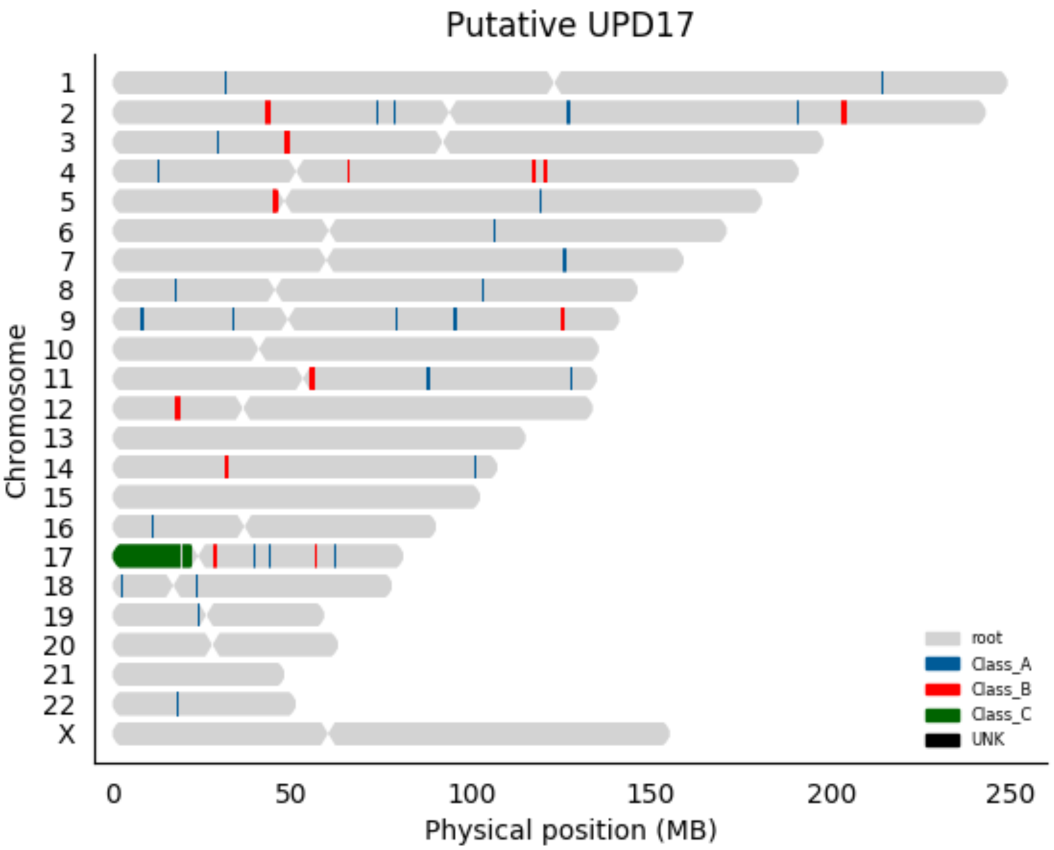

Putative UPD17
Chromosome
X
0
50
100
150
200
250
Physical position (MB)
root
Class_A
Class_B
Class_C
UNK

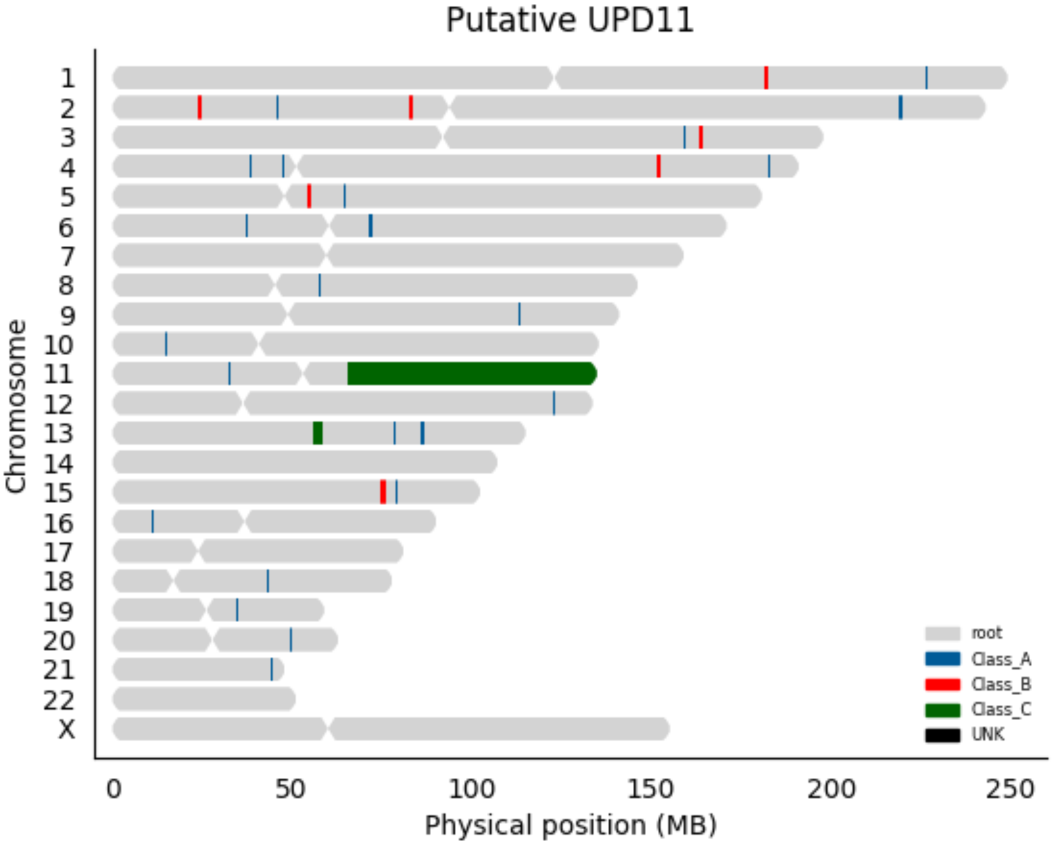

Putative UPD11
Chromosome
X
0
50
100
150
200
250
Physical position (MB)
root
Class_A
Class_B
Class_C
UNK

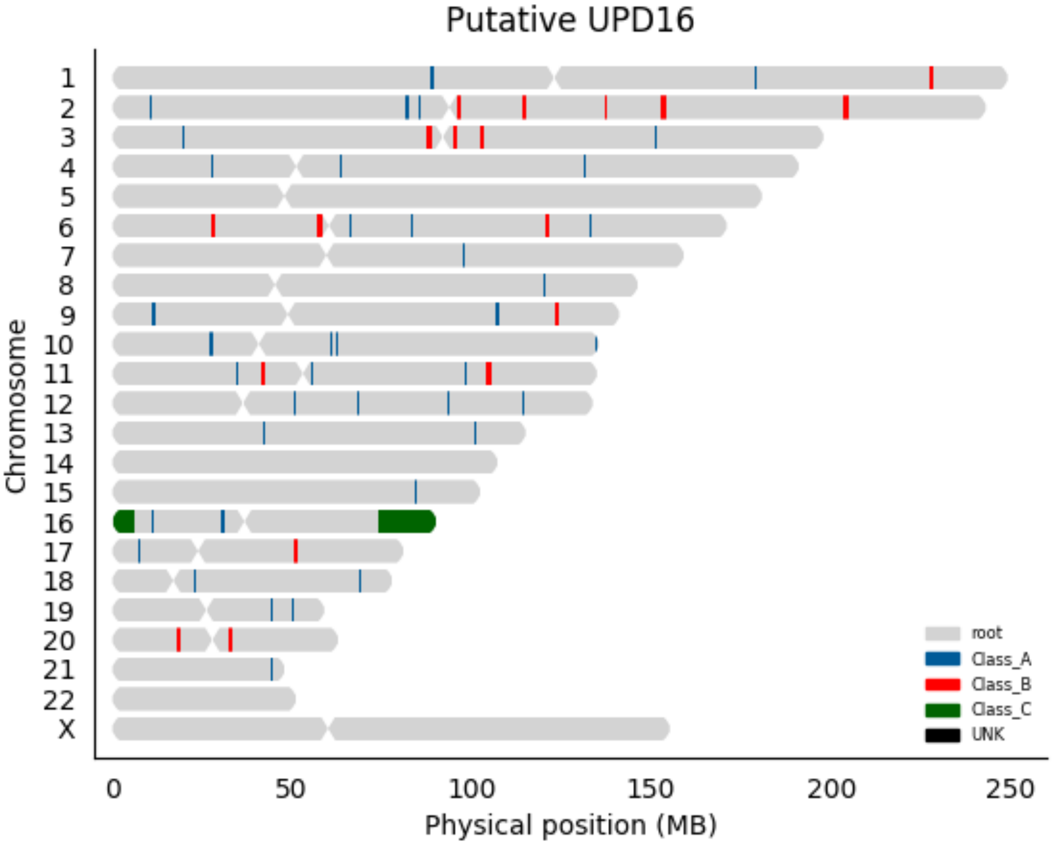

Putative UPD16
Chromosome
X
0
50
100
150
200
250
Physical position (MB)
root
Class_A
Class_B
Class_C
UNK

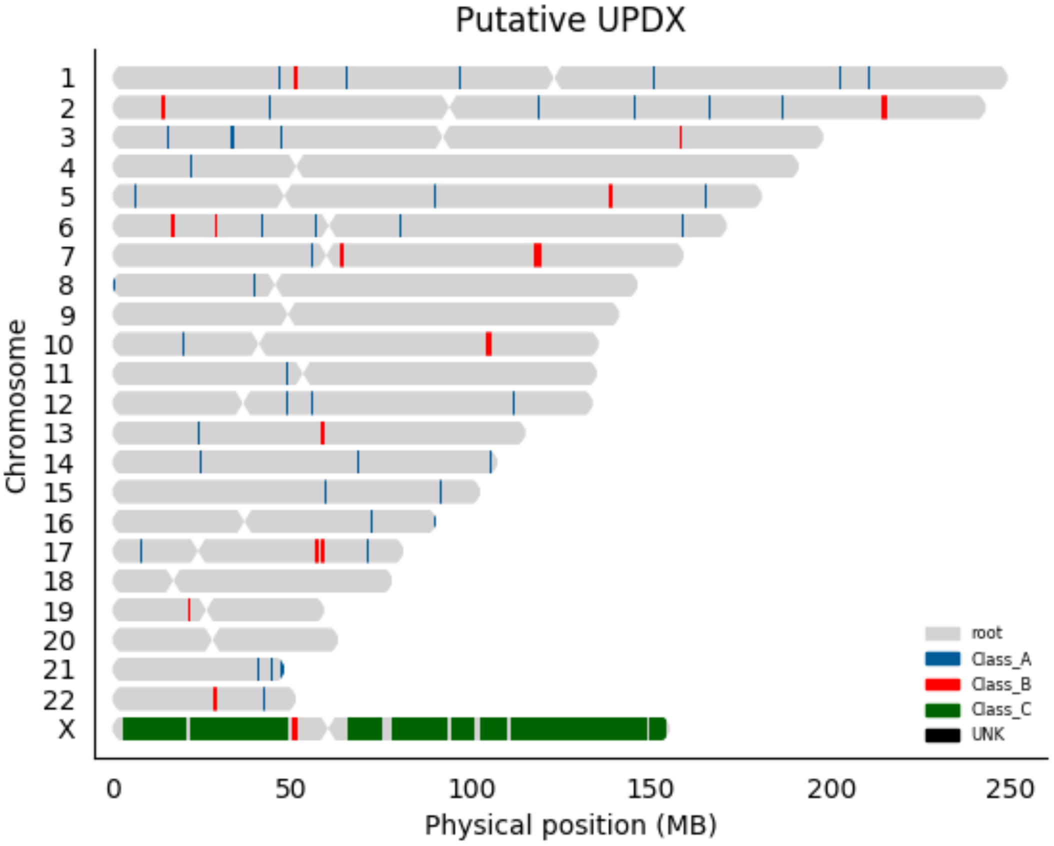

Putative UPDX
Chromosome
X
0
50
100
150
200
250
Physical position (MB)
root
Class_A
Class_B
Class_C
UNK

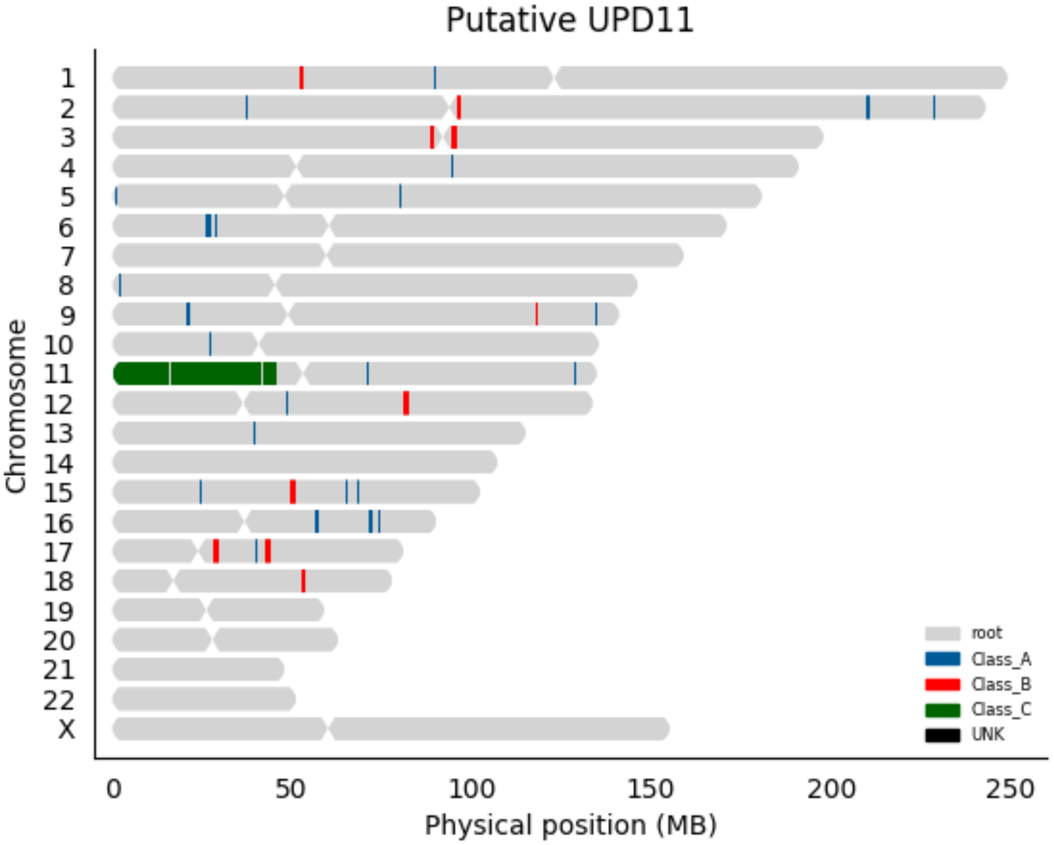

Putative UPD11
Chromosome
X
0
50
100
150
200
250
Physical position (MB)
root
Class_A
Class_B
Class_C
UNK

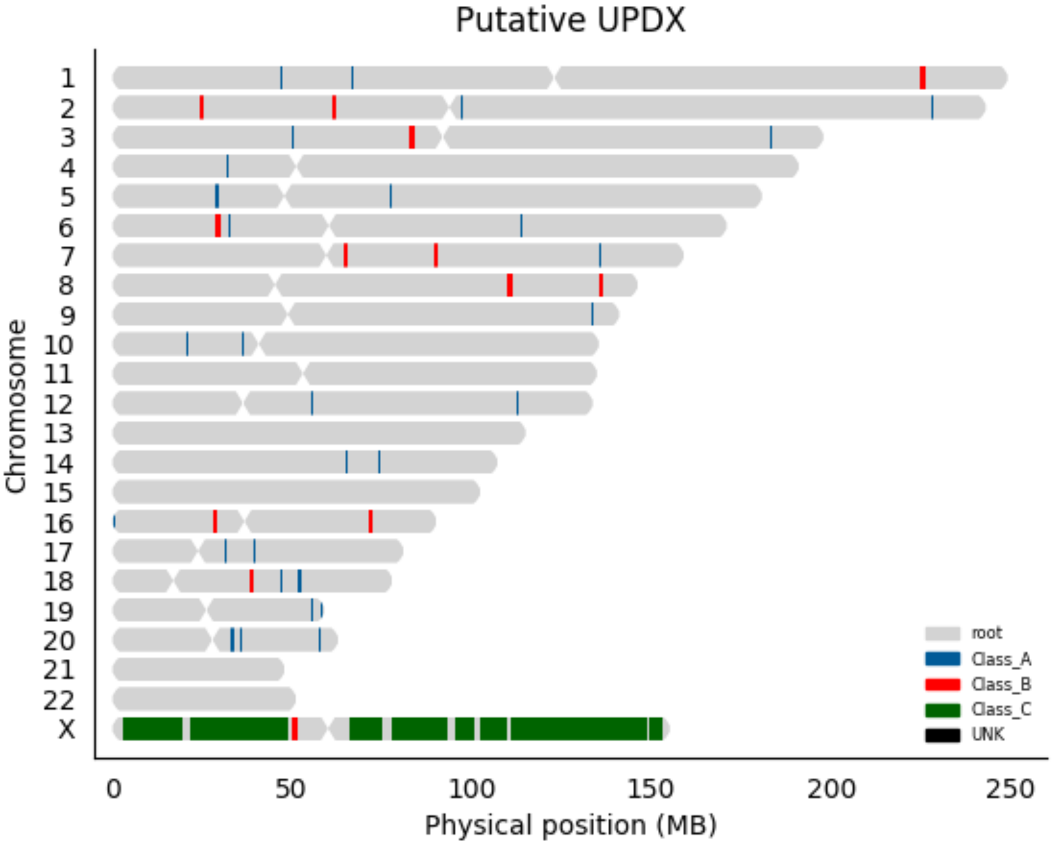

Putative UPDX
X
Chromosome
0
50
100
150
200
250
Physical position (MB)
root
Class_A
Class_B
Class_C
UNK

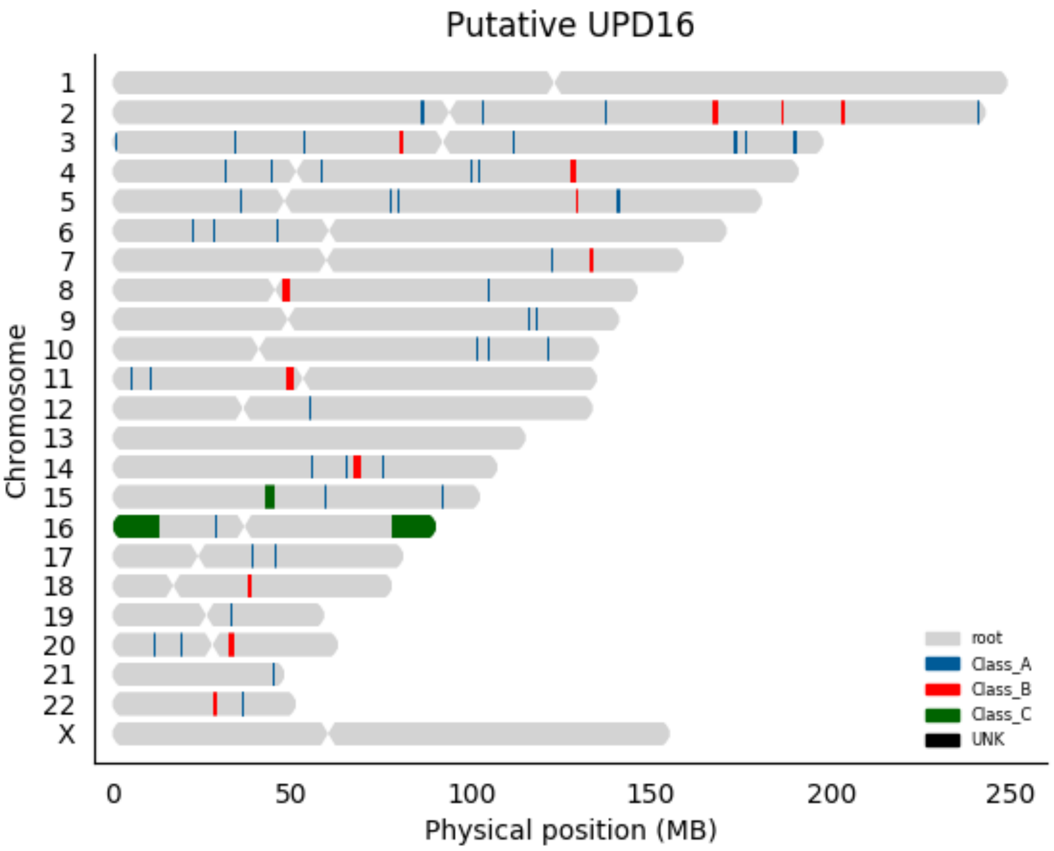
Putative UPD16
Chromosome
X
0
50
100
150
200
250
Physical position (MB)
root
Class_A
Class_B
Class_C
UNK

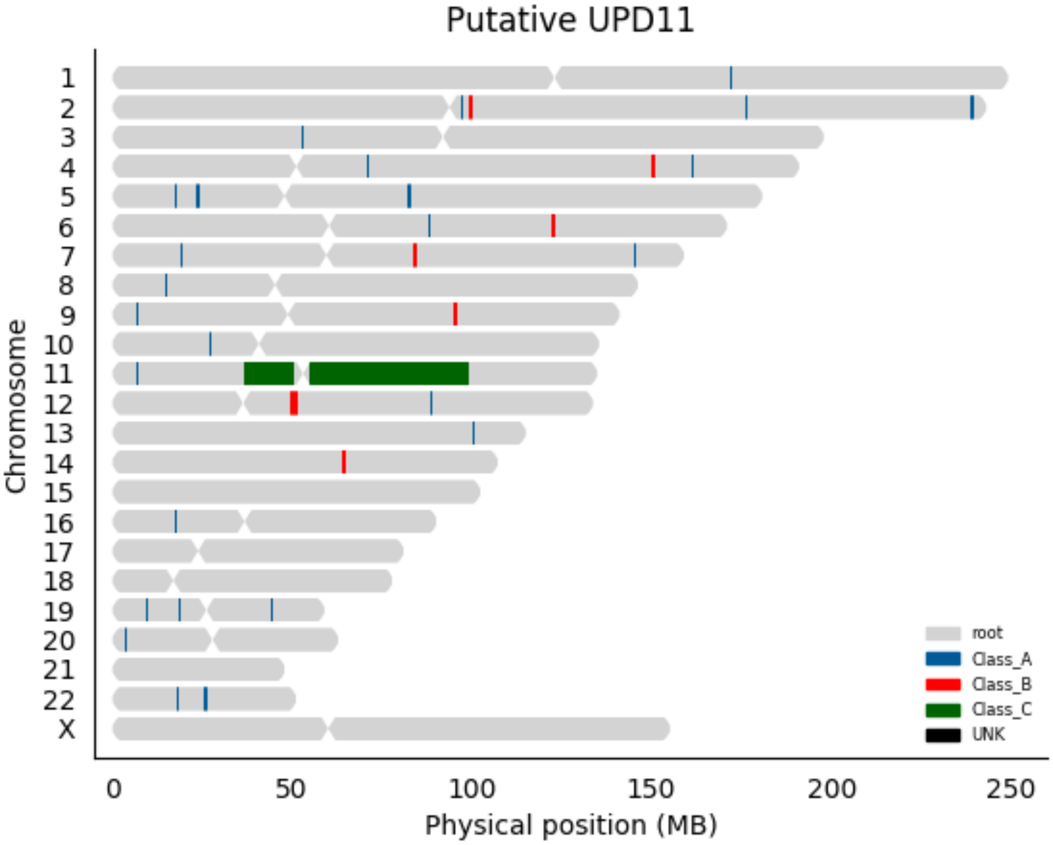

Putative UPD11
Chromosome
X
0
50
100
150
200
250
Physical position (MB)
root
Class_A
Class_B
Class_C
UNK

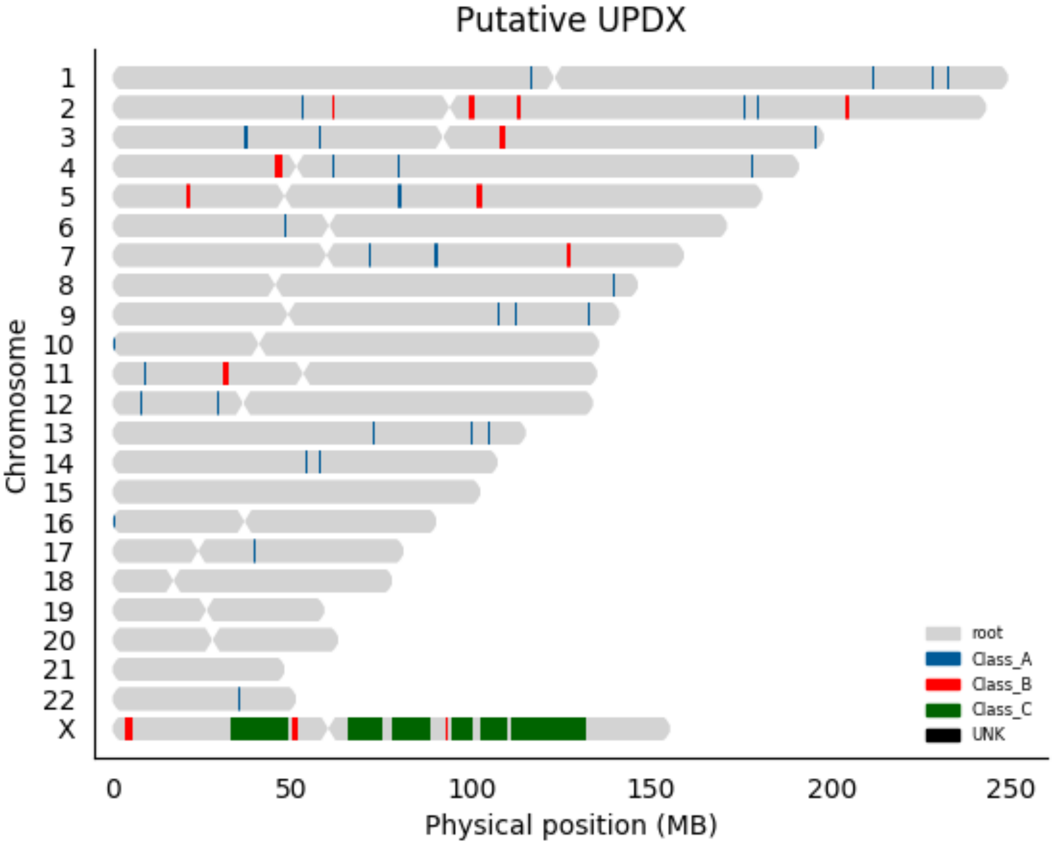

Putative UPDX
Chromosome
X
0
50
100
150
200
250
Physical position (MB)
root
Class_A
Class_B
Class_C
UNK

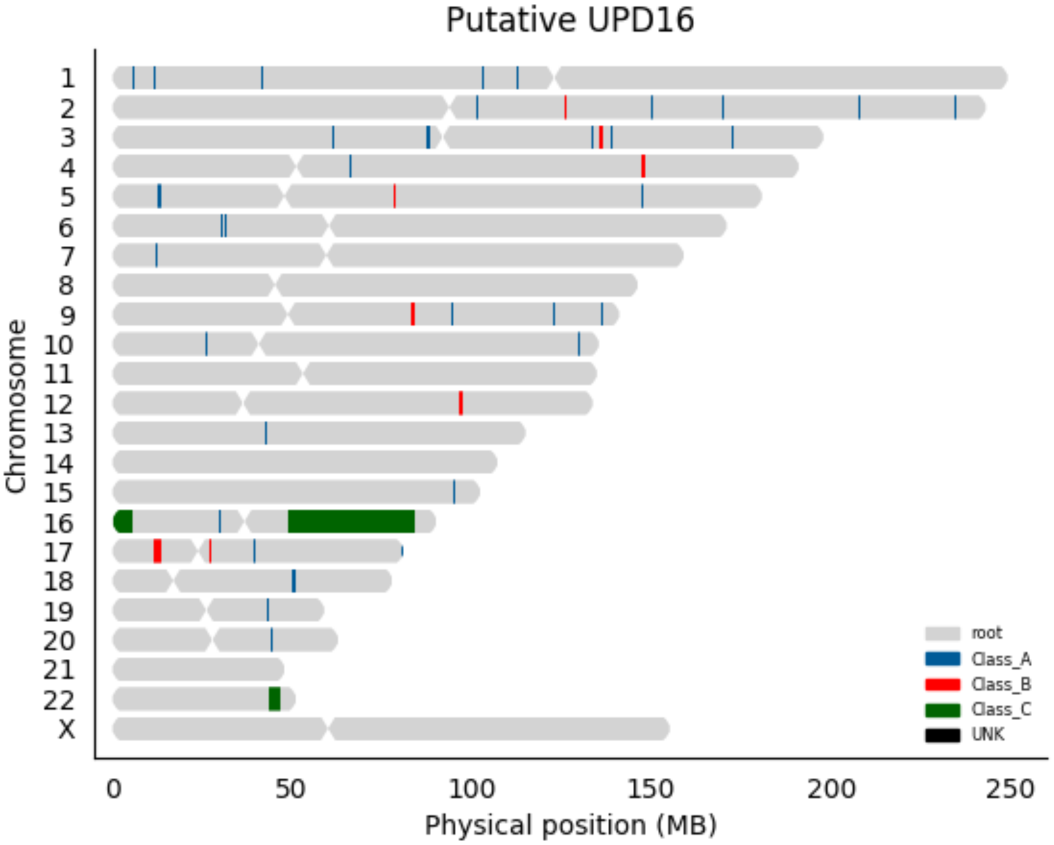

Putative UPD16
Chromosome
X
0
50
100
150
200
250
Physical position (MB)
root
Class_A
Class_B
Class_C
UNK

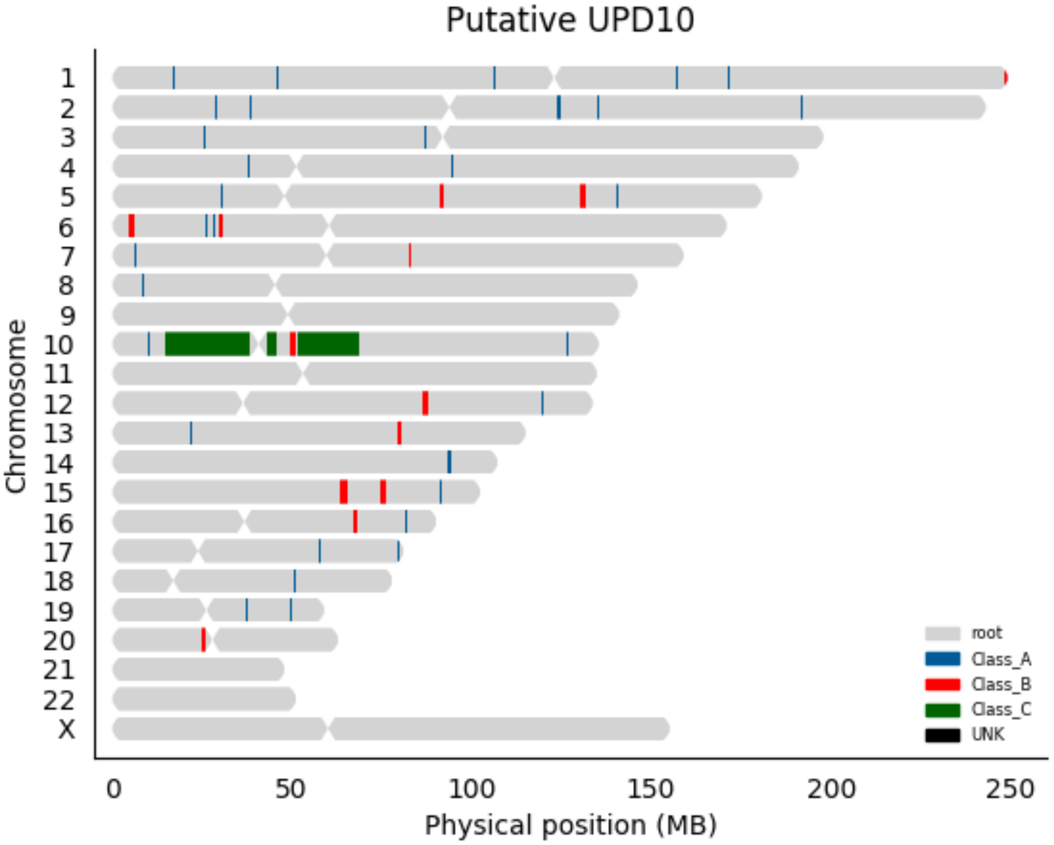

Putative UPD10
Chromosome
X
0
50
100
150
200
250
Physical position (MB)
root
Class_A
Class_B
Class_C
UNK

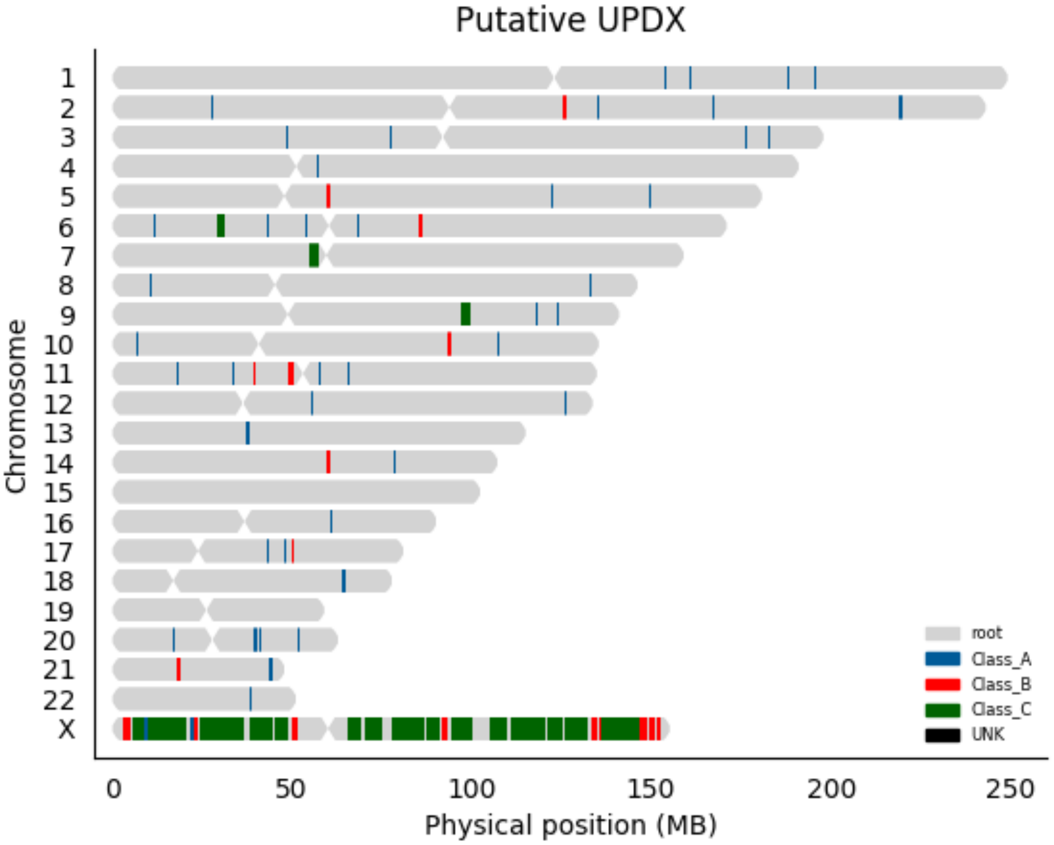
Putative UPDX
Chromosome
X
root
Class_A
Class_B
Class_C
UNK
0
50
100
150
200
250
Physical position (MB)

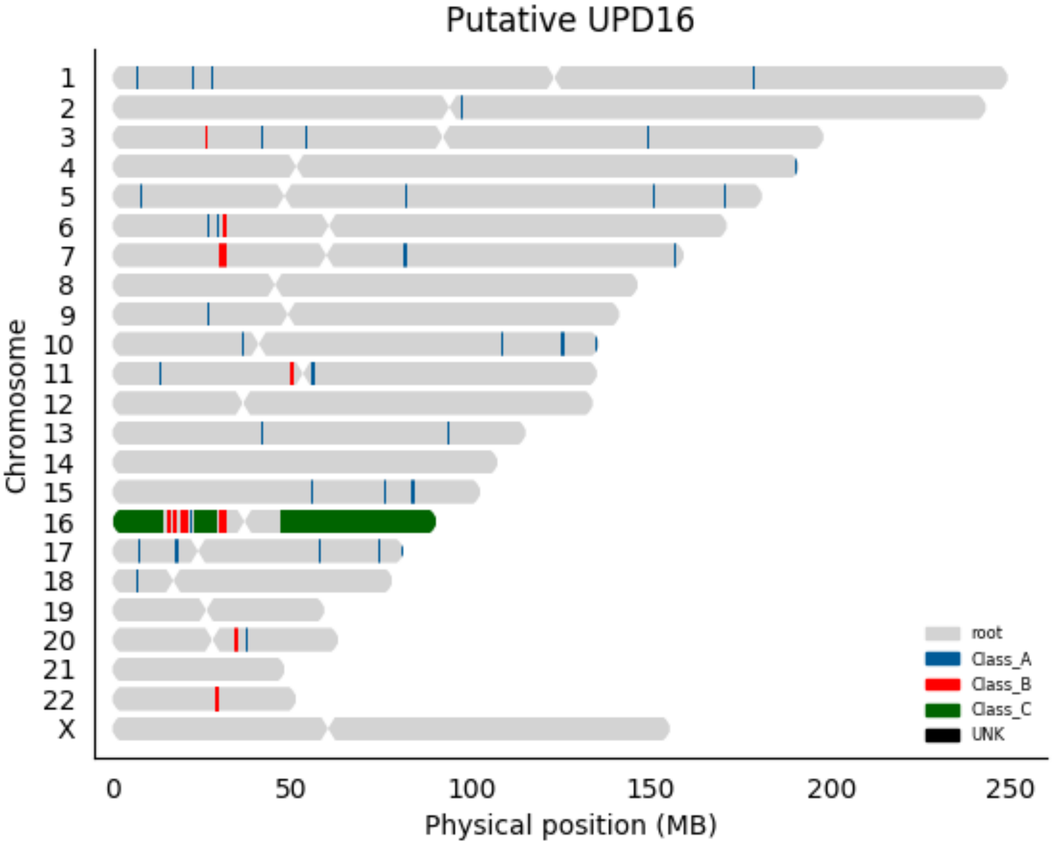

Putative UPD16
Chromosome
X
0
50
100
150
200
250
Physical position (MB)
root
Class_A
Class_B
Class_C
UNK

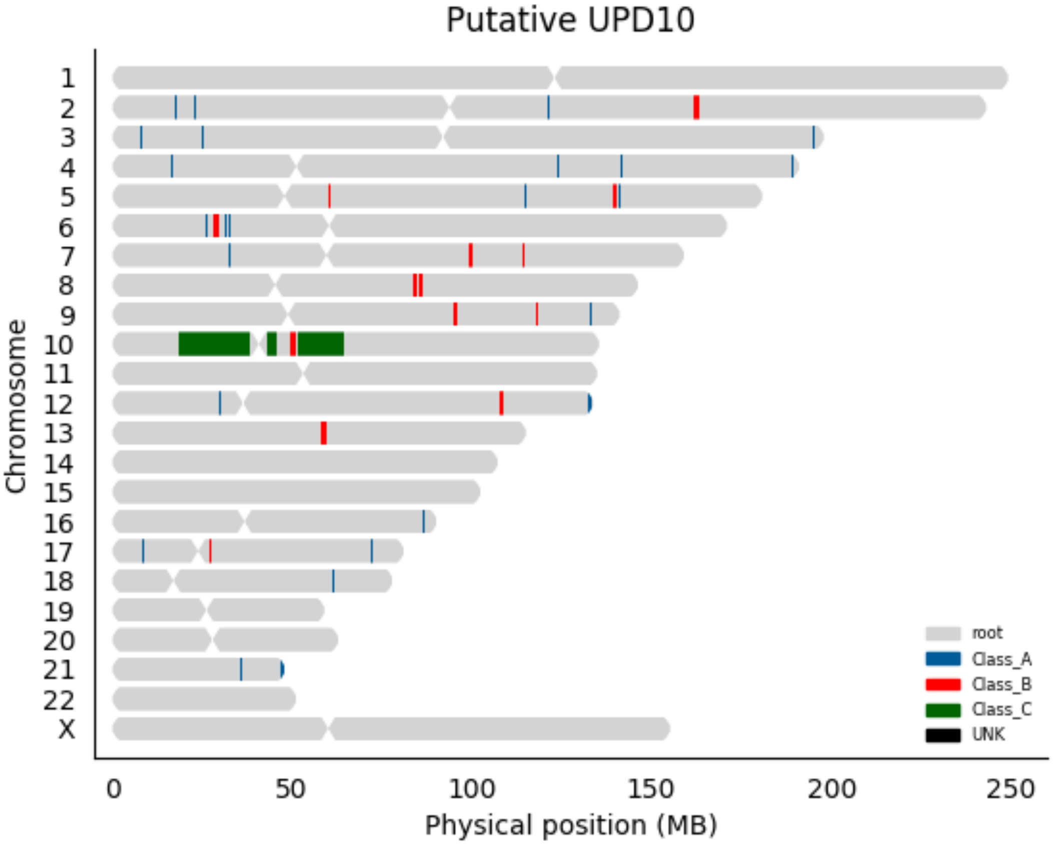
Putative UPD10
Chromosome
X
0
50
100
150
200
250
Physical position (MB)
root
Class_A
Class_B
Class_C
UNK

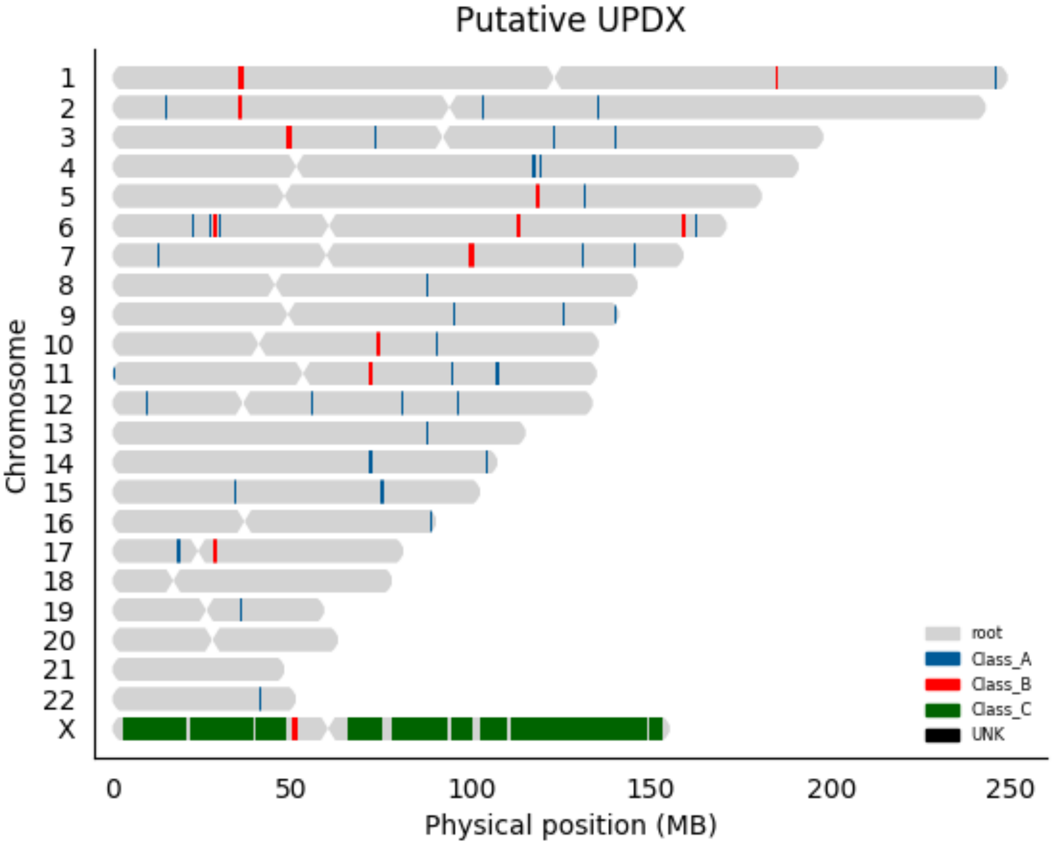

Putative UPDX
Chromosome
X
0
50
100
150
200
250
Physical position (MB)
root
Class_A
Class_B
Class_C
UNK

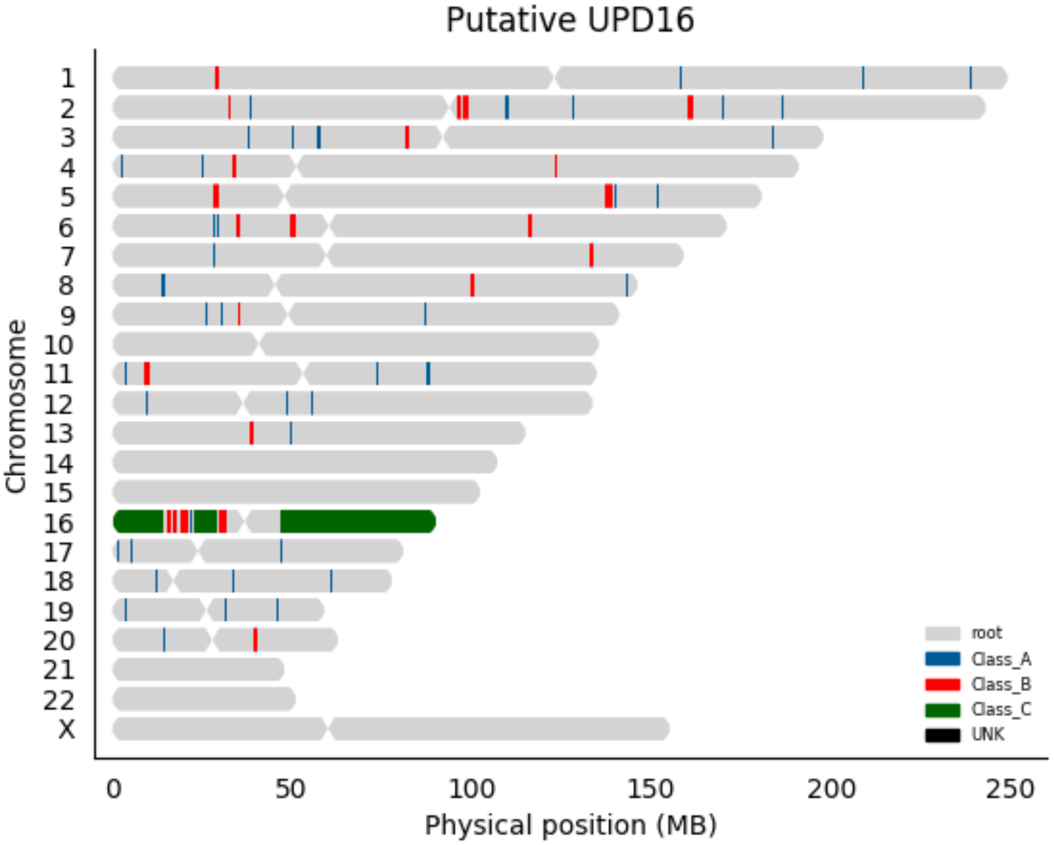

Putative UPD16
Chromosome
X
0
50
100
150
200
250
Physical position (MB)
root
Class_A
Class_B
Class_C
UNK

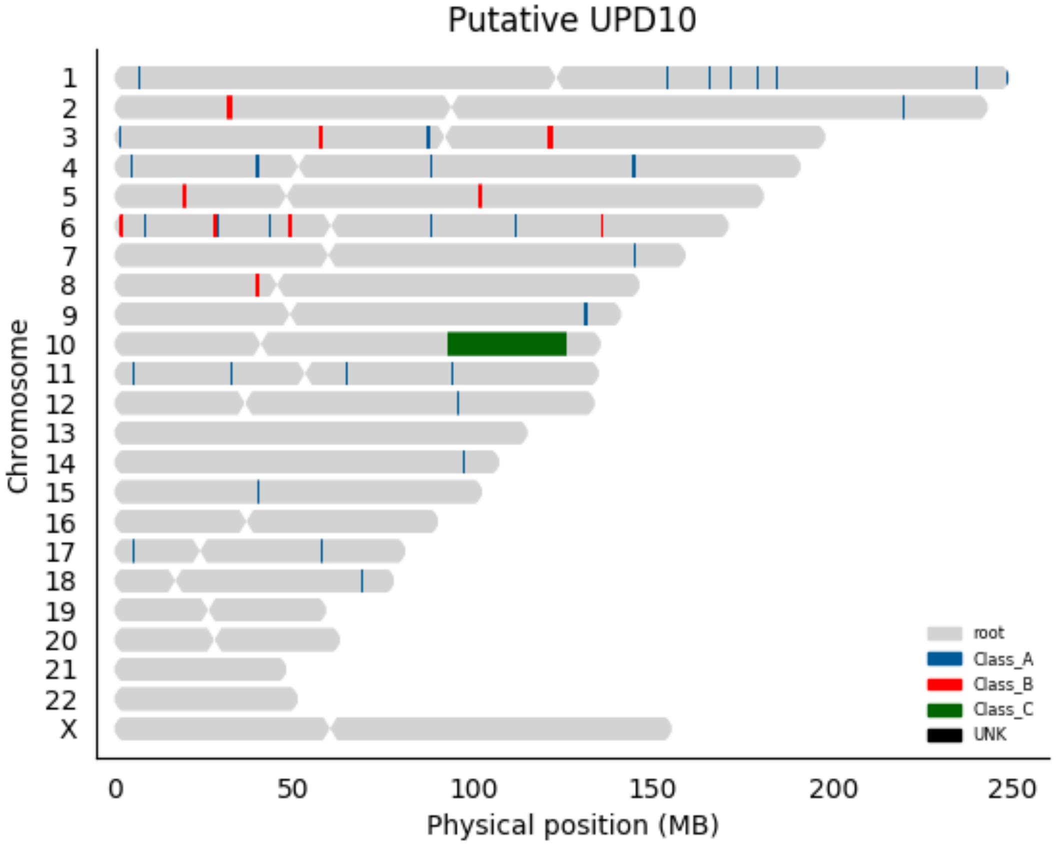

Putative UPD10
X
Chromosome
0
50
100
150
200
250
Physical position (MB)
root
Class_A
Class_B
Class_C
UNK

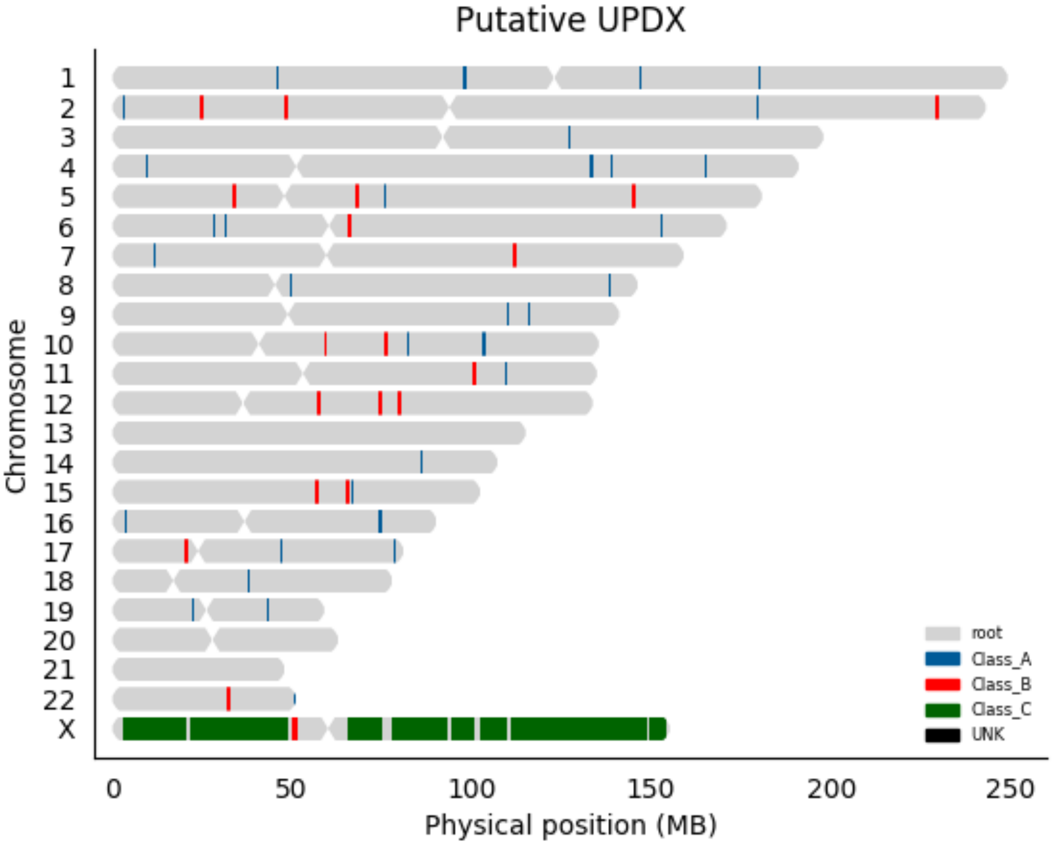
Putative UPDX
Chromosome
X
0
50
100
150
200
250
Physical position (MB)
root
Class_A
Class_B
Class_C
UNK

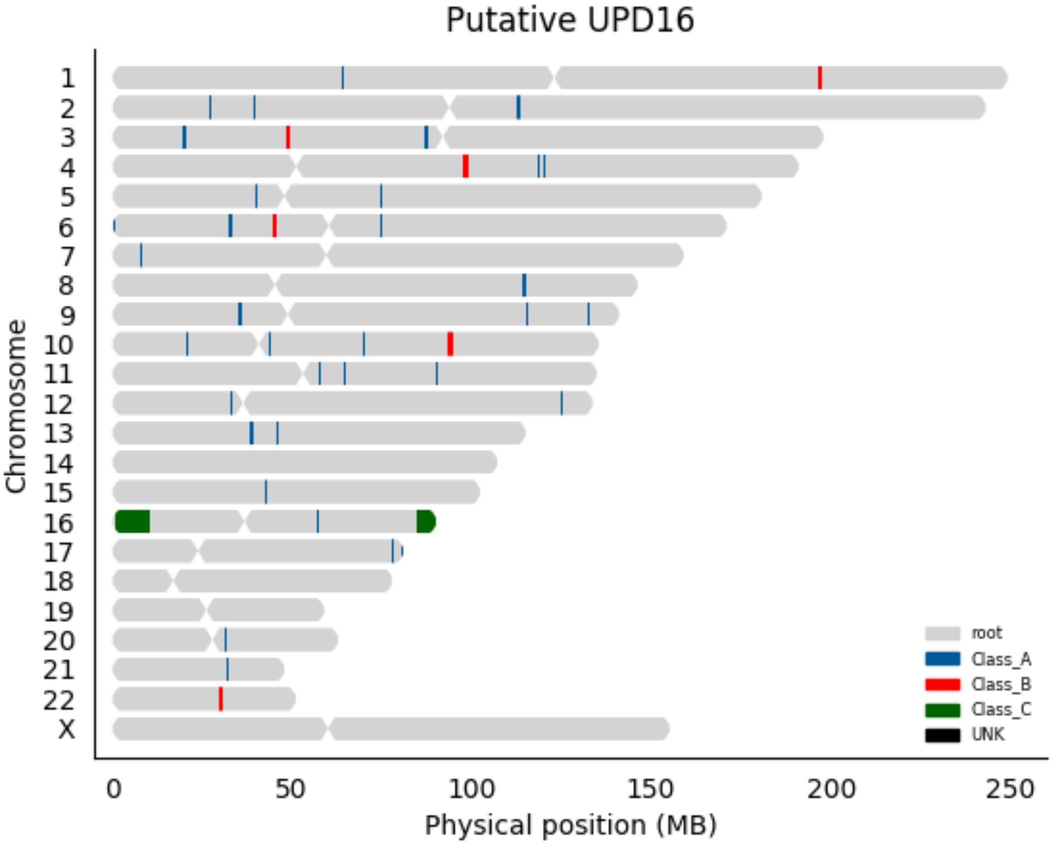

Putative UPD16
Chromosome
X
0
50
100
150
200
250
Physical position (MB)
root
Class_A
Class_B
Class_C
UNK

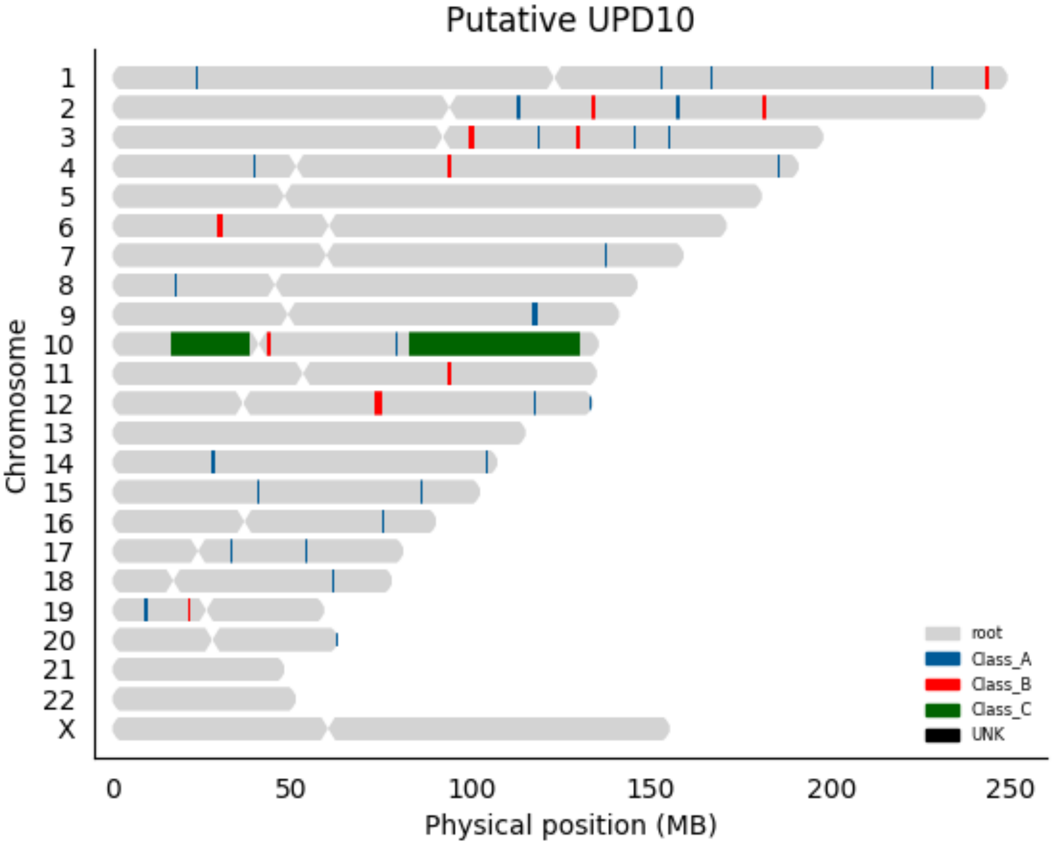

Putative UPD10
Chromosome
X
0
50
100
150
200
250
Physical position (MB)
root
Class_A
Class_B
Class_C
UNK
